# Evidence for the major role of PH4αEFB in the prolyl 4-hydroxylation of *Drosophila* collagen IV

**DOI:** 10.1101/2025.08.05.668786

**Authors:** Yoshihiro Ishikawa, Melissa A. Toups, Marwan Elkrewi, Allison L. Zajac, Sally Horne-Badovinac, Yutaka Matsubayashi

## Abstract

Collagens are fundamental components of extracellular matrices, requiring precise intracellular post-translational modifications for proper function. Among the modifications, prolyl 4-hydroxylation is critical to stabilise the collagen triple helix. In humans, this reaction is mediated by collagen prolyl 4-hydroxylases (P4Hs). While humans possess three genes encoding these enzymes (P4H⍺s), *Drosophila melanogaster* harbour at least 26 candidates for collagen P4H⍺s despite its simple genome, and it is poorly understood which of them are actually working on collagen in the fly. In this study, we addressed this question by carrying out thorough bioinformatic and biochemical analyses. We demonstrate that among the 26 potential collagen P4H⍺s, PH4⍺EFB shares the highest homology with vertebrate collagen P4H⍺s. Furthermore, while collagen P4Hs and their substrates must exist in the same cells, our transcriptomic analyses at the tissue and single cell levels showed a global co-expression of *PH4*⍺*EFB* but not the other P4H⍺-related genes with the collagen IV genes. Moreover, expression of *PH4*⍺*EFB* during embryogenesis was found to precede that of collagen IV, presumably enabling efficient collagen modification by PH4⍺EFB. Finally, biochemical assays confirm that PH4⍺EFB binds collagen, supporting its direct role in collagen IV modification. Collectively, we identify PH4⍺EFB as the primary and potentially constitutive prolyl 4-hydroxylase responsible for collagen IV biosynthesis in *Drosophila*. Our findings highlight the remarkably simple nature of *Drosophila* collagen IV biosynthesis, which may serve as a blueprint for defining the minimal requirements for collagen engineering.

## Introduction

Collagens are one of the most abundant protein superfamilies and composed of 28 different types of collagen proteins in humans [1, 2]. They play essential functions such as imparting biophysical properties to tissues and acting as signalling scaffolds [3–5]. For example, collagens I, II, III, V, and XI are classified as fibrillar collagens, forming structural frameworks in fibril-rich connective tissues, including those in bone, skin, and the vasculature. In contrast, collagens IV, XVII and XVIII and others form flexible networks present in the basement membranes that support all epithelia and many other tissues such as muscles. For a single type of collagen, multiple isoforms can exist. For instance, there are six collagen IV genes in mammals (*COL4A1* – *COL4A6*), encoding six collagen IV proteins known as ⍺ chains (collagen IV α1 – α6). These six α chains assemble into three heterotrimeric collagen IV isoforms, collagens α1α1α2(IV), α3α4α5(IV) and α5α5α6(IV) [3, 6].

Collagen biosynthesis is a highly orchestrated process and requires more than 20 enzymes and chaperones (hereafter referred to as the ‘collagen molecular ensemble’) residing in the endoplasmic reticulum (ER) [7–11]. Interestingly, similar to the diversity of collagen types and isoforms, many components of this ensemble, particularly collagen modifying enzymes, also exhibit remarkable diversity: for example, in humans there are three prolyl 4-hydroxylases (P4Hs), three prolyl 3-hydroxylases (P3Hs), three lysyl hydroxylases (LHs), and two glycosyl transferase 25 domain enzymes (GLT25Ds) [12–14]. Therefore, collagen post-translational modifications (PTMs) can serve as signatures that distinguish collagen types [8, 9, 15, 16]. This diversity of collagen types, collagen modifying enzymes, and PTM patterns suggest that collagen biosynthesis must facilitate a complex set of parameters essential for the correct combinations of α-chain isoforms and PTMs across the 28 different collagen types. Therefore, understanding collagen functions and designing therapies against collagen-related diseases requires research not only on collagens themselves but also on the components of the collagen molecular ensemble [17, 18].

The fruit fly *Drosophila melanogaster* is a well-established model organism for several reasons: 1) it has a relatively compact genome yet shares a significant portion of its genes with humans, including those associated with diseases, 2) its genome is well-mapped and extensively annotated, providing a wealth of genetic tools and resources, including numerous mutant strains and transgenic lines, and 3) it boasts a strong and supportive research community with extensive collaboration and resources sharing, facilitated by accessible database and stock centres [19]. This toolset would also be useful to boost research on collagen biosynthesis, considering the reasons described below. First, *Drosophila* possesses only a minimum set of collagens, lacking any fibrillar collagens [20]. Second, its predominant collagen is the basement membrane collagen α1α1α2(IV), with Multiplexin (a homologue of human collagen XV/XVIII) and Pericardin (a collagen IV-like protein) being other collagen(-related) molecules [21, 22]. While mammals have six collagen IV α chain genes, *Drosophila* has only two collagen IV α1 and α2, encoded by *Col4a1* and *viking* (*vkg)* genes, respectively [23, 24]. Lastly, the collagen molecular ensemble for *Drosophila* collagen biosynthesis is also simple. For example, *Drosophila* has only one isoform each of LH and GLT25D (FlyBase IDs FBgn0036147 and FBgn0051915), compared to three and two in humans, respectively. Additionally, a critical collagen molecular chaperone HSP47 has not been identified in *Drosophila*, and it lacks a gene encoding P3H. This suggests that *Drosophila* represents a minimal system for collagen biosynthesis, making it an ideal model to study the core mechanisms of this essential process.

Prolyl 4-hydroxylases (P4Hs) are enzymes that hydroxylate proline residues to 4-hydroxylproine (4Hyp). In humans, there are seven P4Hs, three of which work on collagens while the others target different substrates including the transcription factors known as hypoxia-inducible factors (HIFs) [25–28]. Collagen P4Hs exist in three isoforms, each of which is a tetrameric complex composed of two ⍺ and two β subunits. The catalytic ⍺ subunits (P4H⍺s) are specific to each isoform, and encoded by the genes P4HA1, P4HA2, and P4HA3, respectively. The β subunit, which is identical to the enzyme protein disulfide isomerase (PDI), is shared by all three isoforms [29–31]. The formation of 4Hyp by this tetramer confers thermal stability to collagens: without 4Hyp, collagens are unable to form a stable triple helical structure at body temperature [32–35]. Surprisingly, in contrast to the aforementioned cases with collagens, LHs, and GLT25Ds, *Drosophila* possesses a far larger number of genes potentially encoding collagen P4H⍺s than mammals do: 26 genes are currently annotated with a Gene Ontology term suggesting collagen P4H activity, mainly based on their sequences (see Results). This raises the question: how many P4Hs are involved in *Drosophila* collagen α1α1α2(IV) biosynthesis? One possibility is that the large number of P4H enzymes have evolved to modify collagens in a manner that is specific to *Drosophila,* making them poor models for the action of the human collagen P4Hs. Alternatively, only a small number of the P4Hs modify collagens and the others have different substrates, which might make the *Drosophila* P4Hs good models for the human enzymes.

This question has been partially addressed in previous studies. In 2002, Abrams and Andrew reported that their BLAST search identified nineteen P4H⍺-related genes in the *Drosophila* genome. They analysed the expression patterns of ten of them in the embryo and found that only PH4⍺EFB was expressed in hemocytes and the fat body [36], the major collagen IV-producing tissues [37]. Since prolyl 4-hydroxylation of collagen occurs intracellularly [32, 38], expression in these tissues suggests PH4⍺EFB is a likely collagen-modifying P4H⍺. Later, genetic evidence supported this: RNAi knockdown of PH4⍺EFB disrupted collagen IV secretion [39, 40]. In contrast, biochemical studies showed that a recombinant form of another *Drosophila* PH4⍺ that was later named PH4⍺MP [36] can hydroxylate collagen peptides *in vitro* [41]. Although both PH4⍺EFB and MP are strong candidates for collagen modifying P4H⍺s, it remains unclear whether one, both, or additional P4H⍺s contribute to collagen biosynthesis directly in *Drosophila*, as their roles are still not fully defined.

In this study, we carried out bioinformatic and biochemical analyses to identify the *Drosophila* P4H⍺s responsible for collagen modification. Our results suggest that PH4⍺EFB is a central collagen modifying P4H⍺ among the 26 potential candidates in *Drosophila,* highlighting the simplicity of the collagen molecular ensemble in this model organism.

## Results

### The Drosophila genome encodes 26 P4H***α***-related genes

To comprehensively identify fly collagen P4H⍺-related genes, we searched FlyBase for the genes annotated with the Gene Ontology (Molecular Function) term ‘procollagen-proline 4-dioxygenase activity’, which is associated with human P4HA1 (Prolyl 4-hydroxylase subunit ⍺-1, NCBI Gene ID 5033). This analysis revealed 26 genes (Table 1), including all the 10 that have been analysed previously [36]. At the time of the search, no experimental data had been recorded as the evidence of the GO term annotation, which was instead based solely on the sequence of the genes (the reports about PH4⍺MP and EFB [34–36, 41] had not been curated yet). Moreover, the majority of these genes had not been named and are still referred to by ‘CG’ symbols. These facts indicate that the genes have been only minimally characterised.

**Table 1.** *Drosophila* collagen P4H⍺-related genes. Genes annotated with the GO (Molecular Function) term ‘procollagen-proline 4-dioxygenase activity’ were searched for on FlyBase and listed. For their protein products, the isoforms used for the phylogenetic analyses in this study are shown with their SwissProt IDs. Bold letters indicate the genes that have been analysed previously [36]. For all the genes, the GO term was recorded to have been inferred from ‘electronic annotation with InterPro:IPR013547’ and ‘biological aspect of ancestor with PANTHER:PTN004202971’.

| Gene Symbol | FlyBase ID | Analysed isoform(s) | SwissProt ID |
| --- | --- | --- | --- |
| CG11828 | FBgn0039616 | PD | Q9VAR7 |
| **CG15539** | FBgn0039782 | PA | Q4V443 |
|  |  | PB | Q0KHY7 |
|  |  | PC | A0A0B4LHW6 |
| CG15864 | FBgn0040528 | PB | Q9VHU7 |
| CG18231 | FBgn0036796 | PB | Q9VVQ7 |
| CG18233 | FBgn0036795 | PB | Q9VVQ6 |
| CG18234 | FBgn0265268 | PC | Q9VVQ5 |
| CG18749 | FBgn0042182 | PB | Q8MSK0 |
| CG31013 | FBgn0051013 | PA | Q9VA50 |
| CG31016 | FBgn0051016 | PB | Q9VA52 |
| CG31021 | FBgn0051021 | PB | Q8IMH8 |
| CG31371 | FBgn0051371 | PB | Q8IMI4 |
| CG31524 | FBgn0051524 | PA | Q8IMI2 |
| CG32199 | FBgn0052199 | PB | Q9VVQ9 |
| CG32201 | FBgn0052201 | PB | Q8IQS7 |
| CG34041 | FBgn0054041 | PD | D3DMS2 |
|  |  | PE | A0A0B4KHI9 |
|  |  | PF | Q2PDP5 |
| CG34345 | FBgn0085374 | PA | A8DYR4 |
| CG4174 | FBgn0036793 | PD | Q9VVQ4 |
| **CG9698** | FBgn0039784 | PA | Q9VA60 |
| **PH4αEFB** | FBgn0039776 | PA | Q9VA69 |
| **PH4αMP** | FBgn0026190 | PA | Q9VA65 |
| **PH4αNE1** | FBgn0039780 | PA | Q9VA64 |
| **PH4αNE2** | FBgn0039783 | PA | Q9VA61 |
| **PH4αNE3** | FBgn0051017 | PA | Q961I8 |
| **PH4αPV** | FBgn0051015 | PA | Q8T5S8 |
| **PH4αSG1** | FBgn0051014 | PA | Q9VA63 |
| **PH4αSG2** | FBgn0039779 | PA | Q9I7H5 |

### Domain structure is conserved between human collagen P4H***α*** and 25 Drosophila homologues

Each human collagen P4H⍺ protein has four parts: the N-terminal (N) domain, the peptide-substrate-binding (PSB) domain, a linker (L) region, and the C-terminal catalytic (CAT) domain (Figure 1A). This structure enables a P4H⍺ protein to bind to another P4H⍺ via the N-domain and to the β subunit PDI via the CAT domain, forming the ⍺2β2 tetramer[30, 31, 42, 43]. The P4HA-PDI interaction is essential for maintaining the solubility and ER retention of P4HA required for its enzymatic activity [44, 45]. The formation of this tetramer has been proposed to be evolutionarily conserved in a previously identified *Drosophila* collagen P4H⍺-related protein PH4⍺MP [29, 36, 41]. Thus, we examined whether the ‘N-PSB-L-CAT’ structure that enables collagen P4H formation is present in the proteins encoded by the 26 fly collagen P4H⍺-related genes identified in this study. We compared the sequences of the fly proteins with that of human collagen P4HA2; as a control, we also examined human PHD3, a prolyl hydroxylase whose substrate is the transcription factor hypoxia-inducible factor (HIF) rather than collagens. It is known that the structure outside the catalytic domain is unrelated between collagen and HIF P4Hs [26, 46] (Figure 1B). While the sequence alignment confirmed that the structures of P4HA2 and PHD3 are conserved only within the catalytic domains (Table S1), we found that almost all the *Drosophila* collagen P4H⍺-related proteins have the same domain organisation as human collagen P4HAs except for the following two cases. First, about 70% of the N-terminal side of the N domain of CG15539-PA is truncated, while the other protein isoforms encoded by the same gene (CG15539-PB and PC) contain a full-length N-domain (https://flybase.org/reports/FBgn0039782) (Figure 1C, Table S1). Second, all the three products of *CG34041* (PD, PE, and PF) lack the catalytic domain; FlyBase shows that they contain one or two N-domains (https://flybase.org/reports/FBgn0054041) (Figure 1 D and E, Table S1). Thus, *CG34041* does not encode a prolyl hydroxylase. However, for the other 25 fly collagen P4H⍺-related genes, the ‘N-PSB-L-CAT’ domain organisation is conserved between their products and human collagen P4HAs, suggesting the ability of the fly proteins to form tetramers with PDI. This necessitates further investigation to identify which of the fly enzymes work on collagen IV.

**Figure. 1.**
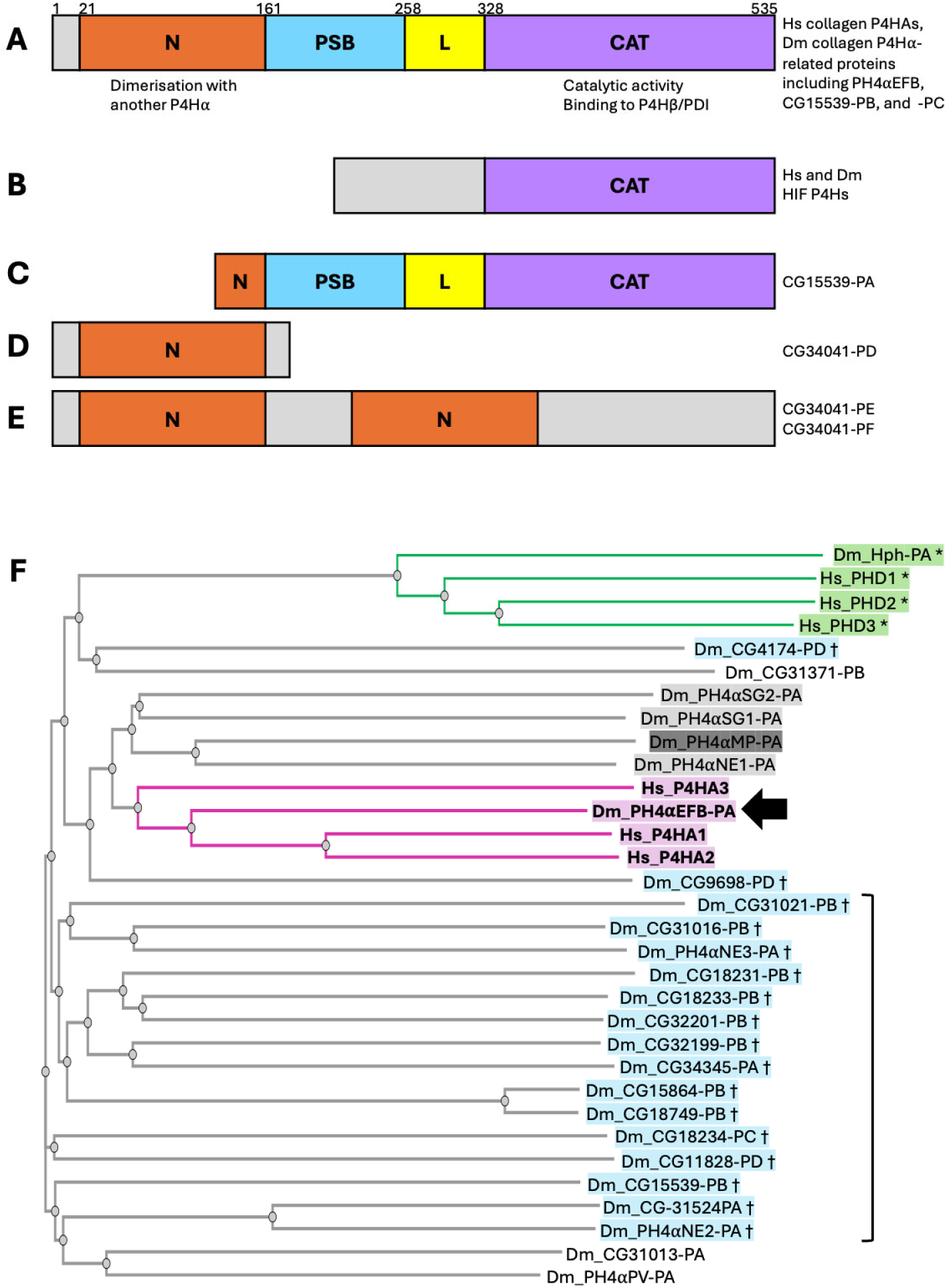
Domain organisation and phylogeny of human (Hs) and *Drosophila* (Dm) prolyl 4-hydroxylases. **(A–E)** Schematic representations of the protein domain structures analysed. Diagrams are not drawn to scale. For full details, see Table S1. **(A)** Canonical domain structure of collagen P4H⍺ proteins, comprising N-terminal (N), peptide-substrate-binding (PSB), linker (L), and catalytic (CAT) domains. This organisation is conserved in all three human collagen P4HAs and in proteins encoded by 25 of the 26 *Drosophila* P4H⍺-related genes including those mentioned at the right. Numbers above the schematic indicate domain boundary positions for human P4HA2. **(B)** Domain structure of HIF prolyl 4-hydroxylases (PHD1–3 and Hph). These are homologous to collagen P4HAs only within the catalytic domain. **(C)** The CG15539-PA isoform has a truncated N-domain, while the other two isoforms (PB and PC) exhibit the full domain organization as in **(A)**. **(D, E)** All three annotated isoforms of CG34041 (PD, PE, and PF) lack the catalytic domain. CG34041-PD contains an N-domain, whereas CG34041-PE and -PF contain two tandem N-domains. **(F)** Phylogenetic tree of P4H⍺s. At the top of the tree, *Drosophila* Hph and human PHD1–3 (green, marked with asterisks [*]) form a distinct clade. In the middle, *Drosophila* PH4⍺EFB (arrow) clusters with the three human collagen prolyl 4-hydroxylases (P4HA1–3), forming a separate clade (magenta, highlighted in bold). PH4⍺MP (dark grey shading) and neighbouring *Drosophila* enzymes (light grey shading) are implicated in tissue-or context-specific collagen modification (see Discussion). Proteins shaded in blue and marked with daggers (†) are highly expressed in the male accessory gland and may hydroxylate the seminal fluid protein Sex Peptide (SP). Sixteen of these 18 accessory gland P4H⍺-related proteins cluster in the most phylogenetically distant region from the magenta and grey clades (bracket; see also Discussion).

### Comprehensive phylogenetic analysis confirms that PH4⍺EFB is closest to human P4Has

Subsequently, we examined the sequence similarity between human and *Drosophila* collagen P4H⍺-related proteins. Since phylogenetic analysis revealed that PH4⍺EFB was closest to human collagen P4H1A1/2 among the 8 P4H⍺-related proteins previously tested [36], we conducted a more comprehensive analysis to determine if unexamined fly proteins show a similar or higher homology to human collagen P4HAs. We constructed a phylogenetic tree using the sequences of human collagen P4HA1, 2, and 3, and the proteins encoded by 25 out of the 26 *Drosophila* collagen P4H⍺-related genes: *CG34041* was excluded from the assay as it was found not to encode a catalytic domain. As a control, we also examined three human and one *Drosophila* HIF prolyl hydroxylases, PHD1-3 [26] and Hph [47], respectively. This analysis first revealed that the HIF prolyl hydroxylases—human PHD1-3 and *Drosophila* Hph—formed a distinct clade (Figure 1F top, green and marked with asterisks), suggesting a correspondence between sequence similarity and substrate specificity. Building on this, the three human collagen P4HA proteins and *Drosophila* PH4⍺EFB formed another clade comprising only these proteins, with the fly enzyme occupying a central position (Figure 1F, magenta and highlighted in bold; arrow points PH4⍺EFB-PA, the only annotated product of the *PH4*⍺*EFB* gene). These results strengthen the previous notion that PH4⍺EFB is closest to human collagen P4HAs among all the fly collagen P4H⍺-related proteins [36].

### PH4***α***EFB shows a spatiotemporal expression pattern compatible with it being the major P4H***α*** for Drosophila collagen IV

As the proline 4-hydroxylation of collagens occurs intracellularly [32, 38], an enzyme mediating this reaction must be co-expressed in the same cell as its substrate collagens. As described in the phylogenetic analysis above, Abrams and Andrew also analysed the expression patterns of the 10 P4H⍺-related genes to identify which enzymes are responsible for the proline 4-hydroxylation in the fly embryo using mRNA in situ hybridisation [36]. Their analysis revealed that *PH4*⍺*EFB* but not the other nine genes is expressed in hemocytes (macrophages) and the fat body, which are the major sources of collagen IV in the embryo [24, 37]. Here, we validate whether PH4⍺EFB remains the sole *Drosophila* PH4⍺ targeting collagen IV as previously suggested [36], utilising newly available materials.

First, we examined the co-expression of all 26 collagen P4H⍺-related genes (Table 1) and the two collagen IV genes, *Col4a1* and *vkg*, in many different postembryonic tissues, cultured cells, and individuals under various environmental perturbations utilising five publicly available transcriptome datasets: FlyAtlas Anatomy Microarray, FlyAtlas2 Anatomy RNA-Seq, and modENCODE Anatomy RNA-Seq for the data from the tissues, modENCODE Cell Lines RNA-Seq for the cultured cells, and modENCODE Treatments RNA-Seq for the environmental perturbations [48–50] (Figure 2, Table 2, Table S2). To quantify the similarity of the expression patterns of *Col4a1* and each of the other genes, we measured the correlation coefficient (*r*), which takes a value between -1 and 1. The value of *r* = 1, 0, and -1 indicate identical, uncorrelated, and complementary expression patterns to *Col4a1*, respectively. In the first four datasets, *vkg* showed a nearly identical expression pattern to *Col4a1* (*r* > 0.95); in the modENCODE Treatments dataset, the correlation was slightly lower but still high (*r* = 0.79). These high correlations presumably reflect the evolutionarily conserved co-expression of the two collagen IV subunit genes under the control of a common promoter [51], supporting the reliability of our assay. Regarding the collagen P4H⍺-related genes, *PH4*⍺*EFB* showed the largest and outstanding *r* values in three datasets (FlyAtlas, FlyAtlas2, and modENCODE Cell Lines) (Figures 2A and B). Indeed, the expression levels of *Col4a1* and *PH4*⍺*EFB* often change in parallel between samples in these datasets (e.g., thick solid bracket in Figure 2C, FlyAtlas2). Even in the remaining two datasets (modENCODE Anatomy and Treatments), *Col4a1* and *PH4*⍺*EFB* levels change almost in parallel, although the graphs do not necessarily exhibit identical numbers of peaks and troughs (e.g., thin solid brackets in Figure 2C). In addition, apart from only two exceptions (Figure 2C>, closed arrows, both from the larval salivary gland) out of a total of 145 samples in all the five datasets, *PH4*⍺*EFB* expression was visible in the samples expressing *Col4a1*.

**Figure. 2.**
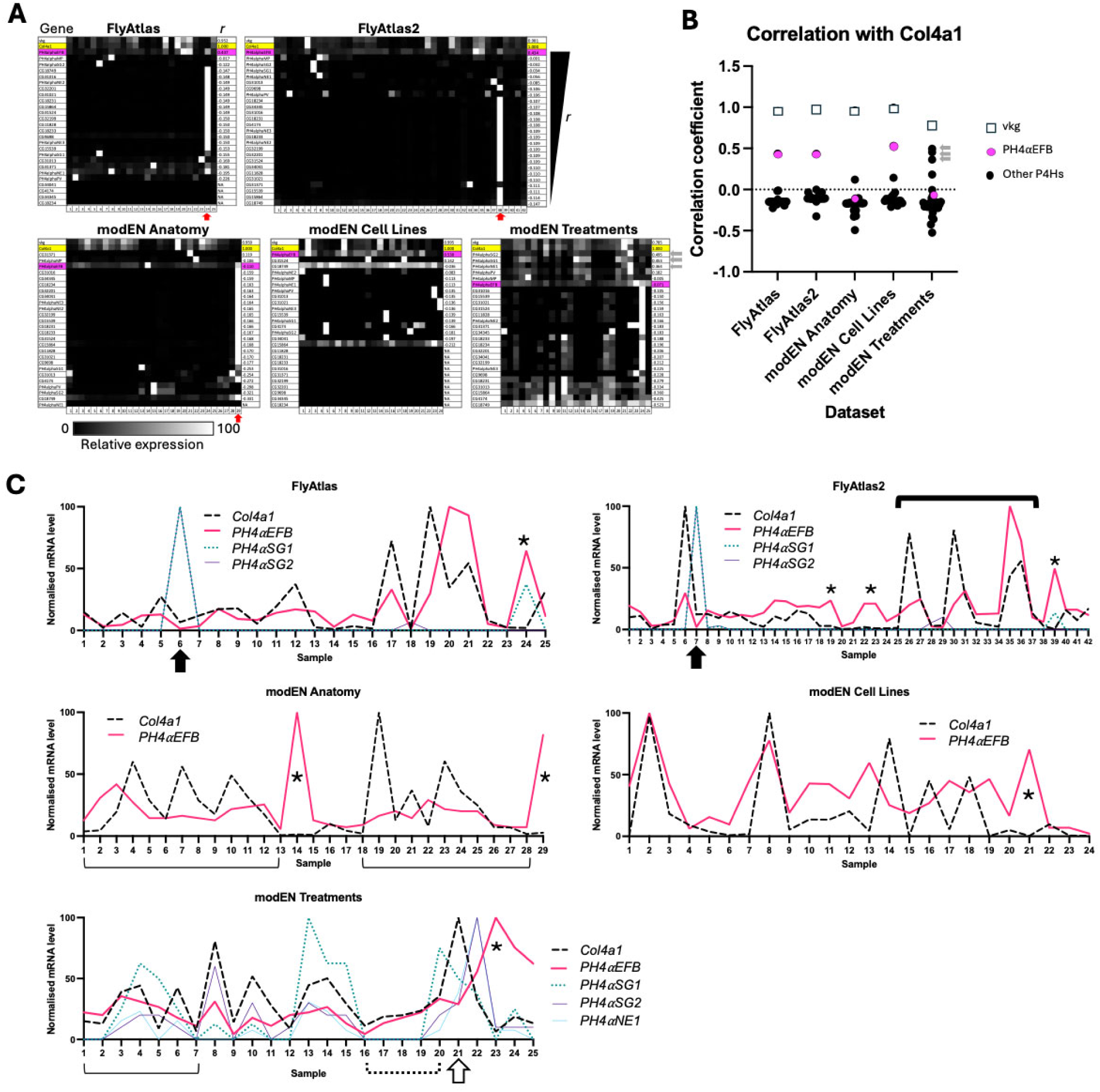
Spatial expression patterns of *Drosophila* collagen IV and PH4α-related genes. **(A)** The expression levels of each gene from the transcriptome dataset indicated are normalised for the maximum level and brightness-coded as in the bar at the bottom. modEN, modENCODE. In all panels, *vkg* is shown at the top. Yellow, *Col4a1*; magenta, *PH4*⍺*EFB*. The PH4⍺-related genes are sorted in the descending order of the values of the correlation coefficient *r* between the expression patterns of *Col4a1* and each of the other genes. For the legends for the sample numbers in **(A)** and **(C)**, see Table 2. For enlarged gene names and the values of gene expression levels, see Table S2, where the genes are sorted in the same order. Grey arrows beside the bottom right panel point the data with *PH4*⍺*SG2*, *SG1*, and *NE1.* Red arrows point accessory gland samples in which multiple PH4⍺-related genes are highly expressed. **(B)** Plot of the *r* values for the genes in each dataset in **(A)**. Grey arrows point the data with *PH4*⍺*SG2*, *SG1*, and *NE1* from top to bottom, respectively. **(C)** For *Col4a1*, *PH4*⍺*EFB, SG1, SG2,* and *NE1*, the values in **(A)** are displayed as line scattered plots. Thick solid bracket, examples of the parallel peaks and troughs of the expression levels of *Col4a1* and *PH4*⍺*EFB*. Thin solid brackets, examples of the cases where *Col4a1* and *PH4*⍺*EFB* levels change largely in parallel, although the graphs do not exhibit identical numbers of peaks and troughs. Closed arrows, samples in which the expression levels of *PH4*⍺*EFB* is low while *Col4a1* is detected. Asterisks, samples in which *PH4*⍺*EFB* level is high while *Col4a1* level is low. Dotted bracket, parallel increase of *Col4a1* and *PH4*⍺*EFB* expression levels. Open arrow, between samples 20 and 21, while the expression level of *Col4a1* increases steeply, that of *PH4*⍺*EFB* slightly decreases. For detail, see Discussion.

**Table 2.** Legend for the sample numbers in the horizontal axes of each panel in Fig. 2A and C.

| FlyAtlas | FlyAtlas2 | modENCODE Anatomy | modENCODE Cell Lines | modENCODE Treatments |
| --- | --- | --- | --- | --- |
| 1. Larval Central Nervous System | 1. 3rd Instar Larval CNS | 1. Imaginal Disc, 3rd Instar Larvae Wandering | 1. Schneider Line 2 S2R+ | 1. Extended Cold, 4-Day Adult |
| 2. Larval Midgut | 2. 3rd Instar Larval Trachea | 2. Central Nervous System, 3rd Instar Larvae | 2. Schneider Line 2 Sg4 | 2. Cold Shock, 4-Day Adult |
| 3. Larval Hindgut | 3. 3rd Instar Larval Midgut | 3. Central Nervous System, Pupae P8 | 3. Embryonic 1182-4H | 3. Heat Shock, 4-Day Adult |
| 4. Larval Malpighian Tubules | 4. 3rd Instar Larval Hindgut | 4. Head, Virgin 1-Day Female | 4. Embryonic GM2 | 4. Cadmium 50 mM 6 Hrs, Larvae L3 |
| 5. Larval Fat Body | 5. 3rd Instar Larval Malpighian Tubule | 5. Head, Virgin 4-Day Female | 5. Embryonic Kc167 | 5. Cadmium 50 mM 12 Hrs, Larvae L3 |
| 6. Larval Salivary Gland | 6. 3rd Instar Larval Fat Body | 6. Head, Virgin 20-Day Female | 6. Embryonic S1 | 6. Cadmium 50 mM 48 Hrs, 4-Day Adult |
| 7. Larval Trachea | 7. 3rd Instar Larval Salivary Gland | 7. Head, Mated 1-Day Female | 7. Embryonic S3 | 7. Cadmium 100 mM 48 Hrs, 4-Day Adult |
| 8. Larval Carcass | 8. 3rd Instar Larval Carcass | 8. Head, Mated 4-Day Female | 8. Leg Disc CME L1 | 8. Copper 0.5 mM 12 Hrs, Larvae L3 |
| 9. Adult Head | 9. 3rd Instar Larval Whole | 9. Head, Mated 20-Day Female | 9. Wing Disc CME-W2 | 9. Copper 15 mM 48 Hrs, 4-Day Adult |
| 10. Adult Eye | 10. Adult Female Head | 10. Head, Mated 1-Day Male | 10. Wing Disc ML-Dmd8 | 10. Zinc 5 mM 12 Hrs, Larvae L3 |
| 11. Adult Brain | 11. Adult Male Head | 11. Head, Mated 4-Day Male | 11. Wing Disc ML-Dmd9 | 11. Zinc 4.5 mM 48 Hrs, 4-Day Adult |
| 12. Adult Thoracic-Abdominal Ganglion | 12. Adult Female Eye | 12. Head, Mated 20-Day Male | 12. Wing Disc ML-Dmd16-C3 | 12. Ethanol 2.5% 3 Hrs, Larvae L3 |
| 13. Adult Crop | 13. Adult Male Eye | 13. Salivary Gland, 3rd Instar Larvae Wandering | 13. Wing Disc ML-Dmd21 | 13. Ethanol 5% 3 Hrs, Larvae L3 |
| 14. Adult Midgut | 14. Adult Female Brain | 14. Salivary Gland, White Prepupae | 14. Wing Disc ML-Dmd32 | 14. Ethanol 10% 3 Hrs, Larvae L3 |
| 15. Adult Hindgut | 15. Adult Male Brain | 15. Digestive System, 3rd Instar Larvae Wandering | 15. Haltere Disc ML-Dmd17-C3 | 15. Caffeine 1.5 mg/ml 4 Hrs, Larvae L3 |
| 16. Adult Malpighian Tubules | 16. Adult Female Thoracico Abdominal Ganglion | 16. Digestive System, 1-Day Adult | 16. Eye-Antennal Disc ML-Dmd11 | 16. Caffeine 2.5 mg/ml 48 Hrs, 4-Day Adult |
| 17. Adult Fat Body | 17. Adult Male Thoracico Abdominal Ganglion | 17. Digestive System, 4-Day Adult | 17. Antennal Disc ML-Dmd20-C5 | 17. Caffeine 25 mg/ml 48 Hrs, 4-Day Adult |
| 18. Adult Salivary Gland | 18. Adult Female Crop | 18. Digestive System, 20-Day Adult | 18. Mixed Discs ML-Dmd4-C1 | 18. Paraquat 5 mM 48 Hrs, 4-Day Adult |
| 19. Adult Heart | 19. Adult Male Crop | 19. Fat Body, 3rd Instar Larvae Wandering | 19. CNS ML-Dmbg1-C1 | 19. Paraquat 10 mM 48 Hrs, 4-Day Adult |
| 20. Adult Virgin Female Spermatheca | 20. Adult Female Midgut | 20. Fat Body, White Prepupae | 20. CNS ML-Dmbg2-C2 | 20. Rotenone 2 µg 12 Hrs, Larvae L3 |
| 21. Adult Inseminated Female Spermatheca | 21. Adult Male Midgut | 21. Fat Body, Pupae P8 | 21. Tumorous Blood Cells Mbn2 | 21. Rotenone 8 µg 12 Hrs, Larvae L3 |
| 22. Adult Ovary | 22. Adult Female Hindgut | 22. Carcass, 3rd Instar Larvae Wandering | 22. Ovary Fgs/OSS | 22. Sindbis Virus, Larval Stage |
| 23. Adult Testis | 23. Adult Male Hindgut | 23. Carcass, 1-Day Adult | 23. Ovary OSC | 23. Sindbis Virus, Pupal Stage |
| 24. Adult Male Accessory Gland | 24. Adult Female Malpighian Tubule | 24. Carcass, 4-Day Adult | 24. Ovary OSS | 24. Sindbis Virus, 4-Day Adult Male |
| 25. Adult Carcass | 25. Adult Male Malpighian Tubule | 25. Carcass, 20-Day Adult |  | 25. Sindbis Virus, 4-Day Adult Female |
|  | 26. Adult Female Fat Body | 26. Ovary, Virgin 4-Day Female |  |  |
|  | 27. Adult Male Fat Body | 27. Ovary, Mated 4-Day Female |  |  |
|  | 28. Adult Female Salivary Gland | 28. Testis, Mated 4-Day Male |  |  |
|  | 29. Adult Male Salivary Gland | 29. Accessory Gland, Mated 4-Day Male |  |  |
|  | 30. Adult Female Heart |  |  |  |
|  | 31. Adult Male Heart |  |  |  |
|  | 32. Adult Female Rectal Pad |  |  |  |
|  | 34. Adult Female Virgin Spermathecum |  |  |  |
|  | 35. Adult Female Mated Spermathecum |  |  |  |
|  | 36. Adult Female Ovary |  |  |  |
|  | 37. Adult Male Testis |  |  |  |
|  | 38. Adult Male Accessory Gland |  |  |  |
|  | 39. Adult Female Carcass |  |  |  |
|  | 40. Adult Male Carcass |  |  |  |
|  | 41. Adult Female Whole |  |  |  |
|  | 42. Adult Male Whole |  |  |  |

Moreover, we also examined the co-expression of *Col4a1*, *vkg* and *P4H*⍺*-*related genes using single-nucleus transcriptome data available from the Fly Cell Atlas [52] in two target tissues ovary and fat body, which are known to express collagen IV [24, 53], as well as the whole-body dataset. To reduce the sparsity of the data, we aggregated neighbouring cells into metacells, where the expression levels of each gene were averaged. Below, all the assays were done using these metacells unless otherwise stated. As in the transcriptome datasets above, we examined the co-expression of *Col4a1*, *vkg* and all the 26 collagen P4H⍺-related genes. In all the three single cell datasets, we confirmed the strong co-expression of *Col4a1* and *vkg* (*r* > 0.90) (Figure 3A, open squares; raw values are in Table S3) and the prominent co-expression of *Col4a1* with *PH4*⍺*EFB* among the 26 P4H⍺*-*related genes (Figure 3A, magenta and black circles) consistent with Figure 2. Collectively, these results indicate a high level of co-expression between *PH4*⍺*EFB* with *Col4a1* at the tissue and single cell levels.

**Figure. 3.**
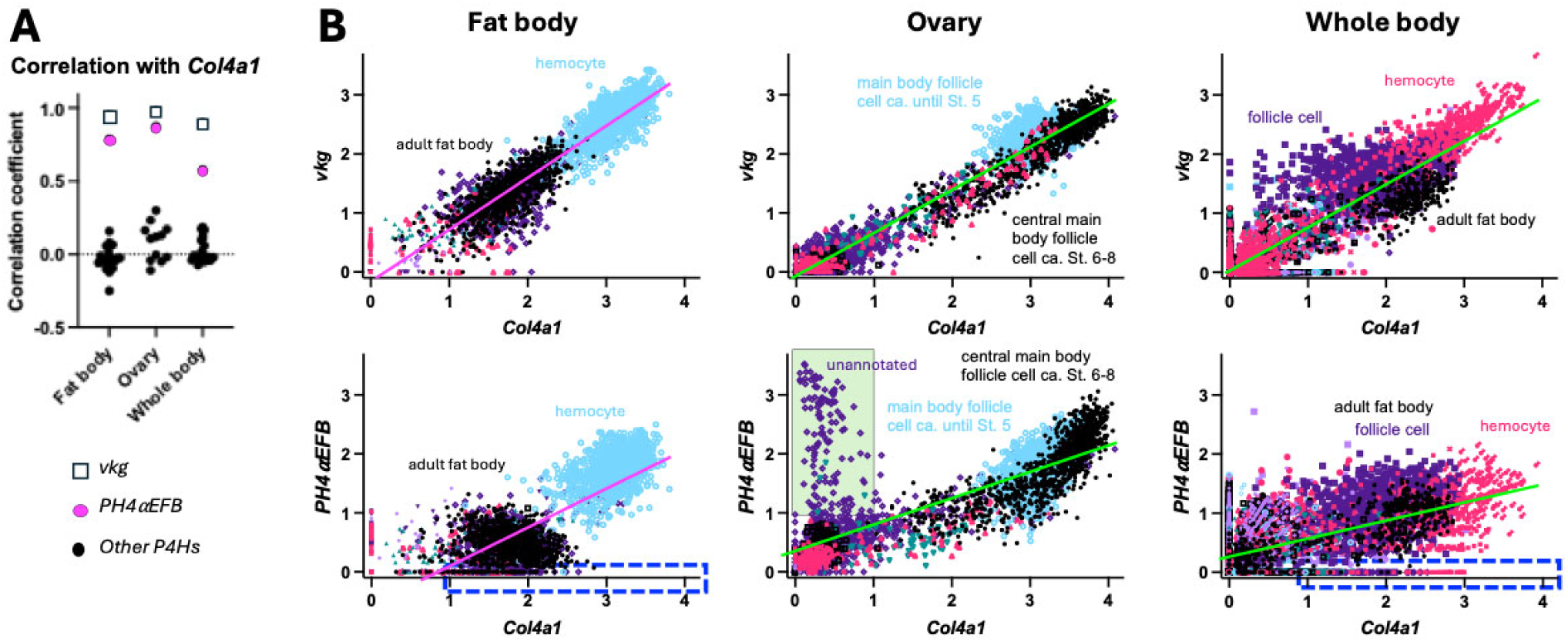
Co-expression of *Col4a1* and *PH4*α*EFB* in single cells. **(A)** Plot of correlation coefficient *r* between *Col4a1* and *vkg* or each of the 26 PH4⍺-related genes in the indicated datasets. **(B)** The expression levels of *Col4a1* vs. *vkg* (top) or *PH4*⍺*EFB* (bottom) in each single metacell were plotted for the indicated datasets. Annotations of representative cell types are shown. For full annotations, see Fig. S1. Regression lines for all the data points in each panel are shown; their colours are altered only for the sake of visibility. n = 3331 (fat body), 3408 (ovary), and 10186 (whole body). Blue dashed-edge rectangles, the area of high *Col4a1* (> 1) and low *PH4*⍺*EFB* (< 0.1) expression; green interior rectangle, the area of low *Col4a1* (< 1) and high *PH4*⍺*EFB* (> 1) expression. The latter rectangle contains 133 metacells, 129 of which are unannotated and the remaining 4 are stretch follicle cells (Cf. Table S3).

Next, to further confirm the co-expression of *PH4*⍺*EFB* with *Col4a1* in these single metacell datasets, we implemented a high-dimensional weighted correlation network analysis in the R package hdWGCNA [54], which detects co-expression modules for each cell type within a given tissue. Across all analyses, the only *P4H*⍺-related gene that shared a co-expression module with the collagen subunits was *PH4*⍺*EFB*. To determine whether the co-occurrence was statistically significant, we performed a Monte Carlo permutation test: module assignments for our candidate and collagen IV subunits for co-expression were randomised 10,000 times, and the number of co-assigned modules was recorded or a randomly selected candidate gene in each iteration. The resulting p-values are the proportion of randomisations where the number of matched module assignments was equal to or exceeded the empirical value. In all three datasets, we detected a significant enrichment of *PH4*⍺*EFB* (Table 3), suggesting potential co-regulation with collagen IV subunits. Therefore, we plotted the expression levels of *Col4a1* vs. *vkg* or *PH4*⍺*EFB* for each single metacell to visualise their co-expression (Figures 3B and S1). In all the fat body, ovary, and whole-body datasets, we found that the plotted points for *Col4a1* vs. *vkg* largely aligned along a single regression line with a positive slope, indicating a prominent positive correlation in their expression (Figures 3B top and Figure S1). Similarly, most of the points for *Col4a1* vs. *PH4*⍺*EFB* also aligned along one regression line with a positive slope in all the three datasets (Figure 3B bottom and Figure S1). These positive correlations between *Col4a1* and *vkg/PH4*⍺*EFB* were also observed in many individual cell types separately analysed (Figs. S2-4). We also visualised the co-expression of the collagen IV genes and PH4⍺EFB on the uniform manifold approximation and projections (UMAPs). Co-expression of *Col4a1* and *vkg* or *PH4*⍺*EFB* was confirmed in cells such as hemocytes, follicle cells, and glia in the fat body, ovary, and whole-body datasets, respectively (closed arrows in Figure S5, Figure 4, and Figure S6, respectively). In UMAPs, co-expression of *Col4a1* and *vkg* or *PH4*⍺*EFB* in cells with lower gene expression (e.g., ‘adult fat body cells’ in the fat body dataset, indicated by open arrows in Figures. 4 and S5) was less clear than in metacell data (Figure 3B), likely due to lower expression levels and higher noise in raw single-cell data. Taken together, we conclude that these metacell plots and UMAPs indicate a high level of co-expression between *PH4*⍺*EFB* with both *Col4a1* and *vkg* at the single cell level in various tissues.

**Table 3.**
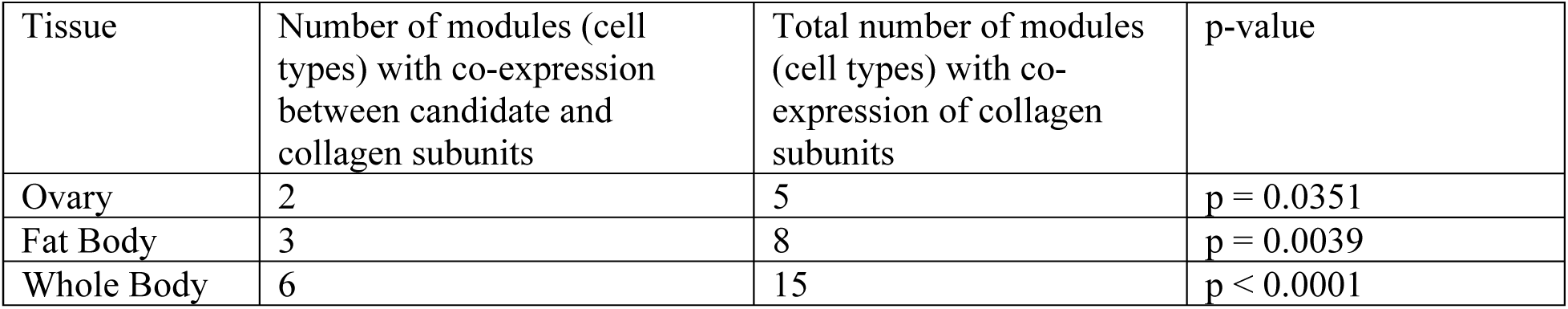
Statistical significance of co-occurrence of *PH4*α*EFB*, *Col4a1* and *vkg* in co-expression modules.

**Figure. 4.**
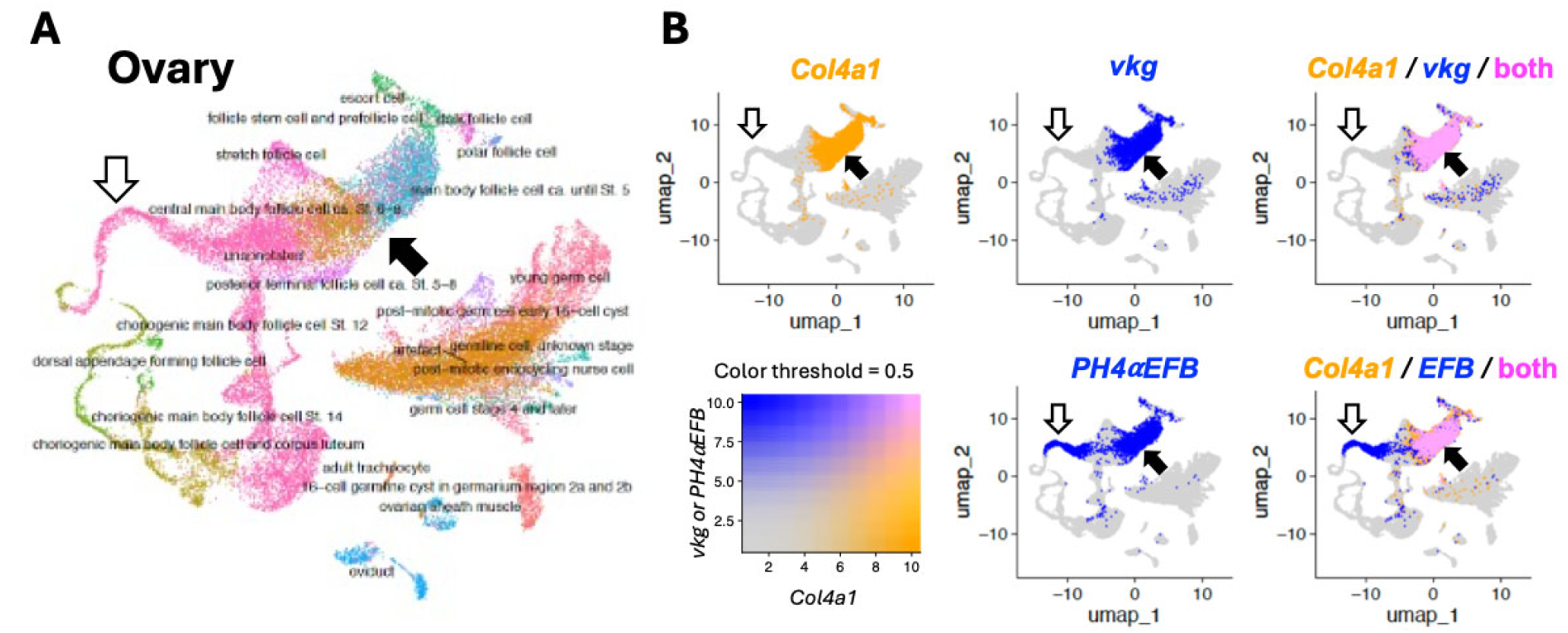
Uniform manifold approximation and projection (UMAP) for the ovary single cell data. **(A)** UMAP showing the entire cells with annotations, with different cell types coded by different colour. **(B)** Expression of *Col4a1*, *vkg,* and *PH4*⍺*EFB (EFB)* colour coded as in the bottom left panel. Top left and middle panels show single gene expression; right panels show overlap. Closed arrows, follicle cells in which the three genes are co-expressed; open arrows, cells that express *PH4*⍺*EFB* but not the collagen IV genes; these cells should correspond to the metacells within the green interior rectangle in Figure. 3B.

It is worth noting that not all cells showed the *Col4a1*-*PH4*⍺*EFB* correlation as mentioned above, although the number of such outliers was not high enough to noticeably affect the overall regression lines or correlation coefficients. For example, there existed ‘low (or no) *PH4*⍺*EFB* but high *Col4a1*’ expression cells in the fat body and whole-body datasets (Figure 3B bottom, dashed-edge rectangles). Conversely, in the metacell plot of the ovary dataset, there was a group of metacells with ‘high *PH4*⍺*EFB* but low *Col4a1’* expression, many of which are unannotated cells (green interior rectangle in Figure 3B bottom centre; Figure S3 bottom right, ‘unannotated’). These metacells should correspond to the cluster of ‘unannotated’ cells with the area of ‘high *PH4*⍺*EFB* but low collagen genes expression’ in the UMAP (Figure 4, open arrows). Potential mechanisms of collagen IV modification and functions of PH4⍺EFB in these ‘low *PH4*⍺*EFB*-high *Col4a1*’ and ‘high *PH4*⍺*EFB*-low *Col4a1*’ expression cells will be explored further in the Discussion.

As the final transcriptome analysis, we compared the expression time courses of the collagen P4H⍺-related and collagen IV genes using the modENCODE Development RNA-Seq data [55]. The expression of *PH4*⍺*EFB* was found to precede that of the collagen IV genes and continue throughout life, suggesting that PH4⍺EFB is always ‘ready’ to hydroxylate newly synthesised collagen IV cains (Figure 5 and Table S4). In conclusion, these spatial and temporal expression patterns are highly suggestive that PH4⍺EFB plays the major and potentially a constitutive role in the proline 4-hydroxylation of collagen IV in *Drosophila*.

**Figure. 5.**
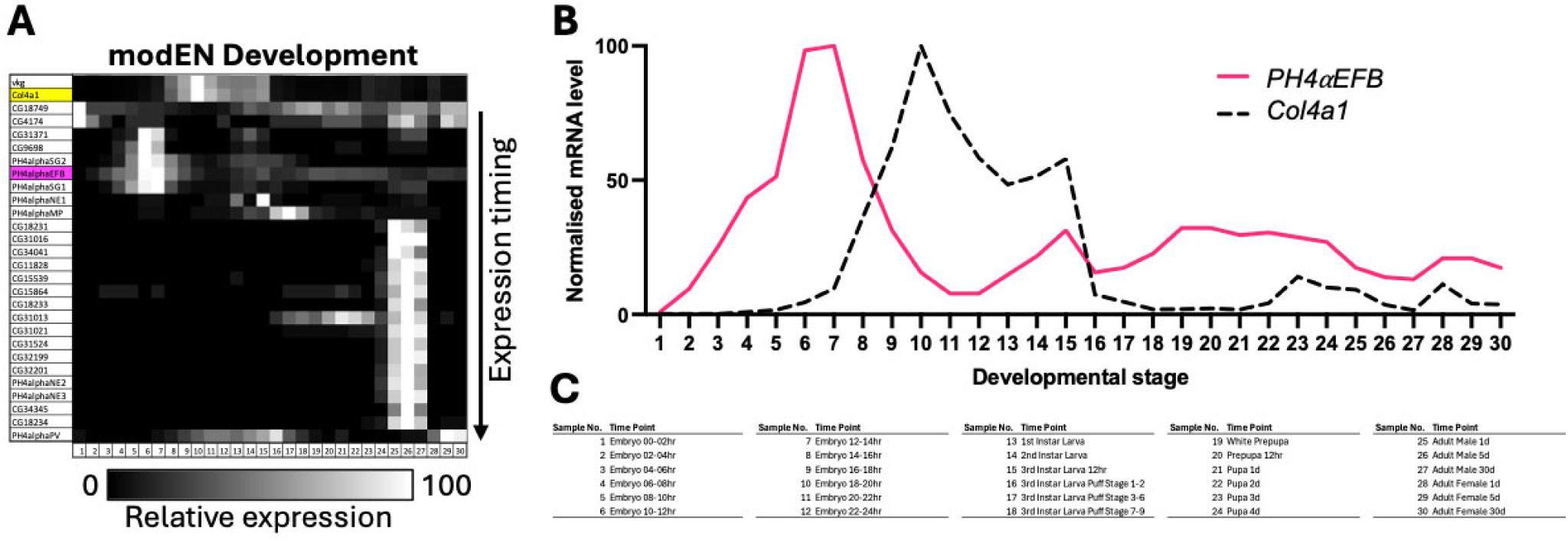
Expression time courses of *Drosophila* collagen IV and PH4⍺-related genes. **(A)** The expression levels of each gene from the modENCODE (modEN) Development transcriptome dataset are normalised for the maximum level and brightness-coded as in the bar at the bottom. *vkg* is at the top. Yellow, *Col4a1*; magenta, *PH4*⍺*EFB*. The PH4⍺-related genes are sorted according to the timing of expression peak: genes with earlier peak are located higher. For enlarged gene names and the values of gene expression levels, see Table S4, where the genes are sorted in the same order. **(B)** Expression time courses of *PH4*⍺*EFB* and *Col4a1* extracted from (A) and displayed in the 2D scattered plot format. **(C)** Legend for the sample numbers in the horizontal axes of **(A)** and **(B)**.

### PH4αEFB is a collagen binding Drosophila P4H***α***

Finally, we examined the collagen-binding ability of PH4⍺EFB using a gelatine pulldown assay. *Drosophila* D17 cells express 6 potential collagen P4H⍺-related genes including *PH4*⍺*EFB* (≥ 1 RPKM in sample 15, modENCODE_Cell_Lines, Sup_Table_1). We incubated D17 cell lysate with gelatine-coupled beads and identified gelatine-bound proteins by protein ID LC-MS analysis. As a positive control, we used mouse PFHR9 cells, which are known to express collagen α1α1α2(IV) and its binding proteins [9, 56] (Figures 6A-C). From the PFHR9 cell extract, many collagen-interacting proteins such as HSP47 and collagen P4HAs were pulled down, demonstrating the reliability of this assay. Importantly, PH4⍺EFB was among the 19 proteins pulled down from the D17 extract (Figures 6C and D), indicating its collagen-binding ability. Importantly, while the structure of PH4⍺EFB predicted its binding to PDI [29] (Table S1, Figure 1A), *Drosophila* PDI indeed co-precipitated PH4⍺EFB (Figures 6C and E). Thus, our result strongly supports that PH4⍺EFB directly binds to collagen and forms a complex with PDI under physiological conditions.

**Figure. 6.**
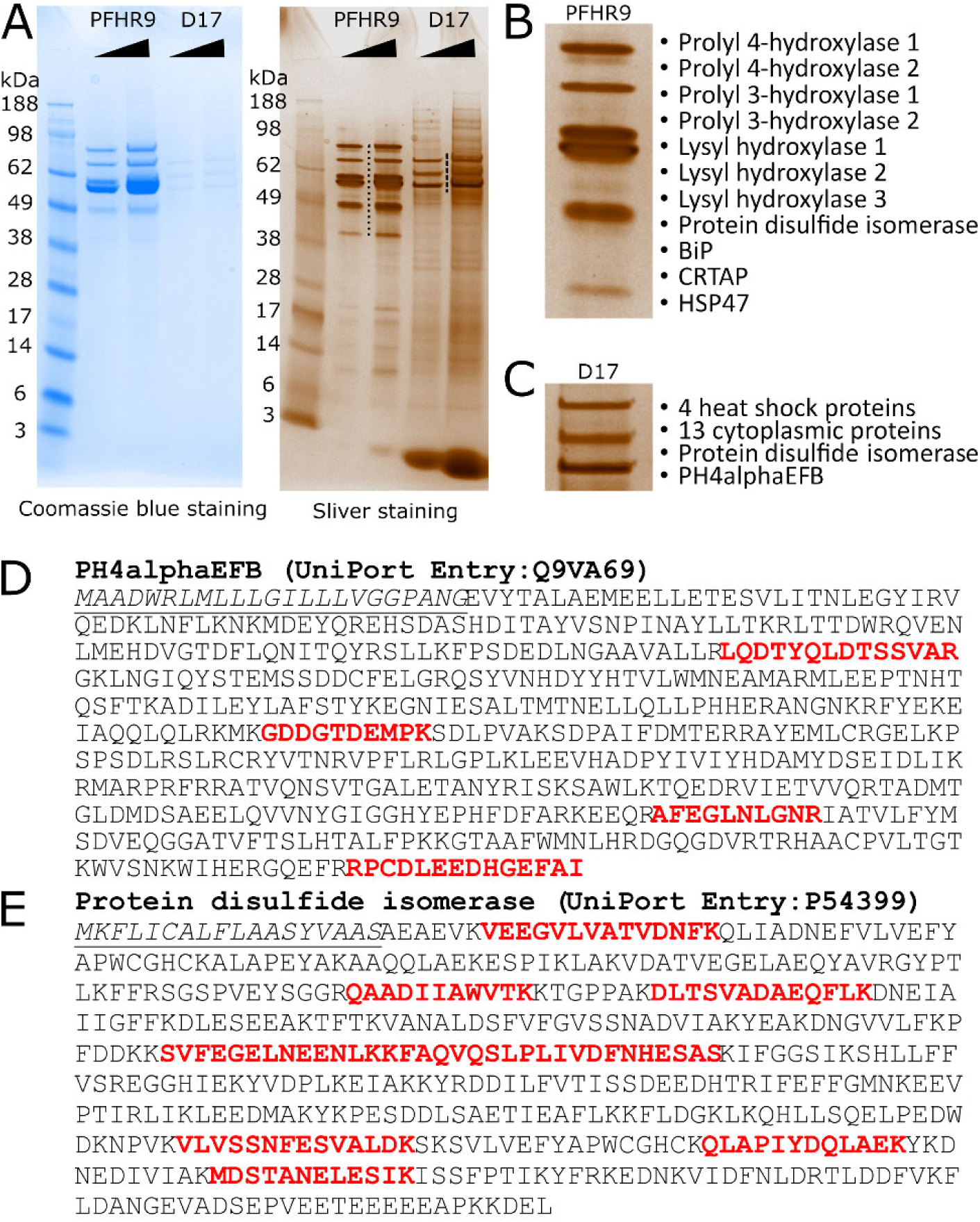
PH4αEFB is captured by gelatine Sepharose from Drosophila D17 cells lysate. **(A)** Mouse and *Drosophila* collagen binding proteins extracted from gelatine Sepharose with SDS sample buffer after extensive NaCl washes. The eluted samples were loaded onto the gels in two different volumes and stained with coomassie blue and silver staining. **(B and C)** The magnified images of the area annotated with dots and dash lines for Mouse **(B)** and *Drosophila* **(C)** collagen binding proteins in the sliver stained gel. Protein names were identified by protein ID LC-MS analyses. The details of protein ID LC-MS results are in Fig. S7. In **(B)**, ‘Prolyl 4 hydroxylase 1’ and ‘2’ correspond to P4HA1 and 2, respectively. **(D and E)** The identified peptides from *Drosophila* PH4⍺EFB **(D)** and protein disulfide isomerase **(E)**, as determined by protein ID LC-MS. The underlined italic and red colour fonts indicate the signal peptide and identified peptides, respectively. The lists of identified proteins by MS (Mascot data) are in Supplementary Source Data.

## Discussion

In *Drosophila*, despite its relatively simple genome, there are at least 26 potential collagen P4H⍺s, and their roles have not been fully elucidated. PH4⍺EFB and PH4⍺MP have been suggested to mediate collagen IV hydroxylation [36, 39–41], which is the only major collagen in *Drosophila*. However, almost nothing is known about the functions of the other potential collagen P4H⍺s, and some of them may also modify collagen IV. In this study, our thorough bioinformatic and biochemical analyses corroborate that PH4⍺EFB plays the major and potentially a constitutive role in the proline 4-hydroxylation of collagen IV. This conclusion is also supported by several other studies. First, there are multiple reports of the enriched expression of *PH4*⍺*EFB* in embryonic macrophages [36, 57], which are the major source of collagen IV in the *Drosophila* embryo [37, 56]. Consistently, mutants lacking macrophages show reduced expression of *PH4*⍺*EFB* [58]. Moreover, knockdown of PH4⍺EFB alone is sufficient to disrupt collagen IV secretion from the fat body and ovarian follicle cells [39, 40].

However, our findings do not entirely rule out the role of P4H⍺s other than PH4⍺EFB in the proline 4-hydroxylation of *Drosophila* collagen IV in specific tissues or contexts. Notably, our sequence analysis revealed that apart from *CG34041*, all the other 25 *Drosophila* collagen P4H⍺-related genes encode proteins that harbour the ‘N-PSB-L-CAT’ domain organisation, a hallmark structure of collagen P4Hs [30, 31, 42, 43] (Figure 1A). Therefore, the products of any of these 25 genes may work on collagen IV (or/and other collagen(-related) molecules such as Multiplexin or Pericardin as discussed later). Indeed, PH4⍺MP hydroxylates collagen peptides *in vitro* [36, 41] and clusters near PH4⍺EFB and human collagen P4HAs in the phylogenetic tree (shaded dark grey in Figure 1F), suggesting potential similarity in substrate specificities. If PH4⍺MP targets collagen IV, other related P4H⍺s including PH4⍺SG1, SG2, and NE1 (shaded light grey in Figure. 1F) might also do so. Consistently, several observations support their possible roles in collagen IV hydroxylation *in vivo*. First, *PH4*⍺*SG1* and *SG2* are highly expressed in the larval salivary gland, where *PH4*⍺*EFB* is nearly absent despite *Col4a1* expression (closed arrows in Figure 2C). While PH4⍺SG1 and SG2 are suggested to have other substrate(s) in the embryonic salivary gland [59], they may target collagen IV in later stages. Moreover, in the *Drosophila* embryo, the expression of *PH4*⍺*SG1* (but not *PH4*⍺*EFB*) is reported to be enriched in caudal visceral mesoderm (CVM) cells, another collagen IV source [57]. These findings suggest a possible role for PH4⍺SG1 and 2 in the salivary gland and for PH4⍺SG1 in CVM cells. Furthermore, in the modENCODE Treatments dataset, *PH4*⍺*SG1, SG2,* and *NE1* expressions show high correlations with *Col4a1*expression (grey arrows in Figures. 2A and B,), similarly to *PH4*⍺*EFB* in other datasets. In Figure 2C, when *Col4a1* expression rises (sample 16 – 20), *PH4*⍺*EFB* also rises (dotted bracket). However, from sample 20 to 21 (open arrow), *Col4a1* rises sharply while *PH4*⍺*EFB* slightly decreases. Here, *PH4*⍺*SG1*, *SG2*, and *NE1* expression increase instead. In the modENCODE Treatments dataset, gene expression changes in response to various environmental perturbations were examined. For example, cadmium (sample 4 in Figure 2C), ethanol (sample 13), and rotenone (sample 20) induced *PH4*⍺*SG1* expression, while sindbis virus (sample 22) induced *PH4*⍺*SG2* and *NE1* expression. Thus, PH4⍺SG1, SG2, and NE1 induced by various stresses may compensate for or support PH4⍺EFB in collagen IV hydroxylation. Nevertheless, we did not detect PH4⍺SG1 binding to gelatine beads in our biochemical analysis using *Drosophila* D17 cells, despite reported *PH4*⍺*SG1* expression in this cell line (Figure 2A and Table S2, sample 15). Possible explanations include loss of *PH4*⍺*SG1* expression in our cell sub-strain, lack of translation, or inability to bind gelatine. These possibilities warrant future investigation.

In the single nucleus data, we detected a small population of metacells with low (or no) *PH4*⍺*EFB* but high *Col4a1* (Figure. 3B). We could not easily pinpoint these cells in UMAPs due to dispersion and noise (Figures. S5 and S6). We hypothesised that other enzymes might be modifying collagen IV in these cells, but could not identify clear candidates, as most of these cells did not express any of the 25 other PH4⍺-related genes (Table S5). We speculate that in these cells, collagen P4H⍺ mRNA, regardless of whether it is for PH4⍺EFB, might have been degraded while the translated enzyme remains active, or collagen IV mRNAs are not translated: the levels of an mRNA and the protein it encodes do not always correlate [60].

While PH4⍺EFB might not be the sole P4H⍺ for *Drosophila* collagen IV, collagen IV might not be the only substrate for PH4⍺EFB either. The correlation between *Col4a1* and *PH4*⍺*EFB* expression is weaker than that between *Col4a1* and *vkg*, which encodes the other subunit of *Drosophila* collagen IV (Figures. 2-4 and S2-S6). Figure 2C shows several *PH4*⍺*EFB* expression peaks accompanied with low *Col4a1* expression (asterisks). Similarly, single nucleus analysis revealed a group of cells with high *PH4*⍺*EFB* but low *Col4a1* (Figures. 3, 4, and S3). Consistent with this, the follicle cells of the egg chamber stop making collagen IV at the end of stage 8 [53, 61, 62], and yet a subset of the follicle cells (border and centripetal cells) continue to express *PH4*⍺*EFB* into stages 9 and 10, where it is required for their migration [63]. Likewise, *PH4*⍺*EFB* is expressed in the embryonic epidermis [36], where collagen IV expression is not detected [37, 56]. These findings suggest that PH4⍺EFB may target other proteins in these cells. Alternatively, some cells might have ceased collagen IV expression and degraded its mRNA, while PH4⍺EFB mRNA persists after fulfilling its function. To explore these possibilities, future studies should examine the temporal expression patterns of *PH4*⍺*EFB* and *Col4a1* at higher resolution and determine PH4⍺EFB’s full substrate spectrum.

In addition to collagen IV, *Drosophila* harbours several collagen(-related) molecules such as Pericardin and Multiplexin [21, 22]. It is intriguing to examine whether PH4⍺EFB also modifies them as a ‘pan *Drosophila* collagen prolyl hydroxylase’, and/or whether any other enzymes modify these proteins. Future experiments exploring the correlation between the phylogeny and substrate specificities of the fly collagen P4H⍺-related proteins in Figure 1F will provide evolutionary insights into the diversification of the ‘collagen molecular ensemble’.

From an evolutionary perspective, it is also noteworthy that some molecules exhibit non-canonical domain organisations. While typical collagen P4H⍺s comprise the N, PSB, L, and CAT (catalytic) domains [30, 31, 42, 43] (Figure 1A), CG15539-PA possesses a truncated N-domain. Moreover, the three isoforms of CG34041 all lack the catalytic domain: CG34041-PD contains one N-domain, whereas CG34041-PE and - PF harbour two tandem N-domains (Figure 1A–E). Currently, the functions of these proteins are totally unknown. Because the N-domain is the interface for the dimerisation of P4H⍺ proteins and the CAT domain is used not only for the enzymatic activity but also for the interaction with PH4β/PDI [31], we speculate that CG15539-PA may have lost the ability to dimerise while still being able to bind to PDI, potentially altering its catalytic activity and/or substrate specificity. Regarding CG34041 isoforms, they may interact with other P4H⍺ molecules via their N-domain(s) either homo- or heterotypically; the consequences of such interactions are difficult to predict. Exploring the functions of these non-canonical collagen P4H⍺-related proteins would shed light on the diversification of regulatory mechanisms governing prolyl hydroxylation reactions.

Our results may also provide some hints toward the search for non-collagen substrates of the collagen P4H⍺-related proteins. For example, 18 out of the 26 *Drosophila* collagen P4H⍺-related genes are highly expressed in the male accessory gland (red arrows in Figure 2A), which produces seminal fluid components and is analogous to the human prostate [64]. Interestingly, a seminal fluid component ‘Sex Peptide (SP)’ is known to be 4-prolyl hydroxylated. SP is transferred to the female during copulation and elicits a wide range of post-mating changes in female physiology and behaviour, such as rejection of further mating, increased food intake, enhanced oviposition, and the augmentation of immune response [65–71]. Among these changes, at least that of the immune response is mediated by the central region of SP containing 4Hyps [67]. Therefore, the accessory gland P4H⍺s may include the hitherto unidentified enzyme(s) responsible for the 4-prolyl hydroxylation of SP. Indeed, in the phylogenetic tree, all 18 accessory gland P4H⍺s (Figure 1F, blue and marked with daggers) fall outside the clade containing PH4⍺EFB (magenta) and the proteins implicated in collagen modification earlier in this study and by others [41] (grey). Moreover, 16 of the accessory gland P4H⍺-like proteins cluster in the region most distant from the magenta and grey clades of (potential) collagen P4H⍺s (Figure 1F, bracket). This pattern may reflect differences in substrate specificity between accessory gland P4H⍺-related proteins and collagen P4Hs. The roles of the accessory gland P4H⍺related proteins in reproduction represent intriguing targets of future research.

In summary, our research identifies PH4⍺EFB as the primary P4H⍺ for *Drosophila* collagen IV, highlighting a remarkably simple enzymatic toolkit for collagen IV biosynthesis in *Drosophila,* involving likely single isoforms of P4H, LH, and GLT25D. Notably, while mammals require additional modifications such as proline 3-hydroxylation, which plays critical roles in the interactions between other ECM proteins with collagen IV and in basement membrane integrity [16, 72–75], *Drosophila* collagen IV biosynthesis proceeds without them, underscoring its minimalistic nature. This minimal *Drosophila* collagen IV biosynthetic machinery offers an exciting avenue for future research. A key direction is to explore whether this simplified system can successfully produce mammalian collagens, especially collagen IV. Conversely, investigating if mammalian machinery can accommodate *Drosophila* collagen IV will reveal crucial insights into the compatibility of these systems. This comparative approach could help clarify which factors, collagen sequences, ER molecular compositions, or both, are critical for optimising *in vitro* collagen biosynthesis. Ultimately, such studies could define the minimal requirements for collagen engineering, a field with significant biomedical and biomaterial implications.

## Experimental procedures

### Identification of Drosophila P4H***⍺***-related genes

In the ‘QuickSearch’ menu at the top page of FlyBase (http://flybase.org/), gene ontology (‘GO’) tab was clicked, ‘molecular function’ was selected from the ‘Data field’ pull-down menu, and the keyword ‘procollagen-proline 4-dioxygenase activity’ was typed into the ‘Enter term’ window. Subsequently, the ‘Search’ button was pressed, to obtain a single match CV (controlled vocabulary) term ‘procollagen-proline 4-dioxygenase activity’. The link on this term was clicked to access its CV Term Report page, which shows that there are 26 genes annotated with this term (Table 1).

### Phylogenetic analysis

Protein sequence comparison and phylogenetic analysis were conducted using Clustal Omega [76]. SwissProt IDs for the annotated protein products of the 26 *Drosophila* P4H⍺-related genes were obtained from FlyBase (Table 1; URLs follow the format: ‘http://flybase.org/reports/’ + ‘FlyBase ID’). For genes annotated with multiple polypeptides, the isoform listed first on the FlyBase page was used in the initial analysis. The same procedure was applied to the *Drosophila* HIF hydroxylase Hph (FlyBase ID: FBgn0264785; SwissProt ID: Q9VN98). SwissProt IDs for the human proteins included in the analysis were as follows: PH4A1 (P13674), PH4A2 (O15460), PH4A3 (Q7Z4N8), PHD1 (Q9GZT9), PHD2 (Q9H6Z9), and PHD3 (Q96KS0).

Preliminary analysis of the 33 proteins above revealed non-canonical domain structures in the protein products of CG15539 and CG34041. To examine whether other annotated products of these genes possess the canonical ‘N-PSB-L-CAT’ structure, a second Clustal Omega analysis was performed including all annotated isoforms for CG15539 and CG34041 (Table 1). For clarity, P4HA1, P4HA3, PHD1, PHD2, and Hph were excluded from the analysis. The resulting multiple sequence alignment was visualised in Table S1 according to <u>this website</u> (https://ouchidekaiseki.com/align.php - note that this website is written in Japanese). Briefly, the file ‘*Alignment in CLUSTAL format with base/residue numbering’* was downloaded from the ‘Results Files’ tab of Clustal Omega and opened in TextEdit (Apple). The contents were copied into Microsoft Excel using the ‘Text Import Wizard’. In the first window of the Wizard, ‘Fixed width’ was selected; in the next, break lines were inserted between residues manually. Upon completion, each amino acid was placed in a separate cell. The cell dimensions were adjusted to form squares, and amino acids were colour-coded using Excel’s ‘Conditional Formatting’. Sequences separated to multiple rows were consolidated into single lines. Human P4HA2 was repositioned to the top of the table, and its domain structure was annotated. Additional formatting (e.g., row numbering, highlights, and colour codes) was applied to enhance clarity.

To construct the phylogenetic tree shown in Figure 1F, a third Clustal Omega run was performed, this time including P4HA1, P4HA3, PHD1, PHD2, and Hph. For clarity, only one protein product with a canonical domain structure was analysed for each *Drosophila* P4Hα-related gene. Specifically, all CG34041 isoforms were excluded, and only the PB isoform of CG15539 was included. The resulting dendrogram was downloaded in SVG format and edited using Adobe Illustrator and Microsoft PowerPoint, preserving branch topology and relative branch lengths.

### Obtaining microarray/RNA-seq data

For each P4H⍺-related gene (Table 1), *Col4a1* (FlyBase ID FBgn0000299) and *vkg* (FBgn0016075), gene report was obtained on FlyBase (http://flybase.org/reports/FBID, where ‘FBID’ is the FlyBase ID of the gene [Table 1]). In the Expression Data > High-Throughput Expression Data section, the following five datasets were opened:

1. FlyAtlas Anatomy Microarray [49]
2. FlyAtlas2 Anatomy RNA-Seq [50]
3. modENCODE Anatomy RNA-Seq [48]
4. modENCODE Development RNA-Seq [55]
5. modENCODE Cell Lines RNA-Seq [48]
6. modENCODE Treatments RNA-Seq [48]

Subsequently, gene expression data were downloaded from the ‘download data (TSV)’ links. From the obtained files, the Mean Affy2 Probeset Expression Values (for FlyAtlas Anatomy Microarray) and RPKM values (for the others) were summarised and analysed in Supplementary Tables 1 (spatial data, 1-3, 5 and 6 above) and 2 (temporal data, 4).

### Brightness coding of gene expression patterns

Normalised gene expression levels (values marked cyan in Supplementary Tables YMS1 and 2) were saved as text files and opened as images with ImageJ/Fiji, using File > Import > ‘Text Image…’ command.

### Single-cell data acquisition and analysis

*Drosophila melanogaster* single-nucleus RNA-sequencing (snRNA-seq) datasets for fat body, ovaries and whole body were obtained from the Fly Cell Atlas [52]. Loom files and H5AD files were integrated and converted to RDS format using SCANPY [77]. Quality control and subsequent analysis were implemented in the R package Seurat [78]. Specifically, we retained cells that expressed a minimum of 200 and a maximum of 2000-3500 genes, depending on tissue type. We then normalised the gene expression measurement for each cell by the total expression, multiplied this result by 10,000 and log-transformed the result. The 2000 most variable genes were selected to aid in detecting biological signal in downstream analysis. These were used for linear transformation of the data (scaling), followed by principal components analysis, retaining the first 50 components. Unsupervised clustering of cells was conducted using the standard Seurat pipeline. This constructs a K-nearest neighbour graph (KNN) based on the Euclidean distance in the PCA space. Clusters were defined using the Louvain algorithm with a resolution of 0.5 and visualised using the non-linear dimensional reduction technique uniform manifold approximation and projection (UMAP). To visualise co-expression, we overlaid expression of *Col4a1*, *vkg*, and *PH4*⍺*EFB* on tissue-specific UMAPs.

For the other analyses, we processed the data as described below to remove noise and gain clearer information. First, we limited the investigation to genes expressed in at least 5% of cells. To reduce the sparsity of the data, we aggregated neighbouring cells with similar expression levels using the MetacellsByGroups function in the R package hdWGCNA (parameters k=25 and max_shared=10, reduction = pca) [54]. We normalised expression values in the metacell expression matrix for downstream analysis. For each cell type in each tissue, we then set the expression matrix. For each cell type, the soft power threshold was automatically selected using the lowest power that meets 0.8 scale-free topology fit [54]. Co-expression networks were then constructed using default parameters. We then computed module eigengenes as well as eigengene connectivity to define co-expression structure. Obtained data were plotted and used to compute Pearson correlations either for the entire dataset or within each cell type[79].

To identify which candidate gene(s) were consistently co-expressed with the two known collagen subunits (*Col4a1* and *vkg*), we performed a high-dimensional weighted correlation network analysis in the R package hdWGCNA [54]. We first identified co-expression modules that included both *Col4a1* and *vkg*. We reasoned that if our gene was involved in the proline 4-hydroxylation of collagen IV, it would co-occur within those modules. For each tissue, we counted the number of times each candidate gene was coexpressed in the same module as *Col4a1*, and *vkg*. We then assessed statistical significance using a Monte Carlo permutation test. Module assignments were randomised across 10,000 replicates. We then selected a single gene at random and computed the number of matches for those modules. The resulting p-values are the proportion of randomisations where the number of matched module assignments was equal to or exceeded the empirical value.

#### Drosophila D17 Cell Culture

A detailed protocol for the culture of *Drosophila* D17 cells is presented in the following reference [80].

#### Mouse PFHR9 Cell Culture

PFHR9 cells were cultured following the referenced protocol [9]. After reaching 80-90 % confluency, to stimulate procollagen biosynthesis, ascorbic acid phosphate (100 μg/ml; Wako Chemicals, 013-12061) was supplemented in Dulbecco’s modified Eagle’s medium (DMEM)/high glucose/pyruvate (Gibco, 11995065) containing 10% (v/v) foetal bovine serum (R&D systems, S11150), penicillin streptomycin glutamine 100× (Gibco, 10378016), and 5 mM Hepes for 1 day. The medium was replaced to the fresh DMEM with ascorbic acid, and the cells were cultured for 2 days. After washing with PBS twice, the cells were scraped and transferred into 15 mL falcon tube. The cell pellets were collected by centrifugation with 4000 rpm using TX-400 rotor (Thermo scientific) for 5 min and stored in -20 C.

#### Gelatine coupled beads pull-down assay

D17 and PFHR9 cell pellets (0.5 g each) were lysed with 7.0 mL of pre-cooled M-PER (Thermo Scientific, 78501) containing Halt Protease Inhibitor Cocktail, EDTAfree (Thermo Scientific, 87785) at 4 °C. Following the manufacturer’s instructions, soluble proteins in the supernatant were incubated with 1.5 mL of gelatine coupled beads, gelatine Sepharose 4B (Cytiva, 17095603), for 2 hours at 4 °C. After washing twice with 10 mL TBS buffer followed by one wash with 25 mM Tris buffer, pH 7.4, containing 1 M NaCl and two additional washes with TBS buffer, the proteins tightly bound to gelatine sepharose beads were eluted with 2X SDS-PAGE sample buffer containing DTT by heating to 95°C for 5 minutes. The eluted proteins were separated on a Bolt 4-12 % Bis-Tris Plus gel (Thermo Scientific) using MES running buffer (Thermo Scientific, B0002). Gels were stained with GelCode Blue Stain Reagent (Thermo Scientific, 24592) and SilverXpress Silver Staining Kit (Thermo Scientific, LC6100). The stained gel images were taken by ChemiDoc MP imaging system (Bio-Rad) using the software Image Lab version 4.0.1 (Bio-Rad). Individual gel bands were carefully excised and analysed by protein identification LC-MS performed by the peptide core facility in the research Department of Shriners Hospitals in Portland OR.

## Data processing, statistics, and presentation

ImageJ/Fiji, R (public domain software), TextEdit (Apple), Excel, PowerPoint (Microsoft), Prism (GraphPad), Illustrator (Adobe), and Inkscape (a free and open-source vector graphics editor) were used.

## Acknowledgements

Computational analyses of single-nucleus transcriptome data were performed on the high performance computer (HPC) at Bournemouth University, the HPC at Institute of Science and Technology Austria, and the high-performance computational resources provided by the Louisiana Optical Network Infrastructure (http://www.loni.org). The authors are grateful to the researchers who published the transcriptome datasets [48, 49, 52, 55] that became the essential bases for this study, to FlyBase for curating the datasets in an easily accessible format, and the Drosophila Genomics Resource Center (DGRC), supported by NIH grant 2P40OD010949, for providing the D17 cell line used in this research. The authors thank Kristian Koski (University of Oulu, Finland) for crucial advice on the domain structure of collagen P4H⍺s, and Ryusuke Niwa and Ryo Hoshino (University of Tsukuba, Japan) for helpful discussions on SP.

## Author contributions

YI and YM were responsible for the overall design of the study. YI, MAT, ME, ALZ, and YM conducted and analysed all experiments, and also prepared the figures. YI, MAT, ME, and YM wrote the original draft. All authors prepared essential materials, reviewed and discussed the results, and were involved in editing the manuscript.

## Additional information

Competing Interests: The authors declare that they have no competing interests related to this work.

## Supplementary Table Legends (separated Excel and PDF files)

**Table S1. Alignment of human P4HA2, PHD3, and *Drosophila* P4Hα-related proteins.**

For human P4HA2, the domain structure and residue numbers at domain boundaries are shown at the top, using the same colour scheme as in Figure 1A. The amino acid length of each protein is indicated at the right end of the sequence. *Hs, Homo sapiens*; Dm, *Drosophila melanogaster*. Colour codes for amino acid residues are shown at the bottom left. Human P4HA2 and PHD3 (1 and 2, highlighted in blue and yellow, respectively) are homologous only within the catalytic domain. *Drosophila* CG15539-PA (22, magenta) lacks approximately the first 70% of the N-domain, whereas the other CG15539 isoforms (23 and 24) contain a complete N-domain. The three CG34041 isoforms (bottom of table, green) share homology with other P4H⍺-related proteins in the N-domain and part of the PSB domain, but lacks the catalytic domain.

**Table S2. Spatial expression patterns of *Drosophila* collagen IV and P4Hα-related genes.**

Each sheet shows the values from the transcriptomics data indicated. Raw values of gene expression levels are summarised at the top left, with the *r* values with *Col4a1* shown in blue letters at the right. Sample legend is at the top right, in the cells shaded grey. *Col4a1* and *PH4*⍺*EFB* are marked yellow and magenta, respectively. In the cells marked cyan at the bottom, the values are normalised for the maximum expression level of each gene. The genes are sorted in the same order as in Fig. 2A.

**Table S3. Expression of the Collagen IV and PH4**⍺**-related Genes in Single Metacells.**

Each sheet shows the results from the indicated dataset. Expression levels of the two collagen IV genes (*Col4a1*, *vkg*) and the 26 collagen PH4⍺-related genes in each metacell are shown, together with other information indicated in the top row.

**Table S4. Expression time courses of *Drosophila* collagen IV and P4H**⍺**-related genes.**

Values from the modENCODE Development data. Raw values of gene expression levels are summarised at the top left. Sample legend is at the top right, in the cells shaded grey. *Col4a1* and *PH4*⍺*EFB* are marked yellow and magenta, respectively. In the cells marked cyan at the bottom, the values are normalised for the maximum expression level of each gene. The genes are sorted in the same order as in Fig. 5A.

**Table S5. Expression of the Collagen IV and PH4α-related Genes in ‘High *Col4a1*, Low *PH4*α*EFB*’ Single Metacells.**

From Table S3, the data of the 374 metacells with high *Col4a1* (> 1) and low *PH4*⍺*EFB* (< 0.1) expression were extracted. Column AE shows the maximum expression level of the 25 PH4⍺-related genes excluding *PH4*⍺*EFB*. The value was zero for 240 metacells (Cell AL2), i.e., no expression of non-*EFB* PH4⍺-related genes was detected in these metacells.

**Supplementary Source Data. Identification of gelatine binding proteins by mass spectrometry.**

Below follows the full MASCOT search result file that includes all proteins found in the gel band from sliver staining SDS-PAGE gel showing top-left.

**Figure S1.**
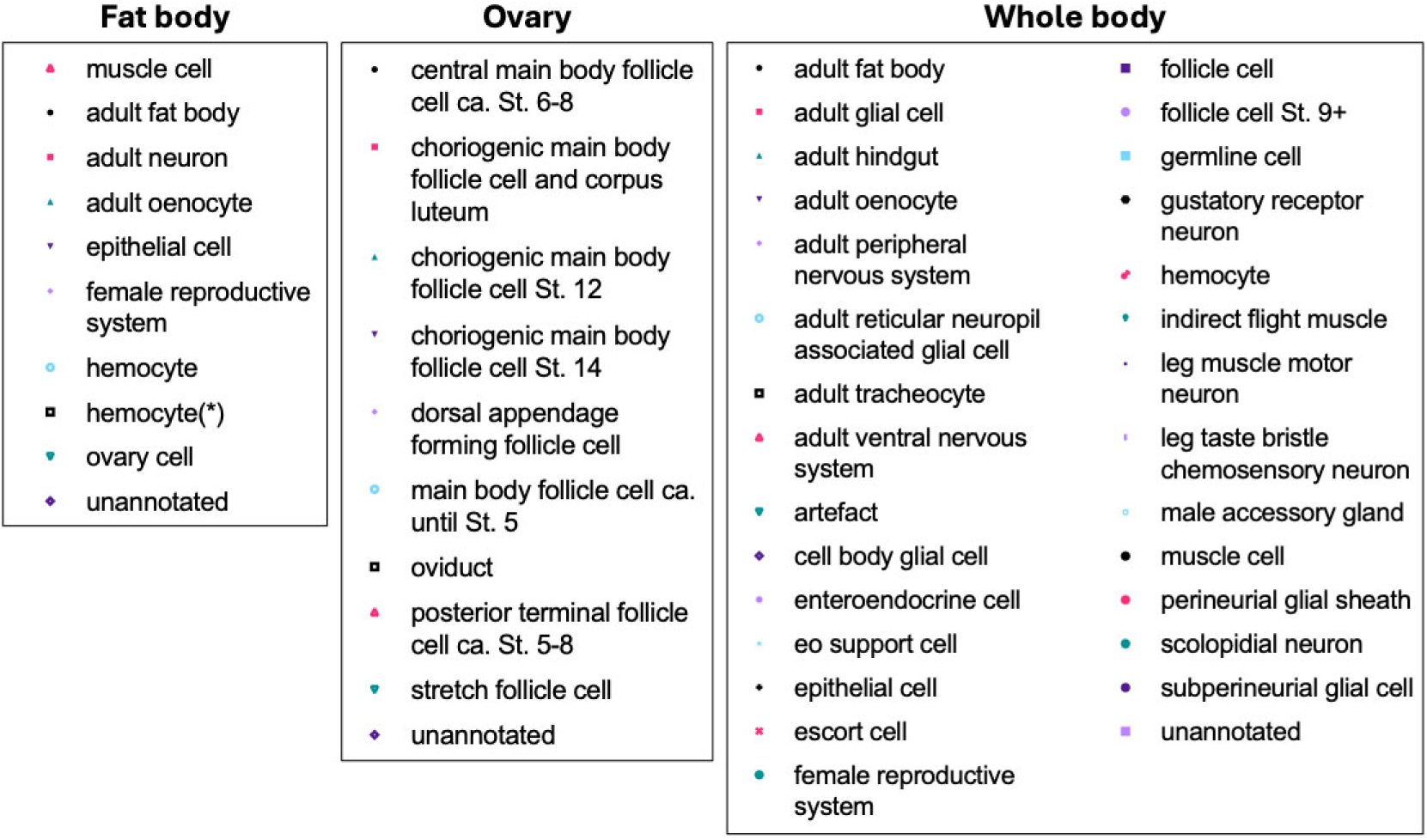
Full legend for the symbols in Figure 3B.

**Figure S2.**
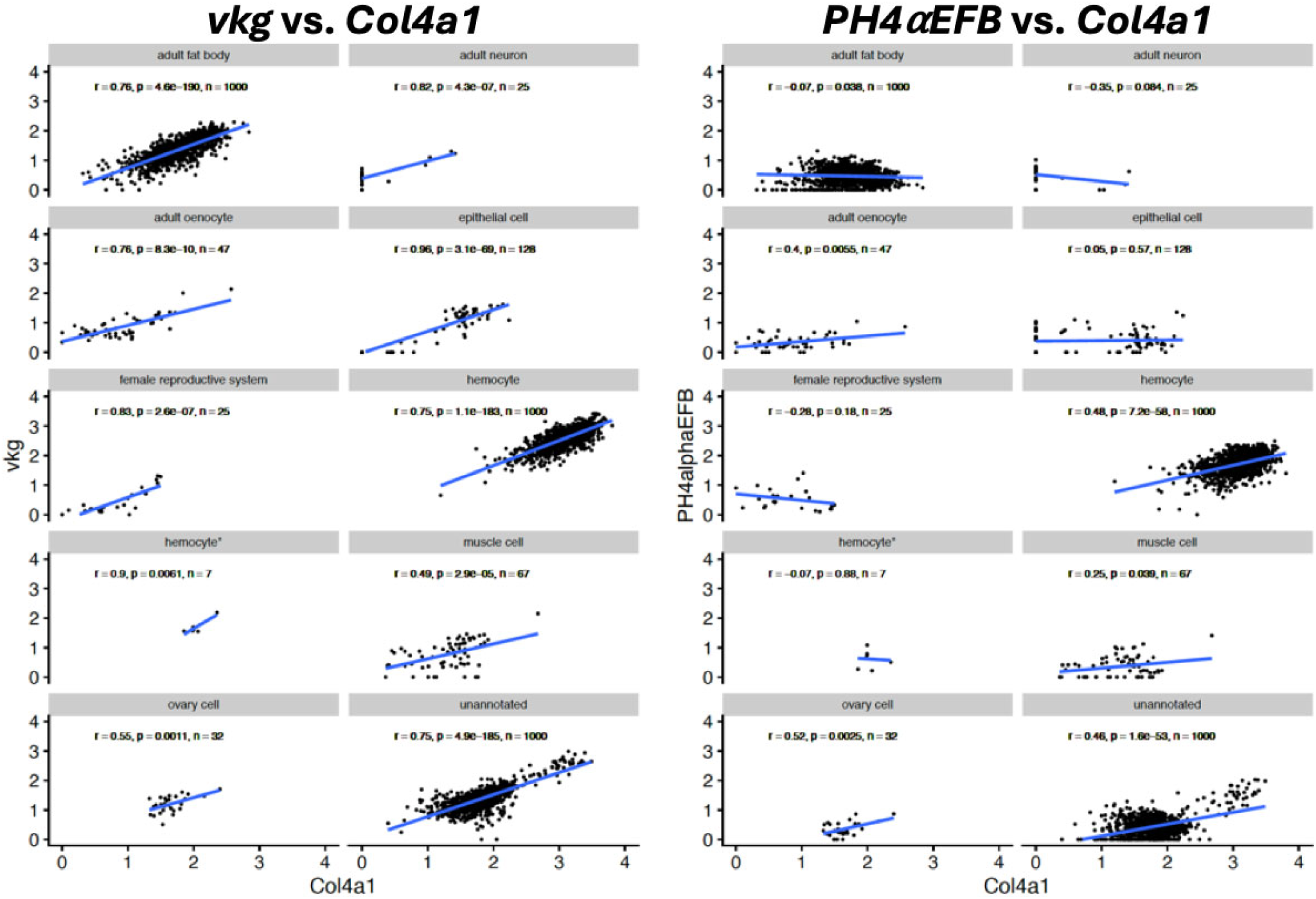
Expression of *Col4a1*, *vkg,* or *PH4*α*EFB* separately analysed for each different cell type in the fat body dataset. Regression lines of the plots, correlation coefficients between the two genes plotted (*r*), the *p* values to obtain the results from the null hypothesis that the slope of the regression lines are zero, and the numbers of metacells analysed (n) are shown.

**Figure S3.**
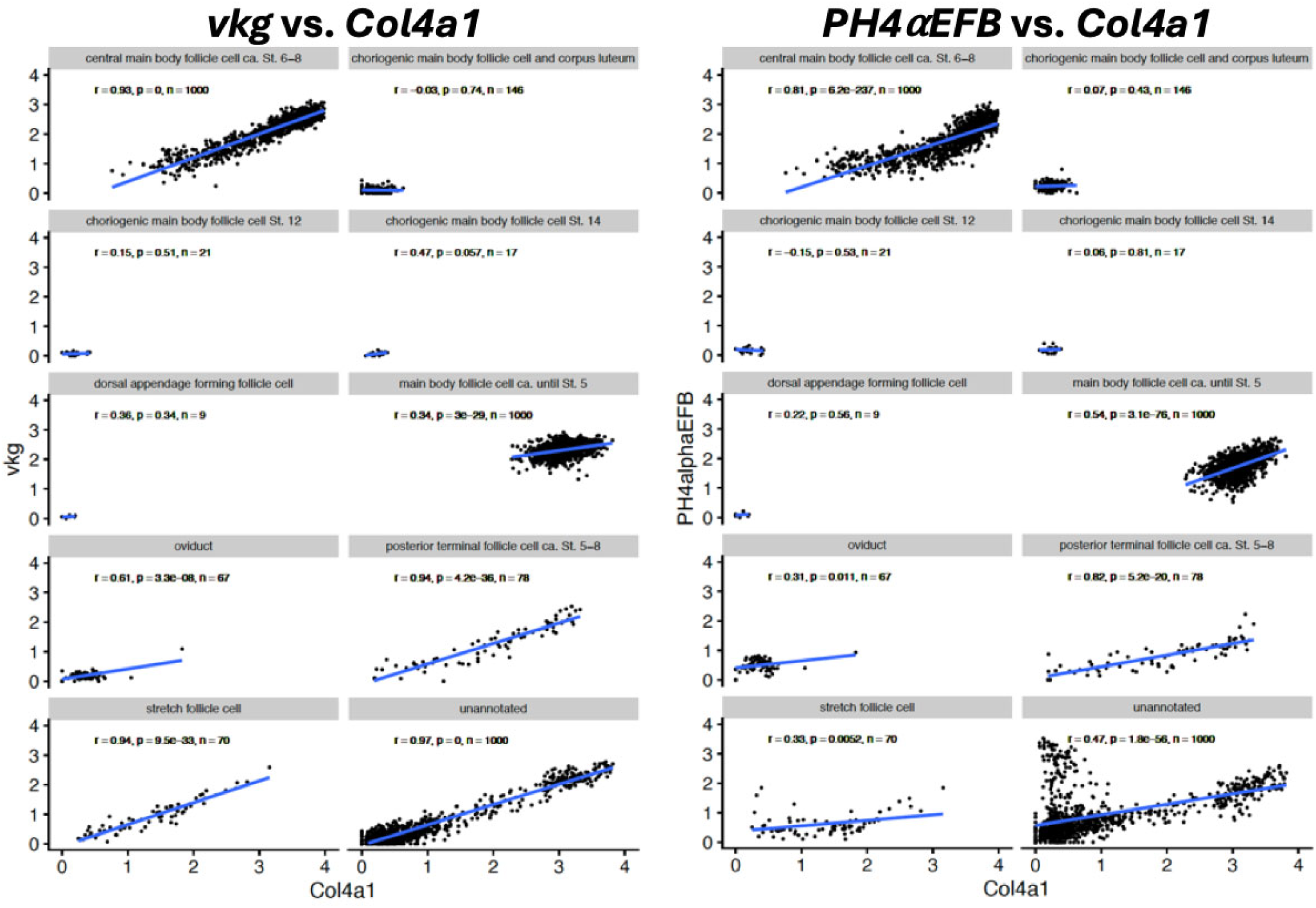
Expression of *Col4a1*, *vkg,* or *PH4*α*EFB* separately analysed for each different cell type in the ovary dataset. Regression lines of the plots, correlation coefficients between the two genes plotted (*r*), the *p* values to obtain the results from the null hypothesis that the slope of the regression lines are zero, and the numbers of metacells analysed (n) are shown.

**Figure S4.**
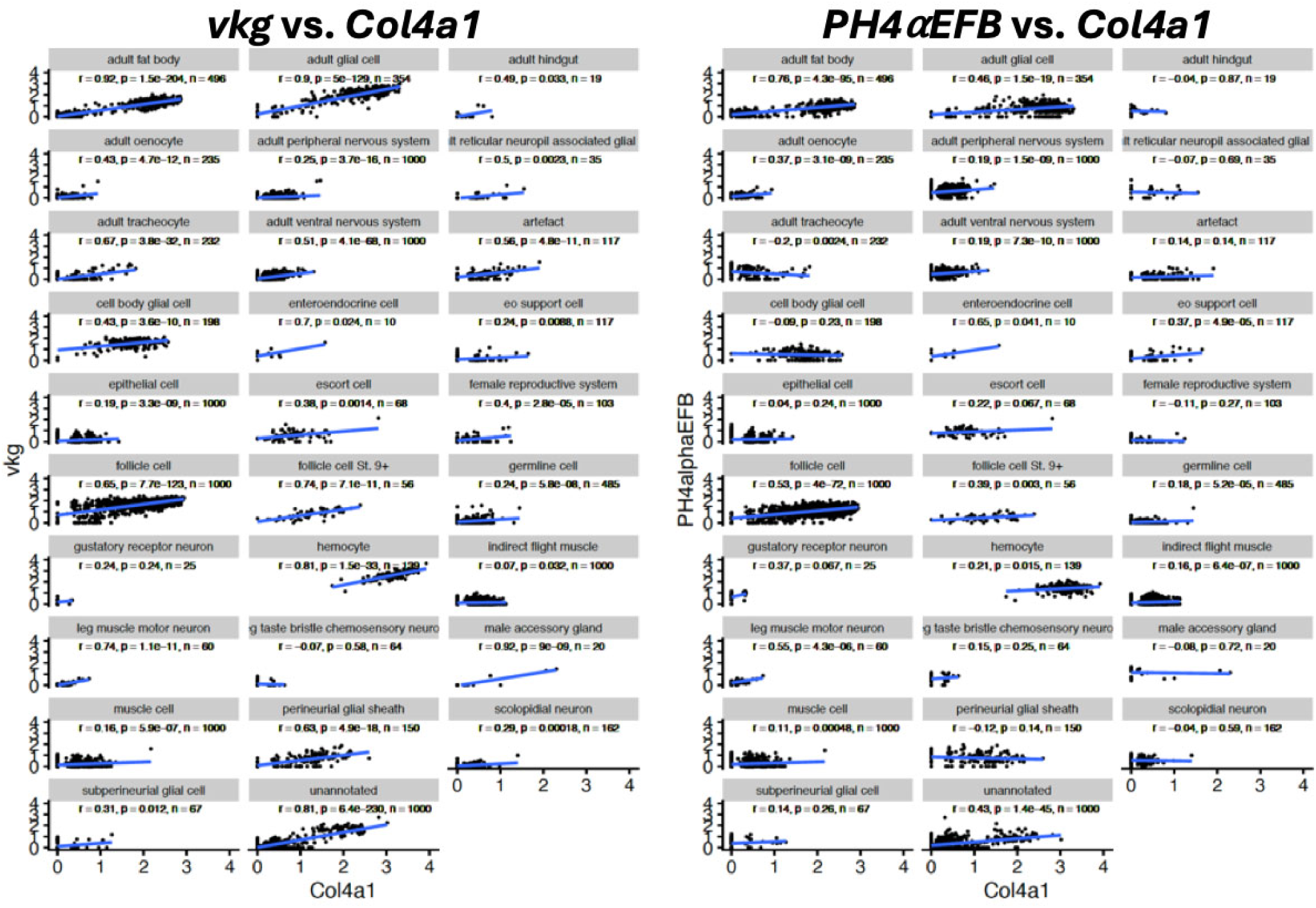
Expression of *Col4a1*, *vkg,* or *PH4*α*EFB* separately analysed for each different cell type in the whole-body dataset. Regression lines of the plots, correlation coefficients between the two genes plotted (*r*), the *p* values to obtain the results from the null hypothesis that the slope of the regression lines are zero, and the numbers of metacells analysed (n) are shown.

**Figure S5.**
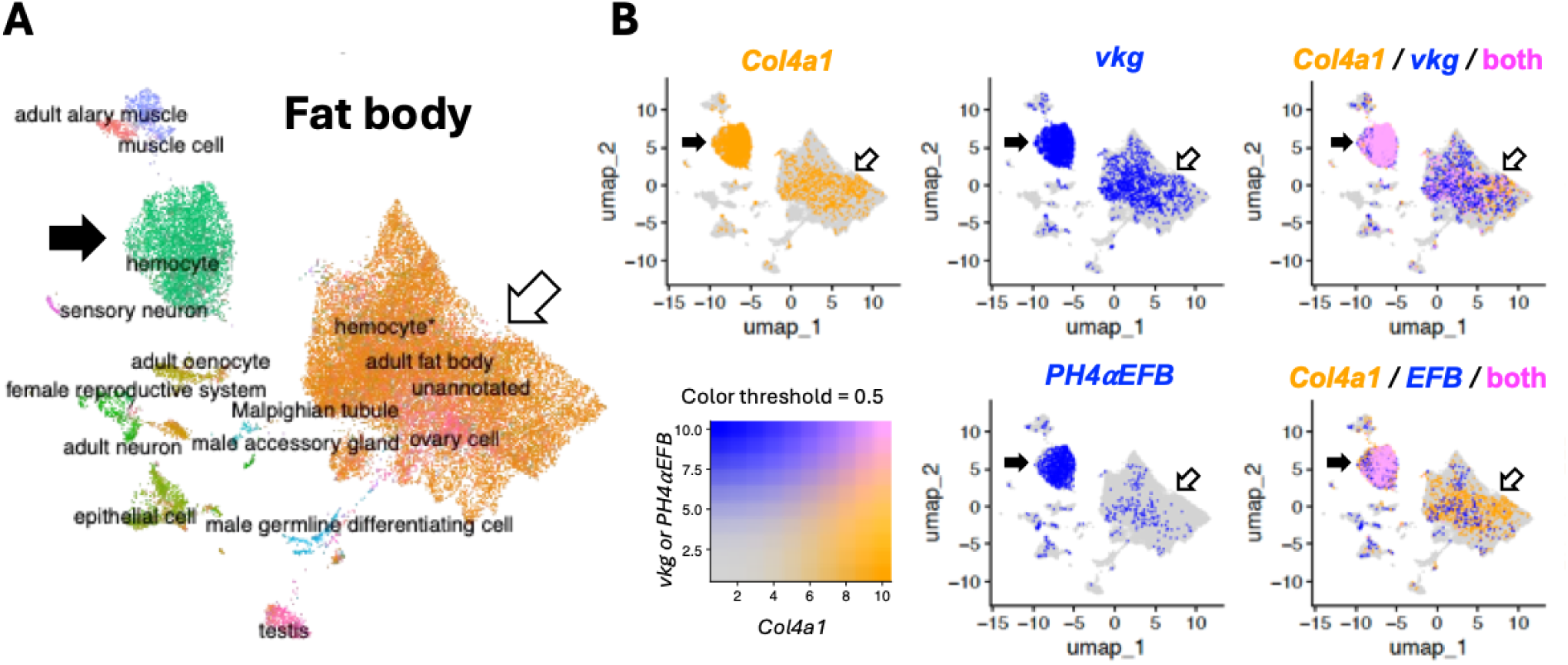
UMAP for the fat body single cell data. **(A)** UMAP showing the entire cells with annotations, with different cell types coded by different colour. **(B)** Expression of *Col4a1*, *vkg,* and *PH4*⍺*EFB (EFB)* colour coded as in the bottom left panel. Top left and middle panels show single gene expression; right panels show overlap. Closed arrows, hemocytes in which the three genes are co-expressed; open arrows, adult fat body cells in which the co-expression of the collagen IV genes and *PH4*⍺*EFB* is not as clear as that in the metacell data in Fig. 3B.

**Figure S6.**
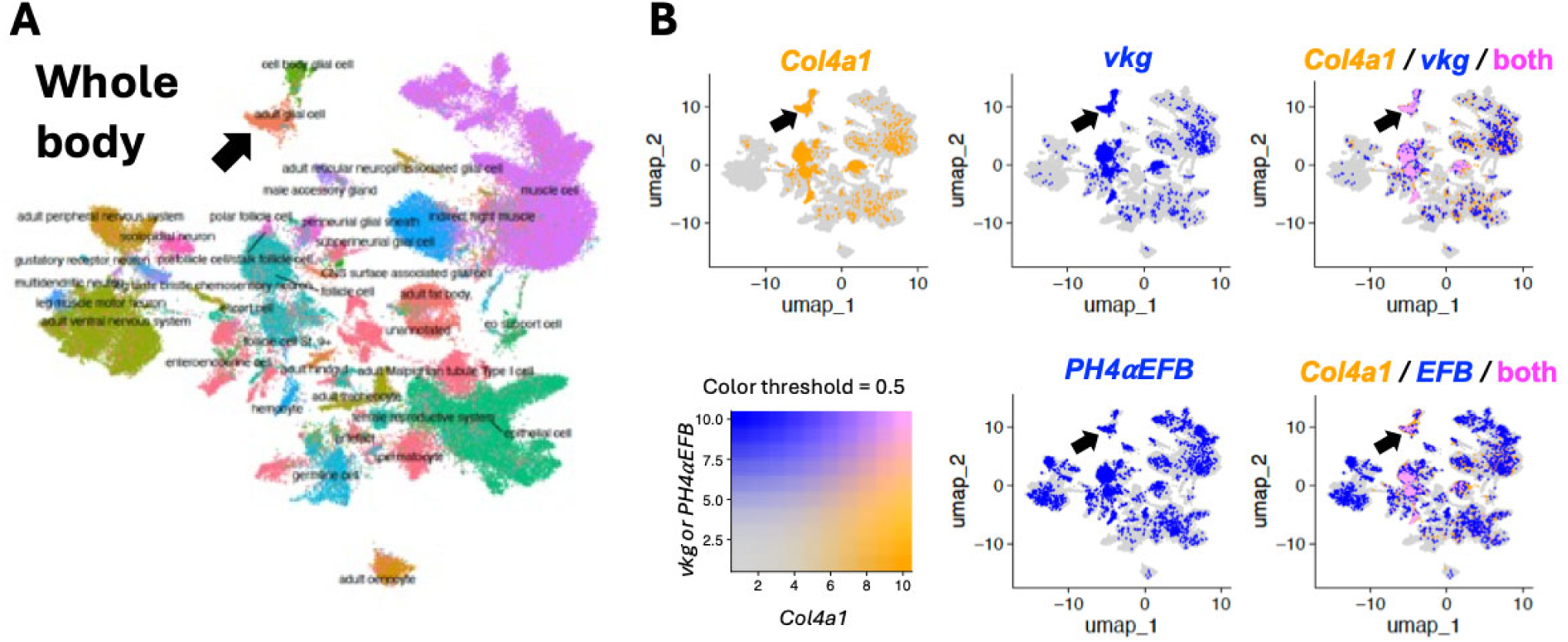
UMAP for the whole-body single cell data. **(A)** UMAP showing the entire cells with annotations, with different cell types coded by different colour. **(B)** Expression of *Col4a1*, *vkg,* and *PH4*⍺*EFB (EFB)* colour coded as in the bottom left panel. Top left and middle panels show single gene expression; right panels show overlap. Closed arrows, glial cells in which the three genes are co-expressed.

