## Supplementary Source Data for "Evidence for the major role of PH4αEFB in the prolyl 4-hydroxylation of *Drosophila* collagen IV"

### Mascot Search Results

PFHR9

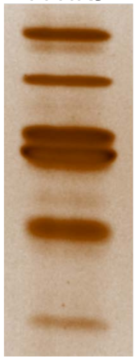

1

User :
Email :
Search title :
MS data file : ZIEN-29Nov2017_1m.mgf
Database : NCBInr 20120419 (17893860 sequences; 6141683785 residues)
Taxonomy : Mus musculus (house mouse) (138831 sequences)
Timestamp : 30 Nov 2017 at 20:13:50 GMT
Enzyme : Trypsin
Variable modifications : Carbamidomethyl (C),Oxidation (M),Oxidation (P)
Mass values : Monoisotopic
Protein Mass : Unrestricted
Peptide Mass Tolerance : ± 1.25 Da
Fragment Mass Tolerance: ± 1.001 Da
Max Missed Cleavages : 1
Instrument type : Default
Number of queries : 16088
Protein hits : gi|23271416 Leprecan 1 [Mus musculus]
gi|5852295 lysyl hydroxylase isoform 2 [Mus musculus]
gi|387397 epidermal keratin subunit I, partial [Mus musculus]
gi|6755106 procollagen-lysine,2-oxoglutarate 5-dioxygenase 1 precursor [Mus musculus]
gi|26345382 unnamed protein product [Mus musculus]
gi|4159806 type II keratin subunit protein [Mus musculus]
gi|1389682 plakoglobin, partial [Mus musculus]
gi|387398 epidermal keratin type I, partial [Mus musculus]
gi|148708988 mCG20427 [Mus musculus]
gi|12859782 unnamed protein product [Mus musculus]
gi|16303309 type II keratin 5 [Mus musculus]
gi|9790161 plakophilin-1 [Mus musculus]
gi|47523977 keratin, type II cytoskeletal 72 [Mus musculus]
gi|47059013 keratin, type II cytoskeletal 73 [Mus musculus]
gi|836898 prolyl 4-hydroxylase alpha(I)-subunit, partial [Mus musculus]
gi|220474 lamin A [Mus musculus]
gi|52785 unnamed protein product [Mus musculus]
gi|6677775 60S ribosomal protein L22 [Mus musculus]
gi|293686 epidermal keratin subunit II [Mus musculus]
gi|14917005 RecName: Full=Stress-70 protein, mitochondrial; AltName: Full=75 kDa glucose-regulated protein; Short=GRP-75; AltName: Full=Heat shock 70 kDa protein 9; AltName: Full=
gi|387207 heat shock protein [Mus musculus]
gi|50797 unnamed protein product [Mus musculus]
gi|6755110 procollagen-lysine,2-oxoglutarate 5-dioxygenase 3 precursor [Mus musculus]
gi|817996 class I (Qa) Q2k antigen [Mus musculus]
gi|19923857 thymidine phosphorylase [Mus musculus]
gi|51452 unnamed protein product [Mus musculus]
gi|398168 keratin 2 epidermis [Mus musculus]
gi|15277319 leucine zipper transcription factor-like protein 1 [Mus musculus]
gi|200765 ribonucleotide reductase subunit M1 [Mus musculus]
gi|191765 alpha-fetoprotein, partial [Mus musculus]
gi|18480346 olfactory receptor MOR175-2 [Mus musculus]
gi|226005 protein 40kD
gi|193645 glucose-regulated protein 78, partial [Mus musculus]
gi|6996913 annexin A2 [Mus musculus]
gi|116283284 Por protein [Mus musculus]
gi|12805465 Dnajc10 protein, partial [Mus musculus]
gi|148678782 mCG1027198, isoform CRA_a [Mus musculus]
gi|16716569 protease, serine, 1 precursor [Mus musculus]
gi|19924302 ankyrin repeat domain-containing protein 6 [Mus musculus]
gi|148680049 mCG147414 [Mus musculus]
gi|26337475 unnamed protein product [Mus musculus]
gi|199938 tumor-specific myb protein [Mus musculus]
gi|22094097 ganglioside-induced differentiation-associated protein 2 [Mus musculus]
gi|198683 unnamed protein product [Mus musculus]
gi|26327761 unnamed protein product [Mus musculus]
gi|51092301 type II keratin Kb14 [Mus musculus]
gi|74208129 unnamed protein product [Mus musculus]
gi|148685252 mCG142044 [Mus musculus]
gi|148671989 mCG1027581 [Mus musculus]
gi|26349347 unnamed protein product [Mus musculus]
gi|6755893 trypsin 4 precursor [Mus musculus]
gi|6754976 peroxiredoxin-1 [Mus musculus]

### Select Summary Report

Format As | Select Summary (protein hits) | Help

Significance threshold p< 0.05 | Max. number of hits AUTO

Standard scoring ○ MudPIT scoring ◉ Ions score or expect cut-off 43 | Show sub-sets 0

Show pop-ups ◉ Suppress pop-ups ○ | Require bold red ☐

Re-Search ◉ All queries ○ Unassigned ○ Below homology threshold ○ Below identity threshold

1. gi|23271416 **Mass:** 83540 **Score:** 243 **Matches:** 11(10) **Sequences:** 8(8) **emPAI:** 0.47
Leprecan 1 [Mus musculus]

| Query | Observed | Mr(expt) | Mr(calc) | Delta | Miss | Score | Expect | Rank | Unique | Peptide |
|---|---|---|---|---|---|---|---|---|---|---|
| 11282 | 1089.6395 | 1088.6322 | 1088.5601 | 0.0721 | 0 | 44 | 0.056 | 1 | U | K.EIETLVEEK.T |
| 11786 | 567.4809 | 1132.9471 | 1132.5360 | 0.4112 | 0 | 50 | 0.0094 | 1 | U | R.TAIEESQAER.K |
| 13314 | 644.9944 | 1287.9741 | 1287.6241 | 0.3500 | 0 | 60 | 0.00092 | 1 | U | K.TVTAEVQPQCGR.A |
| 14077 | 686.7367 | 1371.4588 | 1371.6605 | -0.2017 | 0 | 61 | 0.0017 | 1 | U | R.GDWPGVVLNMER.A 14085 |
| 14212 | 692.1389 | 1382.2633 | 1381.7677 | 0.4955 | 1 | 49 | 0.02 | 1 | U | K.LGQEGKVPLQSAR.M |
| 15268 | 743.7113 | 1485.4080 | 1485.7212 | -0.3132 | 0 | 89 | 2.6e-006 | 1 | U | R.AVGFSSGTENPHGVK.A |
| 15660 | 773.9469 | 1545.8792 | 1545.7133 | 0.1659 | 0 | 61 | 0.0018 | 1 | U | R.QNLDYYQTMSGVK.E |
| 15748 | 781.8881 | 1561.7616 | 1561.7083 | 0.0533 | 0 | (55) | 0.0066 | 1 | U | R.QNLDYYQTMSGVK.E |
| 14531 | 705.4221 | 2113.2444 | 2113.9826 | -0.7382 | 0 | 68 | 0.00038 | 1 | U | K.AVAAAHTFFVGNPEHMEMR.Q |
| 14649 | 710.9669 | 2129.8789 | 2129.9775 | -0.0986 | 0 | (45) | 0.068 | 1 | U | K.AVAAAHTFFVGNPEHMEMR.Q |

**Proteins matching the same set of peptides:**
gi|26326437 **Mass:** 83476 **Score:** 243 **Matches:** 11(10) **Sequences:** 8(8)
unnamed protein product [Mus musculus]
gi|74205527 **Mass:** 84439 **Score:** 243 **Matches:** 11(10) **Sequences:** 8(8)
unnamed protein product [Mus musculus]
gi|109150433 **Mass:** 83598 **Score:** 243 **Matches:** 11(10) **Sequences:** 8(8)
prolyl 3-hydroxylase 1 isoform 2 precursor [Mus musculus]
gi|109150437 **Mass:** 84426 **Score:** 243 **Matches:** 11(10) **Sequences:** 8(8)
prolyl 3-hydroxylase 1 isoform 1 precursor [Mus musculus]
gi|148698509 **Mass:** 84378 **Score:** 243 **Matches:** 11(10) **Sequences:** 8(8)
leprecan 1, isoform CRA_a [Mus musculus]
gi|148698510 **Mass:** 83550 **Score:** 243 **Matches:** 11(10) **Sequences:** 8(8)
leprecan 1, isoform CRA_b [Mus musculus]
gi|12846125 **Mass:** 84033 **Score:** 243 **Matches:** 11(10) **Sequences:** 8(8)
unnamed protein product [Mus musculus]

---

2. gi|5852295 **Mass:** 84473 **Score:** 198 **Matches:** 9(9) **Sequences:** 7(7) **emPAI:** 0.31
lysyl hydroxylase isoform 2 [Mus musculus]

| Query | Observed | Mr(expt) | Mr(calc) | Delta | Miss | Score | Expect | Rank | Unique | Peptide |
|---|---|---|---|---|---|---|---|---|---|---|
| 10580 | 1017.5937 | 1016.5864 | 1016.5906 | -0.0042 | 0 | 46 | 0.035 | 1 | U | R.EFIAPVTLK.V |
| 14359 | 697.9323 | 1393.8501 | 1393.6474 | 0.2027 | 0 | 77 | 2.2e-005 | 1 | U | R.SEDYVDIVQGNR.V 14365 |
| 14882 | 723.4815 | 1444.9483 | 1443.7762 | 1.1721 | 0 | 52 | 0.012 | 1 | U | K.IVFAADGLLWPDK.R |
| 15863 | 800.6402 | 1599.2658 | 1599.8773 | -0.6115 | 1 | 81 | 1.5e-005 | 1 | U | K.IVFAADGLLWPDKR.L 15868 |
| 15907 | 809.6948 | 1617.3751 | 1616.8774 | 0.4977 | 0 | 55 | 0.0063 | 1 | U | K.IFQALNGATDEVVLK.F |
| 15920 | 813.1936 | 1624.3726 | 1623.8885 | 0.4841 | 0 | 46 | 0.039 | 1 | U | K.QIGLENVWLHFIR.E |
| 15034 | 731.9530 | 2192.8372 | 2192.1477 | 0.6895 | 1 | 58 | 0.0041 | 1 | U | K.IFQALNGATDEVVLKFENGK.S |

**Proteins matching the same set of peptides:**
gi|18204027 **Mass:** 84389 **Score:** 198 **Matches:** 9(9) **Sequences:** 7(7)
Procollagen lysine, 2-oxoglutarate 5-dioxygenase 2 [Mus musculus]
gi|148688975 **Mass:** 84401 **Score:** 198 **Matches:** 9(9) **Sequences:** 7(7)
procollagen lysine, 2-oxoglutarate 5-dioxygenase 2, isoform CRA_a [Mus musculus]
gi|148688976 **Mass:** 86812 **Score:** 198 **Matches:** 9(9) **Sequences:** 7(7)

procollagen lysine, 2-oxoglutarate 5-dioxygenase 2, isoform CRA_b [Mus musculus]
gi|218931165 **Mass:** 84435 **Score:** 198 **Matches:** 9(9) **Sequences:** 7(7)
procollagen-lysine,2-oxoglutarate 5-dioxygenase 2 isoform 2 precursor [Mus musculus]
gi|218931167 **Mass:** 86846 **Score:** 198 **Matches:** 9(9) **Sequences:** 7(7)
procollagen-lysine,2-oxoglutarate 5-dioxygenase 2 isoform 1 precursor [Mus musculus]

---

**3.** gi|387397 **Mass:** 57807 **Score:** 143 **Matches:** 3(3) **Sequences:** 2(2) **emPAI:** 0.12
epidermal keratin subunit I, partial [Mus musculus]

| Query | Observed | Mr(expt) | Mr(calc) | Delta | Miss | Score | Expect | Rank | Unique | Peptide |
|---|---|---|---|---|---|---|---|---|---|---|
| 14204 | 691.6060 | 1381.1974 | 1380.6408 | 0.5566 | 0 | 74 | 3e-005 | 1 | U | R.ALEESNYELEGK.I 14193 |
| 14318 | 695.9275 | 1389.8404 | 1389.6736 | 0.1669 | 0 | 78 | 4.2e-005 | 1 | U | K.QSLEASLAETEGR.Y |

**Proteins matching the same set of peptides:**
gi|12852157 **Mass:** 58587 **Score:** 143 **Matches:** 3(3) **Sequences:** 2(2)
unnamed protein product [Mus musculus]
gi|26349141 **Mass:** 52653 **Score:** 143 **Matches:** 3(3) **Sequences:** 2(2)
unnamed protein product [Mus musculus]
gi|26349459 **Mass:** 49471 **Score:** 143 **Matches:** 3(3) **Sequences:** 2(2)
unnamed protein product [Mus musculus]
gi|112983636 **Mass:** 57007 **Score:** 143 **Matches:** 3(3) **Sequences:** 2(2)
keratin, type I cytoskeletal 10 [Mus musculus]
gi|116242600 **Mass:** 57735 **Score:** 143 **Matches:** 3(3) **Sequences:** 2(2)
RecName: Full=Keratin, type I cytoskeletal 10; AltName: Full=56 kDa cytokeratin; AltName: Full=Cytokeratin-10; Short=CK-10; AltName: Full=Keratin, type I cytoskeletal 59 kDa; AltName: Full=Keratin-1(

---

**4.** gi|6755106 **Mass:** 83542 **Score:** 105 **Matches:** 2(1) **Sequences:** 2(1) **emPAI:** 0.08
procollagen-lysine,2-oxoglutarate 5-dioxygenase 1 precursor [Mus musculus]

| Query | Observed | Mr(expt) | Mr(calc) | Delta | Miss | Score | Expect | Rank | Unique | Peptide |
|---|---|---|---|---|---|---|---|---|---|---|
| 14836 | 721.0568 | 1440.0991 | 1439.7370 | 0.3621 | 0 | 45 | 0.068 | 1 | U | K.FLVEYIAPMTEK.L |
| 15866 | 801.0905 | 1600.1665 | 1599.8297 | 0.3368 | 0 | 101 | 1.6e-007 | 1 | U | R.FLGSGGFIGYAPSLSK.L |

**Proteins matching the same set of peptides:**
gi|74207958 **Mass:** 83514 **Score:** 105 **Matches:** 2(1) **Sequences:** 2(1)
unnamed protein product [Mus musculus]

---

**5.** gi|26345382 **Mass:** 46296 **Score:** 89 **Matches:** 2(1) **Sequences:** 2(1) **emPAI:** 0.15
unnamed protein product [Mus musculus]

| Query | Observed | Mr(expt) | Mr(calc) | Delta | Miss | Score | Expect | Rank | Unique | Peptide |
|---|---|---|---|---|---|---|---|---|---|---|
| 15662 | 774.1068 | 1546.1991 | 1545.7246 | 0.4745 | 0 | 87 | 4.4e-006 | 1 | U | R.MISFSSGGENPHGVK.A |
| 15901 | 807.7070 | 2420.0993 | 2420.1437 | -0.0444 | 0 | 45 | 0.066 | 1 | U | K.LAAEGLGFSYAEPNYWISYGGR.Q |

**Proteins matching the same set of peptides:**
gi|148665269 **Mass:** 61664 **Score:** 89 **Matches:** 2(1) **Sequences:** 2(1)
leprecan-like 1 [Mus musculus]
gi|27734076 **Mass:** 80104 **Score:** 89 **Matches:** 2(1) **Sequences:** 2(1)
prolyl 3-hydroxylase 2 precursor [Mus musculus]

---

**6.** gi|4159806 **Mass:** 65183 **Score:** 87 **Matches:** 5(5) **Sequences:** 3(3) **emPAI:** 0.16
type II keratin subunit protein [Mus musculus]

| Query | Observed | Mr(expt) | Mr(calc) | Delta | Miss | Score | Expect | Rank | Unique | Peptide |
|---|---|---|---|---|---|---|---|---|---|---|
| 11211 | 542.0804 | 1082.1462 | 1081.5920 | 0.5541 | 1 | 49 | 0.011 | 1 | | K.FASFIDKVR.F |
| 14216 | 692.2543 | 1382.4940 | 1383.7034 | -1.2094 | 1 | 47 | 0.042 | 1 | | K.SLNDKFASFIDK.V 14237 14244 |
| 15155 | 738.6367 | 1475.2588 | 1474.8144 | 0.4444 | 1 | 61 | 0.0015 | 2 | U | R.FLEQQNKVLQTK.W |

---

**7.** gi|1389682 **Mass:** 68068 **Score:** 87 **Matches:** 2(2) **Sequences:** 2(2) **emPAI:** 0.10
plakoglobin, partial [Mus musculus]

| Query | Observed | Mr(expt) | Mr(calc) | Delta | Miss | Score | Expect | Rank | Unique | Peptide |
|---|---|---|---|---|---|---|---|---|---|---|
| 14556 | 706.2423 | 1410.4699 | 1410.8017 | -0.3317 | 0 | 71 | 0.00016 | 1 | U | R.ALMGSPQLVAAVVR.T |
| 14926 | 725.5305 | 1449.0465 | 1448.8140 | 0.2325 | 0 | 61 | 0.0013 | 1 | U | K.LLNQPNQWPLVK.A |

**Proteins matching the same set of peptides:**
gi|28395018 **Mass:** 81749 **Score:** 87 **Matches:** 2(2) **Sequences:** 2(2)
junction plakoglobin [Mus musculus]

---

**8.** gi|387398 **Mass:** 41454 **Score:** 87 **Matches:** 2(2) **Sequences:** 2(2) **emPAI:** 0.17
epidermal keratin type I, partial [Mus musculus]

| Query | Observed | Mr(expt) | Mr(calc) | Delta | Miss | Score | Expect | Rank | Unique | Peptide |
|---|---|---|---|---|---|---|---|---|---|---|
| 10673 | **514.9652** | **1027.9159** | **1028.5866** | **-0.6707** | **0** | **54** | **0.0037** | **1** | **U** | **R.VLDELTLAR.A** |
| 13972 | **681.9645** | **1361.9144** | **1360.6834** | **1.2310** | **0** | **69** | **0.00028** | **1** | **U** | **R.EVATNSELVQSGK.S** |

**Proteins matching the same set of peptides:**
gi|7106335 **Mass:** 48132 **Score:** 87 **Matches:** 2(2) **Sequences:** 2(2)
keratin, type I cytoskeletal 17 [Mus musculus]
gi|13097093 **Mass:** 30629 **Score:** 87 **Matches:** 2(2) **Sequences:** 2(2)
Krt14 protein [Mus musculus]
gi|21489935 **Mass:** 52834 **Score:** 87 **Matches:** 2(2) **Sequences:** 2(2)
keratin, type I cytoskeletal 14 [Mus musculus]
gi|148670626 **Mass:** 54668 **Score:** 87 **Matches:** 2(2) **Sequences:** 2(2)
mCG144006 [Mus musculus]

---

**9.** gi|148708988 **Mass:** 320241 **Score:** 85 **Matches:** 2(2) **Sequences:** 2(2) **emPAI:** 0.02
mCG20427 [Mus musculus]

| Query | Observed | Mr(expt) | Mr(calc) | Delta | Miss | Score | Expect | Rank | Unique | Peptide |
|---|---|---|---|---|---|---|---|---|---|---|
| 11741 | **565.5758** | **1129.1370** | **1128.6026** | **0.5344** | **0** | **59** | **0.001** | **1** | **U** | **K.IEVLEEELR.L** |
| 14281 | **694.6082** | **1387.2019** | **1386.7103** | **0.4916** | **0** | **68** | **0.00033** | **1** | **U** | **R.LNDSILQATEQR.R** |

**Proteins matching the same set of peptides:**
gi|190194418 **Mass:** 332706 **Score:** 85 **Matches:** 2(2) **Sequences:** 2(2)
desmoplakin [Mus musculus]

---

**10.** gi|12859782 **Mass:** 65586 **Score:** 84 **Matches:** 5(5) **Sequences:** 3(3) **emPAI:** 0.16
unnamed protein product [Mus musculus]

| Query | Observed | Mr(expt) | Mr(calc) | Delta | Miss | Score | Expect | Rank | Unique | Peptide |
|---|---|---|---|---|---|---|---|---|---|---|
| 11211 | 542.0804 | 1082.1462 | 1081.5920 | 0.5541 | 1 | 49 | 0.011 | 1 | | K.FASFIDKVR.F |
| 14216 | 692.2543 | 1382.4940 | 1383.7034 | -1.2094 | 1 | 47 | 0.042 | 1 | | K.SLNDKFASFIDK.V 14237 14244 |
| 15155 | 738.6367 | 1475.2588 | 1474.7780 | 0.4808 | 0 | 58 | 0.0034 | 3 | | R.FLEQQNQVLQTK.W |

**Proteins matching the same set of peptides:**
gi|126116585 **Mass:** 65565 **Score:** 84 **Matches:** 5(5) **Sequences:** 3(3)
keratin, type II cytoskeletal 1 [Mus musculus]

---

**11.** gi|16303309 **Mass:** 61743 **Score:** 84 **Matches:** 3(3) **Sequences:** 2(2) **emPAI:** 0.17
type II keratin 5 [Mus musculus]

| Query | Observed | Mr(expt) | Mr(calc) | Delta | Miss | Score | Expect | Rank | Unique | Peptide |
|---|---|---|---|---|---|---|---|---|---|---|
| 11211 | 542.0804 | 1082.1462 | 1081.5920 | 0.5541 | 1 | 49 | 0.011 | 1 | | K.FASFIDKVR.F |
| 14704 | **713.6212** | **1425.2279** | **1424.6680** | **0.5599** | **0** | **69** | **0.00024** | **1** | **U** | **R.VDALMDEINFMK.M** |
| 14999 | **729.5875** | **1457.1605** | **1456.6578** | **0.5027** | **0** | **(49)** | **0.024** | **1** | **U** | **R.VDALMDEINFMK.M** |

**Proteins matching the same set of peptides:**
gi|20911031 **Mass:** 61729 **Score:** 84 **Matches:** 3(3) **Sequences:** 2(2)
keratin, type II cytoskeletal 5 [Mus musculus]

---

**12.** gi|9790161 **Mass:** 80844 **Score:** 80 **Matches:** 1(1) **Sequences:** 1(1) **emPAI:** 0.04
plakophilin-1 [Mus musculus]

| Query | Observed | Mr(expt) | Mr(calc) | Delta | Miss | Score | Expect | Rank | Unique | Peptide |
|---|---|---|---|---|---|---|---|---|---|---|

| | | | | | | | | | | |
|---|---|---|---|---|---|---|---|---|---|---|
| 15543 | 762.9679 | 1523.9212 | 1523.7804 | 0.1408 | 0 | 80 | 1.4e-005 | 1 | U | R.SPNQNVQQAAAGALR.N |

**Proteins matching the same set of peptides:**
gi|148707614 **Mass:** 78853 **Score:** 80 **Matches:** 1(1) **Sequences:** 1(1)
plakophilin 1 [Mus musculus]

---

**13.** gi|47523977 **Mass:** 56715 **Score:** 71 **Matches:** 2(2) **Sequences:** 2(2) **emPAI:** 0.12
keratin, type II cytoskeletal 72 [Mus musculus]

| Query | Observed | Mr(expt) | Mr(calc) | Delta | Miss | Score | Expect | Rank | Unique | Peptide |
|---|---|---|---|---|---|---|---|---|---|---|
| 11211 | 542.0804 | 1082.1462 | 1081.5920 | 0.5541 | 1 | 49 | 0.011 | 1 | | K.FASFIDKVR.F |
| 15155 | 738.6367 | 1475.2588 | 1475.7620 | -0.5032 | 0 | 66 | 0.0005 | 1 | U | R.FLEQQNQVLETK.W |

---

**14.** gi|47059013 **Mass:** 58875 **Score:** 65 **Matches:** 3(2) **Sequences:** 3(2) **emPAI:** 0.18
keratin, type II cytoskeletal 73 [Mus musculus]

| Query | Observed | Mr(expt) | Mr(calc) | Delta | Miss | Score | Expect | Rank | Unique | Peptide |
|---|---|---|---|---|---|---|---|---|---|---|
| 11211 | 542.0804 | 1082.1462 | 1081.5920 | 0.5541 | 1 | 49 | 0.011 | 1 | | K.FASFIDKVR.F |
| 13933 | 679.6329 | 1357.2512 | 1356.7249 | 0.5263 | 0 | 43 | 0.1 | 1 | U | R.NLDLDSIIAEVR.A |
| 15155 | 738.6367 | 1475.2588 | 1474.7780 | 0.4808 | 0 | 58 | 0.0034 | 3 | | R.FLEQQNQVLQTK.W |

---

**15.** gi|836898 **Mass:** 59960 **Score:** 64 **Matches:** 1(1) **Sequences:** 1(1) **emPAI:** 0.05
prolyl 4-hydroxylase alpha(I)-subunit, partial [Mus musculus]

| Query | Observed | Mr(expt) | Mr(calc) | Delta | Miss | Score | Expect | Rank | Unique | Peptide |
|---|---|---|---|---|---|---|---|---|---|---|
| 15762 | 783.5424 | 1565.0702 | 1564.7522 | 0.3180 | 0 | 64 | 0.00069 | 1 | U | K.SAWLSGYEDPVVSR.I |

**Proteins matching the same set of peptides:**
gi|26336999 **Mass:** 60848 **Score:** 64 **Matches:** 1(1) **Sequences:** 1(1)
unnamed protein product [Mus musculus]
gi|33859596 **Mass:** 60872 **Score:** 64 **Matches:** 1(1) **Sequences:** 1(1)
prolyl 4-hydroxylase subunit alpha-1 precursor [Mus musculus]
gi|74148153 **Mass:** 51722 **Score:** 64 **Matches:** 1(1) **Sequences:** 1(1)
unnamed protein product [Mus musculus]
gi|74224984 **Mass:** 60886 **Score:** 64 **Matches:** 1(1) **Sequences:** 1(1)
unnamed protein product [Mus musculus]
gi|74225936 **Mass:** 63769 **Score:** 64 **Matches:** 1(1) **Sequences:** 1(1)
unnamed protein product [Mus musculus]

---

**16.** gi|220474 **Mass:** 47506 **Score:** 62 **Matches:** 1(1) **Sequences:** 1(1) **emPAI:** 0.07
lamin A [Mus musculus]

| Query | Observed | Mr(expt) | Mr(calc) | Delta | Miss | Score | Expect | Rank | Unique | Peptide |
|---|---|---|---|---|---|---|---|---|---|---|
| 10676 | 515.0694 | 1028.1242 | 1027.5662 | 0.5581 | 0 | 62 | 0.00055 | 1 | U | R.LADALQELR.A |

**Proteins matching the same set of peptides:**
gi|52865 **Mass:** 65367 **Score:** 62 **Matches:** 1(1) **Sequences:** 1(1)
unnamed protein product [Mus musculus]
gi|9506843 **Mass:** 52620 **Score:** 62 **Matches:** 1(1) **Sequences:** 1(1)
prelamin-A/C isoform C2 [Mus musculus]
gi|12835914 **Mass:** 74255 **Score:** 62 **Matches:** 1(1) **Sequences:** 1(1)
unnamed protein product [Mus musculus]
gi|74148166 **Mass:** 74150 **Score:** 62 **Matches:** 1(1) **Sequences:** 1(1)
unnamed protein product [Mus musculus]
gi|74204231 **Mass:** 65451 **Score:** 62 **Matches:** 1(1) **Sequences:** 1(1)
unnamed protein product [Mus musculus]
gi|74219492 **Mass:** 65397 **Score:** 62 **Matches:** 1(1) **Sequences:** 1(1)
unnamed protein product [Mus musculus]
gi|74219697 **Mass:** 65335 **Score:** 62 **Matches:** 1(1) **Sequences:** 1(1)
unnamed protein product [Mus musculus]

gi|74220666 **Mass:** 72427 **Score:** 62 **Matches:** 1(1) **Sequences:** 1(1)
unnamed protein product [Mus musculus]
gi|74220835 **Mass:** 65398 **Score:** 62 **Matches:** 1(1) **Sequences:** 1(1)
unnamed protein product [Mus musculus]
gi|161760667 **Mass:** 65407 **Score:** 62 **Matches:** 1(1) **Sequences:** 1(1)
prelamin-A/C isoform C [Mus musculus]
gi|162287370 **Mass:** 74193 **Score:** 62 **Matches:** 1(1) **Sequences:** 1(1)
prelamin-A/C isoform A [Mus musculus]

---

**17.** gi|52785 **Mass:** 56409 **Score:** 60 **Matches:** 2(2) **Sequences:** 2(2) **emPAI:** 0.12
unnamed protein product [Mus musculus]

| Query | Observed | Mr(expt) | Mr(calc) | Delta | Miss | Score | Expect | Rank | Unique | Peptide |
|---|---|---|---|---|---|---|---|---|---|---|
| 11211 | 542.0804 | 1082.1462 | 1081.5920 | 0.5541 | 1 | 49 | 0.011 | 1 | | K.FASFIDKVR.F |
| 15155 | 738.6367 | 1475.2588 | 1475.7984 | -0.5396 | 1 | 55 | 0.0058 | 4 | U | R.FLEQQNKVLETK.W |

**Proteins matching the same set of peptides:**
gi|22164776 **Mass:** 57517 **Score:** 60 **Matches:** 2(2) **Sequences:** 2(2)
keratin, type II cytoskeletal 79 [Mus musculus]
gi|26324736 **Mass:** 58230 **Score:** 60 **Matches:** 2(2) **Sequences:** 2(2)
unnamed protein product [Mus musculus]
gi|29789317 **Mass:** 59704 **Score:** 60 **Matches:** 2(2) **Sequences:** 2(2)
keratin, type II cytoskeletal 75 [Mus musculus]
gi|39850082 **Mass:** 57817 **Score:** 60 **Matches:** 2(2) **Sequences:** 2(2)
Krt4 protein [Mus musculus]
gi|85701680 **Mass:** 62806 **Score:** 60 **Matches:** 2(2) **Sequences:** 2(2)
keratin, type II cytoskeletal 2 oral [Mus musculus]
gi|109735018 **Mass:** 58204 **Score:** 60 **Matches:** 2(2) **Sequences:** 2(2)
RIKEN cDNA 4732456N10 gene [Mus musculus]
gi|133778953 **Mass:** 56248 **Score:** 60 **Matches:** 2(2) **Sequences:** 2(2)
keratin, type II cytoskeletal 4 [Mus musculus]
gi|148672069 **Mass:** 59231 **Score:** 60 **Matches:** 2(2) **Sequences:** 2(2)
cDNA sequence BC031593 [Mus musculus]
gi|148672071 **Mass:** 57541 **Score:** 60 **Matches:** 2(2) **Sequences:** 2(2)
keratin 4, isoform CRA_b [Mus musculus]
gi|148672072 **Mass:** 63519 **Score:** 60 **Matches:** 2(2) **Sequences:** 2(2)
mCG144546 [Mus musculus]
gi|148672089 **Mass:** 62241 **Score:** 60 **Matches:** 2(2) **Sequences:** 2(2)
keratin 75 [Mus musculus]
gi|148672090 **Mass:** 58890 **Score:** 60 **Matches:** 2(2) **Sequences:** 2(2)
RIKEN cDNA 4732456N10 [Mus musculus]
gi|269914154 **Mass:** 58188 **Score:** 60 **Matches:** 2(2) **Sequences:** 2(2)
uncharacterized protein LOC239673 [Mus musculus]

---

**18.** gi|6677775 **Mass:** 14750 **Score:** 59 **Matches:** 1(1) **Sequences:** 1(1) **emPAI:** 0.23
60S ribosomal protein L22 [Mus musculus]

| Query | Observed | Mr(expt) | Mr(calc) | Delta | Miss | Score | Expect | Rank | Unique | Peptide |
|---|---|---|---|---|---|---|---|---|---|---|
| 12933 | **622.0134** | **1242.0122** | **1241.6728** | **0.3394** | **0** | **59** | **0.001** | **1** | **U** | **K.AGNLGGGVVTIER.S** |

---

**19.** gi|293686 **Mass:** 59414 **Score:** 58 **Matches:** 2(2) **Sequences:** 2(2) **emPAI:** 0.11
epidermal keratin subunit II [Mus musculus]

| Query | Observed | Mr(expt) | Mr(calc) | Delta | Miss | Score | Expect | Rank | Unique | Peptide |
|---|---|---|---|---|---|---|---|---|---|---|
| 11211 | 542.0804 | 1082.1462 | 1081.5920 | 0.5541 | 1 | 49 | 0.011 | 1 | | K.FASFIDKVR.F |
| 13133 | **634.0348** | **1266.0550** | **1265.6139** | **0.4411** | **0** | **50** | **0.021** | **1** | **U** | **R.TDAENEFVTLK.K** |

---

**20.** gi|14917005 **Mass:** 73483 **Score:** 58 **Matches:** 2(2) **Sequences:** 2(2) **emPAI:** 0.09
RecName: Full=Stress-70 protein, mitochondrial; AltName: Full=75 kDa glucose-regulated protein; Short=GRP-75; AltName: Full=Heat shock 70 kDa protein 9; AltName: Full=Mortalin; AltName: Full=Peptide

| Query | Observed | Mr(expt) | Mr(calc) | Delta | Miss | Score | Expect | Rank | Unique | Peptide |
|---|---|---|---|---|---|---|---|---|---|---|
| 12939 | **622.4032** | **1242.7918** | **1241.6728** | **1.1191** | **0** | **49** | **0.03** | **1** | **U** | **K.DAGQISGLNVLR.V** |

| | | | | | | | | | | |
|---|---|---|---|---|---|---|---|---|---|---|
| 14049 | 685.8440 | 2054.5101 | 2054.9545 | -0.4444 | 0 | 55 | 0.0067 | 1 | U | K.STNGDTFLGGEDFDQALLR.H |

**Proteins matching the same set of peptides:**
gi|74204605 **Mass:** 73446 **Score:** 58 **Matches:** 2(2) **Sequences:** 2(2)
unnamed protein product [Mus musculus]
gi|74205924 **Mass:** 63924 **Score:** 58 **Matches:** 2(2) **Sequences:** 2(2)
unnamed protein product [Mus musculus]
gi|74225724 **Mass:** 73431 **Score:** 58 **Matches:** 2(2) **Sequences:** 2(2)
unnamed protein product [Mus musculus]
gi|162461907 **Mass:** 73416 **Score:** 58 **Matches:** 2(2) **Sequences:** 2(2)
stress-70 protein, mitochondrial [Mus musculus]

---

**21.** gi|387207 **Mass:** 69698 **Score:** 57 **Matches:** 1(1) **Sequences:** 1(1) **emPAI:** 0.05
heat shock protein [Mus musculus]

| Query | Observed | Mr(expt) | Mr(calc) | Delta | Miss | Score | Expect | Rank | Unique | Peptide |
|---|---|---|---|---|---|---|---|---|---|---|
| 13034 | 627.9152 | 1253.8158 | 1252.6088 | 1.2070 | 0 | 57 | 0.0041 | 1 | U | R.FEELNADLFR.G |

**Proteins matching the same set of peptides:**
gi|31560686 **Mass:** 69599 **Score:** 57 **Matches:** 1(1) **Sequences:** 1(1)
heat shock-related 70 kDa protein 2 [Mus musculus]
gi|74142040 **Mass:** 70841 **Score:** 57 **Matches:** 1(1) **Sequences:** 1(1)
unnamed protein product [Mus musculus]
gi|74179642 **Mass:** 43517 **Score:** 57 **Matches:** 1(1) **Sequences:** 1(1)
unnamed protein product [Mus musculus]
gi|74214304 **Mass:** 56662 **Score:** 57 **Matches:** 1(1) **Sequences:** 1(1)
unnamed protein product [Mus musculus]
gi|77415383 **Mass:** 62117 **Score:** 57 **Matches:** 1(1) **Sequences:** 1(1)
Hspa8 protein, partial [Mus musculus]
gi|215260057 **Mass:** 69681 **Score:** 57 **Matches:** 1(1) **Sequences:** 1(1)
heat shock protein 70-2 [Mus musculus]
gi|309319 **Mass:** 70793 **Score:** 57 **Matches:** 1(1) **Sequences:** 1(1)
heat shock protein 70 cognate [Mus musculus]
gi|13242237 **Mass:** 70827 **Score:** 57 **Matches:** 1(1) **Sequences:** 1(1)
heat shock cognate 71 kDa protein [Rattus norvegicus]
gi|42542422 **Mass:** 70828 **Score:** 57 **Matches:** 1(1) **Sequences:** 1(1)
Heat shock protein 8 [Mus musculus]
gi|63101351 **Mass:** 68736 **Score:** 57 **Matches:** 1(1) **Sequences:** 1(1)
Hspa8 protein [Mus musculus]
gi|74142813 **Mass:** 50433 **Score:** 57 **Matches:** 1(1) **Sequences:** 1(1)
unnamed protein product [Mus musculus]
gi|74143862 **Mass:** 70813 **Score:** 57 **Matches:** 1(1) **Sequences:** 1(1)
unnamed protein product [Mus musculus]
gi|74181586 **Mass:** 70855 **Score:** 57 **Matches:** 1(1) **Sequences:** 1(1)
unnamed protein product [Mus musculus]
gi|74181633 **Mass:** 70841 **Score:** 57 **Matches:** 1(1) **Sequences:** 1(1)
unnamed protein product [Mus musculus]
gi|74184057 **Mass:** 70857 **Score:** 57 **Matches:** 1(1) **Sequences:** 1(1)
unnamed protein product [Mus musculus]
gi|74186087 **Mass:** 70739 **Score:** 57 **Matches:** 1(1) **Sequences:** 1(1)
unnamed protein product [Mus musculus]
gi|74190799 **Mass:** 70813 **Score:** 57 **Matches:** 1(1) **Sequences:** 1(1)
unnamed protein product [Mus musculus]
gi|74191381 **Mass:** 70859 **Score:** 57 **Matches:** 1(1) **Sequences:** 1(1)
unnamed protein product [Mus musculus]
gi|74198858 **Mass:** 70852 **Score:** 57 **Matches:** 1(1) **Sequences:** 1(1)
unnamed protein product [Mus musculus]
gi|74198978 **Mass:** 70903 **Score:** 57 **Matches:** 1(1) **Sequences:** 1(1)
unnamed protein product [Mus musculus]

gi|74208631 **Mass:** 70827 **Score:** 57 **Matches:** 1(1) **Sequences:** 1(1)
unnamed protein product [Mus musculus]
gi|74211333 **Mass:** 68074 **Score:** 57 **Matches:** 1(1) **Sequences:** 1(1)
unnamed protein product [Mus musculus]
gi|74211667 **Mass:** 60764 **Score:** 57 **Matches:** 1(1) **Sequences:** 1(1)
unnamed protein product [Mus musculus]
gi|74214176 **Mass:** 70828 **Score:** 57 **Matches:** 1(1) **Sequences:** 1(1)
unnamed protein product [Mus musculus]
gi|74220416 **Mass:** 70755 **Score:** 57 **Matches:** 1(1) **Sequences:** 1(1)
unnamed protein product [Mus musculus]
gi|74220592 **Mass:** 70648 **Score:** 57 **Matches:** 1(1) **Sequences:** 1(1)
unnamed protein product [Mus musculus]
gi|74225511 **Mass:** 70783 **Score:** 57 **Matches:** 1(1) **Sequences:** 1(1)
unnamed protein product [Mus musculus]
gi|148693577 **Mass:** 35174 **Score:** 57 **Matches:** 1(1) **Sequences:** 1(1)
mCG5074, isoform CRA_a [Mus musculus]
gi|215261181 **Mass:** 42317 **Score:** 57 **Matches:** 1(1) **Sequences:** 1(1)
Chain A, Chaperone Complex

---

**22.** gi|50797 **Mass:** 50018 **Score:** 55 **Matches:** 2(1) **Sequences:** 1(1) **emPAI:** 0.07
unnamed protein product [Mus musculus]

| Query | Observed | Mr(expt) | Mr(calc) | Delta | Miss | Score | Expect | Rank | Unique | Peptide |
|---|---|---|---|---|---|---|---|---|---|---|
| 10649 | 513.1775 | 1024.3404 | 1024.6030 | -0.2625 | 0 | 52 | 0.013 | 1 | U | K.IGGIGTVPVGR.V 10648 |

**Proteins matching the same set of peptides:**
gi|556301 **Mass:** 50132 **Score:** 55 **Matches:** 2(1) **Sequences:** 1(1)
elongation factor Tu [Mus musculus]
gi|6681273 **Mass:** 50422 **Score:** 55 **Matches:** 2(1) **Sequences:** 1(1)
elongation factor 1-alpha 2 [Mus musculus]
gi|13278382 **Mass:** 50140 **Score:** 55 **Matches:** 2(1) **Sequences:** 1(1)
Eukaryotic translation elongation factor 1 alpha 1 [Mus musculus]
gi|26328693 **Mass:** 50024 **Score:** 55 **Matches:** 2(1) **Sequences:** 1(1)
unnamed protein product [Mus musculus]
gi|26345590 **Mass:** 50086 **Score:** 55 **Matches:** 2(1) **Sequences:** 1(1)
unnamed protein product [Mus musculus]
gi|28460696 **Mass:** 50082 **Score:** 55 **Matches:** 2(1) **Sequences:** 1(1)
elongation factor 1-alpha 1 [Rattus norvegicus]
gi|74195737 **Mass:** 50081 **Score:** 55 **Matches:** 2(1) **Sequences:** 1(1)
unnamed protein product [Mus musculus]
gi|74204203 **Mass:** 50072 **Score:** 55 **Matches:** 2(1) **Sequences:** 1(1)
unnamed protein product [Mus musculus]
gi|74227478 **Mass:** 50034 **Score:** 55 **Matches:** 2(1) **Sequences:** 1(1)
unnamed protein product [Mus musculus]
gi|148694454 **Mass:** 38144 **Score:** 55 **Matches:** 2(1) **Sequences:** 1(1)
mCG15232, isoform CRA_b [Mus musculus]
gi|148694456 **Mass:** 32313 **Score:** 55 **Matches:** 2(1) **Sequences:** 1(1)
mCG15232, isoform CRA_d [Mus musculus]

---

**23.** gi|6755110 **Mass:** 84869 **Score:** 54 **Matches:** 1(1) **Sequences:** 1(1) **emPAI:** 0.04
procollagen-lysine,2-oxoglutarate 5-dioxygenase 3 precursor [Mus musculus]

| Query | Observed | Mr(expt) | Mr(calc) | Delta | Miss | Score | Expect | Rank | Unique | Peptide |
|---|---|---|---|---|---|---|---|---|---|---|
| 14416 | 700.0134 | 1398.0123 | 1397.7150 | 0.2973 | 0 | 54 | 0.0092 | 1 | U | K.LVGPEEALSAGEAR.D |

**Proteins matching the same set of peptides:**
gi|12850403 **Mass:** 84909 **Score:** 54 **Matches:** 1(1) **Sequences:** 1(1)
unnamed protein product [Mus musculus]

---

**24.** gi|817996 **Mass:** 40730 **Score:** 53 **Matches:** 1(1) **Sequences:** 1(1) **emPAI:** 0.08
class I (Qa) Q2k antigen [Mus musculus]

| Query | Observed | Mr(expt) | Mr(calc) | Delta | Miss | Score | Expect | Rank | Unique | Peptide |
|---|---|---|---|---|---|---|---|---|---|---|
| 11186 | 1081.2339 | 1080.2266 | 1079.4883 | 0.7383 | 0 | 53 | 0.0038 | 1 | U | R.FDSDAEKPR.Y |

**Proteins matching the same set of peptides:**

gi|148674754 **Mass:** 40746 **Score:** 53 **Matches:** 1(1) **Sequences:** 1(1)
mCG4996 [Mus musculus]

---

**25.** gi|19923857 **Mass:** 49305 **Score:** 52 **Matches:** 1(1) **Sequences:** 1(1) **emPAI:** 0.07
thymidine phosphorylase [Mus musculus]

| Query | Observed | Mr(expt) | Mr(calc) | Delta | Miss | Score | Expect | Rank | Unique | Peptide |
|---|---|---|---|---|---|---|---|---|---|---|
| 15416 | 754.5466 | 1507.0786 | 1507.7419 | -0.6634 | 1 | 52 | 0.0098 | 1 | U | K.FGGAAVFPDQEKAR.E |

---

**26.** gi|51452 **Mass:** 58833 **Score:** 52 **Matches:** 1(1) **Sequences:** 1(1) **emPAI:** 0.06
unnamed protein product [Mus musculus]

| Query | Observed | Mr(expt) | Mr(calc) | Delta | Miss | Score | Expect | Rank | Unique | Peptide |
|---|---|---|---|---|---|---|---|---|---|---|
| 15865 | 801.0338 | 1600.0531 | 1599.9018 | 0.1513 | 1 | 52 | 0.015 | 1 | U | R.RGVMLAVDAVIAELK.K |

**Proteins matching the same set of peptides:**

gi|148677951 **Mass:** 17541 **Score:** 52 **Matches:** 1(1) **Sequences:** 1(1)
mCG1033306 [Mus musculus]

gi|148700389 **Mass:** 49482 **Score:** 52 **Matches:** 1(1) **Sequences:** 1(1)
mCG116284 [Mus musculus]

gi|51455 **Mass:** 60903 **Score:** 52 **Matches:** 1(1) **Sequences:** 1(1)
heat shock protein 65 [Mus musculus]

gi|26353954 **Mass:** 60918 **Score:** 52 **Matches:** 1(1) **Sequences:** 1(1)
unnamed protein product [Mus musculus]

gi|76779273 **Mass:** 59388 **Score:** 52 **Matches:** 1(1) **Sequences:** 1(1)
Hspd1 protein, partial [Mus musculus]

gi|183396771 **Mass:** 60917 **Score:** 52 **Matches:** 1(1) **Sequences:** 1(1)
60 kDa heat shock protein, mitochondrial [Mus musculus]

---

**27.** gi|398168 **Mass:** 70934 **Score:** 52 **Matches:** 2(1) **Sequences:** 2(1) **emPAI:** 0.09
keratin 2 epidermis [Mus musculus]

| Query | Observed | Mr(expt) | Mr(calc) | Delta | Miss | Score | Expect | Rank | Unique | Peptide |
|---|---|---|---|---|---|---|---|---|---|---|
| 11211 | 542.0804 | 1082.1462 | 1081.5920 | 0.5541 | 1 | 49 | 0.011 | 1 | | K.FASFIDKVR.F |
| 13037 | 628.1775 | 1254.3404 | 1253.6001 | 0.7404 | 0 | 44 | 0.077 | 1 | U | R.GFSSGSAVVSGGSR.R |

**Proteins matching the same set of peptides:**

gi|111308159 **Mass:** 70880 **Score:** 52 **Matches:** 2(1) **Sequences:** 2(1)
Keratin 2 [Mus musculus]

gi|124487419 **Mass:** 70880 **Score:** 52 **Matches:** 2(1) **Sequences:** 2(1)
keratin, type II cytoskeletal 2 epidermal [Mus musculus]

gi|148672075 **Mass:** 71783 **Score:** 52 **Matches:** 2(1) **Sequences:** 2(1)
mCG17605, isoform CRA_a [Mus musculus]

gi|148672076 **Mass:** 72140 **Score:** 52 **Matches:** 2(1) **Sequences:** 2(1)
mCG17605, isoform CRA_b [Mus musculus]

---

**28.** gi|15277319 **Mass:** 34752 **Score:** 51 **Matches:** 1(1) **Sequences:** 1(1) **emPAI:** 0.10
leucine zipper transcription factor-like protein 1 [Mus musculus]

| Query | Observed | Mr(expt) | Mr(calc) | Delta | Miss | Score | Expect | Rank | Unique | Peptide |
|---|---|---|---|---|---|---|---|---|---|---|
| 12441 | 595.5423 | 1189.0701 | 1189.5939 | -0.5238 | 1 | 51 | 0.0072 | 1 | U | K.TLNDKTENQK.S |

**Proteins matching the same set of peptides:**

gi|26325604 **Mass:** 31688 **Score:** 51 **Matches:** 1(1) **Sequences:** 1(1)
unnamed protein product [Mus musculus]

gi|26326047 **Mass:** 34639 **Score:** 51 **Matches:** 1(1) **Sequences:** 1(1)
unnamed protein product [Mus musculus]

gi|26334199 **Mass:** 34694 **Score:** 51 **Matches:** 1(1) **Sequences:** 1(1)
unnamed protein product [Mus musculus]
gi|74181149 **Mass:** 33628 **Score:** 51 **Matches:** 1(1) **Sequences:** 1(1)
unnamed protein product [Mus musculus]

---

**29.** gi|200765 **Mass:** 90163 **Score:** 51 **Matches:** 1(1) **Sequences:** 1(1) **emPAI:** 0.04
ribonucleotide reductase subunit M1 [Mus musculus]

| Query | Observed | Mr(expt) | Mr(calc) | Delta | Miss | Score | Expect | Rank | Unique | Peptide |
|---|---|---|---|---|---|---|---|---|---|---|
| 15601 | 768.6637 | 1535.3128 | 1534.7667 | 0.5461 | 0 | 51 | 0.016 | 1 | U | R.YPFESPEAQLLNK.Q |

**Proteins matching the same set of peptides:**
gi|31982026 **Mass:** 90153 **Score:** 51 **Matches:** 1(1) **Sequences:** 1(1)
ribonucleoside-diphosphate reductase large subunit [Mus musculus]
gi|42490975 **Mass:** 90093 **Score:** 51 **Matches:** 1(1) **Sequences:** 1(1)
Ribonucleotide reductase M1 [Mus musculus]
gi|116283230 **Mass:** 86782 **Score:** 51 **Matches:** 1(1) **Sequences:** 1(1)
Rrm1 protein [Mus musculus]

---

**30.** gi|191765 **Mass:** 47195 **Score:** 51 **Matches:** 1(1) **Sequences:** 1(1) **emPAI:** 0.07
alpha-fetoprotein, partial [Mus musculus]

| Query | Observed | Mr(expt) | Mr(calc) | Delta | Miss | Score | Expect | Rank | Unique | Peptide |
|---|---|---|---|---|---|---|---|---|---|---|
| 15188 | 740.1958 | 1478.3770 | 1478.7881 | -0.4111 | 0 | 51 | 0.015 | 1 | U | K.LGEYGFQNAILVR.Y |

**Proteins matching the same set of peptides:**
gi|74137565 **Mass:** 68688 **Score:** 51 **Matches:** 1(1) **Sequences:** 1(1)
unnamed protein product [Mus musculus]
gi|26340966 **Mass:** 68678 **Score:** 51 **Matches:** 1(1) **Sequences:** 1(1)
unnamed protein product [Mus musculus]
gi|26341396 **Mass:** 64961 **Score:** 51 **Matches:** 1(1) **Sequences:** 1(1)
unnamed protein product [Mus musculus]
gi|163310765 **Mass:** 68648 **Score:** 51 **Matches:** 1(1) **Sequences:** 1(1)
serum albumin precursor [Mus musculus]

---

**31.** gi|18480346 **Score:** 50 **Matches:** 1(1) **Sequences:** 1(1) **emPAI:** 0.09
olfactory receptor MOR175-2 [Mus musculus]

| Query | Observed | Mr(expt) | Mr(calc) | Delta | Miss | Score | Expect | Rank | Unique | Peptide |
|---|---|---|---|---|---|---|---|---|---|---|
| 6979 | 366.0798 | 730.1451 | 731.3562 | -1.2111 | 0 | 50 | 0.013 | 1 | U | K.INSADGR.R |

**Proteins matching the same set of peptides:**
gi|121583707 **Score:** 50 **Matches:** 1(1) **Sequences:** 1(1)

---

**32.** gi|226005 **Mass:** 32732 **Score:** 50 **Matches:** 1(1) **Sequences:** 1(1) **emPAI:** 0.10
protein 40kD

| Query | Observed | Mr(expt) | Mr(calc) | Delta | Miss | Score | Expect | Rank | Unique | Peptide |
|---|---|---|---|---|---|---|---|---|---|---|
| 12564 | 602.6827 | 1203.3509 | 1202.6408 | 0.7102 | 0 | 50 | 0.025 | 1 | U | K.FAAATGATPIAGR.F |

**Proteins matching the same set of peptides:**
gi|293694 **Mass:** 32698 **Score:** 50 **Matches:** 1(1) **Sequences:** 1(1)
laminin receptor [Mus musculus]
gi|12846904 **Mass:** 32840 **Score:** 50 **Matches:** 1(1) **Sequences:** 1(1)
unnamed protein product [Mus musculus]
gi|26344606 **Mass:** 10928 **Score:** 50 **Matches:** 1(1) **Sequences:** 1(1)
unnamed protein product [Mus musculus]
gi|26350123 **Mass:** 32893 **Score:** 50 **Matches:** 1(1) **Sequences:** 1(1)
unnamed protein product [Mus musculus]
gi|62024907 **Mass:** 32821 **Score:** 50 **Matches:** 1(1) **Sequences:** 1(1)
Ribosomal protein SA [Mus musculus]

gi|82905443 **Mass:** 32936 **Score:** 50 **Matches:** 1(1) **Sequences:** 1(1)
PREDICTED: 40S ribosomal protein SA-like isoform 1 [Mus musculus]
gi|148237408 **Mass:** 32817 **Score:** 50 **Matches:** 1(1) **Sequences:** 1(1)
uncharacterized protein LOC100037080 [Xenopus laevis]
gi|148680000 **Mass:** 32817 **Score:** 50 **Matches:** 1(1) **Sequences:** 1(1)
mCG2650 [Mus musculus]
gi|148691799 **Mass:** 32640 **Score:** 50 **Matches:** 1(1) **Sequences:** 1(1)
mCG68071 [Mus musculus]
gi|171948782 **Mass:** 32829 **Score:** 50 **Matches:** 1(1) **Sequences:** 1(1)
laminin receptor [Mus musculus]

---

**33.** gi|193645 **Mass:** 18932 **Score:** 50 **Matches:** 1(1) **Sequences:** 1(1) **emPAI:** 0.18
glucose-regulated protein 78, partial [Mus musculus]

| Query | Observed | Mr(expt) | Mr(calc) | Delta | Miss | Score | Expect | Rank | Unique | Peptide |
|---|---|---|---|---|---|---|---|---|---|---|
| 14399 | 699.2704 | 1396.5262 | 1396.7813 | -0.2551 | 0 | 50 | 0.025 | 1 | U | K.ELEEIVQPIISK.L |

**Proteins matching the same set of peptides:**
gi|387113 **Mass:** 16164 **Score:** 50 **Matches:** 1(1) **Sequences:** 1(1)
immunoglobulin heavy chain binding protein, partial [Mus musculus]
gi|1304157 **Mass:** 72412 **Score:** 50 **Matches:** 1(1) **Sequences:** 1(1)
78 kDa glucose-regulated protein [Mus musculus]
gi|2598562 **Mass:** 72433 **Score:** 50 **Matches:** 1(1) **Sequences:** 1(1)
BiP [Mus musculus]
gi|12835845 **Mass:** 72378 **Score:** 50 **Matches:** 1(1) **Sequences:** 1(1)
unnamed protein product [Mus musculus]
gi|74188814 **Mass:** 72305 **Score:** 50 **Matches:** 1(1) **Sequences:** 1(1)
unnamed protein product [Mus musculus]
gi|74198293 **Mass:** 72419 **Score:** 50 **Matches:** 1(1) **Sequences:** 1(1)
unnamed protein product [Mus musculus]
gi|74198974 **Mass:** 72361 **Score:** 50 **Matches:** 1(1) **Sequences:** 1(1)
unnamed protein product [Mus musculus]
gi|74207492 **Mass:** 72301 **Score:** 50 **Matches:** 1(1) **Sequences:** 1(1)
unnamed protein product [Mus musculus]
gi|74225394 **Mass:** 72337 **Score:** 50 **Matches:** 1(1) **Sequences:** 1(1)
unnamed protein product [Mus musculus]
gi|148676670 **Mass:** 56257 **Score:** 50 **Matches:** 1(1) **Sequences:** 1(1)
heat shock 70kD protein 5 (glucose-regulated protein), isoform CRA_b [Mus musculus]
gi|254540166 **Mass:** 72377 **Score:** 50 **Matches:** 1(1) **Sequences:** 1(1)
78 kDa glucose-regulated protein precursor [Mus musculus]

---

**34.** gi|6996913 **Mass:** 38652 **Score:** 50 **Matches:** 1(1) **Sequences:** 1(1) **emPAI:** 0.09
annexin A2 [Mus musculus]

| Query | Observed | Mr(expt) | Mr(calc) | Delta | Miss | Score | Expect | Rank | Unique | Peptide |
|---|---|---|---|---|---|---|---|---|---|---|
| 12728 | 611.6130 | 1221.2114 | 1221.5877 | -0.3763 | 0 | 50 | 0.023 | 1 | U | K.TPAQYDASELK.A |

**Proteins matching the same set of peptides:**
gi|12849385 **Mass:** 38585 **Score:** 50 **Matches:** 1(1) **Sequences:** 1(1)
unnamed protein product [Mus musculus]
gi|74151782 **Mass:** 38636 **Score:** 50 **Matches:** 1(1) **Sequences:** 1(1)
unnamed protein product [Mus musculus]

---

**35.** gi|116283284 **Mass:** 72885 **Score:** 49 **Matches:** 1(1) **Sequences:** 1(1) **emPAI:** 0.05
Por protein [Mus musculus]

| Query | Observed | Mr(expt) | Mr(calc) | Delta | Miss | Score | Expect | Rank | Unique | Peptide |
|---|---|---|---|---|---|---|---|---|---|---|
| 13383 | 649.1569 | 1296.2993 | 1296.6363 | -0.3370 | 0 | 49 | 0.023 | 1 | U | R.WAAAPAGWAAPGR.R |

---

**36.** gi|12805465 **Mass:** 54890 **Score:** 48 **Matches:** 1(1) **Sequences:** 1(1) **emPAI:** 0.06
Dnajc10 protein, partial [Mus musculus]

| Query | Observed | Mr(expt) | Mr(calc) | Delta | Miss | Score | Expect | Rank | Unique | Peptide |
|---|---|---|---|---|---|---|---|---|---|---|
| 13357 | 647.5936 | 1293.1727 | 1292.6976 | 0.4751 | 0 | 48 | 0.033 | 1 | U | K.SSVLFLNSLDAK.E |

**Proteins matching the same set of peptides:**

gi|12835910 **Mass:** 90547 **Score:** 48 **Matches:** 1(1) **Sequences:** 1(1)
unnamed protein product [Mus musculus]
gi|23270977 **Mass:** 90441 **Score:** 48 **Matches:** 1(1) **Sequences:** 1(1)
DnaJ (Hsp40) homolog, subfamily C, member 10 [Mus musculus]
gi|25140581 **Mass:** 90768 **Score:** 48 **Matches:** 1(1) **Sequences:** 1(1)
ER-resident protein ERdj5 [Mus musculus]
gi|33340135 **Mass:** 90521 **Score:** 48 **Matches:** 1(1) **Sequences:** 1(1)
endoplasmic reticulum DnaJ-PDI fusion protein 1 precursor [Mus musculus]
gi|119508443 **Mass:** 90525 **Score:** 48 **Matches:** 1(1) **Sequences:** 1(1)
dnaJ homolog subfamily C member 10 precursor [Mus musculus]
gi|148695313 **Mass:** 80660 **Score:** 48 **Matches:** 1(1) **Sequences:** 1(1)
mCG12166 [Mus musculus]
gi|329665912 **Mass:** 88825 **Score:** 48 **Matches:** 1(1) **Sequences:** 1(1)
Chain A, Crystal Structure Of Full-Length Erdj5

---

**37.** gi|148678782 **Mass:** 9063 **Score:** 48 **Matches:** 1(1) **Sequences:** 1(1) **emPAI:** 0.38
mCG1027198, isoform CRA_a [Mus musculus]

| Query | Observed | Mr(expt) | Mr(calc) | Delta | Miss | Score | Expect | Rank | Unique | Peptide |
|---|---|---|---|---|---|---|---|---|---|---|
| 11795 | 568.0061 | 1133.9976 | 1134.5451 | -0.5475 | 1 | 48 | 0.015 | 1 | U | K.ERMESSGAIR.I |

---

**38.** gi|16716569 **Mass:** 26118 **Score:** 48 **Matches:** 1(1) **Sequences:** 1(1) **emPAI:** 0.13
protease, serine, 1 precursor [Mus musculus]

| Query | Observed | Mr(expt) | Mr(calc) | Delta | Miss | Score | Expect | Rank | Unique | Peptide |
|---|---|---|---|---|---|---|---|---|---|---|
| 15127 | 737.7551 | 2210.2436 | 2210.0967 | 0.1468 | 0 | 48 | 0.04 | 1 | U | R.LGEHNINVLEGNEQFIDAAK.I |

---

**39.** gi|19924302 **Mass:** 77934 **Score:** 47 **Matches:** 1(1) **Sequences:** 1(1) **emPAI:** 0.04
ankyrin repeat domain-containing protein 6 [Mus musculus]

| Query | Observed | Mr(expt) | Mr(calc) | Delta | Miss | Score | Expect | Rank | Unique | Peptide |
|---|---|---|---|---|---|---|---|---|---|---|
| 12288 | 588.4644 | 1174.9142 | 1174.5951 | 0.3192 | 1 | 47 | 0.02 | 1 | U | K.VMQAPIKGCR.C |

**Proteins matching the same set of peptides:**

gi|50510725 **Mass:** 78081 **Score:** 47 **Matches:** 1(1) **Sequences:** 1(1)
mKIAA0957 protein [Mus musculus]

---

**40.** gi|148680049 **Mass:** 5496 **Score:** 47 **Matches:** 1(1) **Sequences:** 1(1) **emPAI:** 0.65
mCG147414 [Mus musculus]

| Query | Observed | Mr(expt) | Mr(calc) | Delta | Miss | Score | Expect | Rank | Unique | Peptide |
|---|---|---|---|---|---|---|---|---|---|---|
| 15915 | 811.5035 | 1620.9925 | 1619.8770 | 1.1155 | 0 | 47 | 0.046 | 1 | U | M.YPLAVGATSGLVTLDK.Q |

---

**41.** gi|26337475 **Mass:** 33168 **Score:** 47 **Matches:** 1(1) **Sequences:** 1(1) **emPAI:** 0.10
unnamed protein product [Mus musculus]

| Query | Observed | Mr(expt) | Mr(calc) | Delta | Miss | Score | Expect | Rank | Unique | Peptide |
|---|---|---|---|---|---|---|---|---|---|---|
| 7984 | 789.6132 | 788.6059 | 789.3617 | -0.7558 | 0 | 47 | 0.033 | 1 | U | R.SGSSPSPR.A |

**Proteins matching the same set of peptides:**

gi|309095 **Mass:** 135080 **Score:** 47 **Matches:** 1(1) **Sequences:** 1(1)
AE3 protein [Mus musculus]
gi|51328345 **Mass:** 136564 **Score:** 47 **Matches:** 1(1) **Sequences:** 1(1)
Slc4a3 protein [Mus musculus]
gi|165377246 **Mass:** 135288 **Score:** 47 **Matches:** 1(1) **Sequences:** 1(1)
anion exchange protein 3 [Mus musculus]

---

**42.** gi|199938 **Mass:** 65935 **Score:** 46 **Matches:** 1(1) **Sequences:** 1(1) **emPAI:** 0.05
tumor-specific myb protein [Mus musculus]

| Query | Observed | Mr(expt) | Mr(calc) | Delta | Miss | Score | Expect | Rank | Unique | Peptide |
|---|---|---|---|---|---|---|---|---|---|---|
| 11505 | 555.3855 | 1108.7564 | 1108.4715 | 0.2849 | 0 | 46 | 0.025 | 1 | U | -.MGAPLNCPMK.S |

**Proteins matching the same set of peptides:**
gi|199929 **Mass:** 79128 **Score:** 46 **Matches:** 1(1) **Sequences:** 1(1)
myb protein [Mus musculus]

---

**43.** gi|22094097 **Mass:** 56233 **Score:** 46 **Matches:** 1(1) **Sequences:** 1(1) **emPAI:** 0.06
ganglioside-induced differentiation-associated protein 2 [Mus musculus]

| Query | Observed | Mr(expt) | Mr(calc) | Delta | Miss | Score | Expect | Rank | Unique | Peptide |
|---|---|---|---|---|---|---|---|---|---|---|
| 7268 | 747.2545 | 746.2473 | 747.4028 | -1.1555 | 0 | 46 | 0.04 | 1 | U | K.GFNLAAR.F |

**Proteins matching the same set of peptides:**
gi|26350139 **Mass:** 60978 **Score:** 46 **Matches:** 1(1) **Sequences:** 1(1)
unnamed protein product [Mus musculus]
gi|148675693 **Mass:** 60979 **Score:** 46 **Matches:** 1(1) **Sequences:** 1(1)
ganglioside-induced differentiation-associated-protein 2, isoform CRA_a [Mus musculus]

---

**44.** gi|198683 **Mass:** 35858 **Score:** 45 **Matches:** 1(1) **Sequences:** 1(1) **emPAI:** 0.09
unnamed protein product [Mus musculus]

| Query | Observed | Mr(expt) | Mr(calc) | Delta | Miss | Score | Expect | Rank | Unique | Peptide |
|---|---|---|---|---|---|---|---|---|---|---|
| 9577 | 457.5262 | 913.0378 | 912.5756 | 0.4622 | 0 | 45 | 0.024 | 1 | U | K.LVIITAGAR.M |

**Proteins matching the same set of peptides:**
gi|6754524 **Mass:** 36475 **Score:** 45 **Matches:** 1(1) **Sequences:** 1(1)
L-lactate dehydrogenase A chain isoform 1 [Mus musculus]
gi|7305229 **Mass:** 35889 **Score:** 45 **Matches:** 1(1) **Sequences:** 1(1)
L-lactate dehydrogenase C chain [Mus musculus]
gi|13529599 **Mass:** 34481 **Score:** 45 **Matches:** 1(1) **Sequences:** 1(1)
Ldha protein, partial [Mus musculus]
gi|44890285 **Mass:** 11164 **Score:** 45 **Matches:** 1(1) **Sequences:** 1(1)
Ldha protein [Mus musculus]
gi|74204388 **Mass:** 36461 **Score:** 45 **Matches:** 1(1) **Sequences:** 1(1)
unnamed protein product [Mus musculus]
gi|74208131 **Mass:** 36242 **Score:** 45 **Matches:** 1(1) **Sequences:** 1(1)
unnamed protein product [Mus musculus]
gi|74212250 **Mass:** 36476 **Score:** 45 **Matches:** 1(1) **Sequences:** 1(1)
unnamed protein product [Mus musculus]
gi|74217959 **Mass:** 36476 **Score:** 45 **Matches:** 1(1) **Sequences:** 1(1)
unnamed protein product [Mus musculus]
gi|157835307 **Mass:** 35801 **Score:** 45 **Matches:** 1(1) **Sequences:** 1(1)
Chain A, Characterization Of The Antigenic Sites On The Refined 3- Angstroms Resolution Structure Of Mouse Testicular Lactate Dehydrogenase C4
gi|257743039 **Mass:** 39733 **Score:** 45 **Matches:** 1(1) **Sequences:** 1(1)
L-lactate dehydrogenase A chain isoform 2 [Mus musculus]

---

**45.** gi|26327761 **Mass:** 90833 **Score:** 45 **Matches:** 1(1) **Sequences:** 1(1) **emPAI:** 0.04
unnamed protein product [Mus musculus]

| Query | Observed | Mr(expt) | Mr(calc) | Delta | Miss | Score | Expect | Rank | Unique | Peptide |
|---|---|---|---|---|---|---|---|---|---|---|
| 11852 | 571.0167 | 1140.0188 | 1139.6550 | 0.3638 | 0 | 45 | 0.026 | 1 | U | K.LALPADSVNIK.I |

**Proteins matching the same set of peptides:**
gi|26332879 **Mass:** 40136 **Score:** 45 **Matches:** 1(1) **Sequences:** 1(1)
unnamed protein product [Mus musculus]
gi|38257756 **Mass:** 90916 **Score:** 45 **Matches:** 1(1) **Sequences:** 1(1)
RecName: Full=Cullin-5; Short=CUL-5

gi|239051067 **Mass:** 98979 **Score:** 45 **Matches:** 1(1) **Sequences:** 1(1)
cullin-5 isoform 1 [Mus musculus]
gi|239051082 **Mass:** 95790 **Score:** 45 **Matches:** 1(1) **Sequences:** 1(1)
cullin-5 isoform 2 [Mus musculus]

---

**46.** gi|51092301 **Mass:** 60191 **Score:** 44 **Matches:** 1(0) **Sequences:** 1(0) **emPAI:** 0.05
type II keratin Kb14 [Mus musculus]

| Query | Observed | Mr(expt) | Mr(calc) | Delta | Miss | Score | Expect | Rank | Unique | Peptide |
|---|---|---|---|---|---|---|---|---|---|---|
| 13359 | 647.7845 | 1940.3318 | 1940.9996 | -0.6678 | 1 | 44 | 0.072 | 1 | U | R.FASFIDKVQFLEQQNK.V |

**Proteins matching the same set of peptides:**
gi|148672085 **Mass:** 38119 **Score:** 44 **Matches:** 1(0) **Sequences:** 1(0)
mCG144996 [Mus musculus]

---

**47.** gi|74208129 **Mass:** 95473 **Score:** 44 **Matches:** 1(1) **Sequences:** 1(1) **emPAI:** 0.03
unnamed protein product [Mus musculus]

| Query | Observed | Mr(expt) | Mr(calc) | Delta | Miss | Score | Expect | Rank | Unique | Peptide |
|---|---|---|---|---|---|---|---|---|---|---|
| 11982 | 577.0237 | 1152.0328 | 1152.4758 | -0.4430 | 0 | 44 | 0.028 | 1 | U | R.SPSYPPPGCGK.S |

**Proteins matching the same set of peptides:**
gi|74211952 **Mass:** 35351 **Score:** 44 **Matches:** 1(1) **Sequences:** 1(1)
unnamed protein product [Mus musculus]
gi|116248568 **Mass:** 95499 **Score:** 44 **Matches:** 1(1) **Sequences:** 1(1)
RecName: Full=Poly [ADP-ribose] polymerase 8; Short=PARP-8
gi|124486598 **Mass:** 99423 **Score:** 44 **Matches:** 1(1) **Sequences:** 1(1)
poly [ADP-ribose] polymerase 8 [Mus musculus]
gi|148686417 **Mass:** 92421 **Score:** 44 **Matches:** 1(1) **Sequences:** 1(1)
mCG119138 [Mus musculus]

---

**48.** gi|148685252 **Mass:** 396271 **Score:** 44 **Matches:** 1(1) **Sequences:** 1(1) **emPAI:** 0.01
mCG142044 [Mus musculus]

| Query | Observed | Mr(expt) | Mr(calc) | Delta | Miss | Score | Expect | Rank | Unique | Peptide |
|---|---|---|---|---|---|---|---|---|---|---|
| 11130 | 537.3717 | 1072.7289 | 1071.5924 | 1.1365 | 1 | 44 | 0.051 | 1 | U | R.AELKLVEDR.L |

**Proteins matching the same set of peptides:**
gi|182637561 **Mass:** 467476 **Score:** 44 **Matches:** 1(1) **Sequences:** 1(1)
RecName: Full=Dynein heavy chain 3, axonemal; AltName: Full=Axonemal beta dynein heavy chain 3; AltName: Full=Ciliary dynein heavy chain 3

---

**49.** gi|148671989 **Score:** 44 **Matches:** 1(1) **Sequences:** 1(1) **emPAI:** 0.24
mCG1027581 [Mus musculus]

| Query | Observed | Mr(expt) | Mr(calc) | Delta | Miss | Score | Expect | Rank | Unique | Peptide |
|---|---|---|---|---|---|---|---|---|---|---|
| 7739 | 387.8668 | 773.7190 | 774.4236 | -0.7045 | 0 | 44 | 0.041 | 1 | U | K.ISSPSLR.I |

---

**50.** gi|26349347 **Mass:** 135092 **Score:** 44 **Matches:** 1(1) **Sequences:** 1(1) **emPAI:** 0.02
unnamed protein product [Mus musculus]

| Query | Observed | Mr(expt) | Mr(calc) | Delta | Miss | Score | Expect | Rank | Unique | Peptide |
|---|---|---|---|---|---|---|---|---|---|---|
| 13447 | 652.5267 | 1303.0388 | 1302.6238 | 0.4150 | 0 | 44 | 0.039 | 1 | U | -.MVEAAPAGSGPLR.R |

**Proteins matching the same set of peptides:**
gi|148689974 **Mass:** 136815 **Score:** 44 **Matches:** 1(1) **Sequences:** 1(1)
RIKEN cDNA 2310057J16, isoform CRA_a [Mus musculus]
gi|255069774 **Mass:** 135191 **Score:** 44 **Matches:** 1(1) **Sequences:** 1(1)
calmodulin-regulated spectrin-associated protein 3 isoform 1 [Mus musculus]
gi|255069776 **Mass:** 135091 **Score:** 44 **Matches:** 1(1) **Sequences:** 1(1)
calmodulin-regulated spectrin-associated protein 3 isoform 2 [Mus musculus]

---

**51.** gi|6755893 **Mass:** 26257 **Score:** 43 **Matches:** 1(0) **Sequences:** 1(0) **emPAI:** 0.13

trypsin 4 precursor [Mus musculus]

| Query | Observed | Mr(expt) | Mr(calc) | Delta | Miss | Score | Expect | Rank | Unique | Peptide |
|---|---|---|---|---|---|---|---|---|---|---|
| 15139 | 738.1550 | 2211.4433 | 2211.0920 | 0.3513 | 0 | 43 | 0.079 | 1 | U | R.LGEHNINVLEGNEQFVNSAK.I |

**Proteins matching the same set of peptides:**

gi|51010909 **Mass:** 26260 **Score:** 43 **Matches:** 1(0) **Sequences:** 1(0)
trypsin 5 precursor [Mus musculus]

gi|74203392 **Mass:** 27146 **Score:** 43 **Matches:** 1(0) **Sequences:** 1(0)
unnamed protein product [Mus musculus]

---

**52.** gi|6754976 **Mass:** 22162 **Score:** 43 **Matches:** 1(1) **Sequences:** 1(1) **emPAI:** 0.15
peroxiredoxin-1 [Mus musculus]

| Query | Observed | Mr(expt) | Mr(calc) | Delta | Miss | Score | Expect | Rank | Unique | Peptide |
|---|---|---|---|---|---|---|---|---|---|---|
| 11476 | 554.4471 | 1106.8797 | 1106.5972 | 0.2826 | 0 | 43 | 0.039 | 1 | U | R.TIAQDYGVLK.A |

**Proteins matching the same set of peptides:**

gi|12846314 **Mass:** 22222 **Score:** 43 **Matches:** 1(1) **Sequences:** 1(1)
unnamed protein product [Mus musculus]

gi|74198890 **Mass:** 22130 **Score:** 43 **Matches:** 1(1) **Sequences:** 1(1)
unnamed protein product [Mus musculus]

gi|148665674 **Mass:** 22248 **Score:** 43 **Matches:** 1(1) **Sequences:** 1(1)
mCG127770 [Mus musculus]

---

**Mascot:** http://www.matrixscience.com/

PFHR9

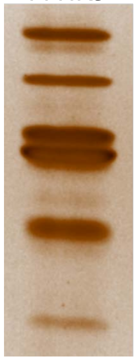

2

### Mascot Search Results

```
User                   :
Email                  :
Search title           :
MS data file           : ZIEN-29Nov2017_2m.mgf
Database               : NCBInr 20120419 (17893860 sequences; 6141683785 residues)
Taxonomy               : Mus musculus (house mouse) (138831 sequences)
Timestamp              : 30 Nov 2017 at 20:24:37 GMT
Enzyme                 : Trypsin
Variable modifications : Carbamidomethyl (C),Oxidation (P),Oxidation (M)
Mass values            : Monoisotopic
Protein Mass           : Unrestricted
Peptide Mass Tolerance : ± 1.25 Da
Fragment Mass Tolerance: ± 1.001 Da
Max Missed Cleavages   : 1
Instrument type        : Default
Number of queries      : 16468
Protein hits           : gi|2598562   BiP [Mus musculus]
                         gi|14917005  RecName: Full=Stress-70 protein, mitochondrial; AltName: Full=75 kDa glucose-regulated protein; Short=GRP-75; AltName: Full=Heat shock 70 kDa protein 9; AltName: Full=
                         gi|74211667  unnamed protein product [Mus musculus]
                         gi|2498741   RecName: Full=Prolyl 4-hydroxylase subunit alpha-2; Short=4-PH alpha-2; AltName: Full=Procollagen-proline,2-oxoglutarate-4-dioxygenase subunit alpha-2; Flags: Precursc
                         gi|387397    epidermal keratin subunit I, partial [Mus musculus]
                         gi|46485130  TPA_exp: keratin Kb40 [Mus musculus]
                         gi|85701680  keratin, type II cytoskeletal 2 oral [Mus musculus]
                         gi|387399    epidermal keratin type I, partial [Mus musculus]
                         gi|16303309  type II keratin 5 [Mus musculus]
                         gi|26349293  unnamed protein product [Mus musculus]
                         gi|6753620   ATP-dependent RNA helicase DDX3X [Mus musculus]
                         gi|47523977  keratin, type II cytoskeletal 72 [Mus musculus]
                         gi|16716569  protease, serine, 1 precursor [Mus musculus]
                         gi|1389682   plakoglobin, partial [Mus musculus]
                         gi|19354359  Ranbp10 protein [Mus musculus]
                         gi|52789     unnamed protein product [Mus musculus]
                         gi|54777     unnamed protein product [Mus musculus]
                         gi|20070408  glycine dehydrogenase [decarboxylating], mitochondrial precursor [Mus musculus]
                         gi|3642689   immunoglobulin heavy chain-binding protein, partial [Mus musculus]
                         gi|26344548  unnamed protein product [Mus musculus]
                         gi|836898    prolyl 4-hydroxylase alpha(I)-subunit, partial [Mus musculus]
                         gi|4159806   type II keratin subunit protein [Mus musculus]
                         gi|37935731  ATP-binding cassette transporter sub-family A member 16 [Mus musculus]
                         gi|21595163  Glycosyltransferase 25 domain containing 1 [Mus musculus]
                         gi|12847276  unnamed protein product [Mus musculus]
                         gi|74191500  unnamed protein product [Mus musculus]
                         gi|81875970  RecName: Full=Alkyldihydroxyacetonephosphate synthase, peroxisomal; Short=Alkyl-DHAP synthase; AltName: Full=Alkylglycerone-phosphate synthase; Flags: Precursor
                         gi|7513694   IgG Fc binding protein - mouse (fragment)
                         gi|148664969 RIKEN cDNA 2900011O08, isoform CRA_a [Mus musculus]
                         gi|12855410  unnamed protein product [Mus musculus]
```

#### Select Summary Report

Format As | Select Summary (protein hits) | Help

Significance threshold p< 0.05 | Max. number of hits AUTO

Standard scoring ○ MudPIT scoring ◉ | Ions score or expect cut-off 43 | Show sub-sets 0

Show pop-ups ◉ Suppress pop-ups ○ | Require bold red ☐

Re-Search | ◉ All queries ○ Unassigned ○ Below homology threshold ○ Below identity threshold

1. gi|2598562 **Mass:** 72433 **Score:** 183 **Matches:** 10(8) **Sequences:** 6(6) **emPAI:** 0.30

BiP [Mus musculus]

| Query | Observed | Mr(expt) | Mr(calc) | Delta | Miss | Score | Expect | Rank | Unique | Peptide |
|---|---|---|---|---|---|---|---|---|---|---|
| 7641 | 758.4543 | 757.4470 | 757.4334 | 0.0136 | 0 | 51 | 0.011 | 1 | U | R.NTVVPTK.K |
| 10764 | 499.1691 | 996.3237 | 996.5101 | -0.1863 | 0 | 49 | 0.012 | 1 | U | R.ALSSQHQAR.I |
| 13124 | 609.0809 | 1216.1473 | 1216.6234 | -0.4761 | 0 | 52 | 0.011 | 1 | U | K.DAGTIAGLNVMR.I 13134 |

| | | | | | | | | | | |
|---|---|---|---|---|---|---|---|---|---|---|
| 13280 | 615.1929 | 1228.3712 | 1227.6207 | 0.7504 | 0 | 66 | 0.00052 | 1 | | R.VEIIANDQGNR.I |
| 14833 | 699.2899 | 1396.5653 | 1396.7813 | -0.2160 | 0 | 60 | 0.0025 | 1 | U | K.ELEEIVQPIISK.L 14827 |
| 15459 | 731.0733 | 1460.1320 | 1459.7518 | 0.3802 | 0 | 79 | 2.8e-005 | 1 | U | K.SDIDEIVLVGGSTR.I 15458 15465 |

**Proteins matching the same set of peptides:**
gi|12835845 **Mass:** 72378 **Score:** 183 **Matches:** 10(8) **Sequences:** 6(6)
unnamed protein product [Mus musculus]
gi|74188814 **Mass:** 72305 **Score:** 183 **Matches:** 10(8) **Sequences:** 6(6)
unnamed protein product [Mus musculus]
gi|74198293 **Mass:** 72419 **Score:** 183 **Matches:** 10(8) **Sequences:** 6(6)
unnamed protein product [Mus musculus]
gi|74198974 **Mass:** 72361 **Score:** 183 **Matches:** 10(8) **Sequences:** 6(6)
unnamed protein product [Mus musculus]
gi|74207492 **Mass:** 72301 **Score:** 183 **Matches:** 10(8) **Sequences:** 6(6)
unnamed protein product [Mus musculus]
gi|74225394 **Mass:** 72337 **Score:** 183 **Matches:** 10(8) **Sequences:** 6(6)
unnamed protein product [Mus musculus]
gi|254540166 **Mass:** 72377 **Score:** 183 **Matches:** 10(8) **Sequences:** 6(6)
78 kDa glucose-regulated protein precursor [Mus musculus]

---

**2.** gi|14917005 **Mass:** 73483 **Score:** 174 **Matches:** 10(10) **Sequences:** 7(7) **emPAI:** 0.36
RecName: Full=Stress-70 protein, mitochondrial; AltName: Full=75 kDa glucose-regulated protein; Short=GRP-75; AltName: Full=Heat shock 70 kDa protein 9; AltName: Full=Mortalin; AltName: Full=Peptide

| Query | Observed | Mr(expt) | Mr(calc) | Delta | Miss | Score | Expect | Rank | Unique | Peptide |
|---|---|---|---|---|---|---|---|---|---|---|
| 10606 | 978.6167 | 977.6095 | 977.4488 | 0.1607 | 0 | 54 | 0.0056 | 1 | U | K.AMQDAEVSK.S |
| 13304 | 616.1958 | 1230.3770 | 1230.6568 | -0.2797 | 0 | 48 | 0.034 | 1 | U | R.QAASSLQQASLK.L |
| 13836 | 645.6029 | 1289.1913 | 1289.6728 | -0.4816 | 0 | 56 | 0.0048 | 1 | U | K.VQQTVQDLFGR.A 13846 13847 |
| 14194 | 667.1742 | 1332.3338 | 1332.7289 | -0.3951 | 0 | 56 | 0.0025 | 1 | U | R.AQFEGIVTDLIK.R |
| 15362 | 726.0390 | 1450.0635 | 1449.7100 | 0.3535 | 0 | 60 | 0.00088 | 1 | U | R.TTPSVVAFTADGER.L 15364 |
| 16185 | 786.0322 | 1570.0499 | 1568.8272 | 1.2227 | 0 | 46 | 0.053 | 1 | U | K.LYSPSQIGAFVLMK.M |
| 15239 | 720.0499 | 2157.1280 | 2157.0372 | 0.0908 | 1 | 61 | 0.0018 | 1 | U | K.ERVEAVNMAEGIIHDTETK.M |

**Proteins matching the same set of peptides:**
gi|74204605 **Mass:** 73446 **Score:** 174 **Matches:** 10(10) **Sequences:** 7(7)
unnamed protein product [Mus musculus]
gi|74225724 **Mass:** 73431 **Score:** 174 **Matches:** 10(10) **Sequences:** 7(7)
unnamed protein product [Mus musculus]
gi|162461907 **Mass:** 73416 **Score:** 174 **Matches:** 10(10) **Sequences:** 7(7)
stress-70 protein, mitochondrial [Mus musculus]

---

**3.** gi|74211667 **Mass:** 60764 **Score:** 169 **Matches:** 9(8) **Sequences:** 9(8) **emPAI:** 0.61
unnamed protein product [Mus musculus]

| Query | Observed | Mr(expt) | Mr(calc) | Delta | Miss | Score | Expect | Rank | Unique | Peptide |
|---|---|---|---|---|---|---|---|---|---|---|
| 7734 | 765.3373 | 764.3300 | 764.4068 | -0.0768 | 0 | 48 | 0.016 | 1 | U | K.VQVEYK.G |
| 13280 | 615.1929 | 1228.3712 | 1227.6207 | 0.7504 | 0 | 66 | 0.00052 | 1 | | K.VEIIANDQGNR.T |
| 13478 | 627.0481 | 1252.0816 | 1252.6088 | -0.5271 | 0 | 47 | 0.039 | 1 | U | R.FEELNADLFR.G |
| 13632 | 634.9739 | 1267.9332 | 1267.6482 | 0.2850 | 1 | 54 | 0.0091 | 1 | U | K.MKEIAEAYLGK.T |
| 13950 | 652.5933 | 1303.1720 | 1302.5914 | 0.5806 | 0 | 43 | 0.094 | 1 | U | K.NSLESYAFNMK.A |
| 15105 | 713.9242 | 1425.8338 | 1425.6558 | 0.1780 | 1 | 66 | 0.00051 | 1 | U | R.RFDDAVVQSDMK.H |
| 15642 | 741.6528 | 1481.2910 | 1480.7998 | 0.4912 | 0 | 70 | 0.00019 | 1 | U | K.SQIHDIVLVGGSTR.I |
| 16318 | 809.0371 | 1616.0595 | 1615.7804 | 0.2792 | 0 | 63 | 0.0011 | 1 | U | K.SFYPEEVSSMVLTK.M |
| 15925 | 759.1199 | 2274.3378 | 2275.1332 | -0.7954 | 0 | 49 | 0.023 | 1 | U | K.SINPDEAVAYGAAVQAAILSGDK.S |

**Proteins matching the same set of peptides:**
gi|309319 **Mass:** 70793 **Score:** 169 **Matches:** 9(8) **Sequences:** 9(8)
heat shock protein 70 cognate [Mus musculus]
gi|13242237 **Mass:** 70827 **Score:** 169 **Matches:** 9(8) **Sequences:** 9(8)
heat shock cognate 71 kDa protein [Rattus norvegicus]
gi|42542422 **Mass:** 70828 **Score:** 169 **Matches:** 9(8) **Sequences:** 9(8)

Heat shock protein 8 [Mus musculus]
gi|63101351 **Mass:** 68736 **Score:** 169 **Matches:** 9(8) **Sequences:** 9(8)
Hspa8 protein [Mus musculus]
gi|74143862 **Mass:** 70813 **Score:** 169 **Matches:** 9(8) **Sequences:** 9(8)
unnamed protein product [Mus musculus]
gi|74181586 **Mass:** 70855 **Score:** 169 **Matches:** 9(8) **Sequences:** 9(8)
unnamed protein product [Mus musculus]
gi|74184057 **Mass:** 70857 **Score:** 169 **Matches:** 9(8) **Sequences:** 9(8)
unnamed protein product [Mus musculus]
gi|74186087 **Mass:** 70739 **Score:** 169 **Matches:** 9(8) **Sequences:** 9(8)
unnamed protein product [Mus musculus]
gi|74190799 **Mass:** 70813 **Score:** 169 **Matches:** 9(8) **Sequences:** 9(8)
unnamed protein product [Mus musculus]
gi|74191381 **Mass:** 70859 **Score:** 169 **Matches:** 9(8) **Sequences:** 9(8)
unnamed protein product [Mus musculus]
gi|74198858 **Mass:** 70852 **Score:** 169 **Matches:** 9(8) **Sequences:** 9(8)
unnamed protein product [Mus musculus]
gi|74198978 **Mass:** 70903 **Score:** 169 **Matches:** 9(8) **Sequences:** 9(8)
unnamed protein product [Mus musculus]
gi|74208631 **Mass:** 70827 **Score:** 169 **Matches:** 9(8) **Sequences:** 9(8)
unnamed protein product [Mus musculus]
gi|74214176 **Mass:** 70828 **Score:** 169 **Matches:** 9(8) **Sequences:** 9(8)
unnamed protein product [Mus musculus]
gi|74220416 **Mass:** 70755 **Score:** 169 **Matches:** 9(8) **Sequences:** 9(8)
unnamed protein product [Mus musculus]
gi|74220592 **Mass:** 70648 **Score:** 169 **Matches:** 9(8) **Sequences:** 9(8)
unnamed protein product [Mus musculus]
gi|74225511 **Mass:** 70783 **Score:** 169 **Matches:** 9(8) **Sequences:** 9(8)
unnamed protein product [Mus musculus]

---

**4.** gi|2498741 **Mass:** 60964 **Score:** 120 **Matches:** 2(2) **Sequences:** 2(2) **emPAI:** 0.11
RecName: Full=Prolyl 4-hydroxylase subunit alpha-2; Short=4-PH alpha-2; AltName: Full=Procollagen-proline,2-oxoglutarate-4-dioxygenase subunit alpha-2; Flags: Precursor

| Query | Observed | Mr(expt) | Mr(calc) | Delta | Miss | Score | Expect | Rank | Unique | Peptide |
|---|---|---|---|---|---|---|---|---|---|---|
| 11452 | 534.0728 | 1066.1309 | 1065.5819 | 0.5491 | 0 | 74 | 6.3e-005 | 1 | U | K.TGVLTVASYR.V |
| 16319 | 809.4653 | 1616.9160 | 1616.7318 | 0.1842 | 0 | 91 | 1.9e-006 | 1 | U | K.SSWLEEDDDPVVAR.V |

**Proteins matching the same set of peptides:**
gi|74216495 **Mass:** 21055 **Score:** 120 **Matches:** 2(2) **Sequences:** 2(2)
unnamed protein product [Mus musculus]
gi|148701597 **Mass:** 57417 **Score:** 120 **Matches:** 2(2) **Sequences:** 2(2)
procollagen-proline, 2-oxoglutarate 4-dioxygenase (proline 4-hydroxylase), alpha II polypeptide, isoform CRA_b [Mus musculus]
gi|148701598 **Mass:** 21273 **Score:** 120 **Matches:** 2(2) **Sequences:** 2(2)
procollagen-proline, 2-oxoglutarate 4-dioxygenase (proline 4-hydroxylase), alpha II polypeptide, isoform CRA_c [Mus musculus]
gi|148701599 **Mass:** 61292 **Score:** 120 **Matches:** 2(2) **Sequences:** 2(2)
procollagen-proline, 2-oxoglutarate 4-dioxygenase (proline 4-hydroxylase), alpha II polypeptide, isoform CRA_d [Mus musculus]
gi|148701600 **Mass:** 66870 **Score:** 120 **Matches:** 2(2) **Sequences:** 2(2)
procollagen-proline, 2-oxoglutarate 4-dioxygenase (proline 4-hydroxylase), alpha II polypeptide, isoform CRA_e [Mus musculus]
gi|209862961 **Mass:** 60774 **Score:** 120 **Matches:** 2(2) **Sequences:** 2(2)
prolyl 4-hydroxylase subunit alpha-2 isoform 1 precursor [Mus musculus]
gi|226874876 **Mass:** 60977 **Score:** 120 **Matches:** 2(2) **Sequences:** 2(2)
prolyl 4-hydroxylase subunit alpha-2 isoform 2 precursor [Mus musculus]

---

**5.** gi|387397 **Mass:** 57807 **Score:** 100 **Matches:** 2(2) **Sequences:** 2(2) **emPAI:** 0.12
epidermal keratin subunit I, partial [Mus musculus]

| Query | Observed | Mr(expt) | Mr(calc) | Delta | Miss | Score | Expect | Rank | Unique | Peptide |
|---|---|---|---|---|---|---|---|---|---|---|
| 14668 | 691.2913 | 1380.5680 | 1380.6408 | -0.0729 | 0 | 68 | 0.00035 | 1 | U | R.ALEESNYELEGK.I |
| 14765 | 696.1694 | 1390.3243 | 1389.6736 | 0.6508 | 0 | 71 | 0.00013 | 1 | U | K.QSLEASLAETEGR.Y |

**Proteins matching the same set of peptides:**
gi|12852157 **Mass:** 58587 **Score:** 100 **Matches:** 2(2) **Sequences:** 2(2)
unnamed protein product [Mus musculus]
gi|26349141 **Mass:** 52653 **Score:** 100 **Matches:** 2(2) **Sequences:** 2(2)
unnamed protein product [Mus musculus]
gi|26349459 **Mass:** 49471 **Score:** 100 **Matches:** 2(2) **Sequences:** 2(2)
unnamed protein product [Mus musculus]
gi|112983636 **Mass:** 57007 **Score:** 100 **Matches:** 2(2) **Sequences:** 2(2)
keratin, type I cytoskeletal 10 [Mus musculus]
gi|116242600 **Mass:** 57735 **Score:** 100 **Matches:** 2(2) **Sequences:** 2(2)
RecName: Full=Keratin, type I cytoskeletal 10; AltName: Full=56 kDa cytokeratin; AltName: Full=Cytokeratin-10; Short=CK-10; AltName: Full=Keratin, type I cytoskeletal 59 kDa; AltName: Full=Keratin-1(

---

**6.** gi|46485130 **Mass:** 85186 **Score:** 79 **Matches:** 2(2) **Sequences:** 2(2) **emPAI:** 0.08
TPA_exp: keratin Kb40 [Mus musculus]

| Query | Observed | Mr(expt) | Mr(calc) | Delta | Miss | Score | Expect | Rank | Unique | Peptide |
|---|---|---|---|---|---|---|---|---|---|---|
| 14685 | 692.1215 | 1382.2284 | 1382.6830 | -0.4546 | 0 | 57 | 0.0039 | 1 | U | R.SLNNQFASFIDK.V |
| 15600 | 739.0794 | 1476.1442 | 1475.7984 | 0.3459 | 1 | 60 | 0.0008 | 1 | | R.FLEQQNKVLETK.W |

**Proteins matching the same set of peptides:**
gi|111185567 **Mass:** 54730 **Score:** 79 **Matches:** 2(2) **Sequences:** 2(2)
Krt78 protein [Mus musculus]
gi|111185722 **Mass:** 56744 **Score:** 79 **Matches:** 2(2) **Sequences:** 2(2)
Krt78 protein [Mus musculus]
gi|119850791 **Mass:** 54739 **Score:** 79 **Matches:** 2(2) **Sequences:** 2(2)
Krt78 protein [Mus musculus]
gi|145580629 **Mass:** 112194 **Score:** 79 **Matches:** 2(2) **Sequences:** 2(2)
keratin Kb40 [Mus musculus]

---

**7.** gi|85701680 **Mass:** 62806 **Score:** 72 **Matches:** 2(2) **Sequences:** 2(2) **emPAI:** 0.11
keratin, type II cytoskeletal 2 oral [Mus musculus]

| Query | Observed | Mr(expt) | Mr(calc) | Delta | Miss | Score | Expect | Rank | Unique | Peptide |
|---|---|---|---|---|---|---|---|---|---|---|
| 11433 | 1065.4515 | 1064.4442 | 1064.5138 | -0.0696 | 0 | 54 | 0.0051 | 1 | U | K.AQYEDIAQK.S |
| 15600 | 739.0794 | 1476.1442 | 1475.7984 | 0.3459 | 1 | 60 | 0.0008 | 1 | | R.FLEQQNKVLETK.W |

**Proteins matching the same set of peptides:**
gi|148672072 **Mass:** 63519 **Score:** 72 **Matches:** 2(2) **Sequences:** 2(2)
mCG144546 [Mus musculus]

---

**8.** gi|387399 **Mass:** 10655 **Score:** 64 **Matches:** 1(1) **Sequences:** 1(1) **emPAI:** 0.32
epidermal keratin type I, partial [Mus musculus]

| Query | Observed | Mr(expt) | Mr(calc) | Delta | Miss | Score | Expect | Rank | Unique | Peptide |
|---|---|---|---|---|---|---|---|---|---|---|
| 16047 | 770.4561 | 2308.3463 | 2308.0567 | 0.2896 | 0 | 64 | 0.00083 | 1 | U | R.LLEGEDAHLSSSQFSSGSQSSR.D |

---

**9.** gi|16303309 **Mass:** 61743 **Score:** 62 **Matches:** 1(1) **Sequences:** 1(1) **emPAI:** 0.05
type II keratin 5 [Mus musculus]

| Query | Observed | Mr(expt) | Mr(calc) | Delta | Miss | Score | Expect | Rank | Unique | Peptide |
|---|---|---|---|---|---|---|---|---|---|---|
| 15266 | 721.0629 | 1440.1112 | 1440.6629 | -0.5517 | 0 | 62 | 0.0013 | 1 | U | R.VDALMDEINFMK.M |

**Proteins matching the same set of peptides:**
gi|20911031 **Mass:** 61729 **Score:** 62 **Matches:** 1(1) **Sequences:** 1(1)
keratin, type II cytoskeletal 5 [Mus musculus]

---

**10.** gi|26349293 **Mass:** 76506 **Score:** 61 **Matches:** 1(1) **Sequences:** 1(1) **emPAI:** 0.04
unnamed protein product [Mus musculus]

| Query | Observed | Mr(expt) | Mr(calc) | Delta | Miss | Score | Expect | Rank | Unique | Peptide |
|---|---|---|---|---|---|---|---|---|---|---|
| 13424 | 624.1697 | 1246.3248 | 1245.6717 | 0.6531 | 0 | 61 | 0.0014 | 1 | U | R.AFINLEAAGVGGK.E |

**Proteins matching the same set of peptides:**
gi|37360558 **Mass:** 99638 **Score:** 61 **Matches:** 1(1) **Sequences:** 1(1)
mKIAA1815 protein [Mus musculus]
gi|74209380 **Mass:** 100058 **Score:** 61 **Matches:** 1(1) **Sequences:** 1(1)
unnamed protein product [Mus musculus]
gi|124487057 **Mass:** 100084 **Score:** 61 **Matches:** 1(1) **Sequences:** 1(1)
endoplasmic reticulum metallopeptidase 1 [Mus musculus]
gi|148709741 **Mass:** 94565 **Score:** 61 **Matches:** 1(1) **Sequences:** 1(1)
mCG124990, isoform CRA_a [Mus musculus]
gi|148709742 **Mass:** 102045 **Score:** 61 **Matches:** 1(1) **Sequences:** 1(1)
mCG124990, isoform CRA_b [Mus musculus]

---

**11.** gi|6753620 **Mass:** 73056 **Score:** 61 **Matches:** 1(1) **Sequences:** 1(1) **emPAI:** 0.04
ATP-dependent RNA helicase DDX3X [Mus musculus]

| Query | Observed | Mr(expt) | Mr(calc) | Delta | Miss | Score | Expect | Rank | Unique | Peptide |
|---|---|---|---|---|---|---|---|---|---|---|
| 14436 | 679.2409 | 1356.4673 | 1356.7765 | -0.3093 | 0 | 61 | 0.0017 | 1 | U | K.QYPISLVLAPTR.E |

**Proteins matching the same set of peptides:**
gi|14861844 **Mass:** 73095 **Score:** 61 **Matches:** 1(1) **Sequences:** 1(1)
putative ATP-dependent RNA helicase Pl10 [Mus musculus]
gi|25141235 **Mass:** 73382 **Score:** 61 **Matches:** 1(1) **Sequences:** 1(1)
ATP-dependent RNA helicase DDX3Y [Mus musculus]
gi|74181660 **Mass:** 73339 **Score:** 61 **Matches:** 1(1) **Sequences:** 1(1)
unnamed protein product [Mus musculus]
gi|148706200 **Mass:** 71999 **Score:** 61 **Matches:** 1(1) **Sequences:** 1(1)
DEAD (Asp-Glu-Ala-Asp) box polypeptide 3, Y-linked, isoform CRA_a [Mus musculus]
gi|219521150 **Mass:** 73027 **Score:** 61 **Matches:** 1(1) **Sequences:** 1(1)
Ddx3x protein [Mus musculus]
gi|223462261 **Mass:** 72969 **Score:** 61 **Matches:** 1(1) **Sequences:** 1(1)
Ddx3x protein [Mus musculus]

---

**12.** gi|47523977 **Mass:** 56715 **Score:** 60 **Matches:** 1(1) **Sequences:** 1(1) **emPAI:** 0.06
keratin, type II cytoskeletal 72 [Mus musculus]

| Query | Observed | Mr(expt) | Mr(calc) | Delta | Miss | Score | Expect | Rank | Unique | Peptide |
|---|---|---|---|---|---|---|---|---|---|---|
| 15600 | 739.0794 | 1476.1442 | 1475.7620 | 0.3822 | 0 | 60 | 0.00096 | 2 | U | R.FLEQQNQVLETK.W |

---

**13.** gi|16716569 **Mass:** 26118 **Score:** 59 **Matches:** 3(3) **Sequences:** 1(1) **emPAI:** 0.13
protease, serine, 1 precursor [Mus musculus]

| Query | Observed | Mr(expt) | Mr(calc) | Delta | Miss | Score | Expect | Rank | Unique | Peptide |
|---|---|---|---|---|---|---|---|---|---|---|
| 15580 | 738.0673 | 2211.1800 | 2210.0967 | 1.0832 | 0 | 52 | 0.014 | 1 | U | R.LGEHNINVLEGNEQFIDAAK.I 15575 15577 |

---

**14.** gi|1389682 **Mass:** 68068 **Score:** 59 **Matches:** 1(1) **Sequences:** 1(1) **emPAI:** 0.05
plakoglobin, partial [Mus musculus]

| Query | Observed | Mr(expt) | Mr(calc) | Delta | Miss | Score | Expect | Rank | Unique | Peptide |
|---|---|---|---|---|---|---|---|---|---|---|
| 15662 | 742.6564 | 1483.2982 | 1482.7467 | 0.5515 | 0 | 59 | 0.0027 | 1 | U | R.NEGTATYAAAVLFR.I |

**Proteins matching the same set of peptides:**
gi|28395018 **Mass:** 81749 **Score:** 59 **Matches:** 1(1) **Sequences:** 1(1)
junction plakoglobin [Mus musculus]

---

**15.** gi|19354359 **Mass:** 54698 **Score:** 59 **Matches:** 1(1) **Sequences:** 1(1) **emPAI:** 0.06
Ranbp10 protein [Mus musculus]

| Query | Observed | Mr(expt) | Mr(calc) | Delta | Miss | Score | Expect | Rank | Unique | Peptide |
|---|---|---|---|---|---|---|---|---|---|---|
| 11862 | 552.3564 | 1102.6982 | 1102.5618 | 0.1363 | 0 | 59 | 0.0018 | 1 | U | R.VGEAIETTQR.F |

**Proteins matching the same set of peptides:**
gi|50510939 **Mass:** 66515 **Score:** 59 **Matches:** 1(1) **Sequences:** 1(1)

mKIAA1464 protein [Mus musculus]
gi|353526275 **Mass:** 67146 **Score:** 59 **Matches:** 1(1) **Sequences:** 1(1)
RecName: Full=Ran-binding protein 10; Short=RanBP10
gi|40804757 **Mass:** 70044 **Score:** 59 **Matches:** 1(1) **Sequences:** 1(1)
ran-binding protein 10 [Mus musculus]

---

**16.** gi|52789 **Mass:** 54415 **Score:** 57 **Matches:** 1(1) **Sequences:** 1(1) **emPAI:** 0.06
unnamed protein product [Mus musculus]

| Query | Observed | Mr(expt) | Mr(calc) | Delta | Miss | Score | Expect | Rank | Unique | Peptide |
|---|---|---|---|---|---|---|---|---|---|---|
| 14685 | 692.1215 | 1382.2284 | 1382.7194 | -0.4910 | 1 | 57 | 0.0039 | 1 | U | K.SLNNKFASFIDK.V |

**Proteins matching the same set of peptides:**
gi|309215 **Mass:** 53210 **Score:** 57 **Matches:** 1(1) **Sequences:** 1(1)
EndoA' cytokeratin (5' end put.); putative [Mus musculus]
gi|511654 **Mass:** 54220 **Score:** 57 **Matches:** 1(1) **Sequences:** 1(1)
keratin type II [Mus musculus]
gi|74177777 **Mass:** 54546 **Score:** 57 **Matches:** 1(1) **Sequences:** 1(1)
unnamed protein product [Mus musculus]
gi|74219975 **Mass:** 54459 **Score:** 57 **Matches:** 1(1) **Sequences:** 1(1)
unnamed protein product [Mus musculus]
gi|76779293 **Mass:** 54514 **Score:** 57 **Matches:** 1(1) **Sequences:** 1(1)
Keratin 8 [Mus musculus]
gi|114145561 **Mass:** 54531 **Score:** 57 **Matches:** 1(1) **Sequences:** 1(1)
keratin, type II cytoskeletal 8 [Mus musculus]

---

**17.** gi|54777 **Mass:** 57108 **Score:** 56 **Matches:** 1(1) **Sequences:** 1(1) **emPAI:** 0.06
unnamed protein product [Mus musculus]

| Query | Observed | Mr(expt) | Mr(calc) | Delta | Miss | Score | Expect | Rank | Unique | Peptide |
|---|---|---|---|---|---|---|---|---|---|---|
| 13198 | **611.9623** | **1221.9101** | **1221.6162** | **0.2939** | **0** | **56** | **0.0059** | **1** | **U** | **R.LITLEEEMTK.Y** |

**Proteins matching the same set of peptides:**
gi|42415475 **Mass:** 57023 **Score:** 56 **Matches:** 1(1) **Sequences:** 1(1)
protein disulfide-isomerase precursor [Mus musculus]
gi|74141920 **Mass:** 57053 **Score:** 56 **Matches:** 1(1) **Sequences:** 1(1)
unnamed protein product [Mus musculus]
gi|74149659 **Mass:** 43005 **Score:** 56 **Matches:** 1(1) **Sequences:** 1(1)
unnamed protein product [Mus musculus]
gi|74178069 **Mass:** 57099 **Score:** 56 **Matches:** 1(1) **Sequences:** 1(1)
unnamed protein product [Mus musculus]
gi|74190076 **Mass:** 56963 **Score:** 56 **Matches:** 1(1) **Sequences:** 1(1)
unnamed protein product [Mus musculus]
gi|74198312 **Mass:** 57051 **Score:** 56 **Matches:** 1(1) **Sequences:** 1(1)
unnamed protein product [Mus musculus]
gi|74198706 **Mass:** 57000 **Score:** 56 **Matches:** 1(1) **Sequences:** 1(1)
unnamed protein product [Mus musculus]
gi|74203945 **Mass:** 56566 **Score:** 56 **Matches:** 1(1) **Sequences:** 1(1)
unnamed protein product [Mus musculus]
gi|74212231 **Mass:** 57053 **Score:** 56 **Matches:** 1(1) **Sequences:** 1(1)
unnamed protein product [Mus musculus]
gi|74219772 **Mass:** 57008 **Score:** 56 **Matches:** 1(1) **Sequences:** 1(1)
unnamed protein product [Mus musculus]
gi|74220649 **Mass:** 57067 **Score:** 56 **Matches:** 1(1) **Sequences:** 1(1)
unnamed protein product [Mus musculus]
gi|148702818 **Mass:** 58984 **Score:** 56 **Matches:** 1(1) **Sequences:** 1(1)
prolyl 4-hydroxylase, beta polypeptide, isoform CRA_a [Mus musculus]
gi|148702819 **Mass:** 61492 **Score:** 56 **Matches:** 1(1) **Sequences:** 1(1)
prolyl 4-hydroxylase, beta polypeptide, isoform CRA_b [Mus musculus]

---

**18.** gi|20070408 **Mass:** 113195 **Score:** 55 **Matches:** 1(1) **Sequences:** 1(1) **emPAI:** 0.03

glycine dehydrogenase [decarboxylating], mitochondrial precursor [Mus musculus]

| Query | Observed | Mr(expt) | Mr(calc) | Delta | Miss | Score | Expect | Rank | Unique | Peptide |
|---|---|---|---|---|---|---|---|---|---|---|
| 11734 | 546.4593 | 1090.9039 | 1091.4228 | -0.5189 | 0 | 55 | 0.0027 | 1 | U | R.DSSGGGGGGGGGDR.G |

**Proteins matching the same set of peptides:**

gi\|148709754 **Mass:** 30341 **Score:** 55 **Matches:** 1(1) **Sequences:** 1(1)
glycine decarboxylase, isoform CRA_c [Mus musculus]
gi\|148709755 **Mass:** 114180 **Score:** 55 **Matches:** 1(1) **Sequences:** 1(1)
glycine decarboxylase, isoform CRA_d [Mus musculus]
gi\|148709756 **Mass:** 98319 **Score:** 55 **Matches:** 1(1) **Sequences:** 1(1)
glycine decarboxylase, isoform CRA_e [Mus musculus]

---

**19.** gi|3642689 **Mass:** 9181 **Score:** 54 **Matches:** 1(1) **Sequences:** 1(1) **emPAI:** 0.38
immunoglobulin heavy chain-binding protein, partial [Mus musculus]

| Query | Observed | Mr(expt) | Mr(calc) | Delta | Miss | Score | Expect | Rank | Unique | Peptide |
|---|---|---|---|---|---|---|---|---|---|---|
| 15298 | 722.9624 | 1443.9102 | 1443.6994 | 0.2108 | 0 | 54 | 0.0046 | 1 | U | -.TWNDPTVQQDIK.F |

---

**20.** gi|26344548 **Mass:** 58526 **Score:** 54 **Matches:** 1(1) **Sequences:** 1(1) **emPAI:** 0.06
unnamed protein product [Mus musculus]

| Query | Observed | Mr(expt) | Mr(calc) | Delta | Miss | Score | Expect | Rank | Unique | Peptide |
|---|---|---|---|---|---|---|---|---|---|---|
| 11185 | 521.0774 | 1040.1402 | 1039.6390 | 0.5013 | 0 | 54 | 0.003 | 1 | U | K.VLEALQVLR.C |

**Proteins matching the same set of peptides:**

gi|35193112 **Mass:** 55286 **Score:** 54 **Matches:** 1(1) **Sequences:** 1(1)
Wdr26 protein [Mus musculus]
gi|148681181 **Mass:** 58625 **Score:** 54 **Matches:** 1(1) **Sequences:** 1(1)
WD repeat domain 26 [Mus musculus]
gi|264681550 **Mass:** 70499 **Score:** 54 **Matches:** 1(1) **Sequences:** 1(1)
WD repeat-containing protein 26 [Mus musculus]

---

**21.** gi|836898 **Mass:** 59960 **Score:** 54 **Matches:** 2(1) **Sequences:** 2(1) **emPAI:** 0.11
prolyl 4-hydroxylase alpha(I)-subunit, partial [Mus musculus]

| Query | Observed | Mr(expt) | Mr(calc) | Delta | Miss | Score | Expect | Rank | Unique | Peptide |
|---|---|---|---|---|---|---|---|---|---|---|
| 12213 | 567.0950 | 1132.1754 | 1131.5924 | 0.5830 | 0 | 49 | 0.011 | 1 | U | K.GIAVDYLPER.Q |
| 13453 | 625.1786 | 1248.3426 | 1248.6462 | -0.3036 | 0 | 43 | 0.091 | 1 | U | K.LLELDPEHQR.A |

**Proteins matching the same set of peptides:**

gi|26336999 **Mass:** 60848 **Score:** 54 **Matches:** 2(1) **Sequences:** 2(1)
unnamed protein product [Mus musculus]
gi|33859596 **Mass:** 60872 **Score:** 54 **Matches:** 2(1) **Sequences:** 2(1)
prolyl 4-hydroxylase subunit alpha-1 precursor [Mus musculus]
gi|74224984 **Mass:** 60886 **Score:** 54 **Matches:** 2(1) **Sequences:** 2(1)
unnamed protein product [Mus musculus]
gi|74225936 **Mass:** 63769 **Score:** 54 **Matches:** 2(1) **Sequences:** 2(1)
unnamed protein product [Mus musculus]

---

**22.** gi|4159806 **Mass:** 65183 **Score:** 52 **Matches:** 1(1) **Sequences:** 1(1) **emPAI:** 0.05
type II keratin subunit protein [Mus musculus]

| Query | Observed | Mr(expt) | Mr(calc) | Delta | Miss | Score | Expect | Rank | Unique | Peptide |
|---|---|---|---|---|---|---|---|---|---|---|
| 13588 | 633.1061 | 1264.1977 | 1264.6299 | -0.4322 | 0 | 52 | 0.01 | 1 | U | R.TNAENEFVTIK.K |

**Proteins matching the same set of peptides:**

gi|12859782 **Mass:** 65586 **Score:** 52 **Matches:** 1(1) **Sequences:** 1(1)
unnamed protein product [Mus musculus]
gi|126116585 **Mass:** 65565 **Score:** 52 **Matches:** 1(1) **Sequences:** 1(1)
keratin, type II cytoskeletal 1 [Mus musculus]

---

**23.** gi|37935731 **Mass:** 192078 **Score:** 52 **Matches:** 1(1) **Sequences:** 1(1) **emPAI:** 0.02
ATP-binding cassette transporter sub-family A member 16 [Mus musculus]

| Query | Observed | Mr(expt) | Mr(calc) | Delta | Miss | Score | Expect | Rank | Unique | Peptide |
|---|---|---|---|---|---|---|---|---|---|---|
| 13521 | 629.4885 | 1256.9625 | 1256.4835 | 0.4790 | 0 | 52 | 0.012 | 1 | U | R.MEECEALCTR.L |

**Proteins matching the same set of peptides:**
gi|148685261 **Mass:** 167887 **Score:** 52 **Matches:** 1(1) **Sequences:** 1(1)
ATP-binding cassette, sub-family A (ABC1), member 16 [Mus musculus]

**24.** gi|21595163 **Mass:** 71011 **Score:** 48 **Matches:** 1(1) **Sequences:** 1(1) **emPAI:** 0.05
Glycosyltransferase 25 domain containing 1 [Mus musculus]

| Query | Observed | Mr(expt) | Mr(calc) | Delta | Miss | Score | Expect | Rank | Unique | Peptide |
|---|---|---|---|---|---|---|---|---|---|---|
| 9480 | 435.0096 | 868.0046 | 867.5905 | 0.4140 | 0 | 48 | 0.0098 | 1 | U | R.VLIALLAR.N |

**Proteins matching the same set of peptides:**
gi|148697003 **Mass:** 54889 **Score:** 48 **Matches:** 1(1) **Sequences:** 1(1)
glycosyltransferase 25 domain containing 1, isoform CRA_b [Mus musculus]
gi|170784829 **Mass:** 71015 **Score:** 48 **Matches:** 1(1) **Sequences:** 1(1)
procollagen galactosyltransferase 1 precursor [Mus musculus]
gi|74217150 **Mass:** 70997 **Score:** 48 **Matches:** 1(1) **Sequences:** 1(1)
unnamed protein product [Mus musculus]

**25.** gi|12847276 **Score:** 48 **Matches:** 1(1) **Sequences:** 1(1) **emPAI:** 0.07
unnamed protein product [Mus musculus]

| Query | Observed | Mr(expt) | Mr(calc) | Delta | Miss | Score | Expect | Rank | Unique | Peptide |
|---|---|---|---|---|---|---|---|---|---|---|
| 15600 | 739.0794 | 1476.1442 | 1475.7442 | 0.4000 | 1 | 48 | 0.013 | 3 | U | R.YEPKCPLGVDISR.E |

**Proteins matching the same set of peptides:**
gi|31560081 **Score:** 48 **Matches:** 1(1) **Sequences:** 1(1)
gi|148667576 **Score:** 48 **Matches:** 1(1) **Sequences:** 1(1)

**26.** gi|74191500 **Mass:** 57022 **Score:** 46 **Matches:** 1(1) **Sequences:** 1(1) **emPAI:** 0.06
unnamed protein product [Mus musculus]

| Query | Observed | Mr(expt) | Mr(calc) | Delta | Miss | Score | Expect | Rank | Unique | Peptide |
|---|---|---|---|---|---|---|---|---|---|---|
| 13198 | 611.9623 | 1221.9101 | 1220.6686 | 1.2415 | 1 | 46 | 0.05 | 2 | U | R.LITLEEKMTK.Y |

**27.** gi|81875970 **Mass:** 71638 **Score:** 45 **Matches:** 1(0) **Sequences:** 1(0) **emPAI:** 0.05
RecName: Full=Alkyldihydroxyacetonephosphate synthase, peroxisomal; Short=Alkyl-DHAP synthase; AltName: Full=Alkylglycerone-phosphate synthase; Flags: Precursor

| Query | Observed | Mr(expt) | Mr(calc) | Delta | Miss | Score | Expect | Rank | Unique | Peptide |
|---|---|---|---|---|---|---|---|---|---|---|
| 15186 | 717.2085 | 2148.6037 | 2149.1127 | -0.5090 | 1 | 45 | 0.06 | 1 | U | R.RAASAAGASPAATPAAPESGTIPK.K |

**Proteins matching the same set of peptides:**
gi|148695250 **Mass:** 53219 **Score:** 45 **Matches:** 1(0) **Sequences:** 1(0)
alkylglycerone phosphate synthase, isoform CRA_a [Mus musculus]
gi|295444834 **Mass:** 74296 **Score:** 45 **Matches:** 1(0) **Sequences:** 1(0)
alkyldihydroxyacetonephosphate synthase, peroxisomal [Mus musculus]

**28.** gi|7513694 **Mass:** 109137 **Score:** 44 **Matches:** 1(0) **Sequences:** 1(0) **emPAI:** 0.03
IgG Fc binding protein - mouse (fragment)

| Query | Observed | Mr(expt) | Mr(calc) | Delta | Miss | Score | Expect | Rank | Unique | Peptide |
|---|---|---|---|---|---|---|---|---|---|---|
| 9539 | 874.3927 | 873.3854 | 873.4920 | -0.1066 | 0 | 44 | 0.06 | 1 | U | R.ISVINGGSK.A |

**Proteins matching the same set of peptides:**
gi|148692210 **Mass:** 147962 **Score:** 44 **Matches:** 1(0) **Sequences:** 1(0)
mCG145390, isoform CRA_a [Mus musculus]
gi|148692211 **Mass:** 146729 **Score:** 44 **Matches:** 1(0) **Sequences:** 1(0)

mCG145390, isoform CRA_b [Mus musculus]
gi|169790797 **Mass:** 275058 **Score:** 44 **Matches:** 1(0) **Sequences:** 1(0)
Fc fragment of IgG binding protein precursor [Mus musculus]
gi|257467625 **Mass:** 280044 **Score:** 44 **Matches:** 1(0) **Sequences:** 1(0)
Fc fragment of IgG binding protein-like precursor [Mus musculus]

---

**29.** gi|148664969 **Mass:** 26715 **Score:** 44 **Matches:** 1(1) **Sequences:** 1(1) **emPAI:** 0.12
RIKEN cDNA 2900011O08, isoform CRA_a [Mus musculus]

| Query | Observed | Mr(expt) | Mr(calc) | Delta | Miss | Score | Expect | Rank | Unique | Peptide |
|---|---|---|---|---|---|---|---|---|---|---|
| 12819 | 593.2423 | 1184.4701 | 1185.5812 | -1.1111 | 0 | 44 | 0.053 | 1 | U | -.NTADLSCLPPR.S |

---

**30.** gi|12855410 **Mass:** 22073 **Score:** 43 **Matches:** 1(1) **Sequences:** 1(1) **emPAI:** 0.15
unnamed protein product [Mus musculus]

| Query | Observed | Mr(expt) | Mr(calc) | Delta | Miss | Score | Expect | Rank | Unique | Peptide |
|---|---|---|---|---|---|---|---|---|---|---|
| 13489 | 627.3521 | 1252.6896 | 1252.6526 | 0.0370 | 1 | 43 | 0.056 | 1 | U | R.EKMVVFFPPK.E |

---

**Mascot:** http://www.matrixscience.com/

PFHR9

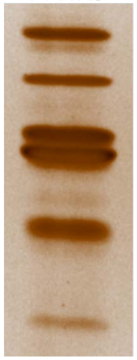

3

### Mascot Search Results

| | | |
|---|---|---|
| User | : | |
| Email | : | |
| Search title | : | |
| MS data file | : | ZIEN-29Nov2017_3m.mgf |
| Database | : | NCBInr 20120419 (17893860 sequences; 6141683785 residues) |
| Taxonomy | : | Mus musculus (house mouse) (138831 sequences) |
| Timestamp | : | 30 Nov 2017 at 20:39:15 GMT |
| Enzyme | : | Trypsin |
| Variable modifications | : | Carbamidomethyl (C),Oxidation (M),Oxidation (P) |
| Mass values | : | Monoisotopic |
| Protein Mass | : | Unrestricted |
| Peptide Mass Tolerance | : | ± 1.25 Da |
| Fragment Mass Tolerance | : | ± 1.001 Da |
| Max Missed Cleavages | : | 1 |
| Instrument type | : | Default |
| Number of queries | : | 19332 |
| Protein hits | : | gi\|54777 unnamed protein product [Mus musculus] |

| | |
|---|---|
| gi\|54777 | unnamed protein product [Mus musculus] |
| gi\|148701597 | procollagen-proline, 2-oxoglutarate 4-dioxygenase (proline 4-hydroxylase), alpha II polypeptide, isoform CRA_b [Mus musculus] |
| gi\|148701600 | procollagen-proline, 2-oxoglutarate 4-dioxygenase (proline 4-hydroxylase), alpha II polypeptide, isoform CRA_e [Mus musculus] |
| gi\|74225936 | unnamed protein product [Mus musculus] |
| gi\|16716569 | protease, serine, 1 precursor [Mus musculus] |
| gi\|347839 | matricin [Mus musculus] |
| gi\|21312570 | protein ERGIC-53 precursor [Mus musculus] |
| gi\|387398 | epidermal keratin type I, partial [Mus musculus] |
| gi\|51452 | unnamed protein product [Mus musculus] |
| gi\|74213832 | unnamed protein product [Mus musculus] |
| gi\|22164776 | keratin, type II cytoskeletal 79 [Mus musculus] |
| gi\|12843914 | unnamed protein product [Mus musculus] |
| gi\|5295992 | chaperonin containing TCP-1 theta subunit [Mus musculus] |
| gi\|1419012 | CCT zeta2, zeta2 subunit of the chaperonin containing TCP-1 (CCT) [Mus musculus] |
| gi\|7710044 | kelch-like ECH-associated protein 1 [Mus musculus] |
| gi\|2623222 | ATP synthase beta-subunit [Mus musculus] |
| gi\|192262 | pro-alpha-1 type I collagen, partial [Mus musculus] |
| gi\|551295 | pyruvate kinase M [Mus musculus] |
| gi\|52789 | unnamed protein product [Mus musculus] |
| gi\|46485130 | TPA_exp: keratin Kb40 [Mus musculus] |
| gi\|1304157 | 78 kDa glucose-regulated protein [Mus musculus] |
| gi\|4159806 | type II keratin subunit protein [Mus musculus] |
| gi\|293686 | epidermal keratin subunit II [Mus musculus] |
| gi\|3687320 | cartilage-associated protein (CASP) [Mus musculus] |
| gi\|74203983 | unnamed protein product [Mus musculus] |
| gi\|468546 | CCT (chaperonin containing TCP-1) beta subunit [Mus musculus] |
| gi\|22297268 | AID, partial [Homo sapiens] |
| gi\|51450 | heat shock protein [Mus musculus] |
| gi\|6680748 | ATP synthase subunit alpha, mitochondrial precursor [Mus musculus] |
| gi\|148688671 | mCG16097 [Mus musculus] |
| gi\|6755893 | trypsin 4 precursor [Mus musculus] |
| gi\|148707748 | nuclear casein kinase and cyclin-dependent kinase substrate 1, isoform CRA_b [Mus musculus] |
| gi\|53830774 | lactase [Mus musculus] |
| gi\|26324736 | unnamed protein product [Mus musculus] |
| gi\|7513694 | IgG Fc binding protein - mouse (fragment) |
| gi\|111599858 | Dehydrogenase E1 and transketolase domain containing 1 [Mus musculus] |
| gi\|74224513 | unnamed protein product [Mus musculus] |
| gi\|21410133 | Kidins220 protein [Mus musculus] |
| gi\|26333649 | unnamed protein product [Mus musculus] |
| gi\|986918 | A10 [Mus musculus] |
| gi\|21313084 | chromodomain Y-like protein 2 [Mus musculus] |
| gi\|124504325 | Armcx4 protein [Mus musculus] |
| gi\|12843046 | unnamed protein product [Mus musculus] |
| gi\|12836661 | unnamed protein product [Mus musculus] |

gi|148681245 mCG145199 [Mus musculus]
gi|148667455 TEA domain family member 4, isoform CRA_b [Mus musculus]
gi|642072 fibrillin-1, partial [Mus musculus]
gi|12836791 unnamed protein product [Mus musculus]
gi|14250408 Aspartyl-tRNA synthetase [Mus musculus]
gi|17512475 Ankyrin repeat and SOCS box-containing 6 [Mus musculus]
gi|309271872 PREDICTED: zinc finger protein 469 [Mus musculus]
gi|26380166 unnamed protein product [Mus musculus]

#### Select Summary Report

Format As | Select Summary (protein hits) | Help

Significance threshold p< 0.05 Max. number of hits AUTO

Standard scoring ○ MudPIT scoring ◉ Ions score or expect cut-off 42 Show sub-sets 0

Show pop-ups ◉ Suppress pop-ups ○ Require bold red ☐

Re-Search ◉ All queries ○ Unassigned ○ Below homology threshold ○ Below identity threshold

**1.** gi|54777 **Mass:** 57108 **Score:** 426 **Matches:** 28(25) **Sequences:** 16(15) **emPAI:** 1.90
unnamed protein product [Mus musculus]

| Query | Observed | Mr(expt) | Mr(calc) | Delta | Miss | Score | Expect | Rank | Unique | Peptide |
|---|---|---|---|---|---|---|---|---|---|---|
| 10277 | 381.1882 | 760.3618 | 760.3715 | -0.0097 | 0 | 50 | 0.015 | 1 | U | K.AEGSEIR.L |
| 10911 | 395.3300 | 788.6454 | 788.3664 | 0.2790 | 0 | 59 | 0.0018 | 1 | U | K.NGDTASPK.E |
| 13577 | 483.6877 | 965.3609 | 965.5586 | -0.1977 | 0 | 55 | 0.0037 | 1 | U | R.ILEFFGLK.K |
| 14327 | 1051.4263 | 1050.4190 | 1050.4982 | -0.0791 | 0 | 51 | 0.0088 | 1 | U | R.NNFEGEITK.E |
| 14499 | 1067.7178 | 1066.7105 | 1065.5091 | 1.2014 | 0 | 45 | 0.033 | 1 | U | R.TVIDYNGER.T |
| 14641 | 1081.6606 | 1080.6533 | 1080.6695 | -0.0162 | 0 | 52 | 0.007 | 1 | U | K.THILLFLPK.S 14649 |
| 14648 | 541.6124 | 1081.2102 | 1080.6695 | 0.5406 | 0 | (48) | 0.012 | 1 | U | K.THILLFLPK.S 14646 |
| 16005 | 1203.5746 | 1202.5673 | 1201.5979 | 0.9694 | 0 | 49 | 0.017 | 1 | U | R.EADDIVNWLK.K |
| 16043 | 604.0506 | 1206.0866 | 1205.6213 | 0.4653 | 0 | 51 | 0.0064 | 1 | U | R.LITLEEEMTK.Y 16046 |
| 16166 | 608.9402 | 1215.8658 | 1215.5884 | 0.2775 | 0 | 47 | 0.025 | 1 | U | K.SNFEEALAAHK.Y |
| 16232 | 611.6342 | 1221.2538 | 1221.6162 | -0.3625 | 0 | (44) | 0.088 | 1 | U | R.LITLEEEMTK.Y |
| 16991 | 647.9220 | 1293.8294 | 1292.5918 | 1.2376 | 0 | 84 | 4.2e-006 | 1 | U | K.MDSTANEVEAVK.V 16980 16985 16986 16990 |
| 17115 | 654.5397 | 1307.0648 | 1307.6357 | -0.5709 | 1 | 46 | 0.043 | 1 | U | R.NNFEGEITKEK.L |
| 18095 | 475.6379 | 1423.8917 | 1423.7711 | 0.1206 | 1 | (46) | 0.031 | 1 | U | K.YQLDKDGVVLFK.K |
| 18099 | 713.1129 | 1424.2113 | 1423.7711 | 0.4402 | 1 | 55 | 0.0069 | 1 | U | K.YQLDKDGVVLFK.K 18100 |
| 18159 | 717.4308 | 1432.8470 | 1431.6882 | 1.1589 | 1 | 63 | 0.0011 | 1 | U | K.SVSDYDGKLSSFK.R |
| 15815 | 594.4030 | 1780.1872 | 1779.8275 | 0.3597 | 0 | (61) | 0.0019 | 1 | U | K.VDATEESDLAQQYGVR.G |
| 19294 | 891.2250 | 1780.4354 | 1779.8275 | 0.6079 | 0 | 78 | 2.7e-005 | 1 | U | K.VDATEESDLAQQYGVR.G |
| 16231 | 611.6050 | 1831.7933 | 1832.9057 | -1.1124 | 0 | 43 | 0.1 | 1 | U | K.ILFIFIDSDHTDNQR.I |
| 19326 | 1493.5814 | 2985.1482 | 2984.4866 | 0.6616 | 0 | 98 | 2.4e-007 | 1 | U | R.TGPAATTLSDTAAAESLVDSSEVTVIGFFK.D |

**Proteins matching the same set of peptides:**
gi|74212231 **Mass:** 57053 **Score:** 426 **Matches:** 28(25) **Sequences:** 16(15)
unnamed protein product [Mus musculus]
gi|74190076 **Mass:** 56963 **Score:** 426 **Matches:** 28(25) **Sequences:** 16(15)
unnamed protein product [Mus musculus]
gi|42415475 **Mass:** 57023 **Score:** 426 **Matches:** 28(25) **Sequences:** 16(15)
protein disulfide-isomerase precursor [Mus musculus]
gi|74141920 **Mass:** 57053 **Score:** 426 **Matches:** 28(25) **Sequences:** 16(15)
unnamed protein product [Mus musculus]

gi|74178069 **Mass:** 57099 **Score:** 426 **Matches:** 28(25) **Sequences:** 16(15)
unnamed protein product [Mus musculus]
gi|74198312 **Mass:** 57051 **Score:** 426 **Matches:** 28(25) **Sequences:** 16(15)
unnamed protein product [Mus musculus]
gi|74198706 **Mass:** 57000 **Score:** 426 **Matches:** 28(25) **Sequences:** 16(15)
unnamed protein product [Mus musculus]
gi|74219772 **Mass:** 57008 **Score:** 426 **Matches:** 28(25) **Sequences:** 16(15)
unnamed protein product [Mus musculus]
gi|148702818 **Mass:** 58984 **Score:** 426 **Matches:** 28(25) **Sequences:** 16(15)
prolyl 4-hydroxylase, beta polypeptide, isoform CRA_a [Mus musculus]
gi|148702819 **Mass:** 61492 **Score:** 426 **Matches:** 28(25) **Sequences:** 16(15)
prolyl 4-hydroxylase, beta polypeptide, isoform CRA_b [Mus musculus]

---

**2.** gi|148701597 **Mass:** 57417 **Score:** 223 **Matches:** 17(15) **Sequences:** 10(10) **emPAI:** 0.75
procollagen-proline, 2-oxoglutarate 4-dioxygenase (proline 4-hydroxylase), alpha II polypeptide, isoform CRA_b [Mus musculus]

| Query | Observed | Mr(expt) | Mr(calc) | Delta | Miss | Score | Expect | Rank | Unique | Peptide |
|---|---|---|---|---|---|---|---|---|---|---|
| 13332 | **470.4389** | **938.8633** | **938.3981** | **0.4651** | **0** | **51** | **0.006** | **1** | **U** | **R.SDEQDAFK.R** |
| 14757 | **1093.6360** | **1092.6287** | **1092.5703** | **0.0585** | **0** | **44** | **0.054** | **1** | | **K.EYILVEEAK.L** 14753 |
| 14783 | **548.5247** | **1095.0348** | **1094.4992** | **0.5356** | **1** | **44** | **0.031** | **1** | **U** | **R.SDEQDAFKR.L** |
| 15163 | **1125.7443** | **1124.7370** | **1124.6958** | **0.0413** | **0** | **46** | **0.028** | **1** | | **R.VPQLLIAPFK.E** |
| 15611 | **583.9658** | **1165.9171** | **1165.6091** | **0.3080** | **0** | **48** | **0.013** | **1** | | **R.LLSLDPSHER.A** |
| 16687 | **631.0879** | **1260.1612** | **1260.6198** | **-0.4585** | **0** | **68** | **0.00026** | **1** | | **K.QLDAGEEATVTK.S** 16695 16699 |
| 18396 | **734.0641** | **1466.1136** | **1466.8034** | **-0.6898** | **1** | **59** | **0.0027** | **1** | | **K.KGTAVFWYNLLR.S** 18398 |
| 18833 | **774.6111** | **1547.2077** | **1547.6450** | **-0.4372** | **0** | **55** | **0.0061** | **1** | | **R.YYDVMSDEEIER.I** 18840 18842 |
| 19065 | **805.0440** | **1608.0735** | **1607.7250** | **0.3485** | **0** | **47** | **0.046** | **1** | | **R.GQEFLRPCGTTEVD.-** 19066 |
| 19092 | **809.5747** | **1617.1349** | **1616.7318** | **0.4030** | **0** | **69** | **0.00021** | **1** | | **K.SSWLEEDDDPVVAR.V** |

**Proteins matching the same set of peptides:**
gi|226874876 **Mass:** 60977 **Score:** 223 **Matches:** 17(15) **Sequences:** 10(10)
prolyl 4-hydroxylase subunit alpha-2 isoform 2 precursor [Mus musculus]

---

**3.** gi|148701600 **Mass:** 66870 **Score:** 216 **Matches:** 16(14) **Sequences:** 9(9) **emPAI:** 0.54
procollagen-proline, 2-oxoglutarate 4-dioxygenase (proline 4-hydroxylase), alpha II polypeptide, isoform CRA_e [Mus musculus]

| Query | Observed | Mr(expt) | Mr(calc) | Delta | Miss | Score | Expect | Rank | Unique | Peptide |
|---|---|---|---|---|---|---|---|---|---|---|
| 13143 | **460.4962** | **918.9779** | **918.4923** | **0.4856** | **0** | **47** | **0.016** | **1** | **U** | **R.RPFDSGLK.T** |
| 14757 | 1093.6360 | 1092.6287 | 1092.5703 | 0.0585 | 0 | 44 | 0.054 | 1 | | K.EYILVEEAK.L 14753 |
| 15163 | 1125.7443 | 1124.7370 | 1124.6958 | 0.0413 | 0 | 46 | 0.028 | 1 | | R.VPQLLIAPFK.E |
| 15611 | 583.9658 | 1165.9171 | 1165.6091 | 0.3080 | 0 | 48 | 0.013 | 1 | | R.LLSLDPSHER.A |
| 16687 | 631.0879 | 1260.1612 | 1260.6198 | -0.4585 | 0 | 68 | 0.00026 | 1 | | K.QLDAGEEATVTK.S 16695 16699 |
| 18396 | 734.0641 | 1466.1136 | 1466.8034 | -0.6898 | 1 | 59 | 0.0027 | 1 | | K.KGTAVFWYNLLR.S 18398 |
| 18833 | 774.6111 | 1547.2077 | 1547.6450 | -0.4372 | 0 | 55 | 0.0061 | 1 | | R.YYDVMSDEEIER.I 18840 18842 |
| 19065 | 805.0440 | 1608.0735 | 1607.7250 | 0.3485 | 0 | 47 | 0.046 | 1 | | R.GQEFLRPCGTTEVD.- 19066 |
| 19092 | 809.5747 | 1617.1349 | 1616.7318 | 0.4030 | 0 | 69 | 0.00021 | 1 | | K.SSWLEEDDDPVVAR.V |

**Proteins matching the same set of peptides:**
gi|209862961 **Mass:** 60774 **Score:** 216 **Matches:** 16(14) **Sequences:** 9(9)
prolyl 4-hydroxylase subunit alpha-2 isoform 1 precursor [Mus musculus]

---

**4.** gi|74225936 **Mass:** 63769 **Score:** 160 **Matches:** 10(8) **Sequences:** 8(6) **emPAI:** 0.42
unnamed protein product [Mus musculus]

| Query | Observed | Mr(expt) | Mr(calc) | Delta | Miss | Score | Expect | Rank | Unique | Peptide |
|---|---|---|---|---|---|---|---|---|---|---|
| 13568 | 483.3983 | 964.7821 | 964.5090 | 0.2731 | 0 | 50 | 0.0085 | 1 | U | K.GNLPGVQHK.S |
| 14305 | 524.3110 | 1046.6075 | 1045.5590 | 1.0485 | 0 | 44 | 0.11 | 1 | U | R.RPCTLSELK.S |
| 15244 | 1132.6241 | 1131.6168 | 1131.5924 | 0.0244 | 0 | 44 | 0.05 | 1 | U | K.GIAVDYLPER.Q |
| 15607 | 583.4688 | 1164.9231 | 1164.6074 | 0.3157 | 0 | 68 | 0.00014 | 1 | U | R.HAACPVLVGNK.W |
| 16994 | 648.0558 | 1294.0970 | 1293.7180 | 0.3790 | 1 | 47 | 0.017 | 1 | U | K.DLVTSLKDYIK.A |
| 18891 | 783.3275 | 1564.6405 | 1564.7522 | -0.1117 | 0 | 53 | 0.013 | 1 | U | K.SAWLSGYEDPVVSR.I |
| 17262 | 662.6197 | 1984.8374 | 1984.0490 | 0.7884 | 1 | 44 | 0.076 | 1 | U | R.RATISNPVTGALETVHYR.I |
| 18487 | 739.6022 | 2215.7847 | 2215.1274 | 0.6573 | 1 | 58 | 0.0028 | 1 | U | R.LTSTATKDPEGFVGHPVNAFK.L 18476 18483 |

5. gi|16716569 **Mass:** 26118 **Score:** 136 **Matches:** 5(4) **Sequences:** 1(1) **emPAI:** 0.27
protease, serine, 1 precursor [Mus musculus]

| Query | Observed | Mr(expt) | Mr(calc) | Delta | Miss | Score | Expect | Rank | Unique | Peptide |
|---|---|---|---|---|---|---|---|---|---|---|
| 18449 | 737.5288 | 2209.5646 | 2210.0967 | -0.5321 | 0 | (67) | 0.00036 | 1 | U | R.LGEHNINVLEGNEQFIDAAK.I 18450 18452 18459 |
| 19321 | 1106.1886 | 2210.3626 | 2210.0967 | 0.2659 | 0 | 78 | 2.9e-005 | 1 | U | R.LGEHNINVLEGNEQFIDAAK.I |

6. gi|347839 **Mass:** 60450 **Score:** 101 **Matches:** 2(2) **Sequences:** 2(2) **emPAI:** 0.11
matricin [Mus musculus]

| Query | Observed | Mr(expt) | Mr(calc) | Delta | Miss | Score | Expect | Rank | Unique | Peptide |
|---|---|---|---|---|---|---|---|---|---|---|
| 15211 | 565.4988 | 1128.9830 | 1128.6251 | 0.3579 | 1 | 73 | 4.7e-005 | 1 | U | R.KVQSGNINAAK.T |
| 16416 | 619.2510 | 1236.4874 | 1236.6094 | -0.1220 | 0 | 70 | 0.00023 | 1 | U | R.MALDDMISTLK.K |

**Proteins matching the same set of peptides:**
gi|6753320 **Mass:** 60591 **Score:** 101 **Matches:** 2(2) **Sequences:** 2(2)
T-complex protein 1 subunit gamma [Mus musculus]
gi|74142612 **Mass:** 58241 **Score:** 101 **Matches:** 2(2) **Sequences:** 2(2)
unnamed protein product [Mus musculus]
gi|74226937 **Mass:** 60563 **Score:** 101 **Matches:** 2(2) **Sequences:** 2(2)
unnamed protein product [Mus musculus]

7. gi|21312570 **Mass:** 57753 **Score:** 91 **Matches:** 3(3) **Sequences:** 2(2) **emPAI:** 0.12
protein ERGIC-53 precursor [Mus musculus]

| Query | Observed | Mr(expt) | Mr(calc) | Delta | Miss | Score | Expect | Rank | Unique | Peptide |
|---|---|---|---|---|---|---|---|---|---|---|
| 16899 | 642.4886 | 1282.9626 | 1282.6405 | 0.3222 | 0 | 66 | 0.00044 | 1 | U | R.YVSSLTEEISR.R 16889 |
| 18234 | 723.1138 | 2166.3195 | 2165.9978 | 0.3217 | 0 | 50 | 0.019 | 1 | U | K.GHPDLQGQPADDIFESIGDR.E |

**Proteins matching the same set of peptides:**
gi|74152119 **Mass:** 61236 **Score:** 91 **Matches:** 3(3) **Sequences:** 2(2)
unnamed protein product [Mus musculus]

8. gi|387398 **Mass:** 41454 **Score:** 83 **Matches:** 2(2) **Sequences:** 1(1) **emPAI:** 0.08
epidermal keratin type I, partial [Mus musculus]

| Query | Observed | Mr(expt) | Mr(calc) | Delta | Miss | Score | Expect | Rank | Unique | Peptide |
|---|---|---|---|---|---|---|---|---|---|---|
| 14151 | 515.5348 | 1029.0550 | 1028.5866 | 0.4684 | 0 | 66 | 0.00022 | 1 | U | R.VLDELTLAR.A 14152 |

**Proteins matching the same set of peptides:**
gi|741022 **Mass:** 49086 **Score:** 83 **Matches:** 2(2) **Sequences:** 1(1)
keratin 15
gi|904215 **Mass:** 49129 **Score:** 83 **Matches:** 2(2) **Sequences:** 1(1)

cytokeratin 15 [Mus musculus]
gi|1083398 **Mass:** 7958 **Score:** 83 **Matches:** 2(2) **Sequences:** 1(1)
keratin 15, type I, cytoskeletal - mouse (fragments)
gi|6680606 **Mass:** 44515 **Score:** 83 **Matches:** 2(2) **Sequences:** 1(1)
keratin, type I cytoskeletal 19 [Mus musculus]
gi|7106335 **Mass:** 48132 **Score:** 83 **Matches:** 2(2) **Sequences:** 1(1)
keratin, type I cytoskeletal 17 [Mus musculus]
gi|13097093 **Mass:** 30629 **Score:** 83 **Matches:** 2(2) **Sequences:** 1(1)
Krt14 protein [Mus musculus]
gi|21489935 **Mass:** 52834 **Score:** 83 **Matches:** 2(2) **Sequences:** 1(1)
keratin, type I cytoskeletal 14 [Mus musculus]
gi|38503465 **Mass:** 50146 **Score:** 83 **Matches:** 2(2) **Sequences:** 1(1)
keratin 17n [Mus musculus]
gi|83304265 **Mass:** 49107 **Score:** 83 **Matches:** 2(2) **Sequences:** 1(1)
RecName: Full=Keratin, type I cytoskeletal 15; AltName: Full=Cytokeratin-15; Short=CK-15; AltName: Full=Keratin-15; Short=K15
gi|85726518 **Mass:** 49857 **Score:** 83 **Matches:** 2(2) **Sequences:** 1(1)
Krt42 protein [Mus musculus]
gi|148670626 **Mass:** 54668 **Score:** 83 **Matches:** 2(2) **Sequences:** 1(1)
mCG144006 [Mus musculus]
gi|154090941 **Mass:** 50102 **Score:** 83 **Matches:** 2(2) **Sequences:** 1(1)
keratin, type I cytoskeletal 42 [Mus musculus]
gi|156139032 **Mass:** 50162 **Score:** 83 **Matches:** 2(2) **Sequences:** 1(1)
Keratin 42 [Mus musculus]
gi|226823220 **Mass:** 49463 **Score:** 83 **Matches:** 2(2) **Sequences:** 1(1)
keratin, type I cytoskeletal 15 [Mus musculus]

---

**9.** gi|51452 **Mass:** 58833 **Score:** 66 **Matches:** 2(2) **Sequences:** 2(2) **emPAI:** 0.11
unnamed protein product [Mus musculus]

| Query | Observed | Mr(expt) | Mr(calc) | Delta | Miss | Score | Expect | Rank | Unique | Peptide |
|---|---|---|---|---|---|---|---|---|---|---|
| 17855 | 695.5581 | 1389.1017 | 1388.6976 | 0.4041 | 0 | 52 | 0.012 | 1 | U | R.GYISPYFINTSK.G |
| 18659 | 752.6903 | 1503.3661 | 1503.7490 | -0.3830 | 0 | 53 | 0.011 | 1 | U | K.TLNDELEIIEGMK.F |

**Proteins matching the same set of peptides:**
gi|51455 **Mass:** 60903 **Score:** 66 **Matches:** 2(2) **Sequences:** 2(2)
heat shock protein 65 [Mus musculus]
gi|26353954 **Mass:** 60918 **Score:** 66 **Matches:** 2(2) **Sequences:** 2(2)
unnamed protein product [Mus musculus]
gi|76779273 **Mass:** 59388 **Score:** 66 **Matches:** 2(2) **Sequences:** 2(2)
Hspd1 protein, partial [Mus musculus]
gi|183396771 **Mass:** 60917 **Score:** 66 **Matches:** 2(2) **Sequences:** 2(2)
60 kDa heat shock protein, mitochondrial [Mus musculus]

---

**10.** gi|74213832 **Mass:** 52871 **Score:** 65 **Matches:** 1(1) **Sequences:** 1(1) **emPAI:** 0.06
unnamed protein product [Mus musculus]

| Query | Observed | Mr(expt) | Mr(calc) | Delta | Miss | Score | Expect | Rank | Unique | Peptide |
|---|---|---|---|---|---|---|---|---|---|---|
| 17796 | 692.4768 | 1382.9391 | 1382.5772 | 0.3618 | 0 | 65 | 0.00055 | 1 | U | R.GGAEQFMEETER.S |

**Proteins matching the same set of peptides:**
gi|148666718 **Mass:** 43932 **Score:** 65 **Matches:** 1(1) **Sequences:** 1(1)
chaperonin subunit 7 (eta), isoform CRA_c [Mus musculus]
gi|26346713 **Mass:** 59598 **Score:** 65 **Matches:** 1(1) **Sequences:** 1(1)

unnamed protein product [Mus musculus]
gi|74147234 **Mass:** 59610 **Score:** 65 **Matches:** 1(1) **Sequences:** 1(1)
unnamed protein product [Mus musculus]
gi|74151643 **Mass:** 59646 **Score:** 65 **Matches:** 1(1) **Sequences:** 1(1)
unnamed protein product [Mus musculus]
gi|74191272 **Mass:** 59600 **Score:** 65 **Matches:** 1(1) **Sequences:** 1(1)
unnamed protein product [Mus musculus]
gi|74212111 **Mass:** 59614 **Score:** 65 **Matches:** 1(1) **Sequences:** 1(1)
unnamed protein product [Mus musculus]
gi|148666717 **Mass:** 61343 **Score:** 65 **Matches:** 1(1) **Sequences:** 1(1)
chaperonin subunit 7 (eta), isoform CRA_b [Mus musculus]
gi|238814391 **Mass:** 59614 **Score:** 65 **Matches:** 1(1) **Sequences:** 1(1)
T-complex protein 1 subunit eta [Mus musculus]

---

**11.** gi|22164776 **Mass:** 57517 **Score:** 65 **Matches:** 2(1) **Sequences:** 2(1) **emPAI:** 0.06
keratin, type II cytoskeletal 79 [Mus musculus]

| Query | Observed | Mr(expt) | Mr(calc) | Delta | Miss | Score | Expect | Rank | Unique | Peptide |
|---|---|---|---|---|---|---|---|---|---|---|
| 16899 | 642.4886 | 1282.9626 | 1283.5491 | -0.5864 | 0 | 43 | 0.084 | 2 | U | K.GGFSSNSASGGGGSR.M |
| 17315 | **665.6135** | **1329.2125** | **1328.7187** | **0.4938** | **0** | **65** | **0.00064** | **1** | **U** | **R.NLDLDSIIAEVK.A** |

**Proteins matching the same set of peptides:**
gi|148672069 **Mass:** 59231 **Score:** 65 **Matches:** 2(1) **Sequences:** 2(1)
cDNA sequence BC031593 [Mus musculus]

---

**12.** gi|12843914 **Mass:** 29993 **Score:** 63 **Matches:** 2(2) **Sequences:** 1(1) **emPAI:** 0.11
unnamed protein product [Mus musculus]

| Query | Observed | Mr(expt) | Mr(calc) | Delta | Miss | Score | Expect | Rank | Unique | Peptide |
|---|---|---|---|---|---|---|---|---|---|---|
| 17066 | **652.3516** | **1302.6886** | **1301.7078** | **0.9807** | **0** | **56** | **0.0062** | **1** | **U** | **R.SLDLDSIIAEVK.A** 17059 |

**Proteins matching the same set of peptides:**
gi|16303309 **Mass:** 61743 **Score:** 63 **Matches:** 2(2) **Sequences:** 1(1)
type II keratin 5 [Mus musculus]
gi|20911031 **Mass:** 61729 **Score:** 63 **Matches:** 2(2) **Sequences:** 1(1)
keratin, type II cytoskeletal 5 [Mus musculus]
gi|54607171 **Mass:** 59299 **Score:** 63 **Matches:** 2(2) **Sequences:** 1(1)
keratin, type II cytoskeletal 6A [Mus musculus]
gi|59798479 **Mass:** 60285 **Score:** 63 **Matches:** 2(2) **Sequences:** 1(1)
RecName: Full=Keratin, type II cytoskeletal 6B; AltName: Full=Cytokeratin-6B; Short=CK-6B; AltName: Full=Keratin-6-beta; Short=mK6-beta; AltName: Full=Keratin-6B; Short=K6B
gi|110645788 **Mass:** 60236 **Score:** 63 **Matches:** 2(2) **Sequences:** 1(1)
Krt6b protein [Mus musculus]
gi|113195684 **Mass:** 59490 **Score:** 63 **Matches:** 2(2) **Sequences:** 1(1)
keratin, type II cytoskeletal 6B [Mus musculus]
gi|116063325 **Mass:** 60154 **Score:** 63 **Matches:** 2(2) **Sequences:** 1(1)
Krt6b protein [Mus musculus]
gi|148672084 **Mass:** 34775 **Score:** 63 **Matches:** 2(2) **Sequences:** 1(1)
mCG17577 [Mus musculus]
gi|148672085 **Mass:** 38119 **Score:** 63 **Matches:** 2(2) **Sequences:** 1(1)
mCG144996 [Mus musculus]
gi|13272554 **Mass:** 42357 **Score:** 63 **Matches:** 2(2) **Sequences:** 1(1)
cytokeratin KRT2-6HF [Mus musculus]
gi|29789317 **Mass:** 59704 **Score:** 63 **Matches:** 2(2) **Sequences:** 1(1)

keratin, type II cytoskeletal 75 [Mus musculus]
gi|148672089 **Mass:** 62241 **Score:** 63 **Matches:** 2(2) **Sequences:** 1(1)
keratin 75 [Mus musculus]

---

**13.** gi|5295992 **Mass:** 59531 **Score:** 63 **Matches:** 2(2) **Sequences:** 2(2) **emPAI:** 0.11
chaperonin containing TCP-1 theta subunit [Mus musculus]

| Query | Observed | Mr(expt) | Mr(calc) | Delta | Miss | Score | Expect | Rank | Unique | Peptide |
|---|---|---|---|---|---|---|---|---|---|---|
| 13522 | 481.9923 | 961.9701 | 961.5055 | 0.4646 | 0 | 52 | 0.0053 | 1 | U | K.APGFAQMLK.D |
| 15269 | 567.3992 | 1132.7838 | 1131.5772 | 1.2066 | 0 | 53 | 0.012 | 1 | U | R.DVDEVSSLLR.T |

**Proteins matching the same set of peptides:**
gi|50510319 **Mass:** 60190 **Score:** 63 **Matches:** 2(2) **Sequences:** 2(2)
mKIAA0002 protein [Mus musculus]
gi|74219231 **Mass:** 59518 **Score:** 63 **Matches:** 2(2) **Sequences:** 2(2)
unnamed protein product [Mus musculus]
gi|126723461 **Mass:** 59518 **Score:** 63 **Matches:** 2(2) **Sequences:** 2(2)
T-complex protein 1 subunit theta [Mus musculus]

---

**14.** gi|1419012 **Mass:** 58147 **Score:** 63 **Matches:** 1(1) **Sequences:** 1(1) **emPAI:** 0.06
CCT zeta2, zeta2 subunit of the chaperonin containing TCP-1 (CCT) [Mus musculus]

| Query | Observed | Mr(expt) | Mr(calc) | Delta | Miss | Score | Expect | Rank | Unique | Peptide |
|---|---|---|---|---|---|---|---|---|---|---|
| 13828 | 496.0008 | 989.9870 | 989.5216 | 0.4655 | 0 | 63 | 0.00052 | 1 | U | K.MLVSGAGDIK.L |

**Proteins matching the same set of peptides:**
gi|6753324 **Mass:** 57968 **Score:** 63 **Matches:** 1(1) **Sequences:** 1(1)
T-complex protein 1 subunit zeta [Mus musculus]
gi|62948125 **Mass:** 58040 **Score:** 63 **Matches:** 1(1) **Sequences:** 1(1)
Chaperonin containing Tcp1, subunit 6a (zeta) [Mus musculus]
gi|74198471 **Mass:** 58067 **Score:** 63 **Matches:** 1(1) **Sequences:** 1(1)
unnamed protein product [Mus musculus]
gi|74204475 **Mass:** 57967 **Score:** 63 **Matches:** 1(1) **Sequences:** 1(1)
unnamed protein product [Mus musculus]
gi|74204595 **Mass:** 58068 **Score:** 63 **Matches:** 1(1) **Sequences:** 1(1)
unnamed protein product [Mus musculus]
gi|226693361 **Mass:** 58148 **Score:** 63 **Matches:** 1(1) **Sequences:** 1(1)
T-complex protein 1 subunit zeta-2 [Mus musculus]

---

**15.** gi|7710044 **Mass:** 69508 **Score:** 60 **Matches:** 1(1) **Sequences:** 1(1) **emPAI:** 0.05
kelch-like ECH-associated protein 1 [Mus musculus]

| Query | Observed | Mr(expt) | Mr(calc) | Delta | Miss | Score | Expect | Rank | Unique | Peptide |
|---|---|---|---|---|---|---|---|---|---|---|
| 17782 | 691.6884 | 1381.3623 | 1381.6990 | -0.3367 | 0 | 60 | 0.0021 | 1 | U | R.LLYAVGGFDGTNR.L |

**Proteins matching the same set of peptides:**
gi|26337871 **Mass:** 69478 **Score:** 60 **Matches:** 1(1) **Sequences:** 1(1)
unnamed protein product [Mus musculus]
gi|37359786 **Mass:** 71015 **Score:** 60 **Matches:** 1(1) **Sequences:** 1(1)
mKIAA0132 protein [Mus musculus]
gi|74181739 **Mass:** 69492 **Score:** 60 **Matches:** 1(1) **Sequences:** 1(1)
unnamed protein product [Mus musculus]
gi|74200263 **Mass:** 43846 **Score:** 60 **Matches:** 1(1) **Sequences:** 1(1)

unnamed protein product [Mus musculus]
gi|74212473 **Mass:** 69482 **Score:** 60 **Matches:** 1(1) **Sequences:** 1(1)
unnamed protein product [Mus musculus]
gi|93278448 **Mass:** 34652 **Score:** 60 **Matches:** 1(1) **Sequences:** 1(1)
Chain A, Structural Basis For The Defects Of Human Lung Cancer Somatic Mutations In The Repression Activity Of Keap1 On Nrf2
gi|158428176 **Mass:** 34946 **Score:** 60 **Matches:** 1(1) **Sequences:** 1(1)
Chain A, Crystal Structure Of The Keap1 Protein In Complexed With The N-Terminal Region Of The Nrf2 Transcription Factor

---

**16.** gi|2623222 **Mass:** 56344 **Score:** 60 **Matches:** 1(1) **Sequences:** 1(1) **emPAI:** 0.06
ATP synthase beta-subunit [Mus musculus]

| Query | Observed | Mr(expt) | Mr(calc) | Delta | Miss | Score | Expect | Rank | Unique | Peptide |
|---|---|---|---|---|---|---|---|---|---|---|
| 18179 | 718.5294 | 1435.0442 | 1434.7467 | 0.2975 | 0 | 60 | 0.00091 | 1 | U | R.FTQAGSEVSALLGR.I |

**Proteins matching the same set of peptides:**
gi|23272966 **Mass:** 56632 **Score:** 60 **Matches:** 1(1) **Sequences:** 1(1)
Atp5b protein [Mus musculus]
gi|31980648 **Mass:** 56265 **Score:** 60 **Matches:** 1(1) **Sequences:** 1(1)
ATP synthase subunit beta, mitochondrial precursor [Mus musculus]
gi|74139650 **Mass:** 56207 **Score:** 60 **Matches:** 1(1) **Sequences:** 1(1)
unnamed protein product [Mus musculus]
gi|74151565 **Mass:** 56294 **Score:** 60 **Matches:** 1(1) **Sequences:** 1(1)
unnamed protein product [Mus musculus]
gi|74185232 **Mass:** 56278 **Score:** 60 **Matches:** 1(1) **Sequences:** 1(1)
unnamed protein product [Mus musculus]
gi|74197074 **Mass:** 56294 **Score:** 60 **Matches:** 1(1) **Sequences:** 1(1)
unnamed protein product [Mus musculus]
gi|74198645 **Mass:** 56294 **Score:** 60 **Matches:** 1(1) **Sequences:** 1(1)
unnamed protein product [Mus musculus]
gi|74204509 **Mass:** 56294 **Score:** 60 **Matches:** 1(1) **Sequences:** 1(1)
unnamed protein product [Mus musculus]
gi|74220566 **Mass:** 56221 **Score:** 60 **Matches:** 1(1) **Sequences:** 1(1)
unnamed protein product [Mus musculus]
gi|74225421 **Mass:** 56235 **Score:** 60 **Matches:** 1(1) **Sequences:** 1(1)
unnamed protein product [Mus musculus]
gi|89574015 **Mass:** 48047 **Score:** 60 **Matches:** 1(1) **Sequences:** 1(1)
mitochondrial ATP synthase, H+ transporting F1 complex beta subunit [Mus musculus]

---

**17.** gi|192262 **Mass:** 54219 **Score:** 58 **Matches:** 1(1) **Sequences:** 1(1) **emPAI:** 0.06
pro-alpha-1 type I collagen, partial [Mus musculus]

| Query | Observed | Mr(expt) | Mr(calc) | Delta | Miss | Score | Expect | Rank | Unique | Peptide |
|---|---|---|---|---|---|---|---|---|---|---|
| 15700 | 589.5136 | 1177.0126 | 1176.5598 | 0.4528 | 0 | 58 | 0.0014 | 1 | U | R.GQAGVMGFPGPK.G |

**Proteins matching the same set of peptides:**
gi|470674 **Mass:** 137859 **Score:** 58 **Matches:** 1(1) **Sequences:** 1(1)
collagen pro-alpha-1 type I chain [Mus musculus]
gi|34328108 **Mass:** 137948 **Score:** 58 **Matches:** 1(1) **Sequences:** 1(1)
collagen alpha-1(I) chain precursor [Mus musculus]
gi|37589303 **Mass:** 117788 **Score:** 58 **Matches:** 1(1) **Sequences:** 1(1)
Col1a1 protein [Mus musculus]

---

**18.** gi|551295 **Mass:** 57824 **Score:** 57 **Matches:** 1(1) **Sequences:** 1(1) **emPAI:** 0.06

pyruvate kinase M [Mus musculus]

| Query | Observed | Mr(expt) | Mr(calc) | Delta | Miss | Score | Expect | Rank | Unique | Peptide |
|---|---|---|---|---|---|---|---|---|---|---|
| 18372 | 732.1815 | 1462.3485 | 1461.8079 | 0.5406 | 0 | 57 | 0.0032 | 1 | U | K.IYVDDGLISLQVK.E |

**Proteins matching the same set of peptides:**

gi|1405933 **Mass:** 57878 **Score:** 57 **Matches:** 1(1) **Sequences:** 1(1)
M2-type pyruvate kinase [Mus musculus]
gi|31981562 **Mass:** 57808 **Score:** 57 **Matches:** 1(1) **Sequences:** 1(1)
pyruvate kinase isozymes M1/M2 isoform 1 [Mus musculus]
gi|74151988 **Mass:** 57820 **Score:** 57 **Matches:** 1(1) **Sequences:** 1(1)
unnamed protein product [Mus musculus]
gi|74196318 **Mass:** 57807 **Score:** 57 **Matches:** 1(1) **Sequences:** 1(1)
unnamed protein product [Mus musculus]
gi|74212815 **Mass:** 43138 **Score:** 57 **Matches:** 1(1) **Sequences:** 1(1)
unnamed protein product [Mus musculus]
gi|74221210 **Mass:** 57809 **Score:** 57 **Matches:** 1(1) **Sequences:** 1(1)
unnamed protein product [Mus musculus]
gi|359807367 **Mass:** 57948 **Score:** 57 **Matches:** 1(1) **Sequences:** 1(1)
pyruvate kinase isozymes M1/M2 isoform 2 [Mus musculus]

---

**19.** gi|52789 **Mass:** 54415 **Score:** 57 **Matches:** 1(1) **Sequences:** 1(1) **emPAI:** 0.06
unnamed protein product [Mus musculus]

| Query | Observed | Mr(expt) | Mr(calc) | Delta | Miss | Score | Expect | Rank | Unique | Peptide |
|---|---|---|---|---|---|---|---|---|---|---|
| 17798 | 692.5499 | 1383.0852 | 1382.7194 | 0.3659 | 1 | 57 | 0.0016 | 1 | U | K.SLNNKFASFIDK.V |

**Proteins matching the same set of peptides:**

gi|309215 **Mass:** 53210 **Score:** 57 **Matches:** 1(1) **Sequences:** 1(1)
EndoA' cytokeratin (5' end put.); putative [Mus musculus]
gi|511654 **Mass:** 54220 **Score:** 57 **Matches:** 1(1) **Sequences:** 1(1)
keratin type II [Mus musculus]
gi|74177777 **Mass:** 54546 **Score:** 57 **Matches:** 1(1) **Sequences:** 1(1)
unnamed protein product [Mus musculus]
gi|74219975 **Mass:** 54459 **Score:** 57 **Matches:** 1(1) **Sequences:** 1(1)
unnamed protein product [Mus musculus]
gi|76779293 **Mass:** 54514 **Score:** 57 **Matches:** 1(1) **Sequences:** 1(1)
Keratin 8 [Mus musculus]
gi|114145561 **Mass:** 54531 **Score:** 57 **Matches:** 1(1) **Sequences:** 1(1)
keratin, type II cytoskeletal 8 [Mus musculus]

---

**20.** gi|46485130 **Mass:** 85186 **Score:** 57 **Matches:** 1(1) **Sequences:** 1(1) **emPAI:** 0.04
TPA_exp: keratin Kb40 [Mus musculus]

| Query | Observed | Mr(expt) | Mr(calc) | Delta | Miss | Score | Expect | Rank | Unique | Peptide |
|---|---|---|---|---|---|---|---|---|---|---|
| 17798 | 692.5499 | 1383.0852 | 1382.6830 | 0.4022 | 0 | 57 | 0.0017 | 2 | U | R.SLNNQFASFIDK.V |

**Proteins matching the same set of peptides:**

gi|111185567 **Score:** 57 **Matches:** 1(1) **Sequences:** 1(1)
gi|111185722 **Score:** 57 **Matches:** 1(1) **Sequences:** 1(1)
gi|119850791 **Score:** 57 **Matches:** 1(1) **Sequences:** 1(1)
gi|145580629 **Mass:** 112194 **Score:** 57 **Matches:** 1(1) **Sequences:** 1(1)
keratin Kb40 [Mus musculus]

---

21. gi|1304157 **Mass:** 72412 **Score:** 57 **Matches:** 1(1) **Sequences:** 1(1) **emPAI:** 0.05
78 kDa glucose-regulated protein [Mus musculus]

| Query | Observed | Mr(expt) | Mr(calc) | Delta | Miss | Score | Expect | Rank | Unique | Peptide |
|---|---|---|---|---|---|---|---|---|---|---|
| 18139 | 715.6650 | 1429.3155 | 1429.6838 | -0.3682 | 0 | 57 | 0.0043 | 1 | U | R.TWNDPSVQQDIK.F |

**Proteins matching the same set of peptides:**
gi|2598562 **Mass:** 72433 **Score:** 57 **Matches:** 1(1) **Sequences:** 1(1)
BiP [Mus musculus]
gi|12835845 **Mass:** 72378 **Score:** 57 **Matches:** 1(1) **Sequences:** 1(1)
unnamed protein product [Mus musculus]
gi|74143673 **Mass:** 56858 **Score:** 57 **Matches:** 1(1) **Sequences:** 1(1)
unnamed protein product [Mus musculus]
gi|74188814 **Mass:** 72305 **Score:** 57 **Matches:** 1(1) **Sequences:** 1(1)
unnamed protein product [Mus musculus]
gi|74198293 **Mass:** 72419 **Score:** 57 **Matches:** 1(1) **Sequences:** 1(1)
unnamed protein product [Mus musculus]
gi|74198974 **Mass:** 72361 **Score:** 57 **Matches:** 1(1) **Sequences:** 1(1)
unnamed protein product [Mus musculus]
gi|74207492 **Mass:** 72301 **Score:** 57 **Matches:** 1(1) **Sequences:** 1(1)
unnamed protein product [Mus musculus]
gi|74220199 **Mass:** 68455 **Score:** 57 **Matches:** 1(1) **Sequences:** 1(1)
unnamed protein product [Mus musculus]
gi|74225394 **Mass:** 72337 **Score:** 57 **Matches:** 1(1) **Sequences:** 1(1)
unnamed protein product [Mus musculus]
gi|254540166 **Mass:** 72377 **Score:** 57 **Matches:** 1(1) **Sequences:** 1(1)
78 kDa glucose-regulated protein precursor [Mus musculus]

---

22. gi|4159806 **Mass:** 65183 **Score:** 55 **Matches:** 2(2) **Sequences:** 2(2) **emPAI:** 0.10
type II keratin subunit protein [Mus musculus]

| Query | Observed | Mr(expt) | Mr(calc) | Delta | Miss | Score | Expect | Rank | Unique | Peptide |
|---|---|---|---|---|---|---|---|---|---|---|
| 17798 | 692.5499 | 1383.0852 | 1383.7034 | -0.6182 | 1 | 44 | 0.031 | 3 | U | K.SLNDKFASFIDK.V |
| 17884 | 697.6450 | 1393.2755 | 1392.7249 | 0.5506 | 1 | 55 | 0.0067 | 1 | U | R.TNAENEFVTIKK.D |

**Proteins matching the same set of peptides:**
gi|12859782 **Mass:** 65586 **Score:** 55 **Matches:** 2(2) **Sequences:** 2(2)
unnamed protein product [Mus musculus]
gi|126116585 **Mass:** 65565 **Score:** 55 **Matches:** 2(2) **Sequences:** 2(2)
keratin, type II cytoskeletal 1 [Mus musculus]

---

23. gi|293686 **Mass:** 59414 **Score:** 55 **Matches:** 1(1) **Sequences:** 1(1) **emPAI:** 0.06
epidermal keratin subunit II [Mus musculus]

| Query | Observed | Mr(expt) | Mr(calc) | Delta | Miss | Score | Expect | Rank | Unique | Peptide |
|---|---|---|---|---|---|---|---|---|---|---|
| 17884 | 697.6450 | 1393.2755 | 1393.7089 | -0.4334 | 1 | 55 | 0.0067 | 1 | U | R.TDAENEFVTLKK.D |

---

24. gi|3687320 **Mass:** 46137 **Score:** 52 **Matches:** 1(1) **Sequences:** 1(1) **emPAI:** 0.07
cartilage-associated protein (CASP) [Mus musculus]

| Query | Observed | Mr(expt) | Mr(calc) | Delta | Miss | Score | Expect | Rank | Unique | Peptide |
|---|---|---|---|---|---|---|---|---|---|---|
| 19209 | 840.6470 | 1679.2794 | 1678.8488 | 0.4306 | 0 | 52 | 0.01 | 1 | U | R.TSISDMELALPDFLK.A |

**Proteins matching the same set of peptides:**

gi|12857227 **Mass:** 46121 **Score:** 52 **Matches:** 1(1) **Sequences:** 1(1)
unnamed protein product [Mus musculus]
gi|148677365 **Mass:** 46110 **Score:** 52 **Matches:** 1(1) **Sequences:** 1(1)
mCG7119 [Mus musculus]
gi|225543173 **Mass:** 46140 **Score:** 52 **Matches:** 1(1) **Sequences:** 1(1)
cartilage-associated protein precursor [Mus musculus]

---

**25.** gi|74203983 **Mass:** 60183 **Score:** 51 **Matches:** 1(1) **Sequences:** 1(1) **emPAI:** 0.05
unnamed protein product [Mus musculus]

| Query | Observed | Mr(expt) | Mr(calc) | Delta | Miss | Score | Expect | Rank | Unique | Peptide |
|---|---|---|---|---|---|---|---|---|---|---|
| 17926 | 700.4083 | 1398.8021 | 1398.6159 | 0.1862 | 0 | 51 | 0.02 | 1 | U | K.SIDMSWDSVTMK.H |

**Proteins matching the same set of peptides:**
gi|76677895 **Mass:** 58508 **Score:** 51 **Matches:** 1(1) **Sequences:** 1(1)
poly(U)-binding-splicing factor PUF60 isoform b [Mus musculus]
gi|111185612 **Mass:** 26264 **Score:** 51 **Matches:** 1(1) **Sequences:** 1(1)
Puf60 protein [Mus musculus]
gi|148697575 **Mass:** 58380 **Score:** 51 **Matches:** 1(1) **Sequences:** 1(1)
RIKEN cDNA 2410104I19, isoform CRA_c [Mus musculus]
gi|148697576 **Mass:** 58982 **Score:** 51 **Matches:** 1(1) **Sequences:** 1(1)
RIKEN cDNA 2410104I19, isoform CRA_d [Mus musculus]
gi|257196183 **Mass:** 60211 **Score:** 51 **Matches:** 1(1) **Sequences:** 1(1)
poly(U)-binding-splicing factor PUF60 isoform a [Mus musculus]
gi|257196186 **Mass:** 53995 **Score:** 51 **Matches:** 1(1) **Sequences:** 1(1)
poly(U)-binding-splicing factor PUF60 isoform c [Mus musculus]

---

**26.** gi|468546 **Mass:** 57411 **Score:** 50 **Matches:** 1(1) **Sequences:** 1(1) **emPAI:** 0.06
CCT (chaperonin containing TCP-1) beta subunit [Mus musculus]

| Query | Observed | Mr(expt) | Mr(calc) | Delta | Miss | Score | Expect | Rank | Unique | Peptide |
|---|---|---|---|---|---|---|---|---|---|---|
| 18778 | 767.4490 | 1532.8834 | 1531.7916 | 1.0918 | 0 | 50 | 0.023 | 1 | U | R.DAALMVTNDGATILK.N |

**Proteins matching the same set of peptides:**
gi|7670405 **Mass:** 52436 **Score:** 50 **Matches:** 1(1) **Sequences:** 1(1)
unnamed protein product [Mus musculus]
gi|126521835 **Mass:** 57441 **Score:** 50 **Matches:** 1(1) **Sequences:** 1(1)
T-complex protein 1 subunit beta [Mus musculus]
gi|148689878 **Mass:** 44497 **Score:** 50 **Matches:** 1(1) **Sequences:** 1(1)
chaperonin subunit 2 (beta), isoform CRA_b [Mus musculus]

---

**27.** gi|22297268 **Score:** 50 **Matches:** 1(1) **Sequences:** 1(1) **emPAI:** 0.16
AID, partial [Homo sapiens]

| Query | Observed | Mr(expt) | Mr(calc) | Delta | Miss | Score | Expect | Rank | Unique | Peptide |
|---|---|---|---|---|---|---|---|---|---|---|
| 10277 | 381.1882 | 760.3618 | 760.3715 | -0.0097 | 0 | 50 | 0.015 | 2 | U | K.AESEGLR.R |

---

**28.** gi|51450 **Mass:** 46560 **Score:** 49 **Matches:** 1(1) **Sequences:** 1(1) **emPAI:** 0.07
heat shock protein [Mus musculus]

| Query | Observed | Mr(expt) | Mr(calc) | Delta | Miss | Score | Expect | Rank | Unique | Peptide |
|---|---|---|---|---|---|---|---|---|---|---|
| 16265 | 612.9139 | 1223.8133 | 1223.6510 | 0.1623 | 0 | 49 | 0.027 | 1 | U | K.GVVEVTHDLQK.H |

**Proteins matching the same set of peptides:**

gi|200966 **Mass:** 45623 **Score:** 49 **Matches:** 1(1) **Sequences:** 1(1)
put. serine protease inhibitor [Mus musculus]
gi|26345418 **Mass:** 46481 **Score:** 49 **Matches:** 1(1) **Sequences:** 1(1)
unnamed protein product [Mus musculus]
gi|26348007 **Mass:** 46490 **Score:** 49 **Matches:** 1(1) **Sequences:** 1(1)
unnamed protein product [Mus musculus]
gi|74139266 **Mass:** 46490 **Score:** 49 **Matches:** 1(1) **Sequences:** 1(1)
unnamed protein product [Mus musculus]
gi|74191337 **Mass:** 46505 **Score:** 49 **Matches:** 1(1) **Sequences:** 1(1)
unnamed protein product [Mus musculus]
gi|74198254 **Mass:** 46500 **Score:** 49 **Matches:** 1(1) **Sequences:** 1(1)
unnamed protein product [Mus musculus]
gi|148684430 **Mass:** 44954 **Score:** 49 **Matches:** 1(1) **Sequences:** 1(1)
serine (or cysteine) peptidase inhibitor, clade H, member 1, isoform CRA_a [Mus musculus]
gi|161353502 **Mass:** 46504 **Score:** 49 **Matches:** 1(1) **Sequences:** 1(1)
serpin H1 precursor [Mus musculus]

---

**29.** gi|6680748 **Mass:** 59716 **Score:** 49 **Matches:** 1(1) **Sequences:** 1(1) **emPAI:** 0.06
ATP synthase subunit alpha, mitochondrial precursor [Mus musculus]

| Query | Observed | Mr(expt) | Mr(calc) | Delta | Miss | Score | Expect | Rank | Unique | Peptide |
|---|---|---|---|---|---|---|---|---|---|---|
| 9365 | 362.2126 | 722.4106 | 722.4439 | -0.0333 | 0 | 49 | 0.012 | 1 | U | K.APGIIPR.I |

**Proteins matching the same set of peptides:**
gi|74139457 **Mass:** 59656 **Score:** 49 **Matches:** 1(1) **Sequences:** 1(1)
unnamed protein product [Mus musculus]
gi|74146998 **Mass:** 55919 **Score:** 49 **Matches:** 1(1) **Sequences:** 1(1)
unnamed protein product [Mus musculus]
gi|74182195 **Mass:** 32607 **Score:** 49 **Matches:** 1(1) **Sequences:** 1(1)
unnamed protein product [Mus musculus]
gi|74211072 **Mass:** 59730 **Score:** 49 **Matches:** 1(1) **Sequences:** 1(1)
unnamed protein product [Mus musculus]
gi|74211198 **Mass:** 51304 **Score:** 49 **Matches:** 1(1) **Sequences:** 1(1)
unnamed protein product [Mus musculus]
gi|74211977 **Mass:** 59746 **Score:** 49 **Matches:** 1(1) **Sequences:** 1(1)
unnamed protein product [Mus musculus]
gi|148677500 **Mass:** 51819 **Score:** 49 **Matches:** 1(1) **Sequences:** 1(1)
ATP synthase, H+ transporting, mitochondrial F1 complex, alpha subunit, isoform 1, isoform CRA_d [Mus musculus]
gi|148677501 **Mass:** 54561 **Score:** 49 **Matches:** 1(1) **Sequences:** 1(1)
ATP synthase, H+ transporting, mitochondrial F1 complex, alpha subunit, isoform 1, isoform CRA_e [Mus musculus]
gi|148677504 **Mass:** 54738 **Score:** 49 **Matches:** 1(1) **Sequences:** 1(1)
ATP synthase, H+ transporting, mitochondrial F1 complex, alpha subunit, isoform 1, isoform CRA_h [Mus musculus]

---

**30.** gi|148688671 **Score:** 49 **Matches:** 1(1) **Sequences:** 1(1) **emPAI:** 0.30
mCG16097 [Mus musculus]

| Query | Observed | Mr(expt) | Mr(calc) | Delta | Miss | Score | Expect | Rank | Unique | Peptide |
|---|---|---|---|---|---|---|---|---|---|---|
| 9365 | 362.2126 | 722.4106 | 722.4439 | -0.0333 | 0 | 49 | 0.012 | 1 | U | K.APGLLPR.L |

---

**31.** gi|6755893 **Mass:** 26257 **Score:** 48 **Matches:** 2(2) **Sequences:** 1(1) **emPAI:** 0.27
trypsin 4 precursor [Mus musculus]

| Query | Observed | Mr(expt) | Mr(calc) | Delta | Miss | Score | Expect | Rank | Unique | Peptide |
|---|---|---|---|---|---|---|---|---|---|---|
| 18452 | 737.6481 | 2209.9224 | 2211.0920 | -1.1696 | 0 | 47 | 0.039 | 2 | U | R.LGEHNINVLEGNEQFVNSAK.I |

| | | | | | | | | | | |
|---|---|---|---|---|---|---|---|---|---|---|
| 19321 | 1106.1886 | 2210.3626 | 2211.0920 | -0.7293 | 0 | (47) | 0.044 | 2 | U | R.LGEHNINVLEGNEQFVNSAK.I |

**Proteins matching the same set of peptides:**
gi|51010909 **Mass:** 26260 **Score:** 48 **Matches:** 2(2) **Sequences:** 1(1)
trypsin 5 precursor [Mus musculus]
gi|74203392 **Mass:** 27146 **Score:** 48 **Matches:** 2(2) **Sequences:** 1(1)
unnamed protein product [Mus musculus]

---

**32.** gi|148707748 **Score:** 47 **Matches:** 1(1) **Sequences:** 1(1) **emPAI:** 0.10
nuclear casein kinase and cyclin-dependent kinase substrate 1, isoform CRA_b [Mus musculus]

| Query | Observed | Mr(expt) | Mr(calc) | Delta | Miss | Score | Expect | Rank | Unique | Peptide |
|---|---|---|---|---|---|---|---|---|---|---|
| 18778 | 767.4490 | 1532.8834 | 1531.6525 | 1.2309 | 1 | 47 | 0.044 | 2 | U | K.EEDEEAESPPEKK.S |

**Proteins matching the same set of peptides:**
gi|224809559 **Score:** 47 **Matches:** 1(1) **Sequences:** 1(1)
gi|224809563 **Score:** 47 **Matches:** 1(1) **Sequences:** 1(1)

---

**33.** gi|53830774 **Mass:** 34092 **Score:** 47 **Matches:** 1(1) **Sequences:** 1(1) **emPAI:** 0.10
lactase [Mus musculus]

| Query | Observed | Mr(expt) | Mr(calc) | Delta | Miss | Score | Expect | Rank | Unique | Peptide |
|---|---|---|---|---|---|---|---|---|---|---|
| 14788 | **548.6042** | **1095.1938** | **1094.5132** | **0.6806** | **0** | **47** | **0.016** | **1** | **U** | **R.LPEFTESEK.K** |

**Proteins matching the same set of peptides:**
gi|74192292 **Mass:** 139170 **Score:** 47 **Matches:** 1(1) **Sequences:** 1(1)
unnamed protein product [Mus musculus]
gi|124487297 **Mass:** 217665 **Score:** 47 **Matches:** 1(1) **Sequences:** 1(1)
lactase-phlorizin hydrolase preproprotein [Mus musculus]
gi|148707805 **Mass:** 217693 **Score:** 47 **Matches:** 1(1) **Sequences:** 1(1)
mCG128560 [Mus musculus]

---

**34.** gi|26324736 **Score:** 47 **Matches:** 1(1) **Sequences:** 1(1) **emPAI:** 0.06
unnamed protein product [Mus musculus]

| Query | Observed | Mr(expt) | Mr(calc) | Delta | Miss | Score | Expect | Rank | Unique | Peptide |
|---|---|---|---|---|---|---|---|---|---|---|
| 17059 | 651.8716 | 1301.7286 | 1300.7238 | 1.0048 | 0 | 47 | 0.029 | 2 | U | R.SLNLDSIIAEVK.A |

**Proteins matching the same set of peptides:**
gi|109735018 **Score:** 47 **Matches:** 1(1) **Sequences:** 1(1)
gi|148672090 **Score:** 47 **Matches:** 1(1) **Sequences:** 1(1)
gi|269914154 **Score:** 47 **Matches:** 1(1) **Sequences:** 1(1)

---

**35.** gi|7513694 **Mass:** 109137 **Score:** 46 **Matches:** 1(1) **Sequences:** 1(1) **emPAI:** 0.03
IgG Fc binding protein - mouse (fragment)

| Query | Observed | Mr(expt) | Mr(calc) | Delta | Miss | Score | Expect | Rank | Unique | Peptide |
|---|---|---|---|---|---|---|---|---|---|---|
| 12529 | **437.9564** | **873.8983** | **873.4920** | **0.4063** | **0** | **46** | **0.022** | **1** | **U** | **R.ISVINGGSK.A** |

**Proteins matching the same set of peptides:**
gi|148692210 **Mass:** 147962 **Score:** 46 **Matches:** 1(1) **Sequences:** 1(1)
mCG145390, isoform CRA_a [Mus musculus]
gi|148692211 **Mass:** 146729 **Score:** 46 **Matches:** 1(1) **Sequences:** 1(1)
mCG145390, isoform CRA_b [Mus musculus]

gi|169790797 **Mass:** 275058 **Score:** 46 **Matches:** 1(1) **Sequences:** 1(1)
Fc fragment of IgG binding protein precursor [Mus musculus]
gi|257467625 **Mass:** 280044 **Score:** 46 **Matches:** 1(1) **Sequences:** 1(1)
Fc fragment of IgG binding protein-like precursor [Mus musculus]

---

**36.** gi|111599858 **Score:** 46 **Matches:** 1(1) **Sequences:** 1(1) **emPAI:** 0.03
Dehydrogenase E1 and transketolase domain containing 1 [Mus musculus]

| Query | Observed | Mr(expt) | Mr(calc) | Delta | Miss | Score | Expect | Rank | Unique | Peptide |
|---|---|---|---|---|---|---|---|---|---|---|
| 10277 | 381.1882 | 760.3618 | 759.3875 | 0.9743 | 1 | 46 | 0.034 | 3 | U | K.AKSGEPR.G |

**Proteins matching the same set of peptides:**
gi|124487485 **Score:** 46 **Matches:** 1(1) **Sequences:** 1(1)

---

**37.** gi|74224513 **Mass:** 37387 **Score:** 44 **Matches:** 1(1) **Sequences:** 1(1) **emPAI:** 0.09
unnamed protein product [Mus musculus]

| Query | Observed | Mr(expt) | Mr(calc) | Delta | Miss | Score | Expect | Rank | Unique | Peptide |
|---|---|---|---|---|---|---|---|---|---|---|
| 11939 | **419.3358** | **836.6570** | **835.5280** | **1.1290** | **0** | **44** | **0.035** | **1** | **U** | **R.VAAPPLLR.V** |

---

**38.** gi|21410133 **Score:** 44 **Matches:** 1(1) **Sequences:** 1(1) **emPAI:** 0.13
Kidins220 protein [Mus musculus]

| Query | Observed | Mr(expt) | Mr(calc) | Delta | Miss | Score | Expect | Rank | Unique | Peptide |
|---|---|---|---|---|---|---|---|---|---|---|
| 10277 | 381.1882 | 760.3618 | 759.3875 | 0.9743 | 1 | 44 | 0.055 | 4 | U | K.AEGKAER.V |

**Proteins matching the same set of peptides:**
gi|28385979 **Mass:** 172572 **Score:** 44 **Matches:** 1(1) **Sequences:** 1(1)
Kidins220 protein [Mus musculus]
gi|28972688 **Mass:** 188137 **Score:** 44 **Matches:** 1(1) **Sequences:** 1(1)
mKIAA1250 protein [Mus musculus]
gi|62826025 **Score:** 44 **Matches:** 1(1) **Sequences:** 1(1)
gi|124487039 **Mass:** 199059 **Score:** 44 **Matches:** 1(1) **Sequences:** 1(1)
kinase D-interacting substrate of 220 kDa [Mus musculus]
gi|148705033 **Mass:** 190650 **Score:** 44 **Matches:** 1(1) **Sequences:** 1(1)
mCG18896 [Mus musculus]
gi|187957160 **Mass:** 185440 **Score:** 44 **Matches:** 1(1) **Sequences:** 1(1)
Kidins220 protein [Mus musculus]

---

**39.** gi|26333649 **Score:** 44 **Matches:** 1(1) **Sequences:** 1(1) **emPAI:** 0.07
unnamed protein product [Mus musculus]

| Query | Observed | Mr(expt) | Mr(calc) | Delta | Miss | Score | Expect | Rank | Unique | Peptide |
|---|---|---|---|---|---|---|---|---|---|---|
| 13828 | 496.0008 | 989.9870 | 989.5182 | 0.4689 | 1 | 44 | 0.04 | 2 | U | K.FPKDAGLDK.L |

**Proteins matching the same set of peptides:**
gi|74146722 **Score:** 44 **Matches:** 1(1) **Sequences:** 1(1)
gi|74192493 **Score:** 44 **Matches:** 1(1) **Sequences:** 1(1)
gi|113865995 **Score:** 44 **Matches:** 1(1) **Sequences:** 1(1)
gi|116325981 **Score:** 44 **Matches:** 1(1) **Sequences:** 1(1)
gi|148669182 **Score:** 44 **Matches:** 1(1) **Sequences:** 1(1)
gi|148707033 **Score:** 44 **Matches:** 1(1) **Sequences:** 1(1)
gi|254847433 **Score:** 44 **Matches:** 1(1) **Sequences:** 1(1)

gi|254847435 Score: 44 Matches: 1(1) Sequences: 1(1)
gi|254847438 Score: 44 Matches: 1(1) Sequences: 1(1)
gi|377837285 Score: 44 Matches: 1(1) Sequences: 1(1)
gi|377837287 Score: 44 Matches: 1(1) Sequences: 1(1)

---

**40.** gi|986918 **Mass:** 51416 **Score:** 44 **Matches:** 1(1) **Sequences:** 1(1) **emPAI:** 0.06
A10 [Mus musculus]

| Query | Observed | Mr(expt) | Mr(calc) | Delta | Miss | Score | Expect | Rank | Unique | Peptide |
|---|---|---|---|---|---|---|---|---|---|---|
| 15221 | 566.0025 | 1129.9904 | 1129.5979 | 0.3925 | 0 | 44 | 0.036 | 1 | U | K.VTADVINAAEK.L |

**Proteins matching the same set of peptides:**
gi|26345686 **Mass:** 56546 **Score:** 44 **Matches:** 1(1) **Sequences:** 1(1)
unnamed protein product [Mus musculus]
gi|52353955 **Mass:** 56549 **Score:** 44 **Matches:** 1(1) **Sequences:** 1(1)
D-3-phosphoglycerate dehydrogenase [Mus musculus]

---

**41.** gi|21313084 **Mass:** 56107 **Score:** 44 **Matches:** 1(1) **Sequences:** 1(1) **emPAI:** 0.06
chromodomain Y-like protein 2 [Mus musculus]

| Query | Observed | Mr(expt) | Mr(calc) | Delta | Miss | Score | Expect | Rank | Unique | Peptide |
|---|---|---|---|---|---|---|---|---|---|---|
| 15639 | 585.4729 | 1168.9312 | 1168.6676 | 0.2636 | 1 | 44 | 0.035 | 1 | U | K.RVNSPLSRPK.K |

**Proteins matching the same set of peptides:**
gi|32479356 **Mass:** 56107 **Score:** 44 **Matches:** 1(1) **Sequences:** 1(1)
chromodomain Y-like protein 2 [Mus musculus]
gi|74206467 **Mass:** 56077 **Score:** 44 **Matches:** 1(1) **Sequences:** 1(1)
unnamed protein product [Mus musculus]

---

**42.** gi|124504325 **Mass:** 128405 **Score:** 44 **Matches:** 1(0) **Sequences:** 1(0) **emPAI:** 0.03
Armcx4 protein [Mus musculus]

| Query | Observed | Mr(expt) | Mr(calc) | Delta | Miss | Score | Expect | Rank | Unique | Peptide |
|---|---|---|---|---|---|---|---|---|---|---|
| 12706 | 887.4671 | 886.4599 | 886.4621 | -0.0022 | 1 | 44 | 0.071 | 1 | U | K.GNRNSVPK.A |

**Proteins matching the same set of peptides:**
gi|363989837 **Mass:** 242806 **Score:** 44 **Matches:** 1(0) **Sequences:** 1(0)
armadillo repeat-containing X-linked protein 4 [Mus musculus]

---

**43.** gi|12843046 **Mass:** 26390 **Score:** 43 **Matches:** 1(0) **Sequences:** 1(0) **emPAI:** 0.13
unnamed protein product [Mus musculus]

| Query | Observed | Mr(expt) | Mr(calc) | Delta | Miss | Score | Expect | Rank | Unique | Peptide |
|---|---|---|---|---|---|---|---|---|---|---|
| 18063 | 711.1305 | 1420.2464 | 1419.7146 | 0.5318 | 0 | 43 | 0.081 | 1 | U | K.YVNWIQQTIAAN.- |

**Proteins matching the same set of peptides:**
gi|71043961 **Mass:** 26405 **Score:** 43 **Matches:** 1(0) **Sequences:** 1(0)
trypsinogen 7 precursor [Mus musculus]
gi|148681611 **Mass:** 25182 **Score:** 43 **Matches:** 1(0) **Sequences:** 1(0)
RIKEN cDNA 2210010C04, isoform CRA_a [Mus musculus]

---

**44.** gi|12836661 **Mass:** 117980 **Score:** 43 **Matches:** 1(1) **Sequences:** 1(1) **emPAI:** 0.03
unnamed protein product [Mus musculus]

| Query | Observed | Mr(expt) | Mr(calc) | Delta | Miss | Score | Expect | Rank | Unique | Peptide |
|---|---|---|---|---|---|---|---|---|---|---|

| | | | | | | | | | | |
|---|---|---|---|---|---|---|---|---|---|---|
| 14417 | 530.9701 | 1059.9256 | 1059.5093 | 0.4164 | 0 | 43 | 0.045 | 1 | U | R.CPPPLACALK.A |

**Proteins matching the same set of peptides:**

gi\|16741201 **Mass:** 70327 **Score:** 43 **Matches:** 1(1) **Sequences:** 1(1)
Pelp1 protein [Mus musculus]

gi\|37046775 **Mass:** 91728 **Score:** 43 **Matches:** 1(1) **Sequences:** 1(1)
Pelp1 protein, partial [Mus musculus]

gi\|257900472 **Mass:** 117995 **Score:** 43 **Matches:** 1(1) **Sequences:** 1(1)
proline-, glutamic acid- and leucine-rich protein 1 [Mus musculus]

---

**45.** gi\|148681245 **Mass:** 9487 **Score:** 43 **Matches:** 1(0) **Sequences:** 1(0) **emPAI:** 0.37
mCG145199 [Mus musculus]

| Query | Observed | Mr(expt) | Mr(calc) | Delta | Miss | Score | Expect | Rank | Unique | Peptide |
|---|---|---|---|---|---|---|---|---|---|---|
| 11940 | 419.3534 | 836.6922 | 836.4729 | 0.2193 | 1 | 43 | 0.085 | 1 | U | R.LAGHGRAR.S |

---

**46.** gi\|148667455 **Mass:** 58456 **Score:** 43 **Matches:** 1(1) **Sequences:** 1(1) **emPAI:** 0.06
TEA domain family member 4, isoform CRA_b [Mus musculus]

| Query | Observed | Mr(expt) | Mr(calc) | Delta | Miss | Score | Expect | Rank | Unique | Peptide |
|---|---|---|---|---|---|---|---|---|---|---|
| 14488 | 534.0168 | 1066.0190 | 1066.5995 | -0.5805 | 1 | 43 | 0.041 | 1 | U | R.ARGPAAVAAAGR.L |

**Proteins matching the same set of peptides:**

gi\|148667459 **Mass:** 62771 **Score:** 43 **Matches:** 1(1) **Sequences:** 1(1)
TEA domain family member 4, isoform CRA_f [Mus musculus]

gi\|148667454 **Mass:** 59545 **Score:** 43 **Matches:** 1(1) **Sequences:** 1(1)
TEA domain family member 4, isoform CRA_a [Mus musculus]

---

**47.** gi\|642072 **Mass:** 118421 **Score:** 43 **Matches:** 1(0) **Sequences:** 1(0) **emPAI:** 0.03
fibrillin-1, partial [Mus musculus]

| Query | Observed | Mr(expt) | Mr(calc) | Delta | Miss | Score | Expect | Rank | Unique | Peptide |
|---|---|---|---|---|---|---|---|---|---|---|
| 14550 | 536.8515 | 1071.6884 | 1070.4451 | 1.2433 | 0 | 43 | 0.076 | 1 | U | R.CVNTDGSYR.C |

**Proteins matching the same set of peptides:**

gi\|726324 **Mass:** 312194 **Score:** 43 **Matches:** 1(0) **Sequences:** 1(0)
fibrillin-1 [Mus musculus]

gi\|2494284 **Mass:** 312051 **Score:** 43 **Matches:** 1(0) **Sequences:** 1(0)
RecName: Full=Fibrillin-1; Flags: Precursor

gi\|3688648 **Mass:** 418019 **Score:** 43 **Matches:** 1(0) **Sequences:** 1(0)
mutant fibrillin-1 [Mus musculus]

gi\|118197277 **Mass:** 312083 **Score:** 43 **Matches:** 1(0) **Sequences:** 1(0)
fibrillin-1 precursor [Mus musculus]

gi\|148696192 **Mass:** 312056 **Score:** 43 **Matches:** 1(0) **Sequences:** 1(0)
fibrillin 1, isoform CRA_a [Mus musculus]

---

**48.** gi\|12836791 **Mass:** 23191 **Score:** 43 **Matches:** 1(0) **Sequences:** 1(0) **emPAI:** 0.14
unnamed protein product [Mus musculus]

| Query | Observed | Mr(expt) | Mr(calc) | Delta | Miss | Score | Expect | Rank | Unique | Peptide |
|---|---|---|---|---|---|---|---|---|---|---|
| 16636 | 629.0249 | 1256.0352 | 1255.5478 | 0.4874 | 0 | 43 | 0.11 | 1 | U | -.MPPGPCAWPPR.A |

**Proteins matching the same set of peptides:**

gi|47059145 **Mass:** 23306 **Score:** 43 **Matches:** 1(0) **Sequences:** 1(0)
fibronectin type III domain-containing protein 5 precursor [Mus musculus]

---

**49.** gi|14250408 **Mass:** 57081 **Score:** 42 **Matches:** 1(0) **Sequences:** 1(0) **emPAI:** 0.06
Aspartyl-tRNA synthetase [Mus musculus]

| Query | Observed | Mr(expt) | Mr(calc) | Delta | Miss | Score | Expect | Rank | Unique | Peptide |
|---|---|---|---|---|---|---|---|---|---|---|
| 15565 | 581.3489 | 1160.6833 | 1159.6601 | 1.0232 | 0 | 42 | 0.077 | 1 | U | K.IYVISLAEPR.L |

**Proteins matching the same set of peptides:**
gi|26346400 **Mass:** 57111 **Score:** 42 **Matches:** 1(0) **Sequences:** 1(0)
unnamed protein product [Mus musculus]
gi|74139282 **Mass:** 57145 **Score:** 42 **Matches:** 1(0) **Sequences:** 1(0)
unnamed protein product [Mus musculus]
gi|74141887 **Mass:** 57067 **Score:** 42 **Matches:** 1(0) **Sequences:** 1(0)
unnamed protein product [Mus musculus]
gi|74226918 **Mass:** 57109 **Score:** 42 **Matches:** 1(0) **Sequences:** 1(0)
unnamed protein product [Mus musculus]
gi|148707802 **Mass:** 64317 **Score:** 42 **Matches:** 1(0) **Sequences:** 1(0)
aspartyl-tRNA synthetase [Mus musculus]
gi|211065507 **Mass:** 57111 **Score:** 42 **Matches:** 1(0) **Sequences:** 1(0)
aspartate--tRNA ligase, cytoplasmic isoform 1 [Mus musculus]

---

**50.** gi|17512475 **Mass:** 46233 **Score:** 42 **Matches:** 1(0) **Sequences:** 1(0) **emPAI:** 0.07
Ankyrin repeat and SOCS box-containing 6 [Mus musculus]

| Query | Observed | Mr(expt) | Mr(calc) | Delta | Miss | Score | Expect | Rank | Unique | Peptide |
|---|---|---|---|---|---|---|---|---|---|---|
| 18446 | 736.9879 | 2207.9419 | 2208.1902 | -0.2483 | 1 | 42 | 0.14 | 1 | U | K.IQALHASLRQLESYPPPLK.H |

**Proteins matching the same set of peptides:**
gi|19111154 **Mass:** 46247 **Score:** 42 **Matches:** 1(0) **Sequences:** 1(0)
ankyrin repeat and SOCS box protein 6 [Mus musculus]
gi|26354010 **Mass:** 46217 **Score:** 42 **Matches:** 1(0) **Sequences:** 1(0)
unnamed protein product [Mus musculus]

---

**51.** gi|309271872 **Mass:** 403142 **Score:** 42 **Matches:** 1(1) **Sequences:** 1(1) **emPAI:** 0.01
PREDICTED: zinc finger protein 469 [Mus musculus]

| Query | Observed | Mr(expt) | Mr(calc) | Delta | Miss | Score | Expect | Rank | Unique | Peptide |
|---|---|---|---|---|---|---|---|---|---|---|
| 14830 | 551.5288 | 1101.0431 | 1101.5051 | -0.4620 | 0 | 42 | 0.054 | 1 | U | R.SSPGQPSSSPR.L |

---

**52.** gi|26380166 **Mass:** 13950 **Score:** 42 **Matches:** 1(1) **Sequences:** 1(1) **emPAI:** 0.24
unnamed protein product [Mus musculus]

| Query | Observed | Mr(expt) | Mr(calc) | Delta | Miss | Score | Expect | Rank | Unique | Peptide |
|---|---|---|---|---|---|---|---|---|---|---|
| 17965 | 703.0179 | 1404.0212 | 1404.6885 | -0.6673 | 0 | 42 | 0.056 | 1 | U | K.GPSEGAYDVILPR.A |

**Proteins matching the same set of peptides:**
gi|46395891 **Mass:** 48390 **Score:** 42 **Matches:** 1(1) **Sequences:** 1(1)
RecName: Full=G-protein coupled receptor family C group 5 member C; AltName: Full=Retinoic acid-induced gene 3 protein; Short=RAIG-3; Flags: Precursor
gi|148669181 **Mass:** 48467 **Score:** 42 **Matches:** 1(1) **Sequences:** 1(1)
G protein-coupled receptor, family C, group 5, member C [Mus musculus]
gi|160333231 **Mass:** 48453 **Score:** 42 **Matches:** 1(1) **Sequences:** 1(1)
G-protein coupled receptor family C group 5 member C isoform a precursor [Mus musculus]

gi|160333233 **Mass:** 48416 **Score:** 42 **Matches:** 1(1) **Sequences:** 1(1)
G-protein coupled receptor family C group 5 member C isoform b precursor [Mus musculus]
gi|215406578 **Mass:** 11983 **Score:** 42 **Matches:** 1(1) **Sequences:** 1(1)
G protein-coupled receptor, family C, group 5, member C [Mus musculus]

---

**Mascot:** http://www.matrixscience.com/

PFHR9

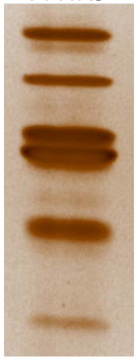

4

### Mascot Search Results

```
User                   :
Email                  :
Search title           :
MS data file           : ZIEN-29Nov2017_4m.mgf
Database               : NCBInr 20120419 (17893860 sequences; 6141683785 residues)
Taxonomy               : Mus musculus (house mouse) (138831 sequences)
Timestamp              : 30 Nov 2017 at 20:51:28 GMT
Enzyme                 : Trypsin
Variable modifications : Oxidation (M),Carbamidomethyl (C),Oxidation (P)
Mass values            : Monoisotopic
Protein Mass           : Unrestricted
Peptide Mass Tolerance : ± 1.25 Da
Fragment Mass Tolerance: ± 1.001 Da
Max Missed Cleavages   : 1
Instrument type        : Default
Number of queries      : 16414
Protein hits           : gi|22164776  keratin, type II cytoskeletal 79 [Mus musculus]
                         gi|51450     heat shock protein [Mus musculus]
                         gi|7638398   epidermal keratin 10 [Mus musculus]
                         gi|16716569  protease, serine, 1 precursor [Mus musculus]
                         gi|12843914  unnamed protein product [Mus musculus]
                         gi|9910294   keratin, type II cytoskeletal 71 [Mus musculus]
                         gi|2498741   RecName: Full=Prolyl 4-hydroxylase subunit alpha-2; Short=4-PH alpha-2; AltName: Full=Procollagen-proline,2-oxoglutarate-4-dioxygenase subunit alpha-2; Flags: Precursc
                         gi|148672076 mCG17605, isoform CRA_b [Mus musculus]
                         gi|51092303  Try10-like trypsinogen precursor [Mus musculus]
                         gi|4159806   type II keratin subunit protein [Mus musculus]
                         gi|6166378   growth suppressor 1L [Mus musculus]
                         gi|295841496 tubulin beta 3 [Mus musculus]
                         gi|438134    DNA-directed DNA polymerase [Mus musculus]
                         gi|6680888   CD63 antigen [Mus musculus]
                         gi|202210    alpha-tubulin isotype M-alpha-2 [Mus musculus]
                         gi|3687320   cartilage-associated protein (CASP) [Mus musculus]
                         gi|85701680  keratin, type II cytoskeletal 2 oral [Mus musculus]
                         gi|21489935  keratin, type I cytoskeletal 14 [Mus musculus]
                         gi|468546    CCT (chaperonin containing TCP-1) beta subunit [Mus musculus]
                         gi|6755893   trypsin 4 precursor [Mus musculus]
                         gi|27370092  elongation factor Tu, mitochondrial isoform 1 [Mus musculus]
                         gi|293686    epidermal keratin subunit II [Mus musculus]
                         gi|3212116   prefoldin subunit 2 [Mus musculus]
                         gi|193645    glucose-regulated protein 78, partial [Mus musculus]
                         gi|53830774  lactase [Mus musculus]
                         gi|81872882  RecName: Full=Adenomatous polyposis coli protein 2
                         gi|191765    alpha-fetoprotein, partial [Mus musculus]
                         gi|54777     unnamed protein product [Mus musculus]
                         gi|26390362  unnamed protein product [Mus musculus]
```

#### Select Summary Report

Format As | Select Summary (protein hits) | Help

Significance threshold p< 0.05 | Max. number of hits AUTO

Standard scoring ○ MudPIT scoring ● Ions score or expect cut-off 43 | Show sub-sets 0

Show pop-ups ● Suppress pop-ups ○ | Require bold red ☐

Re-Search ● All queries ○ Unassigned ○ Below homology threshold ○ Below identity threshold

**1.** gi|22164776 **Mass:** 57517 **Score:** 102 **Matches:** 2(2) **Sequences:** 1(1) **emPAI:** 0.06

keratin, type II cytoskeletal 79 [Mus musculus]

| Query | Observed | Mr(expt) | Mr(calc) | Delta | Miss | Score | Expect | Rank | Unique | Peptide |
|---|---|---|---|---|---|---|---|---|---|---|
| 14241 | 665.9768 | 1329.9391 | 1328.7187 | 1.2204 | 0 | 79 | 1.2e-005 | 1 | U | R.NLDLDSIIAEVK.A 14234 |

**Proteins matching the same set of peptides:**

gi|148672069 **Mass:** 59231 **Score:** 102 **Matches:** 2(2) **Sequences:** 1(1)
cDNA sequence BC031593 [Mus musculus]

---

2. gi|51450 **Mass:** 46560 **Score:** 102 **Matches:** 4(3) **Sequences:** 4(3) **emPAI:** 0.32
heat shock protein [Mus musculus]

| Query | Observed | Mr(expt) | Mr(calc) | Delta | Miss | Score | Expect | Rank | Unique | Peptide |
|---|---|---|---|---|---|---|---|---|---|---|
| 14017 | 653.5609 | 1305.1073 | 1305.6677 | -0.5604 | 0 | 67 | 0.00035 | 1 | U | R.DNQSGSLLFIGR.L |
| 14227 | 665.1298 | 1328.2451 | 1327.6628 | 0.5823 | 0 | 52 | 0.013 | 1 | U | K.LQMVEMPLAHK.L |
| 14757 | 690.9507 | 1379.8868 | 1379.5881 | 0.2987 | 0 | 44 | 0.097 | 1 | U | R.TGLYNYYDDEK.E |
| 16290 | 830.6638 | 1659.3131 | 1658.7941 | 0.5190 | 0 | 59 | 0.0027 | 1 | U | R.LYGPSSVSFADDFVR.S |

**Proteins matching the same set of peptides:**
gi|200966 **Mass:** 45623 **Score:** 102 **Matches:** 4(3) **Sequences:** 4(3)
put. serine protease inhibitor [Mus musculus]
gi|26345418 **Mass:** 46481 **Score:** 102 **Matches:** 4(3) **Sequences:** 4(3)
unnamed protein product [Mus musculus]
gi|26348007 **Mass:** 46490 **Score:** 102 **Matches:** 4(3) **Sequences:** 4(3)
unnamed protein product [Mus musculus]
gi|74191337 **Mass:** 46505 **Score:** 102 **Matches:** 4(3) **Sequences:** 4(3)
unnamed protein product [Mus musculus]
gi|74198254 **Mass:** 46500 **Score:** 102 **Matches:** 4(3) **Sequences:** 4(3)
unnamed protein product [Mus musculus]
gi|148684430 **Mass:** 44954 **Score:** 102 **Matches:** 4(3) **Sequences:** 4(3)
serine (or cysteine) peptidase inhibitor, clade H, member 1, isoform CRA_a [Mus musculus]
gi|161353502 **Mass:** 46504 **Score:** 102 **Matches:** 4(3) **Sequences:** 4(3)
serpin H1 precursor [Mus musculus]

---

3. gi|7638398 **Mass:** 34962 **Score:** 85 **Matches:** 2(2) **Sequences:** 2(2) **emPAI:** 0.20
epidermal keratin 10 [Mus musculus]

| Query | Observed | Mr(expt) | Mr(calc) | Delta | Miss | Score | Expect | Rank | Unique | Peptide |
|---|---|---|---|---|---|---|---|---|---|---|
| 11545 | 546.0249 | 1090.0351 | 1089.5237 | 0.5115 | 0 | 46 | 0.023 | 1 | | K.VTMQNLNDR.L |
| 14774 | 691.5409 | 1381.0672 | 1380.6408 | 0.4264 | 0 | 80 | 1.6e-005 | 1 | U | R.ALEESNYELEGK.I |

**Proteins matching the same set of peptides:**
gi|387397 **Mass:** 57807 **Score:** 85 **Matches:** 2(2) **Sequences:** 2(2)
epidermal keratin subunit I, partial [Mus musculus]
gi|12852157 **Mass:** 58587 **Score:** 85 **Matches:** 2(2) **Sequences:** 2(2)
unnamed protein product [Mus musculus]
gi|26349141 **Mass:** 52653 **Score:** 85 **Matches:** 2(2) **Sequences:** 2(2)
unnamed protein product [Mus musculus]
gi|26349459 **Mass:** 49471 **Score:** 85 **Matches:** 2(2) **Sequences:** 2(2)
unnamed protein product [Mus musculus]
gi|112983636 **Mass:** 57007 **Score:** 85 **Matches:** 2(2) **Sequences:** 2(2)
keratin, type I cytoskeletal 10 [Mus musculus]
gi|116242600 **Mass:** 57735 **Score:** 85 **Matches:** 2(2) **Sequences:** 2(2)
RecName: Full=Keratin, type I cytoskeletal 10; AltName: Full=56 kDa cytokeratin; AltName: Full=Cytokeratin-10; Short=CK-10; AltName: Full=Keratin, type I cytoskeletal 59 kDa; AltName: Full=Keratin-1[illegible]

---

4. gi|16716569 **Mass:** 26118 **Score:** 83 **Matches:** 2(1) **Sequences:** 1(1) **emPAI:** 0.13
protease, serine, 1 precursor [Mus musculus]

| Query | Observed | Mr(expt) | Mr(calc) | Delta | Miss | Score | Expect | Rank | Unique | Peptide |
|---|---|---|---|---|---|---|---|---|---|---|
| 15632 | 737.8900 | 2210.6482 | 2210.0967 | 0.5515 | 0 | (44) | 0.083 | 1 | U | R.LGEHNINVLEGNEQFIDAAK.I |
| 16412 | 1106.3871 | 2210.7596 | 2210.0967 | 0.6629 | 0 | 83 | 7.4e-006 | 1 | U | R.LGEHNINVLEGNEQFIDAAK.I |

---

5. gi|12843914 **Mass:** 29993 **Score:** 82 **Matches:** 1(1) **Sequences:** 1(1) **emPAI:** 0.11
unnamed protein product [Mus musculus]

| Query | Observed | Mr(expt) | Mr(calc) | Delta | Miss | Score | Expect | Rank | Unique | Peptide |
|---|---|---|---|---|---|---|---|---|---|---|
| 13984 | 651.6257 | 1301.2369 | 1301.7078 | -0.4709 | 0 | 82 | 1.5e-005 | 1 | U | R.SLDLDSIIAEVK.A |

**Proteins matching the same set of peptides:**
gi|13272554 **Mass:** 42357 **Score:** 82 **Matches:** 1(1) **Sequences:** 1(1)
cytokeratin KRT2-6HF [Mus musculus]
gi|16303309 **Mass:** 61743 **Score:** 82 **Matches:** 1(1) **Sequences:** 1(1)
type II keratin 5 [Mus musculus]
gi|20911031 **Mass:** 61729 **Score:** 82 **Matches:** 1(1) **Sequences:** 1(1)
keratin, type II cytoskeletal 5 [Mus musculus]
gi|29789317 **Mass:** 59704 **Score:** 82 **Matches:** 1(1) **Sequences:** 1(1)
keratin, type II cytoskeletal 75 [Mus musculus]
gi|54607171 **Mass:** 59299 **Score:** 82 **Matches:** 1(1) **Sequences:** 1(1)
keratin, type II cytoskeletal 6A [Mus musculus]
gi|59798479 **Mass:** 60285 **Score:** 82 **Matches:** 1(1) **Sequences:** 1(1)
RecName: Full=Keratin, type II cytoskeletal 6B; AltName: Full=Cytokeratin-6B; Short=CK-6B; AltName: Full=Keratin-6-beta; Short=mK6-beta; AltName: Full=Keratin-6B; Short=K6B
gi|110645788 **Mass:** 60236 **Score:** 82 **Matches:** 1(1) **Sequences:** 1(1)
Krt6b protein [Mus musculus]
gi|113195684 **Mass:** 59490 **Score:** 82 **Matches:** 1(1) **Sequences:** 1(1)
keratin, type II cytoskeletal 6B [Mus musculus]
gi|116063325 **Mass:** 60154 **Score:** 82 **Matches:** 1(1) **Sequences:** 1(1)
Krt6b protein [Mus musculus]
gi|148672084 **Mass:** 34775 **Score:** 82 **Matches:** 1(1) **Sequences:** 1(1)
mCG17577 [Mus musculus]
gi|148672089 **Mass:** 62241 **Score:** 82 **Matches:** 1(1) **Sequences:** 1(1)
keratin 75 [Mus musculus]
gi|148672085 **Mass:** 38119 **Score:** 82 **Matches:** 1(1) **Sequences:** 1(1)
mCG144996 [Mus musculus]

---

**6.** gi|9910294 **Mass:** 57347 **Score:** 75 **Matches:** 2(2) **Sequences:** 2(2) **emPAI:** 0.12
keratin, type II cytoskeletal 71 [Mus musculus]

| Query | Observed | Mr(expt) | Mr(calc) | Delta | Miss | Score | Expect | Rank | Unique | Peptide |
|---|---|---|---|---|---|---|---|---|---|---|
| 11976 | 562.3885 | 1122.7624 | 1122.5557 | 0.2068 | 0 | 49 | 0.029 | 1 | | K.AEAEALYQTK.F |
| 15642 | 738.2304 | 1474.4461 | 1474.7780 | -0.3318 | 0 | 65 | 0.00053 | 1 | U | R.FLEQQNQVLQTK.W |

**Proteins matching the same set of peptides:**
gi|33146295 **Mass:** 57391 **Score:** 75 **Matches:** 2(2) **Sequences:** 2(2)
type II keratin [Mus musculus]
gi|148672081 **Mass:** 43429 **Score:** 75 **Matches:** 2(2) **Sequences:** 2(2)
mCG17589, isoform CRA_b [Mus musculus]

---

**7.** gi|2498741 **Mass:** 60964 **Score:** 66 **Matches:** 1(1) **Sequences:** 1(1) **emPAI:** 0.05
RecName: Full=Prolyl 4-hydroxylase subunit alpha-2; Short=4-PH alpha-2; AltName: Full=Procollagen-proline,2-oxoglutarate-4-dioxygenase subunit alpha-2; Flags: Precursor

| Query | Observed | Mr(expt) | Mr(calc) | Delta | Miss | Score | Expect | Rank | Unique | Peptide |
|---|---|---|---|---|---|---|---|---|---|---|
| 16157 | 785.1390 | 1568.2634 | 1568.6743 | -0.4109 | 0 | 66 | 0.00051 | 1 | U | R.QFFPTDEDESGAAR.A |

**Proteins matching the same set of peptides:**
gi|148701596 **Mass:** 40313 **Score:** 66 **Matches:** 1(1) **Sequences:** 1(1)
procollagen-proline, 2-oxoglutarate 4-dioxygenase (proline 4-hydroxylase), alpha II polypeptide, isoform CRA_a [Mus musculus]
gi|148701597 **Mass:** 57417 **Score:** 66 **Matches:** 1(1) **Sequences:** 1(1)
procollagen-proline, 2-oxoglutarate 4-dioxygenase (proline 4-hydroxylase), alpha II polypeptide, isoform CRA_b [Mus musculus]
gi|148701599 **Mass:** 61292 **Score:** 66 **Matches:** 1(1) **Sequences:** 1(1)
procollagen-proline, 2-oxoglutarate 4-dioxygenase (proline 4-hydroxylase), alpha II polypeptide, isoform CRA_d [Mus musculus]
gi|148701600 **Mass:** 66870 **Score:** 66 **Matches:** 1(1) **Sequences:** 1(1)
procollagen-proline, 2-oxoglutarate 4-dioxygenase (proline 4-hydroxylase), alpha II polypeptide, isoform CRA_e [Mus musculus]
gi|209862961 **Mass:** 60774 **Score:** 66 **Matches:** 1(1) **Sequences:** 1(1)
prolyl 4-hydroxylase subunit alpha-2 isoform 1 precursor [Mus musculus]
gi|226874876 **Mass:** 60977 **Score:** 66 **Matches:** 1(1) **Sequences:** 1(1)
prolyl 4-hydroxylase subunit alpha-2 isoform 2 precursor [Mus musculus]

---

**8.** gi|148672076 **Mass:** 72140 **Score:** 65 **Matches:** 2(1) **Sequences:** 2(1) **emPAI:** 0.09

mCG17605, isoform CRA_b [Mus musculus]

| Query | Observed | Mr(expt) | Mr(calc) | Delta | Miss | Score | Expect | Rank | Unique | Peptide |
|---|---|---|---|---|---|---|---|---|---|---|
| 11007 | 521.2697 | 1040.5249 | 1040.5502 | -0.0253 | 0 | 46 | 0.067 | 1 | U | K.VDPEIQNVK.S |
| 13545 | 628.0165 | 1254.0184 | 1253.6001 | 0.4184 | 0 | 65 | 0.0007 | 1 | U | R.GFSSGSAVVSGGSR.R |

**Proteins matching the same set of peptides:**

gi|398168 **Mass:** 70934 **Score:** 65 **Matches:** 2(1) **Sequences:** 2(1)
keratin 2 epidermis [Mus musculus]
gi|111308159 **Mass:** 70880 **Score:** 65 **Matches:** 2(1) **Sequences:** 2(1)
Keratin 2 [Mus musculus]
gi|124487419 **Mass:** 70880 **Score:** 65 **Matches:** 2(1) **Sequences:** 2(1)
keratin, type II cytoskeletal 2 epidermal [Mus musculus]
gi|148672075 **Mass:** 71783 **Score:** 65 **Matches:** 2(1) **Sequences:** 2(1)
mCG17605, isoform CRA_a [Mus musculus]

---

**9.** gi|51092303 **Mass:** 26514 **Score:** 65 **Matches:** 1(1) **Sequences:** 1(1) **emPAI:** 0.13
Try10-like trypsinogen precursor [Mus musculus]

| Query | Observed | Mr(expt) | Mr(calc) | Delta | Miss | Score | Expect | Rank | Unique | Peptide |
|---|---|---|---|---|---|---|---|---|---|---|
| 12639 | 588.3151 | 1174.6156 | 1174.6267 | -0.0112 | 0 | 65 | 0.00087 | 1 | U | K.TLDNDIMLIK.L |

**Proteins matching the same set of peptides:**

gi|84781771 **Mass:** 26204 **Score:** 65 **Matches:** 1(1) **Sequences:** 1(1)
trypsin 10 precursor [Mus musculus]

---

**10.** gi|4159806 **Mass:** 65183 **Score:** 64 **Matches:** 1(1) **Sequences:** 1(1) **emPAI:** 0.05
type II keratin subunit protein [Mus musculus]

| Query | Observed | Mr(expt) | Mr(calc) | Delta | Miss | Score | Expect | Rank | Unique | Peptide |
|---|---|---|---|---|---|---|---|---|---|---|
| 15642 | 738.2304 | 1474.4461 | 1474.8144 | -0.3682 | 1 | 64 | 0.0007 | 2 | U | R.FLEQQNKVLQTK.W |

---

**11.** gi|6166378 **Mass:** 84758 **Score:** 58 **Matches:** 1(1) **Sequences:** 1(1) **emPAI:** 0.04
growth suppressor 1L [Mus musculus]

| Query | Observed | Mr(expt) | Mr(calc) | Delta | Miss | Score | Expect | Rank | Unique | Peptide |
|---|---|---|---|---|---|---|---|---|---|---|
| 15783 | 747.1336 | 1492.2527 | 1491.7398 | 0.5129 | 0 | 58 | 0.0013 | 1 | U | R.SPYNYLQVAYFK.I |

**Proteins matching the same set of peptides:**

gi|6166380 **Mass:** 61560 **Score:** 58 **Matches:** 1(1) **Sequences:** 1(1)
growth suppressor 1S [Mus musculus]
gi|12846125 **Mass:** 84033 **Score:** 58 **Matches:** 1(1) **Sequences:** 1(1)
unnamed protein product [Mus musculus]
gi|23271416 **Mass:** 83540 **Score:** 58 **Matches:** 1(1) **Sequences:** 1(1)
Leprecan 1 [Mus musculus]
gi|26326437 **Mass:** 83476 **Score:** 58 **Matches:** 1(1) **Sequences:** 1(1)
unnamed protein product [Mus musculus]
gi|74205527 **Mass:** 84439 **Score:** 58 **Matches:** 1(1) **Sequences:** 1(1)
unnamed protein product [Mus musculus]
gi|109150433 **Mass:** 83598 **Score:** 58 **Matches:** 1(1) **Sequences:** 1(1)
prolyl 3-hydroxylase 1 isoform 2 precursor [Mus musculus]
gi|109150437 **Mass:** 84426 **Score:** 58 **Matches:** 1(1) **Sequences:** 1(1)
prolyl 3-hydroxylase 1 isoform 1 precursor [Mus musculus]
gi|148698509 **Mass:** 84378 **Score:** 58 **Matches:** 1(1) **Sequences:** 1(1)
leprecan 1, isoform CRA_a [Mus musculus]
gi|148698510 **Mass:** 83550 **Score:** 58 **Matches:** 1(1) **Sequences:** 1(1)
leprecan 1, isoform CRA_b [Mus musculus]
gi|148698511 **Mass:** 63605 **Score:** 58 **Matches:** 1(1) **Sequences:** 1(1)
leprecan 1, isoform CRA_c [Mus musculus]

---

**12.** gi|295841496 **Mass:** 14518 **Score:** 58 **Matches:** 2(2) **Sequences:** 1(1) **emPAI:** 0.23

tubulin beta 3 [Mus musculus]

| Query | Observed | Mr(expt) | Mr(calc) | Delta | Miss | Score | Expect | Rank | Unique | Peptide |
|---|---|---|---|---|---|---|---|---|---|---|
| 14141 | 660.4166 | 1318.8186 | 1318.6955 | 0.1231 | 0 | 52 | 0.016 | 1 | U | R.IMNTFSVVPSPK.V 14142 |

**Proteins matching the same set of peptides:**

gi|202229 **Mass:** 42212 **Score:** 58 **Matches:** 2(2) **Sequences:** 1(1)
beta-tubulin, partial [Mus musculus]
gi|5174735 **Mass:** 49799 **Score:** 58 **Matches:** 2(2) **Sequences:** 1(1)
tubulin beta-4B chain [Homo sapiens]
gi|7106439 **Mass:** 49639 **Score:** 58 **Matches:** 2(2) **Sequences:** 1(1)
tubulin beta-5 chain [Mus musculus]
gi|12846758 **Mass:** 49608 **Score:** 58 **Matches:** 2(2) **Sequences:** 1(1)
unnamed protein product [Mus musculus]
gi|12851187 **Mass:** 49496 **Score:** 58 **Matches:** 2(2) **Sequences:** 1(1)
unnamed protein product [Mus musculus]
gi|12963615 **Mass:** 50386 **Score:** 58 **Matches:** 2(2) **Sequences:** 1(1)
tubulin beta-3 chain [Mus musculus]
gi|13542680 **Mass:** 49783 **Score:** 58 **Matches:** 2(2) **Sequences:** 1(1)
Tubulin, beta 2C [Mus musculus]
gi|21361322 **Mass:** 49554 **Score:** 58 **Matches:** 2(2) **Sequences:** 1(1)
tubulin beta-4A chain [Homo sapiens]
gi|26355849 **Mass:** 32238 **Score:** 58 **Matches:** 2(2) **Sequences:** 1(1)
unnamed protein product [Mus musculus]
gi|26356799 **Mass:** 34123 **Score:** 58 **Matches:** 2(2) **Sequences:** 1(1)
unnamed protein product [Mus musculus]
gi|74141821 **Mass:** 49667 **Score:** 58 **Matches:** 2(2) **Sequences:** 1(1)
unnamed protein product [Mus musculus]
gi|74144588 **Mass:** 49771 **Score:** 58 **Matches:** 2(2) **Sequences:** 1(1)
unnamed protein product [Mus musculus]
gi|74204140 **Mass:** 49616 **Score:** 58 **Matches:** 2(2) **Sequences:** 1(1)
unnamed protein product [Mus musculus]
gi|74212109 **Mass:** 49620 **Score:** 58 **Matches:** 2(2) **Sequences:** 1(1)
unnamed protein product [Mus musculus]
gi|74223737 **Mass:** 49652 **Score:** 58 **Matches:** 2(2) **Sequences:** 1(1)
unnamed protein product [Mus musculus]
gi|74223783 **Mass:** 50402 **Score:** 58 **Matches:** 2(2) **Sequences:** 1(1)
unnamed protein product [Mus musculus]
gi|148676266 **Mass:** 39342 **Score:** 58 **Matches:** 2(2) **Sequences:** 1(1)
mCG20287 [Mus musculus]
gi|148678613 **Mass:** 49970 **Score:** 58 **Matches:** 2(2) **Sequences:** 1(1)
mCG1424 [Mus musculus]
gi|148682710 **Mass:** 49784 **Score:** 58 **Matches:** 2(2) **Sequences:** 1(1)
mCG49614, isoform CRA_b [Mus musculus]
gi|148691289 **Mass:** 49910 **Score:** 58 **Matches:** 2(2) **Sequences:** 1(1)
tubulin, beta 5 [Mus musculus]

---

**13.** gi|438134 **Mass:** 123705 **Score:** 57 **Matches:** 1(1) **Sequences:** 1(1) **emPAI:** 0.03
DNA-directed DNA polymerase [Mus musculus]

| Query | Observed | Mr(expt) | Mr(calc) | Delta | Miss | Score | Expect | Rank | Unique | Peptide |
|---|---|---|---|---|---|---|---|---|---|---|
| 11720 | 553.2844 | 1104.5543 | 1104.6040 | -0.0497 | 1 | 57 | 0.0047 | 1 | U | R.RQGPGPGVPPK.R |

**Proteins matching the same set of peptides:**

gi|2827903 **Mass:** 123657 **Score:** 57 **Matches:** 1(1) **Sequences:** 1(1)
DNA polymerase delta catalytic subunit [Mus musculus]
gi|14318657 **Mass:** 123666 **Score:** 57 **Matches:** 1(1) **Sequences:** 1(1)
Polymerase (DNA directed), delta 1, catalytic subunit [Mus musculus]
gi|26353290 **Mass:** 123698 **Score:** 57 **Matches:** 1(1) **Sequences:** 1(1)
unnamed protein product [Mus musculus]

gi|148690782 **Mass:** 125906 **Score:** 57 **Matches:** 1(1) **Sequences:** 1(1)
polymerase (DNA directed), delta 1, catalytic subunit, isoform CRA_a [Mus musculus]
gi|148690783 **Mass:** 90830 **Score:** 57 **Matches:** 1(1) **Sequences:** 1(1)
polymerase (DNA directed), delta 1, catalytic subunit, isoform CRA_b [Mus musculus]
gi|148690785 **Mass:** 122966 **Score:** 57 **Matches:** 1(1) **Sequences:** 1(1)
polymerase (DNA directed), delta 1, catalytic subunit, isoform CRA_d [Mus musculus]
gi|148690786 **Mass:** 123682 **Score:** 57 **Matches:** 1(1) **Sequences:** 1(1)
polymerase (DNA directed), delta 1, catalytic subunit, isoform CRA_e [Mus musculus]
gi|148690787 **Mass:** 107166 **Score:** 57 **Matches:** 1(1) **Sequences:** 1(1)
polymerase (DNA directed), delta 1, catalytic subunit, isoform CRA_f [Mus musculus]
gi|254587977 **Mass:** 123712 **Score:** 57 **Matches:** 1(1) **Sequences:** 1(1)
DNA polymerase delta catalytic subunit [Mus musculus]

---

**14.** gi|6680888 **Mass:** 25749 **Score:** 56 **Matches:** 1(1) **Sequences:** 1(1) **emPAI:** 0.13
CD63 antigen [Mus musculus]

| Query | Observed | Mr(expt) | Mr(calc) | Delta | Miss | Score | Expect | Rank | Unique | Peptide |
|---|---|---|---|---|---|---|---|---|---|---|
| 15128 | 708.0230 | 1414.0315 | 1413.6711 | 0.3604 | 0 | 56 | 0.0049 | 1 | U | K.SFQQQMQNYLK.D |

**Proteins matching the same set of peptides:**
gi|26389974 **Mass:** 26766 **Score:** 56 **Matches:** 1(1) **Sequences:** 1(1)
unnamed protein product [Mus musculus]
gi|74151897 **Mass:** 25777 **Score:** 56 **Matches:** 1(1) **Sequences:** 1(1)
unnamed protein product [Mus musculus]

---

**15.** gi|202210 **Mass:** 50134 **Score:** 55 **Matches:** 1(1) **Sequences:** 1(1) **emPAI:** 0.07
alpha-tubulin isotype M-alpha-2 [Mus musculus]

| Query | Observed | Mr(expt) | Mr(calc) | Delta | Miss | Score | Expect | Rank | Unique | Peptide |
|---|---|---|---|---|---|---|---|---|---|---|
| 10750 | 508.1963 | 1014.3781 | 1014.5709 | -0.1928 | 0 | 55 | 0.004 | 1 | U | K.DVNAAIATIK.T |

**Proteins matching the same set of peptides:**
gi|202223 **Mass:** 22209 **Score:** 55 **Matches:** 1(1) **Sequences:** 1(1)
alpha-tubulin, partial [Mus musculus]
gi|202225 **Mass:** 39293 **Score:** 55 **Matches:** 1(1) **Sequences:** 1(1)
alpha-tubulin, partial [Mus musculus]
gi|6678465 **Mass:** 49928 **Score:** 55 **Matches:** 1(1) **Sequences:** 1(1)
tubulin alpha-3 chain [Mus musculus]
gi|6678469 **Mass:** 49877 **Score:** 55 **Matches:** 1(1) **Sequences:** 1(1)
tubulin alpha-1C chain [Mus musculus]
gi|6755901 **Mass:** 50104 **Score:** 55 **Matches:** 1(1) **Sequences:** 1(1)
tubulin alpha-1A chain [Mus musculus]
gi|12839396 **Mass:** 44021 **Score:** 55 **Matches:** 1(1) **Sequences:** 1(1)
unnamed protein product [Mus musculus]
gi|12850141 **Mass:** 38332 **Score:** 55 **Matches:** 1(1) **Sequences:** 1(1)
unnamed protein product [Mus musculus]
gi|13435888 **Mass:** 38746 **Score:** 55 **Matches:** 1(1) **Sequences:** 1(1)
Tuba1b protein [Mus musculus]
gi|34740335 **Mass:** 50120 **Score:** 55 **Matches:** 1(1) **Sequences:** 1(1)
tubulin alpha-1B chain [Mus musculus]
gi|53733821 **Mass:** 50120 **Score:** 55 **Matches:** 1(1) **Sequences:** 1(1)
Tubulin, alpha 1A [Mus musculus]
gi|74181454 **Mass:** 50104 **Score:** 55 **Matches:** 1(1) **Sequences:** 1(1)
unnamed protein product [Mus musculus]
gi|74186501 **Mass:** 50108 **Score:** 55 **Matches:** 1(1) **Sequences:** 1(1)
unnamed protein product [Mus musculus]
gi|74198443 **Mass:** 49907 **Score:** 55 **Matches:** 1(1) **Sequences:** 1(1)
unnamed protein product [Mus musculus]
gi|74198980 **Mass:** 50048 **Score:** 55 **Matches:** 1(1) **Sequences:** 1(1)
unnamed protein product [Mus musculus]
gi|74216624 **Mass:** 50116 **Score:** 55 **Matches:** 1(1) **Sequences:** 1(1)

unnamed protein product [Mus musculus]
gi|74220042 **Mass:** 50105 **Score:** 55 **Matches:** 1(1) **Sequences:** 1(1)
unnamed protein product [Mus musculus]
gi|148672205 **Mass:** 50888 **Score:** 55 **Matches:** 1(1) **Sequences:** 1(1)
mCG18413, isoform CRA_b [Mus musculus]
gi|148704842 **Mass:** 43956 **Score:** 55 **Matches:** 1(1) **Sequences:** 1(1)
mCG125663 [Mus musculus]

---

**16.** gi|3687320 **Mass:** 46137 **Score:** 54 **Matches:** 1(1) **Sequences:** 1(1) **emPAI:** 0.07
cartilage-associated protein (CASP) [Mus musculus]

| Query | Observed | Mr(expt) | Mr(calc) | Delta | Miss | Score | Expect | Rank | Unique | Peptide |
|---|---|---|---|---|---|---|---|---|---|---|
| 11191 | **532.0100** | **1062.0053** | **1062.5458** | **-0.5405** | **0** | **54** | **0.0083** | **1** | **U** | **R.SVLADFQQR.E** |

**Proteins matching the same set of peptides:**
gi|12857227 **Mass:** 46121 **Score:** 54 **Matches:** 1(1) **Sequences:** 1(1)
unnamed protein product [Mus musculus]
gi|148677365 **Mass:** 46110 **Score:** 54 **Matches:** 1(1) **Sequences:** 1(1)
mCG7119 [Mus musculus]
gi|225543173 **Mass:** 46140 **Score:** 54 **Matches:** 1(1) **Sequences:** 1(1)
cartilage-associated protein precursor [Mus musculus]

---

**17.** gi|85701680 **Mass:** 62806 **Score:** 53 **Matches:** 2(2) **Sequences:** 2(2) **emPAI:** 0.11
keratin, type II cytoskeletal 2 oral [Mus musculus]

| Query | Observed | Mr(expt) | Mr(calc) | Delta | Miss | Score | Expect | Rank | Unique | Peptide |
|---|---|---|---|---|---|---|---|---|---|---|
| 11227 | **1065.4674** | **1064.4601** | **1064.5138** | **-0.0537** | **0** | **43** | **0.058** | **1** | **U** | **K.AQYEDIAQK.S** |
| 11976 | 562.3885 | 1122.7624 | 1122.5557 | 0.2068 | 0 | 49 | 0.029 | 1 | | K.AEAEALYQTK.L |

**Proteins matching the same set of peptides:**
gi|148672072 **Mass:** 63519 **Score:** 53 **Matches:** 2(2) **Sequences:** 2(2)
mCG144546 [Mus musculus]

---

**18.** gi|21489935 **Mass:** 52834 **Score:** 52 **Matches:** 2(2) **Sequences:** 2(2) **emPAI:** 0.13
keratin, type I cytoskeletal 14 [Mus musculus]

| Query | Observed | Mr(expt) | Mr(calc) | Delta | Miss | Score | Expect | Rank | Unique | Peptide |
|---|---|---|---|---|---|---|---|---|---|---|
| 10856 | **515.0688** | **1028.1230** | **1028.5866** | **-0.4635** | **0** | **45** | **0.03** | **1** | **U** | **R.VLDELTLAR.A** |
| 11545 | 546.0249 | 1090.0351 | 1089.5237 | 0.5115 | 0 | 46 | 0.023 | 1 | | K.VTMQNLNDR.L |

**Proteins matching the same set of peptides:**
gi|148670626 **Mass:** 54668 **Score:** 52 **Matches:** 2(2) **Sequences:** 2(2)
mCG144006 [Mus musculus]
gi|741022 **Mass:** 49086 **Score:** 52 **Matches:** 2(2) **Sequences:** 2(2)
keratin 15
gi|904215 **Mass:** 49129 **Score:** 52 **Matches:** 2(2) **Sequences:** 2(2)
cytokeratin 15 [Mus musculus]
gi|83304265 **Mass:** 49107 **Score:** 52 **Matches:** 2(2) **Sequences:** 2(2)
RecName: Full=Keratin, type I cytoskeletal 15; AltName: Full=Cytokeratin-15; Short=CK-15; AltName: Full=Keratin-15; Short=K15
gi|226823220 **Mass:** 49463 **Score:** 52 **Matches:** 2(2) **Sequences:** 2(2)
keratin, type I cytoskeletal 15 [Mus musculus]

---

**19.** gi|468546 **Mass:** 57411 **Score:** 51 **Matches:** 1(1) **Sequences:** 1(1) **emPAI:** 0.06
CCT (chaperonin containing TCP-1) beta subunit [Mus musculus]

| Query | Observed | Mr(expt) | Mr(calc) | Delta | Miss | Score | Expect | Rank | Unique | Peptide |
|---|---|---|---|---|---|---|---|---|---|---|
| 16188 | **791.9488** | **1581.8830** | **1581.9090** | **-0.0259** | **0** | **51** | **0.017** | **1** | **U** | **R.QVLLSAAEAAEVILR.V** |

**Proteins matching the same set of peptides:**
gi|7670405 **Mass:** 52436 **Score:** 51 **Matches:** 1(1) **Sequences:** 1(1)
unnamed protein product [Mus musculus]

gi|126521835 **Mass:** 57441 **Score:** 51 **Matches:** 1(1) **Sequences:** 1(1)
T-complex protein 1 subunit beta [Mus musculus]

---

**20.** gi|6755893 **Score:** 50 **Matches:** 1(1) **Sequences:** 1(1) **emPAI:** 0.12
trypsin 4 precursor [Mus musculus]

| Query | Observed | Mr(expt) | Mr(calc) | Delta | Miss | Score | Expect | Rank | Unique | Peptide |
|---|---|---|---|---|---|---|---|---|---|---|
| 16412 | 1106.3871 | 2210.7596 | 2211.0920 | -0.3323 | 0 | 50 | 0.017 | 2 | U | R.LGEHNINVLEGNEQFVNSAK.I |

**Proteins matching the same set of peptides:**
gi|51010909 **Score:** 50 **Matches:** 1(1) **Sequences:** 1(1)
gi|74203392 **Score:** 50 **Matches:** 1(1) **Sequences:** 1(1)

---

**21.** gi|27370092 **Mass:** 49477 **Score:** 50 **Matches:** 1(1) **Sequences:** 1(1) **emPAI:** 0.07
elongation factor Tu, mitochondrial isoform 1 [Mus musculus]

| Query | Observed | Mr(expt) | Mr(calc) | Delta | Miss | Score | Expect | Rank | Unique | Peptide |
|---|---|---|---|---|---|---|---|---|---|---|
| 12768 | **593.2388** | **1184.4630** | **1184.6149** | **-0.1519** | **0** | **50** | **0.027** | **1** | **U** | **R.AEAGDNLGALVR.G** |

**Proteins matching the same set of peptides:**
gi|148685430 **Mass:** 43302 **Score:** 50 **Matches:** 1(1) **Sequences:** 1(1)
mCG22399, isoform CRA_e [Mus musculus]
gi|148696761 **Mass:** 16047 **Score:** 50 **Matches:** 1(1) **Sequences:** 1(1)
mCG1048875, isoform CRA_a [Mus musculus]
gi|148696762 **Mass:** 12433 **Score:** 50 **Matches:** 1(1) **Sequences:** 1(1)
mCG1048875, isoform CRA_b [Mus musculus]
gi|148696763 **Mass:** 49507 **Score:** 50 **Matches:** 1(1) **Sequences:** 1(1)
mCG1048875, isoform CRA_c [Mus musculus]
gi|254911131 **Mass:** 47193 **Score:** 50 **Matches:** 1(1) **Sequences:** 1(1)
elongation factor Tu, mitochondrial isoform 2 [Mus musculus]
gi|148685429 **Mass:** 49613 **Score:** 50 **Matches:** 1(1) **Sequences:** 1(1)
mCG22399, isoform CRA_d [Mus musculus]

---

**22.** gi|293686 **Mass:** 59414 **Score:** 49 **Matches:** 1(1) **Sequences:** 1(1) **emPAI:** 0.06
epidermal keratin subunit II [Mus musculus]

| Query | Observed | Mr(expt) | Mr(calc) | Delta | Miss | Score | Expect | Rank | Unique | Peptide |
|---|---|---|---|---|---|---|---|---|---|---|
| 14931 | **698.0006** | **1393.9867** | **1393.7089** | **0.2778** | **1** | **49** | **0.025** | **1** | **U** | **R.TDAENEFVTLKK.D** |

---

**23.** gi|3212116 **Mass:** 16766 **Score:** 49 **Matches:** 1(1) **Sequences:** 1(1) **emPAI:** 0.20
prefoldin subunit 2 [Mus musculus]

| Query | Observed | Mr(expt) | Mr(calc) | Delta | Miss | Score | Expect | Rank | Unique | Peptide |
|---|---|---|---|---|---|---|---|---|---|---|
| 14659 | **686.9688** | **1371.9231** | **1370.7769** | **1.1462** | **0** | **49** | **0.014** | **1** | **U** | **K.IIETLSQQLQAK.G** |

**Proteins matching the same set of peptides:**
gi|31981577 **Mass:** 16524 **Score:** 49 **Matches:** 1(1) **Sequences:** 1(1)
prefoldin subunit 2 [Mus musculus]
gi|148707143 **Mass:** 11754 **Score:** 49 **Matches:** 1(1) **Sequences:** 1(1)
prefoldin 2, isoform CRA_b [Mus musculus]

---

**24.** gi|193645 **Mass:** 18932 **Score:** 46 **Matches:** 1(1) **Sequences:** 1(1) **emPAI:** 0.18
glucose-regulated protein 78, partial [Mus musculus]

| Query | Observed | Mr(expt) | Mr(calc) | Delta | Miss | Score | Expect | Rank | Unique | Peptide |
|---|---|---|---|---|---|---|---|---|---|---|
| 14961 | **699.5220** | **1397.0295** | **1396.7813** | **0.2482** | **0** | **46** | **0.041** | **1** | **U** | **K.ELEEIVQPIISK.L** |

**Proteins matching the same set of peptides:**
gi|387113 **Mass:** 16164 **Score:** 46 **Matches:** 1(1) **Sequences:** 1(1)
immunoglobulin heavy chain binding protein, partial [Mus musculus]
gi|1304157 **Mass:** 72412 **Score:** 46 **Matches:** 1(1) **Sequences:** 1(1)
78 kDa glucose-regulated protein [Mus musculus]

gi|2598562 **Mass:** 72433 **Score:** 46 **Matches:** 1(1) **Sequences:** 1(1)
BiP [Mus musculus]
gi|12835845 **Mass:** 72378 **Score:** 46 **Matches:** 1(1) **Sequences:** 1(1)
unnamed protein product [Mus musculus]
gi|74188814 **Mass:** 72305 **Score:** 46 **Matches:** 1(1) **Sequences:** 1(1)
unnamed protein product [Mus musculus]
gi|74198293 **Mass:** 72419 **Score:** 46 **Matches:** 1(1) **Sequences:** 1(1)
unnamed protein product [Mus musculus]
gi|74198974 **Mass:** 72361 **Score:** 46 **Matches:** 1(1) **Sequences:** 1(1)
unnamed protein product [Mus musculus]
gi|74207492 **Mass:** 72301 **Score:** 46 **Matches:** 1(1) **Sequences:** 1(1)
unnamed protein product [Mus musculus]
gi|74225394 **Mass:** 72337 **Score:** 46 **Matches:** 1(1) **Sequences:** 1(1)
unnamed protein product [Mus musculus]
gi|148676670 **Mass:** 56257 **Score:** 46 **Matches:** 1(1) **Sequences:** 1(1)
heat shock 70kD protein 5 (glucose-regulated protein), isoform CRA_b [Mus musculus]
gi|254540166 **Mass:** 72377 **Score:** 46 **Matches:** 1(1) **Sequences:** 1(1)
78 kDa glucose-regulated protein precursor [Mus musculus]

---

**25.** gi|53830774 **Mass:** 34092 **Score:** 46 **Matches:** 1(1) **Sequences:** 1(1) **emPAI:** 0.10
lactase [Mus musculus]

| Query | Observed | Mr(expt) | Mr(calc) | Delta | Miss | Score | Expect | Rank | Unique | Peptide |
|---|---|---|---|---|---|---|---|---|---|---|
| 11588 | 548.0338 | 1094.0531 | 1094.5132 | -0.4600 | 0 | 46 | 0.018 | 1 | U | R.LPEFTESEK.K |

**Proteins matching the same set of peptides:**
gi|74192292 **Mass:** 139170 **Score:** 46 **Matches:** 1(1) **Sequences:** 1(1)
unnamed protein product [Mus musculus]
gi|124487297 **Mass:** 217665 **Score:** 46 **Matches:** 1(1) **Sequences:** 1(1)
lactase-phlorizin hydrolase preproprotein [Mus musculus]
gi|148707805 **Mass:** 217693 **Score:** 46 **Matches:** 1(1) **Sequences:** 1(1)
mCG128560 [Mus musculus]

---

**26.** gi|81872882 **Mass:** 242989 **Score:** 46 **Matches:** 1(1) **Sequences:** 1(1) **emPAI:** 0.01
RecName: Full=Adenomatous polyposis coli protein 2

| Query | Observed | Mr(expt) | Mr(calc) | Delta | Miss | Score | Expect | Rank | Unique | Peptide |
|---|---|---|---|---|---|---|---|---|---|---|
| 15985 | 762.1919 | 2283.5538 | 2284.1344 | -0.5806 | 1 | 46 | 0.042 | 1 | U | K.YQAAAMAVSPGTCVPSLYVRK.Q |

**Proteins matching the same set of peptides:**
gi|117938322 **Mass:** 242963 **Score:** 46 **Matches:** 1(1) **Sequences:** 1(1)
adenomatous polyposis coli protein 2 [Mus musculus]

---

**27.** gi|191765 **Mass:** 47195 **Score:** 45 **Matches:** 1(0) **Sequences:** 1(0) **emPAI:** 0.07
alpha-fetoprotein, partial [Mus musculus]

| Query | Observed | Mr(expt) | Mr(calc) | Delta | Miss | Score | Expect | Rank | Unique | Peptide |
|---|---|---|---|---|---|---|---|---|---|---|
| 15680 | 740.7523 | 1479.4900 | 1478.7881 | 0.7018 | 0 | 45 | 0.065 | 1 | U | K.LGEYGFQNAILVR.Y |

**Proteins matching the same set of peptides:**
gi|26340966 **Mass:** 68678 **Score:** 45 **Matches:** 1(0) **Sequences:** 1(0)
unnamed protein product [Mus musculus]
gi|26341396 **Mass:** 64961 **Score:** 45 **Matches:** 1(0) **Sequences:** 1(0)
unnamed protein product [Mus musculus]
gi|74137565 **Mass:** 68688 **Score:** 45 **Matches:** 1(0) **Sequences:** 1(0)
unnamed protein product [Mus musculus]
gi|163310765 **Mass:** 68648 **Score:** 45 **Matches:** 1(0) **Sequences:** 1(0)
serum albumin precursor [Mus musculus]

---

**28.** gi|54777 **Mass:** 57108 **Score:** 45 **Matches:** 1(0) **Sequences:** 1(0) **emPAI:** 0.06
unnamed protein product [Mus musculus]

| Query | Observed | Mr(expt) | Mr(calc) | Delta | Miss | Score | Expect | Rank | Unique | Peptide |
|---|---|---|---|---|---|---|---|---|---|---|
| 13901 | 647.1276 | 1292.2406 | 1292.5918 | -0.3512 | 0 | 45 | 0.055 | 1 | U | K.MDSTANEVEAVK.V |

**Proteins matching the same set of peptides:**

gi\|42415475 **Mass:** 57023 **Score:** 45 **Matches:** 1(0) **Sequences:** 1(0)
protein disulfide-isomerase precursor [Mus musculus]
gi\|74138891 **Mass:** 56951 **Score:** 45 **Matches:** 1(0) **Sequences:** 1(0)
unnamed protein product [Mus musculus]
gi\|74141920 **Mass:** 57053 **Score:** 45 **Matches:** 1(0) **Sequences:** 1(0)
unnamed protein product [Mus musculus]
gi\|74178069 **Mass:** 57099 **Score:** 45 **Matches:** 1(0) **Sequences:** 1(0)
unnamed protein product [Mus musculus]
gi\|74190076 **Mass:** 56963 **Score:** 45 **Matches:** 1(0) **Sequences:** 1(0)
unnamed protein product [Mus musculus]
gi\|74191500 **Mass:** 57022 **Score:** 45 **Matches:** 1(0) **Sequences:** 1(0)
unnamed protein product [Mus musculus]
gi\|74198312 **Mass:** 57051 **Score:** 45 **Matches:** 1(0) **Sequences:** 1(0)
unnamed protein product [Mus musculus]
gi\|74198632 **Mass:** 16308 **Score:** 45 **Matches:** 1(0) **Sequences:** 1(0)
unnamed protein product [Mus musculus]
gi\|74198706 **Mass:** 57000 **Score:** 45 **Matches:** 1(0) **Sequences:** 1(0)
unnamed protein product [Mus musculus]
gi\|74203945 **Mass:** 56566 **Score:** 45 **Matches:** 1(0) **Sequences:** 1(0)
unnamed protein product [Mus musculus]
gi\|74212231 **Mass:** 57053 **Score:** 45 **Matches:** 1(0) **Sequences:** 1(0)
unnamed protein product [Mus musculus]
gi\|74219772 **Mass:** 57008 **Score:** 45 **Matches:** 1(0) **Sequences:** 1(0)
unnamed protein product [Mus musculus]
gi\|74220649 **Mass:** 57067 **Score:** 45 **Matches:** 1(0) **Sequences:** 1(0)
unnamed protein product [Mus musculus]
gi\|148702818 **Mass:** 58984 **Score:** 45 **Matches:** 1(0) **Sequences:** 1(0)
prolyl 4-hydroxylase, beta polypeptide, isoform CRA_a [Mus musculus]
gi\|148702819 **Mass:** 61492 **Score:** 45 **Matches:** 1(0) **Sequences:** 1(0)
prolyl 4-hydroxylase, beta polypeptide, isoform CRA_b [Mus musculus]

---

**29.** gi\|26390362 **Mass:** 39663 **Score:** 45 **Matches:** 1(0) **Sequences:** 1(0) **emPAI:** 0.08
unnamed protein product [Mus musculus]

| Query | Observed | Mr(expt) | Mr(calc) | Delta | Miss | Score | Expect | Rank | Unique | Peptide |
|---|---|---|---|---|---|---|---|---|---|---|
| 12452 | 581.0289 | 1160.0432 | 1159.5622 | 0.4810 | 0 | 45 | 0.068 | 1 | U | R.EGWTVQDVAR.V |

**Proteins matching the same set of peptides:**

gi\|148287013 **Mass:** 39693 **Score:** 45 **Matches:** 1(0) **Sequences:** 1(0)
paraneoplastic antigen Ma1 homolog [Mus musculus]

---

**Mascot:** http://www.matrixscience.com/

PFHR9

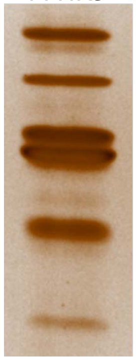

5

### {MATRIX SCIENCE} Mascot Search Results

```
User                     :
Email                    :
Search title             :
MS data file             : ZIEN-29Nov2017_5m.mgf
Database                 : NCBInr 20120419 (17893860 sequences; 6141683785 residues)
Taxonomy                 : Mus musculus (house mouse) (138831 sequences)
Timestamp                : 30 Nov 2017 at 21:31:00 GMT
Enzyme                   : Trypsin
Variable modifications   : Carbamidomethyl (C),Oxidation (M),Oxidation (P)
Mass values              : Monoisotopic
Protein Mass             : Unrestricted
Peptide Mass Tolerance   : ± 1.25 Da
Fragment Mass Tolerance: ± 1.001 Da
Max Missed Cleavages     : 1
Instrument type          : Default
Number of queries        : 15860
Protein hits             : gi|4159806  type II keratin subunit protein [Mus musculus]
                           gi|46485130 TPA_exp: keratin Kb40 [Mus musculus]
                           gi|12859782 unnamed protein product [Mus musculus]
                           gi|47523977 keratin, type II cytoskeletal 72 [Mus musculus]
                           gi|387397   epidermal keratin subunit I, partial [Mus musculus]
                           gi|2498741  RecName: Full=Prolyl 4-hydroxylase subunit alpha-2; Short=4-PH alpha-2; AltName: Full=Procollagen-proline,2-oxoglutarate-4-dioxygenase subunit alpha-2; Flags: Precursor
                           gi|16303309 type II keratin 5 [Mus musculus]
                           gi|1389682  plakoglobin, partial [Mus musculus]
                           gi|16716569 protease, serine, 1 precursor [Mus musculus]
                           gi|26337253 unnamed protein product [Mus musculus]
                           gi|12846468 unnamed protein product [Mus musculus]
                           gi|6755893  trypsin 4 precursor [Mus musculus]
                           gi|3687320  cartilage-associated protein (CASP) [Mus musculus]
                           gi|26336999 unnamed protein product [Mus musculus]
                           gi|53830774 lactase [Mus musculus]
                           gi|51450    heat shock protein [Mus musculus]
                           gi|52789    unnamed protein product [Mus musculus]
                           gi|21489937 lanC-like protein 1 [Mus musculus]
                           gi|12859178 unnamed protein product [Mus musculus]
                           gi|19353667 Pcmtd2 protein, partial [Mus musculus]
```

#### Select Summary Report

Format As | Select Summary (protein hits) | Help

Significance threshold p< 0.05 Max. number of hits AUTO

Standard scoring ○ MudPIT scoring ◉ Ions score or expect cut-off 43 Show sub-sets 0

Show pop-ups ◉ Suppress pop-ups ○ Require bold red ☐

Re-Search ◉ All queries ○ Unassigned ○ Below homology threshold ○ Below identity threshold

1. gi|4159806 **Mass:** 65183 **Score:** 178 **Matches:** 5(5) **Sequences:** 2(2) **emPAI:** 0.10
type II keratin subunit protein [Mus musculus]

| Query | Observed | Mr(expt) | Mr(calc) | Delta | Miss | Score | Expect | Rank | Unique | Peptide |
|---|---|---|---|---|---|---|---|---|---|---|
| 14220 | 692.5523 | 1383.0901 | 1383.7034 | -0.6133 | 1 | 49 | 0.022 | 1 | | K.SLNDKFASFIDK.V |
| 15069 | 738.1921 | 1474.3697 | 1474.8144 | -0.4446 | 1 | 81 | 1.3e-005 | 1 | U | R.FLEQQNKVLQTK.W 15077 15082 15083 |

2. gi|46485130 **Mass:** 85186 **Score:** 171 **Matches:** 6(6) **Sequences:** 2(2) **emPAI:** 0.08
TPA_exp: keratin Kb40 [Mus musculus]

| Query | Observed | Mr(expt) | Mr(calc) | Delta | Miss | Score | Expect | Rank | Unique | Peptide |
|---|---|---|---|---|---|---|---|---|---|---|
| 15082 | 738.6157 | 1475.2169 | 1475.7984 | -0.5815 | 1 | 85 | 2.5e-006 | 1 | U | R.FLEQQNKVLETK.W 15077 15083 15090 15094 |
| 15719 | 820.4082 | 1638.8018 | 1637.8525 | 0.9493 | 1 | 55 | 0.0075 | 1 | U | R.SLNNQFASFIDKVR.F |

**Proteins matching the same set of peptides:**
gi|111185567 **Mass:** 54730 **Score:** 171 **Matches:** 6(6) **Sequences:** 2(2)
Krt78 protein [Mus musculus]
gi|111185722 **Mass:** 56744 **Score:** 171 **Matches:** 6(6) **Sequences:** 2(2)
Krt78 protein [Mus musculus]
gi|119850791 **Mass:** 54739 **Score:** 171 **Matches:** 6(6) **Sequences:** 2(2)
Krt78 protein [Mus musculus]
gi|145580629 **Mass:** 112194 **Score:** 171 **Matches:** 6(6) **Sequences:** 2(2)
keratin Kb40 [Mus musculus]

---

**3.** gi|12859782 **Mass:** 65586 **Score:** 152 **Matches:** 5(5) **Sequences:** 2(2) **emPAI:** 0.10
unnamed protein product [Mus musculus]

| Query | Observed | Mr(expt) | Mr(calc) | Delta | Miss | Score | Expect | Rank | Unique | Peptide |
|---|---|---|---|---|---|---|---|---|---|---|
| 14220 | 692.5523 | 1383.0901 | 1383.7034 | -0.6133 | 1 | 49 | 0.022 | 1 | | K.SLNDKFASFIDK.V |
| 15082 | 738.6157 | 1475.2169 | 1474.7780 | 0.4389 | 0 | 81 | 7.5e-006 | 3 | U | R.FLEQQNQVLQTK.W 15069 15077 15083 |

**Proteins matching the same set of peptides:**
gi|126116585 **Mass:** 65565 **Score:** 152 **Matches:** 5(5) **Sequences:** 2(2)
keratin, type II cytoskeletal 1 [Mus musculus]

---

**4.** gi|47523977 **Mass:** 56715 **Score:** 145 **Matches:** 5(5) **Sequences:** 1(1) **emPAI:** 0.06
keratin, type II cytoskeletal 72 [Mus musculus]

| Query | Observed | Mr(expt) | Mr(calc) | Delta | Miss | Score | Expect | Rank | Unique | Peptide |
|---|---|---|---|---|---|---|---|---|---|---|
| 15082 | 738.6157 | 1475.2169 | 1475.7620 | -0.5451 | 0 | 80 | 7.9e-006 | 4 | U | R.FLEQQNQVLETK.W 15077 15083 15090 15094 |

---

**5.** gi|387397 **Mass:** 57807 **Score:** 122 **Matches:** 3(3) **Sequences:** 2(2) **emPAI:** 0.12
epidermal keratin subunit I, partial [Mus musculus]

| Query | Observed | Mr(expt) | Mr(calc) | Delta | Miss | Score | Expect | Rank | Unique | Peptide |
|---|---|---|---|---|---|---|---|---|---|---|
| 14203 | **691.6326** | **1381.2506** | **1380.6408** | **0.6097** | **0** | **81** | **1.7e-005** | **1** | **U** | **R.ALEESNYELEGK.I** |
| 14287 | **696.0388** | **1390.0630** | **1389.6736** | **0.3894** | **0** | **72** | **0.00013** | **1** | **U** | **K.QSLEASLAETEGR.Y** 14292 |

**Proteins matching the same set of peptides:**
gi|12852157 **Mass:** 58587 **Score:** 122 **Matches:** 3(3) **Sequences:** 2(2)
unnamed protein product [Mus musculus]
gi|26349141 **Mass:** 52653 **Score:** 122 **Matches:** 3(3) **Sequences:** 2(2)
unnamed protein product [Mus musculus]
gi|26349459 **Mass:** 49471 **Score:** 122 **Matches:** 3(3) **Sequences:** 2(2)
unnamed protein product [Mus musculus]
gi|112983636 **Mass:** 57007 **Score:** 122 **Matches:** 3(3) **Sequences:** 2(2)
keratin, type I cytoskeletal 10 [Mus musculus]
gi|116242600 **Mass:** 57735 **Score:** 122 **Matches:** 3(3) **Sequences:** 2(2)
RecName: Full=Keratin, type I cytoskeletal 10; AltName: Full=56 kDa cytokeratin; AltName: Full=Cytokeratin-10; Short=CK-10; AltName: Full=Keratin, type I cytoskeletal 59 kDa; AltName: Full=Keratin-1[illegible]

---

**6.** gi|2498741 **Mass:** 60964 **Score:** 74 **Matches:** 1(1) **Sequences:** 1(1) **emPAI:** 0.05
RecName: Full=Prolyl 4-hydroxylase subunit alpha-2; Short=4-PH alpha-2; AltName: Full=Procollagen-proline,2-oxoglutarate-4-dioxygenase subunit alpha-2; Flags: Precursor

| Query | Observed | Mr(expt) | Mr(calc) | Delta | Miss | Score | Expect | Rank | Unique | Peptide |
|---|---|---|---|---|---|---|---|---|---|---|
| 10969 | **533.6311** | **1065.2477** | **1065.5819** | **-0.3342** | **0** | **74** | **3.2e-005** | **1** | **U** | **K.TGVLTVASYR.V** |

**Proteins matching the same set of peptides:**
gi|74216495 **Mass:** 21055 **Score:** 74 **Matches:** 1(1) **Sequences:** 1(1)
unnamed protein product [Mus musculus]
gi|148701597 **Mass:** 57417 **Score:** 74 **Matches:** 1(1) **Sequences:** 1(1)
procollagen-proline, 2-oxoglutarate 4-dioxygenase (proline 4-hydroxylase), alpha II polypeptide, isoform CRA_b [Mus musculus]
gi|148701598 **Mass:** 21273 **Score:** 74 **Matches:** 1(1) **Sequences:** 1(1)
procollagen-proline, 2-oxoglutarate 4-dioxygenase (proline 4-hydroxylase), alpha II polypeptide, isoform CRA_c [Mus musculus]
gi|148701599 **Mass:** 61292 **Score:** 74 **Matches:** 1(1) **Sequences:** 1(1)
procollagen-proline, 2-oxoglutarate 4-dioxygenase (proline 4-hydroxylase), alpha II polypeptide, isoform CRA_d [Mus musculus]

gi|148701600 **Mass:** 66870 **Score:** 74 **Matches:** 1(1) **Sequences:** 1(1)
procollagen-proline, 2-oxoglutarate 4-dioxygenase (proline 4-hydroxylase), alpha II polypeptide, isoform CRA_e [Mus musculus]
gi|209862961 **Mass:** 60774 **Score:** 74 **Matches:** 1(1) **Sequences:** 1(1)
prolyl 4-hydroxylase subunit alpha-2 isoform 1 precursor [Mus musculus]
gi|226874876 **Mass:** 60977 **Score:** 74 **Matches:** 1(1) **Sequences:** 1(1)
prolyl 4-hydroxylase subunit alpha-2 isoform 2 precursor [Mus musculus]

---

**7.** gi|16303309 **Mass:** 61743 **Score:** 73 **Matches:** 2(2) **Sequences:** 2(2) **emPAI:** 0.11
type II keratin 5 [Mus musculus]

| Query | Observed | Mr(expt) | Mr(calc) | Delta | Miss | Score | Expect | Rank | Unique | Peptide |
|---|---|---|---|---|---|---|---|---|---|---|
| 13007 | 621.5424 | 1241.0702 | 1241.5564 | -0.4862 | 0 | 53 | 0.01 | 1 | U | R.TEAESWYQTK.Y |
| 14648 | 713.6679 | 1425.3211 | 1424.6680 | 0.6532 | 0 | 57 | 0.0037 | 1 | U | R.VDALMDEINFMK.M |

**Proteins matching the same set of peptides:**
gi|20911031 **Mass:** 61729 **Score:** 73 **Matches:** 2(2) **Sequences:** 2(2)
keratin, type II cytoskeletal 5 [Mus musculus]

---

**8.** gi|1389682 **Mass:** 68068 **Score:** 73 **Matches:** 1(1) **Sequences:** 1(1) **emPAI:** 0.05
plakoglobin, partial [Mus musculus]

| Query | Observed | Mr(expt) | Mr(calc) | Delta | Miss | Score | Expect | Rank | Unique | Peptide |
|---|---|---|---|---|---|---|---|---|---|---|
| 15139 | 742.2332 | 1482.4517 | 1482.7467 | -0.2949 | 0 | 73 | 8.8e-005 | 1 | U | R.NEGTATYAAAVLFR.I |

**Proteins matching the same set of peptides:**
gi|28395018 **Mass:** 81749 **Score:** 73 **Matches:** 1(1) **Sequences:** 1(1)
junction plakoglobin [Mus musculus]

---

**9.** gi|16716569 **Mass:** 26118 **Score:** 65 **Matches:** 2(2) **Sequences:** 1(1) **emPAI:** 0.13
protease, serine, 1 precursor [Mus musculus]

| Query | Observed | Mr(expt) | Mr(calc) | Delta | Miss | Score | Expect | Rank | Unique | Peptide |
|---|---|---|---|---|---|---|---|---|---|---|
| 15061 | 738.0139 | 2211.0197 | 2210.0967 | 0.9230 | 0 | 59 | 0.0026 | 1 | U | R.LGEHNINVLEGNEQFIDAAK.I 15060 |

---

**10.** gi|26337253 **Mass:** 53417 **Score:** 63 **Matches:** 1(1) **Sequences:** 1(1) **emPAI:** 0.06
unnamed protein product [Mus musculus]

| Query | Observed | Mr(expt) | Mr(calc) | Delta | Miss | Score | Expect | Rank | Unique | Peptide |
|---|---|---|---|---|---|---|---|---|---|---|
| 14805 | 722.0333 | 1442.0520 | 1441.7235 | 0.3285 | 0 | 63 | 0.0012 | 1 | U | K.GPAGVELNSMQPVK.E |

**Proteins matching the same set of peptides:**
gi|71060037 **Mass:** 53416 **Score:** 63 **Matches:** 1(1) **Sequences:** 1(1)
Slc2a3 [Mus musculus]
gi|261862282 **Mass:** 53444 **Score:** 63 **Matches:** 1(1) **Sequences:** 1(1)
solute carrier family 2, facilitated glucose transporter member 3 [Mus musculus]

---

**11.** gi|12846468 **Mass:** 40872 **Score:** 58 **Matches:** 1(1) **Sequences:** 1(1) **emPAI:** 0.08
unnamed protein product [Mus musculus]

| Query | Observed | Mr(expt) | Mr(calc) | Delta | Miss | Score | Expect | Rank | Unique | Peptide |
|---|---|---|---|---|---|---|---|---|---|---|
| 14387 | 700.5706 | 1399.1266 | 1398.7983 | 0.3283 | 0 | 58 | 0.0028 | 1 | U | K.LLAAFPNLANLSR.T |

**Proteins matching the same set of peptides:**
gi|26341012 **Mass:** 38517 **Score:** 58 **Matches:** 1(1) **Sequences:** 1(1)
unnamed protein product [Mus musculus]
gi|26343781 **Mass:** 33610 **Score:** 58 **Matches:** 1(1) **Sequences:** 1(1)
unnamed protein product [Mus musculus]
gi|26351471 **Mass:** 49011 **Score:** 58 **Matches:** 1(1) **Sequences:** 1(1)
unnamed protein product [Mus musculus]
gi|50510431 **Mass:** 45578 **Score:** 58 **Matches:** 1(1) **Sequences:** 1(1)
mKIAA0265 protein [Mus musculus]
gi|58037463 **Mass:** 48981 **Score:** 58 **Matches:** 1(1) **Sequences:** 1(1)

kelch domain-containing protein 10 [Mus musculus]
gi|62945268 **Mass:** 45351 **Score:** 58 **Matches:** 1(1) **Sequences:** 1(1)
kelch domain-containing protein 10 [Rattus norvegicus]
gi|148681799 **Mass:** 33511 **Score:** 58 **Matches:** 1(1) **Sequences:** 1(1)
RIKEN cDNA 2410127E18, isoform CRA_a [Mus musculus]
gi|148681802 **Mass:** 32113 **Score:** 58 **Matches:** 1(1) **Sequences:** 1(1)
RIKEN cDNA 2410127E18, isoform CRA_c [Mus musculus]

---

**12.** gi|6755893 **Mass:** 26257 **Score:** 57 **Matches:** 1(1) **Sequences:** 1(1) **emPAI:** 0.13
trypsin 4 precursor [Mus musculus]

| Query | Observed | Mr(expt) | Mr(calc) | Delta | Miss | Score | Expect | Rank | Unique | Peptide |
|---|---|---|---|---|---|---|---|---|---|---|
| 15071 | 738.3822 | 2212.1248 | 2211.0920 | 1.0328 | 0 | 57 | 0.0052 | 1 | U | R.LGEHNINVLEGNEQFVNSAK.I |

**Proteins matching the same set of peptides:**
gi|51010909 **Mass:** 26260 **Score:** 57 **Matches:** 1(1) **Sequences:** 1(1)
trypsin 5 precursor [Mus musculus]
gi|74203392 **Mass:** 27146 **Score:** 57 **Matches:** 1(1) **Sequences:** 1(1)
unnamed protein product [Mus musculus]

---

**13.** gi|3687320 **Mass:** 46137 **Score:** 57 **Matches:** 2(2) **Sequences:** 2(2) **emPAI:** 0.15
cartilage-associated protein (CASP) [Mus musculus]

| Query | Observed | Mr(expt) | Mr(calc) | Delta | Miss | Score | Expect | Rank | Unique | Peptide |
|---|---|---|---|---|---|---|---|---|---|---|
| 13636 | 662.2533 | 1322.4920 | 1322.6176 | -0.1256 | 0 | 47 | 0.045 | 1 | U | R.DELMPLESAYR.H |
| 15767 | 840.3280 | 1678.6414 | 1678.8488 | -0.2073 | 0 | 47 | 0.043 | 1 | U | R.TSISDMELALPDFLK.A |

**Proteins matching the same set of peptides:**
gi|12857227 **Mass:** 46121 **Score:** 57 **Matches:** 2(2) **Sequences:** 2(2)
unnamed protein product [Mus musculus]
gi|148677365 **Mass:** 46110 **Score:** 57 **Matches:** 2(2) **Sequences:** 2(2)
mCG7119 [Mus musculus]
gi|225543173 **Mass:** 46140 **Score:** 57 **Matches:** 2(2) **Sequences:** 2(2)
cartilage-associated protein precursor [Mus musculus]

---

**14.** gi|26336999 **Mass:** 60848 **Score:** 55 **Matches:** 1(1) **Sequences:** 1(1) **emPAI:** 0.05
unnamed protein product [Mus musculus]

| Query | Observed | Mr(expt) | Mr(calc) | Delta | Miss | Score | Expect | Rank | Unique | Peptide |
|---|---|---|---|---|---|---|---|---|---|---|
| 15096 | 739.4654 | 2215.3743 | 2215.1274 | 0.2470 | 1 | 55 | 0.006 | 1 | U | R.LTSTATKDPEGFVGHPVNAFK.L |

**Proteins matching the same set of peptides:**
gi|33859596 **Mass:** 60872 **Score:** 55 **Matches:** 1(1) **Sequences:** 1(1)
prolyl 4-hydroxylase subunit alpha-1 precursor [Mus musculus]
gi|74148153 **Mass:** 51722 **Score:** 55 **Matches:** 1(1) **Sequences:** 1(1)
unnamed protein product [Mus musculus]
gi|74224984 **Mass:** 60886 **Score:** 55 **Matches:** 1(1) **Sequences:** 1(1)
unnamed protein product [Mus musculus]
gi|74225936 **Mass:** 63769 **Score:** 55 **Matches:** 1(1) **Sequences:** 1(1)
unnamed protein product [Mus musculus]

---

**15.** gi|53830774 **Mass:** 34092 **Score:** 49 **Matches:** 1(1) **Sequences:** 1(1) **emPAI:** 0.10
lactase [Mus musculus]

| Query | Observed | Mr(expt) | Mr(calc) | Delta | Miss | Score | Expect | Rank | Unique | Peptide |
|---|---|---|---|---|---|---|---|---|---|---|
| 11285 | 547.7469 | 1093.4792 | 1094.5132 | -1.0339 | 0 | 49 | 0.014 | 1 | U | R.LPEFTESEK.K |

**Proteins matching the same set of peptides:**
gi|74192292 **Mass:** 139170 **Score:** 49 **Matches:** 1(1) **Sequences:** 1(1)
unnamed protein product [Mus musculus]
gi|124487297 **Mass:** 217665 **Score:** 49 **Matches:** 1(1) **Sequences:** 1(1)

lactase-phlorizin hydrolase preproprotein [Mus musculus]
gi|148707805 **Mass:** 217693 **Score:** 49 **Matches:** 1(1) **Sequences:** 1(1)
mCG128560 [Mus musculus]

---

**16.** gi|51450 **Mass:** 46560 **Score:** 48 **Matches:** 1(1) **Sequences:** 1(1) **emPAI:** 0.07
heat shock protein [Mus musculus]

| Query | Observed | Mr(expt) | Mr(calc) | Delta | Miss | Score | Expect | Rank | Unique | Peptide |
|---|---|---|---|---|---|---|---|---|---|---|
| 12831 | 613.0960 | 1224.1775 | 1223.6510 | 0.5265 | 0 | 48 | 0.012 | 1 | U | K.GVVEVTHDLQK.H |

**Proteins matching the same set of peptides:**
gi|200966 **Mass:** 45623 **Score:** 48 **Matches:** 1(1) **Sequences:** 1(1)
put. serine protease inhibitor [Mus musculus]
gi|26345418 **Mass:** 46481 **Score:** 48 **Matches:** 1(1) **Sequences:** 1(1)
unnamed protein product [Mus musculus]
gi|26348007 **Mass:** 46490 **Score:** 48 **Matches:** 1(1) **Sequences:** 1(1)
unnamed protein product [Mus musculus]
gi|74139266 **Mass:** 46490 **Score:** 48 **Matches:** 1(1) **Sequences:** 1(1)
unnamed protein product [Mus musculus]
gi|74191337 **Mass:** 46505 **Score:** 48 **Matches:** 1(1) **Sequences:** 1(1)
unnamed protein product [Mus musculus]
gi|74198254 **Mass:** 46500 **Score:** 48 **Matches:** 1(1) **Sequences:** 1(1)
unnamed protein product [Mus musculus]
gi|148684430 **Mass:** 44954 **Score:** 48 **Matches:** 1(1) **Sequences:** 1(1)
serine (or cysteine) peptidase inhibitor, clade H, member 1, isoform CRA_a [Mus musculus]
gi|161353502 **Mass:** 46504 **Score:** 48 **Matches:** 1(1) **Sequences:** 1(1)
serpin H1 precursor [Mus musculus]

---

**17.** gi|52789 **Mass:** 54415 **Score:** 45 **Matches:** 1(1) **Sequences:** 1(1) **emPAI:** 0.06
unnamed protein product [Mus musculus]

| Query | Observed | Mr(expt) | Mr(calc) | Delta | Miss | Score | Expect | Rank | Unique | Peptide |
|---|---|---|---|---|---|---|---|---|---|---|
| 14209 | 692.0989 | 1382.1832 | 1382.7194 | -0.5362 | 1 | 45 | 0.058 | 1 | U | K.SLNNKFASFIDK.V |

**Proteins matching the same set of peptides:**
gi|309215 **Mass:** 53210 **Score:** 45 **Matches:** 1(1) **Sequences:** 1(1)
EndoA' cytokeratin (5' end put.); putative [Mus musculus]
gi|511654 **Mass:** 54220 **Score:** 45 **Matches:** 1(1) **Sequences:** 1(1)
keratin type II [Mus musculus]
gi|74177777 **Mass:** 54546 **Score:** 45 **Matches:** 1(1) **Sequences:** 1(1)
unnamed protein product [Mus musculus]
gi|74219975 **Mass:** 54459 **Score:** 45 **Matches:** 1(1) **Sequences:** 1(1)
unnamed protein product [Mus musculus]
gi|76779293 **Mass:** 54514 **Score:** 45 **Matches:** 1(1) **Sequences:** 1(1)
Keratin 8 [Mus musculus]
gi|114145561 **Mass:** 54531 **Score:** 45 **Matches:** 1(1) **Sequences:** 1(1)
keratin, type II cytoskeletal 8 [Mus musculus]

---

**18.** gi|21489937 **Mass:** 45312 **Score:** 45 **Matches:** 1(1) **Sequences:** 1(1) **emPAI:** 0.07
lanC-like protein 1 [Mus musculus]

| Query | Observed | Mr(expt) | Mr(calc) | Delta | Miss | Score | Expect | Rank | Unique | Peptide |
|---|---|---|---|---|---|---|---|---|---|---|
| 10769 | 524.0907 | 1046.1668 | 1045.5226 | 0.6442 | 0 | 45 | 0.029 | 1 | U | R.ELLQQMER.G |

**Proteins matching the same set of peptides:**
gi|148667828 **Mass:** 45346 **Score:** 45 **Matches:** 1(1) **Sequences:** 1(1)
LanC (bacterial lantibiotic synthetase component C)-like 1, isoform CRA_a [Mus musculus]
gi|148667829 **Mass:** 47990 **Score:** 45 **Matches:** 1(1) **Sequences:** 1(1)
LanC (bacterial lantibiotic synthetase component C)-like 1, isoform CRA_b [Mus musculus]

---

**19.** gi|12859178 **Mass:** 26301 **Score:** 45 **Matches:** 1(0) **Sequences:** 1(0) **emPAI:** 0.13

unnamed protein product [Mus musculus]

| Query | Observed | Mr(expt) | Mr(calc) | Delta | Miss | Score | Expect | Rank | Unique | Peptide |
|---|---|---|---|---|---|---|---|---|---|---|
| 13273 | 639.2183 | 1914.6330 | 1913.8795 | 0.7534 | 1 | 45 | 0.072 | 1 | U | K.DAEEWFFTKTEELNR.E |

**Proteins matching the same set of peptides:**

gi\|38503465 **Mass:** 50146 **Score:** 45 **Matches:** 1(0) **Sequences:** 1(0)
keratin 17n [Mus musculus]

gi\|85726518 **Mass:** 49857 **Score:** 45 **Matches:** 1(0) **Sequences:** 1(0)
Krt42 protein [Mus musculus]

gi\|154090941 **Mass:** 50102 **Score:** 45 **Matches:** 1(0) **Sequences:** 1(0)
keratin, type I cytoskeletal 42 [Mus musculus]

gi\|156139032 **Mass:** 50162 **Score:** 45 **Matches:** 1(0) **Sequences:** 1(0)
Keratin 42 [Mus musculus]

---

**20.** gi\|19353667 **Mass:** 19691 **Score:** 43 **Matches:** 2(1) **Sequences:** 2(1) **emPAI:** 0.17
Pcmtd2 protein, partial [Mus musculus]

| Query | Observed | Mr(expt) | Mr(calc) | Delta | Miss | Score | Expect | Rank | Unique | Peptide |
|---|---|---|---|---|---|---|---|---|---|---|
| 10138 | 489.0460 | 976.0774 | 975.4662 | 0.6112 | 0 | 43 | 0.045 | 1 | U | R.TGPSAWETK.K |
| 11516 | 556.1887 | 1110.3629 | 1110.5631 | -0.2002 | 0 | 43 | 0.1 | 1 | U | R.METIVFLDK.E |

**Proteins matching the same set of peptides:**

gi\|23956398 **Mass:** 40730 **Score:** 43 **Matches:** 2(1) **Sequences:** 2(1)
protein-L-isoaspartate O-methyltransferase domain-containing protein 2 [Mus musculus]

gi\|26348291 **Mass:** 21087 **Score:** 43 **Matches:** 2(1) **Sequences:** 2(1)
unnamed protein product [Mus musculus]

---

**Mascot:** http://www.matrixscience.com/

### Mascot Search Results

D17

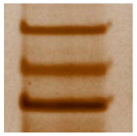

A

User :
Email :
Search title :
MS data file : ZIEN-29Nov2017_flyA.mgf
Database : NCBInr 20120419 (17893860 sequences; 6141683785 residues)
Taxonomy : Drosophila (fruit flies) (222527 sequences)
Timestamp : 30 Nov 2017 at 18:31:47 GMT
Enzyme : Trypsin
Variable modifications : Carbamidomethyl (C),Oxidation (M)
Mass values : Monoisotopic
Protein Mass : Unrestricted
Peptide Mass Tolerance : ± 1.25 Da
Fragment Mass Tolerance: ± 1.001 Da
Max Missed Cleavages : 1
Instrument type : ESI-TRAP
Number of queries : 16095
Protein hits : gi|17737967 heat shock protein cognate 4, isoform A [Drosophila melanogaster]
gi|157658 heat shock protein cognate 72 [Drosophila melanogaster]
gi|157667 heat shock protein cognate 71 [Drosophila melanogaster]
gi|157678 heat shock cognate 70 protein (partial) (at locus 88E), partial [Drosophila melanogaster]
gi|6118254 DEAD-box protein abstrakt [Drosophila melanogaster]
gi|195449894 GK22765 [Drosophila willistoni]
gi|24651407 prolyl-4-hydroxylase-alpha EFB [Drosophila melanogaster]
gi|195341536 GM12882 [Drosophila sechellia]
gi|24644396 CG2082, isoform C [Drosophila melanogaster]
gi|157891 myosin heavy chain [Drosophila melanogaster]
gi|195161163 GL25329 [Drosophila persimilis]
gi|194864176 GG23181 [Drosophila erecta]
gi|194769542 GF19055 [Drosophila ananassae]
gi|194766886 GF22554 [Drosophila ananassae]

#### Select Summary Report

Format As | Select Summary (protein hits) ▼ | Help

Significance threshold p< 0.05 Max. number of hits AUTO

Standard scoring ○ MudPIT scoring ◉ Ions score or expect cut-off 44 Show sub-sets 0

Show pop-ups ◉ Suppress pop-ups ○ Require bold red ☐

Re-Search ◉ All queries ○ Unassigned ○ Below homology threshold ○ Below identity threshold

**1.** gi|17737967 **Mass:** 71087 **Score:** 168 **Matches:** 7(7) **Sequences:** 6(6) **emPAI:** 0.31

heat shock protein cognate 4, isoform A [Drosophila melanogaster]

| Query | Observed | Mr(expt) | Mr(calc) | Delta | Miss | Score | Expect | Rank | Unique | Peptide |
|---|---|---|---|---|---|---|---|---|---|---|
| 12119 | 587.4327 | 1172.8509 | 1172.5826 | 0.2684 | 0 | 51 | 0.011 | 1 | U | K.WLDANQLADK.E |
| 12465 | 600.5588 | 1199.1031 | 1198.6670 | 0.4362 | 0 | 57 | 0.0044 | 1 | U | K.DAGTIAGLNVLR.I 12466 |
| 12751 | 613.9276 | 1225.8405 | 1225.5285 | 0.3120 | 0 | 49 | 0.032 | 1 | | K.FDDAAVQSDMK.H |
| 13970 | 678.0540 | 1354.0934 | 1353.6235 | 0.4699 | 1 | 53 | 0.011 | 1 | | R.KFDDAAVQSDMK.H |
| 15110 | 727.5170 | 1453.0195 | 1451.8096 | 1.2099 | 0 | 76 | 3e-005 | 1 | U | K.SVIHDIVLVGGSTR.I |
| 15964 | 839.5435 | 1677.0724 | 1676.8192 | 0.2532 | 0 | 92 | 1.2e-006 | 1 | U | K.NQVAMNPTQTIFDAK.R |

**Proteins matching the same set of peptides:**

gi|19527633 **Mass:** 71060 **Score:** 168 **Matches:** 7(7) **Sequences:** 6(6)
RH04426p [Drosophila melanogaster]

gi|194900946 **Mass:** 71001 **Score:** 168 **Matches:** 7(7) **Sequences:** 6(6)
GG16900 [Drosophila erecta]

gi|195328813 **Mass:** 70971 **Score:** 168 **Matches:** 7(7) **Sequences:** 6(6)
GM24208 [Drosophila sechellia]

gi|195570732 **Mass:** 70916 **Score:** 168 **Matches:** 7(7) **Sequences:** 6(6)
Hsc70-4 [Drosophila simulans]

---

**2.** gi|157658 **Mass:** 72190 **Score:** 108 **Matches:** 5(5) **Sequences:** 5(5) **emPAI:** 0.25

heat shock protein cognate 72 [Drosophila melanogaster]

| Query | Observed | Mr(expt) | Mr(calc) | Delta | Miss | Score | Expect | Rank | Unique | Peptide |
|---|---|---|---|---|---|---|---|---|---|---|
| 9446 | 478.0703 | 954.1261 | 953.5043 | 0.6218 | 0 | 62 | 0.00066 | 1 | U | R.ALSGSHQVR.I |
| 11066 | 552.6559 | 1103.2972 | 1103.5710 | -0.2738 | 0 | 56 | 0.0035 | 1 | U | K.LESAIDESIK.W |
| 13643 | 659.0761 | 1316.1375 | 1315.6295 | 0.5080 | 0 | 55 | 0.006 | 1 | U | R.NELESYAYSLK.N |
| 15556 | 754.5173 | 1507.0201 | 1505.7800 | 1.2402 | 0 | 51 | 0.015 | 1 | U | K.VFAPEEISAMVLGK.M |
| 15667 | 764.1249 | 1526.2352 | 1525.7413 | 0.4939 | 0 | 48 | 0.028 | 1 | U | R.ITPSYVAFTADGER.L |

**Proteins matching the same set of peptides:**

gi|24641402 **Mass:** 72216 **Score:** 108 **Matches:** 5(5) **Sequences:** 5(5)
heat shock protein cognate 3, isoform A [Drosophila melanogaster]

gi|28557577 **Mass:** 72069 **Score:** 108 **Matches:** 5(5) **Sequences:** 5(5)
RH21402p [Drosophila melanogaster]

gi|125981509 **Mass:** 72261 **Score:** 108 **Matches:** 5(5) **Sequences:** 5(5)
GA17988 [Drosophila pseudoobscura pseudoobscura]

gi|194762782 **Mass:** 72278 **Score:** 108 **Matches:** 5(5) **Sequences:** 5(5)
GF20248 [Drosophila ananassae]
gi|195165202 **Mass:** 72272 **Score:** 108 **Matches:** 5(5) **Sequences:** 5(5)
GL20354 [Drosophila persimilis]
gi|195457222 **Mass:** 72381 **Score:** 108 **Matches:** 5(5) **Sequences:** 5(5)
GK18328 [Drosophila willistoni]
gi|195480743 **Mass:** 72226 **Score:** 108 **Matches:** 5(5) **Sequences:** 5(5)
Hsc70-3 [Drosophila yakuba]

---

**3.** gi|157667 **Mass:** 74162 **Score:** 100 **Matches:** 6(5) **Sequences:** 5(4) **emPAI:** 0.24
heat shock protein cognate 71 [Drosophila melanogaster]

| Query | Observed | Mr(expt) | Mr(calc) | Delta | Miss | Score | Expect | Rank | Unique | Peptide |
|---|---|---|---|---|---|---|---|---|---|---|
| 9395 | **953.4493** | **952.4420** | **952.4865** | **-0.0445** | **0** | **51** | **0.013** | **1** | **U** | **K.DITNLSYK.V** |
| 9494 | **958.5143** | **957.5070** | **957.4879** | **0.0191** | **0** | **45** | **0.064** | **1** | **U** | **K.VIENAEGAR.T** |
| 11337 | **563.4215** | **1124.8284** | **1124.5349** | **0.2935** | **1** | **49** | **0.015** | **1** | **U** | **K.KAEEYATADK.Q** |
| 12759 | **614.1332** | **1226.2518** | **1225.6779** | **0.5739** | **0** | **66** | **0.00049** | **1** | **U** | **K.DAGQIAGLNVLR.V** 12742 |
| 13805 | **668.6088** | **1335.2030** | **1334.6718** | **0.5312** | **0** | **45** | **0.035** | **1** | **U** | **K.ETAEAYLNTPVK.N** |

**Proteins matching the same set of peptides:**
gi|24653595 **Mass:** 74020 **Score:** 100 **Matches:** 6(5) **Sequences:** 5(4)
heat shock protein cognate 5 [Drosophila melanogaster]
gi|194883130 **Mass:** 74029 **Score:** 100 **Matches:** 6(5) **Sequences:** 5(4)
GG22433 [Drosophila erecta]
gi|195334294 **Mass:** 74060 **Score:** 100 **Matches:** 6(5) **Sequences:** 5(4)
GM20220 [Drosophila sechellia]
gi|195485915 **Mass:** 74021 **Score:** 100 **Matches:** 6(5) **Sequences:** 5(4)
GE12323 [Drosophila yakuba]
gi|195583328 **Mass:** 74074 **Score:** 100 **Matches:** 6(5) **Sequences:** 5(4)
GD25692 [Drosophila simulans]
gi|194756900 **Mass:** 73926 **Score:** 100 **Matches:** 6(5) **Sequences:** 5(4)
GF11360 [Drosophila ananassae]

---

**4.** gi|157678 **Mass:** 11411 **Score:** 97 **Matches:** 3(3) **Sequences:** 3(3) **emPAI:** 1.20
heat shock cognate 70 protein (partial) (at locus 88E), partial [Drosophila melanogaster]

| Query | Observed | Mr(expt) | Mr(calc) | Delta | Miss | Score | Expect | Rank | Unique | Peptide |
|---|---|---|---|---|---|---|---|---|---|---|
| 12751 | 613.9276 | 1225.8405 | 1225.5285 | 0.3120 | 0 | 49 | 0.032 | 1 | | K.FDDAAVQSDMK.H |
| 13970 | 678.0540 | 1354.0934 | 1353.6235 | 0.4699 | 1 | 53 | 0.011 | 1 | | R.KFDDAAVQSDMK.H |
| 15964 | 839.5435 | 1677.0724 | 1677.8032 | -0.7308 | 0 | 79 | 2.5e-005 | 2 | U | K.NEVAMNPTQTIFDAK.R |

---

**5.** gi|6118254 **Mass:** 68888 **Score:** 60 **Matches:** 1(1) **Sequences:** 1(1) **emPAI:** 0.05
DEAD-box protein abstrakt [Drosophila melanogaster]

| Query | Observed | Mr(expt) | Mr(calc) | Delta | Miss | Score | Expect | Rank | Unique | Peptide |
|---|---|---|---|---|---|---|---|---|---|---|
| 14629 | 704.5223 | 1407.0300 | 1406.7769 | 0.2531 | 0 | 60 | 0.002 | 1 | U | R.ILVEGETPSPPIR.S |

**Proteins matching the same set of peptides:**
gi|17977678 **Mass:** 69444 **Score:** 60 **Matches:** 1(1) **Sequences:** 1(1)
abstrakt [Drosophila melanogaster]

---

**6.** gi|195449894 **Mass:** 21978 **Score:** 57 **Matches:** 1(1) **Sequences:** 1(1) **emPAI:** 0.15
GK22765 [Drosophila willistoni]

| Query | Observed | Mr(expt) | Mr(calc) | Delta | Miss | Score | Expect | Rank | Unique | Peptide |
|---|---|---|---|---|---|---|---|---|---|---|
| 9376 | 475.9925 | 949.9705 | 950.4127 | -0.4422 | 0 | 57 | 0.0025 | 1 | U | M.TTANACDQK.G |

---

**7.** gi|24651407 **Mass:** 63075 **Score:** 53 **Matches:** 1(1) **Sequences:** 1(1) **emPAI:** 0.05
prolyl-4-hydroxylase-alpha EFB [Drosophila melanogaster]

| Query | Observed | Mr(expt) | Mr(calc) | Delta | Miss | Score | Expect | Rank | Unique | Peptide |
|---|---|---|---|---|---|---|---|---|---|---|
| 10856 | 545.6321 | 1089.2496 | 1089.5567 | -0.3071 | 0 | 53 | 0.012 | 1 | U | R.AFEGLNLGNR.I |

**Proteins matching the same set of peptides:**
gi|125772807 **Mass:** 62641 **Score:** 53 **Matches:** 1(1) **Sequences:** 1(1)
GA15946 [Drosophila pseudoobscura pseudoobscura]
gi|194765194 **Mass:** 62567 **Score:** 53 **Matches:** 1(1) **Sequences:** 1(1)
GF22904 [Drosophila ananassae]
gi|194905436 **Mass:** 62737 **Score:** 53 **Matches:** 1(1) **Sequences:** 1(1)
GG11753 [Drosophila erecta]
gi|195110919 **Mass:** 55326 **Score:** 53 **Matches:** 1(1) **Sequences:** 1(1)
GI24860 [Drosophila mojavensis]
gi|195159323 **Mass:** 55783 **Score:** 53 **Matches:** 1(1) **Sequences:** 1(1)
GL13463 [Drosophila persimilis]
gi|195391754 **Mass:** 55603 **Score:** 53 **Matches:** 1(1) **Sequences:** 1(1)
GJ24502 [Drosophila virilis]
gi|195452726 **Mass:** 62836 **Score:** 53 **Matches:** 1(1) **Sequences:** 1(1)
GK14136 [Drosophila willistoni]
gi|195505190 **Mass:** 55776 **Score:** 53 **Matches:** 1(1) **Sequences:** 1(1)
GE10881 [Drosophila yakuba]
gi|195575089 **Mass:** 62991 **Score:** 53 **Matches:** 1(1) **Sequences:** 1(1)
GD21521 [Drosophila simulans]

**8.** gi|195341536 **Mass:** 63009 **Score:** 53 **Matches:** 1(1) **Sequences:** 1(1) **emPAI:** 0.05
GM12882 [Drosophila sechellia]

| Query | Observed | Mr(expt) | Mr(calc) | Delta | Miss | Score | Expect | Rank | Unique | Peptide |
|---|---|---|---|---|---|---|---|---|---|---|
| 10856 | 545.6321 | 1089.2496 | 1089.5567 | -0.3071 | 0 | 53 | 0.012 | 1 | U | R.AFEGINLGNR.I |

**9.** gi|24644396 **Mass:** 39679 **Score:** 48 **Matches:** 1(1) **Sequences:** 1(1) **emPAI:** 0.08
CG2082, isoform C [Drosophila melanogaster]

| Query | Observed | Mr(expt) | Mr(calc) | Delta | Miss | Score | Expect | Rank | Unique | Peptide |
|---|---|---|---|---|---|---|---|---|---|---|
| 12547 | **604.5787** | **1810.7142** | **1810.8558** | **-0.1416** | **1** | **48** | **0.036** | **1** | **U** | **R.GRAFSTASSGSLEGANGSR.R** |

**Proteins matching the same set of peptides:**

gi|24644398 **Mass:** 39652 **Score:** 48 **Matches:** 1(1) **Sequences:** 1(1)
CG2082, isoform A [Drosophila melanogaster]
gi|62484286 **Mass:** 40038 **Score:** 48 **Matches:** 1(1) **Sequences:** 1(1)
CG2082, isoform B [Drosophila melanogaster]
gi|194745478 **Mass:** 40170 **Score:** 48 **Matches:** 1(1) **Sequences:** 1(1)
GF18647 [Drosophila ananassae]
gi|194898877 **Mass:** 39987 **Score:** 48 **Matches:** 1(1) **Sequences:** 1(1)
GG10854 [Drosophila erecta]
gi|195343799 **Mass:** 41940 **Score:** 48 **Matches:** 1(1) **Sequences:** 1(1)
GM10591 [Drosophila sechellia]
gi|262331602 **Mass:** 19636 **Score:** 48 **Matches:** 1(1) **Sequences:** 1(1)
AT13994p [Drosophila melanogaster]
gi|281360117 **Mass:** 40012 **Score:** 48 **Matches:** 1(1) **Sequences:** 1(1)
CG2082, isoform G [Drosophila melanogaster]
gi|281360119 **Mass:** 39979 **Score:** 48 **Matches:** 1(1) **Sequences:** 1(1)
CG2082, isoform H [Drosophila melanogaster]
gi|327180770 **Mass:** 41129 **Score:** 48 **Matches:** 1(1) **Sequences:** 1(1)
RH63159p [Drosophila melanogaster]
gi|383292515 **Mass:** 36950 **Score:** 48 **Matches:** 1(1) **Sequences:** 1(1)
CG2082, isoform O [Drosophila melanogaster]

**10.** gi|157891 **Score:** 48 **Matches:** 1(1) **Sequences:** 1(1) **emPAI:** 0.01
myosin heavy chain [Drosophila melanogaster]

| Query | Observed | Mr(expt) | Mr(calc) | Delta | Miss | Score | Expect | Rank | Unique | Peptide |
|---|---|---|---|---|---|---|---|---|---|---|
| 9376 | 475.9925 | 949.9705 | 950.4127 | -0.4422 | 0 | 48 | 0.019 | 2 | U | K.MQGETNQK.T |

**Proteins matching the same set of peptides:**

| | | | |
|---|---|---|---|
| gi\|157892 | Score: 48 | Matches: 1(1) | Sequences: 1(1) |
| gi\|2546936 | Score: 48 | Matches: 1(1) | Sequences: 1(1) |
| gi\|2546937 | Score: 48 | Matches: 1(1) | Sequences: 1(1) |
| gi\|2546938 | Score: 48 | Matches: 1(1) | Sequences: 1(1) |
| gi\|2546939 | Score: 48 | Matches: 1(1) | Sequences: 1(1) |
| gi\|24584692 | Score: 48 | Matches: 1(1) | Sequences: 1(1) |
| gi\|24584694 | Score: 48 | Matches: 1(1) | Sequences: 1(1) |
| gi\|24584696 | Score: 48 | Matches: 1(1) | Sequences: 1(1) |
| gi\|24584700 | Score: 48 | Matches: 1(1) | Sequences: 1(1) |
| gi\|24584702 | Score: 48 | Matches: 1(1) | Sequences: 1(1) |
| gi\|24584704 | Score: 48 | Matches: 1(1) | Sequences: 1(1) |
| gi\|24584706 | Score: 48 | Matches: 1(1) | Sequences: 1(1) |
| gi\|24584710 | Score: 48 | Matches: 1(1) | Sequences: 1(1) |
| gi\|24584712 | Score: 48 | Matches: 1(1) | Sequences: 1(1) |
| gi\|24584714 | Score: 48 | Matches: 1(1) | Sequences: 1(1) |
| gi\|24584716 | Score: 48 | Matches: 1(1) | Sequences: 1(1) |
| gi\|28574239 | Score: 48 | Matches: 1(1) | Sequences: 1(1) |
| gi\|110825729 | Score: 48 | Matches: 1(1) | Sequences: 1(1) |
| gi\|194884434 | Score: 48 | Matches: 1(1) | Sequences: 1(1) |
| gi\|195344578 | Score: 48 | Matches: 1(1) | Sequences: 1(1) |
| gi\|195436762 | Score: 48 | Matches: 1(1) | Sequences: 1(1) |
| gi\|195483992 | Score: 48 | Matches: 1(1) | Sequences: 1(1) |
| gi\|198474044 | Score: 48 | Matches: 1(1) | Sequences: 1(1) |
| gi\|219990777 | Score: 48 | Matches: 1(1) | Sequences: 1(1) |
| gi\|281365095 | Score: 48 | Matches: 1(1) | Sequences: 1(1) |
| gi\|281365099 | Score: 48 | Matches: 1(1) | Sequences: 1(1) |
| gi\|383291520 | Score: 48 | Matches: 1(1) | Sequences: 1(1) |
| gi\|383291521 | Score: 48 | Matches: 1(1) | Sequences: 1(1) |
| gi\|383291522 | Score: 48 | Matches: 1(1) | Sequences: 1(1) |
| gi\|383291523 | Score: 48 | Matches: 1(1) | Sequences: 1(1) |
| gi\|383291524 | Score: 48 | Matches: 1(1) | Sequences: 1(1) |

---

**11.** gi|195161163 **Mass:** 58891 **Score:** 47 **Matches:** 1(1) **Sequences:** 1(1) **emPAI:** 0.06

GL25329 [Drosophila persimilis]

| Query | Observed | Mr(expt) | Mr(calc) | Delta | Miss | Score | Expect | Rank | Unique | Peptide |
|---|---|---|---|---|---|---|---|---|---|---|
| 12491 | **601.5865** | **1201.1585** | **1200.5232** | **0.6353** | **0** | **47** | **0.05** | **1** | **U** | **R.QDNGGGGGVGGGGGR.F** |

---

**12.** gi|194864176 **Score:** 46 **Matches:** 1(1) **Sequences:** 1(1) **emPAI:** 0.02

GG23181 [Drosophila erecta]

| Query | Observed | Mr(expt) | Mr(calc) | Delta | Miss | Score | Expect | Rank | Unique | Peptide |
|---|---|---|---|---|---|---|---|---|---|---|
| 12466 | 600.5885 | 1199.1624 | 1198.6670 | 0.4955 | 0 | 46 | 0.058 | 2 | U | R.ETNNVIAIGLR.D |

**13.** gi|194769542 **Mass:** 194372 **Score:** 46 **Matches:** 1(1) **Sequences:** 1(1) **emPAI:** 0.02
GF19055 [Drosophila ananassae]

| Query | Observed | Mr(expt) | Mr(calc) | Delta | Miss | Score | Expect | Rank | Unique | Peptide |
|---|---|---|---|---|---|---|---|---|---|---|
| 9300 | 472.3273 | 942.6400 | 942.5498 | 0.0902 | 0 | 46 | 0.046 | 1 | U | K.VIEISVQR.G |

**Proteins matching the same set of peptides:**

gi|194892726 **Mass:** 191336 **Score:** 46 **Matches:** 1(1) **Sequences:** 1(1)
GG19193 [Drosophila erecta]

gi|195132717 **Mass:** 70104 **Score:** 46 **Matches:** 1(1) **Sequences:** 1(1)
GI21732 [Drosophila mojavensis]

gi|195392652 **Mass:** 186045 **Score:** 46 **Matches:** 1(1) **Sequences:** 1(1)
GJ19114 [Drosophila virilis]

gi|195432565 **Mass:** 215970 **Score:** 46 **Matches:** 1(1) **Sequences:** 1(1)
GK19785 [Drosophila willistoni]

gi|195481506 **Mass:** 198954 **Score:** 46 **Matches:** 1(1) **Sequences:** 1(1)
GE17755 [Drosophila yakuba]

gi|198469723 **Mass:** 196607 **Score:** 46 **Matches:** 1(1) **Sequences:** 1(1)
GA23340 [Drosophila pseudoobscura pseudoobscura]

**14.** gi|194766886 **Score:** 45 **Matches:** 1(1) **Sequences:** 1(1) **emPAI:** 0.01
GF22554 [Drosophila ananassae]

| Query | Observed | Mr(expt) | Mr(calc) | Delta | Miss | Score | Expect | Rank | Unique | Peptide |
|---|---|---|---|---|---|---|---|---|---|---|
| 9376 | 475.9925 | 949.9705 | 950.4742 | -0.5037 | 1 | 45 | 0.034 | 3 | U | K.MKSAELEK.E |

**Mascot:** http://www.matrixscience.com/

### Mascot Search Results

D17

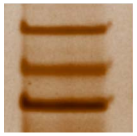

B

User :
Email :
Search title :
MS data file : ZIEN-29Nov2017_flyB.mgf
Database : NCBInr 20120419 (17893860 sequences; 6141683785 residues)
Taxonomy : Drosophila (fruit flies) (222527 sequences)
Timestamp : 30 Nov 2017 at 18:51:10 GMT
Enzyme : Trypsin
Variable modifications : Carbamidomethyl (C),Oxidation (M)
Mass values : Monoisotopic
Protein Mass : Unrestricted
Peptide Mass Tolerance : ± 1.25 Da
Fragment Mass Tolerance: ± 1.001 Da
Max Missed Cleavages : 1
Instrument type : Default
Number of queries : 15798
Protein hits : gi|17647799 protein disulfide isomerase, isoform A [Drosophila melanogaster]
gi|24664683 CG16979 [Drosophila melanogaster]
gi|195451942 GK13309 [Drosophila willistoni]
gi|24644396 CG2082, isoform C [Drosophila melanogaster]
gi|198476294 GA25398 [Drosophila pseudoobscura pseudoobscura]
gi|24640588 CG1440, isoform A [Drosophila melanogaster]
gi|24651407 prolyl-4-hydroxylase-alpha EFB [Drosophila melanogaster]
gi|195161605 GL26392 [Drosophila persimilis]
gi|195450712 GK13603 [Drosophila willistoni]

#### Select Summary Report

Format As | Select Summary (protein hits) | Help

Significance threshold p< 0.05 Max. number of hits AUTO

Standard scoring MudPIT scoring Ions score or expect cut-off 44 Show sub-sets 0

Show pop-ups Suppress pop-ups Require bold red

Re-Search | All queries | Unassigned | Below homology threshold | Below identity threshold

1. gi|17647799 **Mass:** 55746 **Score:** 120 **Matches:** 3(3) **Sequences:** 3(3) **emPAI:** 0.19
protein disulfide isomerase, isoform A [Drosophila melanogaster]

**Query Observed Mr(expt) Mr(calc) Delta Miss Score Expect Rank Unique Peptide**

| | | | | | | | | | | |
|---|---|---|---|---|---|---|---|---|---|---|
| 12211 | 608.3029 | 1214.5912 | 1214.6659 | -0.0747 | 0 | 55 | 0.0097 | 1 | U | R.QAADIIAWVTK.K |
| 13260 | 669.5514 | 1337.0882 | 1336.6180 | 0.4702 | 0 | 86 | 2.5e-006 | 1 | U | K.MDSTANELESIK.I |
| 15162 | 754.9669 | 1507.9192 | 1506.7202 | 1.1990 | 0 | 62 | 0.00091 | 1 | U | K.SVFEGELNEENLK.K |

**Proteins matching the same set of peptides:**

gi|195327719 **Mass:** 55772 **Score:** 120 **Matches:** 3(3) **Sequences:** 3(3)
GM24501 [Drosophila sechellia]

---

**2.** gi|24664683 **Mass:** 68207 **Score:** 66 **Matches:** 1(1) **Sequences:** 1(1) **emPAI:** 0.05
CG16979 [Drosophila melanogaster]

| Query | Observed | Mr(expt) | Mr(calc) | Delta | Miss | Score | Expect | Rank | Unique | Peptide |
|---|---|---|---|---|---|---|---|---|---|---|
| 15088 | 751.1436 | 1500.2727 | 1500.7937 | -0.5210 | 0 | 66 | 0.00048 | 1 | U | R.VADGNLVFNVPQTK.I |

**Proteins matching the same set of peptides:**

gi|194872892 **Mass:** 68503 **Score:** 66 **Matches:** 1(1) **Sequences:** 1(1)
GG15911 [Drosophila erecta]

gi|195327809 **Mass:** 68273 **Score:** 66 **Matches:** 1(1) **Sequences:** 1(1)
GM25541 [Drosophila sechellia]

gi|195477958 **Mass:** 68443 **Score:** 66 **Matches:** 1(1) **Sequences:** 1(1)
GE23132 [Drosophila yakuba]

gi|195590489 **Mass:** 68190 **Score:** 66 **Matches:** 1(1) **Sequences:** 1(1)
GD14556 [Drosophila simulans]

---

**3.** gi|195451942 **Mass:** 253843 **Score:** 53 **Matches:** 1(1) **Sequences:** 1(1) **emPAI:** 0.01
GK13309 [Drosophila willistoni]

| Query | Observed | Mr(expt) | Mr(calc) | Delta | Miss | Score | Expect | Rank | Unique | Peptide |
|---|---|---|---|---|---|---|---|---|---|---|
| 15598 | 801.8900 | 1601.7655 | 1600.7917 | 0.9737 | 0 | 53 | 0.012 | 1 | U | R.NGGLGAAGIVASNSGSNR.V |

---

**4.** gi|24644396 **Mass:** 39679 **Score:** 52 **Matches:** 1(1) **Sequences:** 1(1) **emPAI:** 0.08
CG2082, isoform C [Drosophila melanogaster]

| Query | Observed | Mr(expt) | Mr(calc) | Delta | Miss | Score | Expect | Rank | Unique | Peptide |
|---|---|---|---|---|---|---|---|---|---|---|
| 12135 | 604.6827 | 1811.0264 | 1810.8558 | 0.1706 | 1 | 52 | 0.016 | 1 | U | R.GRAFSTASSGSLEGANGSR.R |

**Proteins matching the same set of peptides:**

gi|24644398 **Mass:** 39652 **Score:** 52 **Matches:** 1(1) **Sequences:** 1(1)
CG2082, isoform A [Drosophila melanogaster]

gi|62484286 **Mass:** 40038 **Score:** 52 **Matches:** 1(1) **Sequences:** 1(1)

CG2082, isoform B [Drosophila melanogaster]
gi|194745478 **Mass:** 40170 **Score:** 52 **Matches:** 1(1) **Sequences:** 1(1)
GF18647 [Drosophila ananassae]
gi|194898877 **Mass:** 39987 **Score:** 52 **Matches:** 1(1) **Sequences:** 1(1)
GG10854 [Drosophila erecta]
gi|195343799 **Mass:** 41940 **Score:** 52 **Matches:** 1(1) **Sequences:** 1(1)
GM10591 [Drosophila sechellia]
gi|262331602 **Mass:** 19636 **Score:** 52 **Matches:** 1(1) **Sequences:** 1(1)
AT13994p [Drosophila melanogaster]
gi|281360117 **Mass:** 40012 **Score:** 52 **Matches:** 1(1) **Sequences:** 1(1)
CG2082, isoform G [Drosophila melanogaster]
gi|281360119 **Mass:** 39979 **Score:** 52 **Matches:** 1(1) **Sequences:** 1(1)
CG2082, isoform H [Drosophila melanogaster]
gi|327180770 **Mass:** 41129 **Score:** 52 **Matches:** 1(1) **Sequences:** 1(1)
RH63159p [Drosophila melanogaster]
gi|383292515 **Mass:** 36950 **Score:** 52 **Matches:** 1(1) **Sequences:** 1(1)
CG2082, isoform O [Drosophila melanogaster]

---

**5.** gi|198476294 **Mass:** 41611 **Score:** 49 **Matches:** 1(1) **Sequences:** 1(1) **emPAI:** 0.08
GA25398 [Drosophila pseudoobscura pseudoobscura]

| Query | Observed | Mr(expt) | Mr(calc) | Delta | Miss | Score | Expect | Rank | Unique | Peptide |
|---|---|---|---|---|---|---|---|---|---|---|
| 14689 | 729.6755 | 1457.3365 | 1456.7885 | 0.5480 | 1 | 49 | 0.025 | 1 | U | R.ETQIKNNIEQIK.D |

---

**6.** gi|24640588 **Mass:** 55121 **Score:** 49 **Matches:** 1(1) **Sequences:** 1(1) **emPAI:** 0.06
CG1440, isoform A [Drosophila melanogaster]

| Query | Observed | Mr(expt) | Mr(calc) | Delta | Miss | Score | Expect | Rank | Unique | Peptide |
|---|---|---|---|---|---|---|---|---|---|---|
| 15015 | 746.5173 | 2236.5302 | 2236.8925 | -0.3624 | 0 | 49 | 0.026 | 1 | U | M.SDNNSGSGGGGSGGGGAGNNNSANSGK.N |

---

**7.** gi|24651407 **Mass:** 63075 **Score:** 47 **Matches:** 2(2) **Sequences:** 2(2) **emPAI:** 0.11
prolyl-4-hydroxylase-alpha EFB [Drosophila melanogaster]

| Query | Observed | Mr(expt) | Mr(calc) | Delta | Miss | Score | Expect | Rank | Unique | Peptide |
|---|---|---|---|---|---|---|---|---|---|---|
| 10488 | 541.1962 | 1080.3779 | 1079.4077 | 0.9702 | 0 | 45 | 0.045 | 1 | U | K.GDDGTDEMPK.S |
| 15695 | 844.5203 | 1687.0260 | 1686.7308 | 0.2952 | 0 | 47 | 0.045 | 1 | U | R.RPCDLEEDHGEFAI.- |

**Proteins matching the same set of peptides:**
gi|195505190 **Mass:** 55776 **Score:** 47 **Matches:** 2(2) **Sequences:** 2(2)
GE10881 [Drosophila yakuba]

---

**8.** gi|195161605 **Mass:** 46663 **Score:** 47 **Matches:** 1(1) **Sequences:** 1(1) **emPAI:** 0.07
GL26392 [Drosophila persimilis]

| Query | Observed | Mr(expt) | Mr(calc) | Delta | Miss | Score | Expect | Rank | Unique | Peptide |
|---|---|---|---|---|---|---|---|---|---|---|
| 14284 | 712.2616 | 1422.5086 | 1422.7428 | -0.2342 | 1 | 47 | 0.042 | 1 | U | K.LIESMEKFLEGK.L |

**Proteins matching the same set of peptides:**
gi|198472755 **Mass:** 46598 **Score:** 47 **Matches:** 1(1) **Sequences:** 1(1)
GA14329 [Drosophila pseudoobscura pseudoobscura]

---

**9.** gi|195450712 **Mass:** 342814 **Score:** 45 **Matches:** 1(0) **Sequences:** 1(0) **emPAI:** 0.01
GK13603 [Drosophila willistoni]

| Query | Observed | Mr(expt) | Mr(calc) | Delta | Miss | Score | Expect | Rank | Unique | Peptide |
|---|---|---|---|---|---|---|---|---|---|---|
| 15099 | 751.6011 | 1501.1876 | 1501.7922 | -0.6046 | 1 | 45 | 0.057 | 1 | U | K.SCELLDIQQRIK.S |

---

**Mascot:** http://www.matrixscience.com/

### {MATRIX SCIENCE} Mascot Search Results

D17

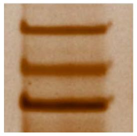

C

| | | |
|---|---|---|
| User | : | |
| Email | : | |
| Search title | : | |
| MS data file | : | ZIEN-29Nov2017_flyC.mgf |
| Database | : | NCBInr 20120419 (17893860 sequences; 6141683785 residues) |
| Taxonomy | : | Drosophila (fruit flies) (222527 sequences) |
| Timestamp | : | 30 Nov 2017 at 19:12:36 GMT |
| Enzyme | : | Trypsin |
| Variable modifications | : | Carbamidomethyl (C),Oxidation (M) |
| Mass values | : | Monoisotopic |
| Protein Mass | : | Unrestricted |
| Peptide Mass Tolerance | : | ± 1.25 Da |
| Fragment Mass Tolerance | : | ± 1.001 Da |
| Max Missed Cleavages | : | 1 |
| Instrument type | : | Default |
| Number of queries | : | 16295 |
| Protein hits | : | gi\|195590397 GD12572 [Drosophila simulans] |
| | | gi\|17981717 catalase [Drosophila melanogaster] |
| | | gi\|195022939 GH14374 [Drosophila grimshawi] |
| | | gi\|8444 unnamed protein product [Drosophila melanogaster] |
| | | gi\|287945 ATP synthase beta subunit [Drosophila melanogaster] |
| | | gi\|19921848 CG8258 [Drosophila melanogaster] |
| | | gi\|24655737 beta-Tubulin at 56D, isoform B [Drosophila melanogaster] |
| | | gi\|24651407 prolyl-4-hydroxylase-alpha EFB [Drosophila melanogaster] |
| | | gi\|195385695 GJ11603 [Drosophila virilis] |
| | | gi\|195438453 GK24169 [Drosophila willistoni] |
| | | gi\|3108349 pyruvate kinase [Drosophila melanogaster] |
| | | gi\|195434210 GK19056 [Drosophila willistoni] |
| | | gi\|21356731 CG5112 [Drosophila melanogaster] |
| | | gi\|195395688 GJ10965 [Drosophila virilis] |
| | | gi\|195434441 GK14789 [Drosophila willistoni] |
| | | gi\|194763631 GF20995 [Drosophila ananassae] |
| | | gi\|194901836 GG18651 [Drosophila erecta] |
| | | gi\|24648478 KaiRIA [Drosophila melanogaster] |
| | | gi\|195023900 GH20904 [Drosophila grimshawi] |
| | | gi\|195488195 GE14061 [Drosophila yakuba] |
| | | gi\|195126431 GI13073 [Drosophila mojavensis] |
| | | gi\|16554779 no on or off transient A [Drosophila virilis] |

#### Select Summary Report

Format As | Select Summary (protein hits) ▼ | Help

Significance threshold p< 0.05 Max. number of hits AUTO

Standard scoring ○ MudPIT scoring ◉ Ions score or expect cut-off 44 Show sub-sets 0

Show pop-ups ◉ Suppress pop-ups ○ Require bold red ☐

Re-Search ◉ All queries ○ Unassigned ○ Below homology threshold ○ Below identity threshold

1. gi|195590397 **Mass:** 55745 **Score:** 238 **Matches:** 12(12) **Sequences:** 8(8) **emPAI:** 0.58

GD12572 [Drosophila simulans]

| Query | Observed | Mr(expt) | Mr(calc) | Delta | Miss | Score | Expect | Rank | Unique | Peptide |
|---|---|---|---|---|---|---|---|---|---|---|
| 12656 | 608.6249 | 1215.2352 | 1214.6659 | 0.5693 | 0 | 58 | 0.0033 | 1 | U | R.QAADIIAWVTK.K 12647 |
| 13773 | 669.4131 | 1336.8116 | 1336.6180 | 0.1936 | 0 | 68 | 0.00042 | 1 | U | K.MDSTANELESIK.I |
| 14382 | 694.6771 | 1387.3397 | 1387.7347 | -0.3950 | 0 | 56 | 0.0062 | 1 | U | K.QLAPIYDQLAEK.Y 14377 |
| 14981 | 719.0861 | 1436.1577 | 1435.7195 | 0.4382 | 0 | 46 | 0.046 | 1 | U | K.DLTSVADAEQFLK.D |
| 15648 | 754.4683 | 1506.9221 | 1506.7930 | 0.1292 | 0 | 57 | 0.0027 | 1 | U | K.VLVSSNFESVALDK.S |
| 15743 | 760.4873 | 1518.9600 | 1518.7930 | 0.1671 | 0 | 54 | 0.0087 | 1 | U | K.VEEGVLVATVDNFK.Q 15736 15740 |
| 16281 | 1115.7028 | 2229.3910 | 2229.1430 | 0.2481 | 0 | 72 | 8.8e-005 | 1 | U | K.FAQVQSLPLIVDFNHESASK.I |
| 16021 | 787.0250 | 2358.0532 | 2357.2379 | 0.8153 | 1 | 58 | 0.0036 | 1 | U | K.KFAQVQSLPLIVDFNHESASK.I |

**Proteins matching the same set of peptides:**

gi|17647799 **Mass:** 55746 **Score:** 238 **Matches:** 12(12) **Sequences:** 8(8)

protein disulfide isomerase, isoform A [Drosophila melanogaster]

gi|194751557 **Mass:** 55553 **Score:** 238 **Matches:** 12(12) **Sequences:** 8(8)

GF10739 [Drosophila ananassae]

gi|194871212 **Mass:** 55744 **Score:** 238 **Matches:** 12(12) **Sequences:** 8(8)

GG13681 [Drosophila erecta]

gi|195327719 **Mass:** 55772 **Score:** 238 **Matches:** 12(12) **Sequences:** 8(8)

GM24501 [Drosophila sechellia]

---

2. gi|17981717 **Mass:** 57113 **Score:** 88 **Matches:** 3(3) **Sequences:** 3(3) **emPAI:** 0.18

catalase [Drosophila melanogaster]

| Query | Observed | Mr(expt) | Mr(calc) | Delta | Miss | Score | Expect | Rank | Unique | Peptide |
|---|---|---|---|---|---|---|---|---|---|---|
| 12231 | 589.4701 | 1176.9256 | 1176.6060 | 0.3196 | 0 | 46 | 0.048 | 1 | U | R.MLTEELNLAK.S |
| 12891 | 619.3951 | 1236.7757 | 1236.5986 | 0.1771 | 0 | 45 | 0.051 | 1 | U | R.DAASNQLIDYK.N |
| 15036 | 721.9436 | 1441.8726 | 1440.6481 | 1.2245 | 0 | 83 | 6.4e-006 | 1 | | R.FSTVGGESGSADTAR.D |

**Proteins matching the same set of peptides:**

gi|194871234 **Mass:** 57183 **Score:** 88 **Matches:** 3(3) **Sequences:** 3(3)

GG13679 [Drosophila erecta]

gi|195352234 **Mass:** 57134 **Score:** 88 **Matches:** 3(3) **Sequences:** 3(3)
Cat [Drosophila sechellia]

gi|195494497 **Mass:** 57064 **Score:** 88 **Matches:** 3(3) **Sequences:** 3(3)
GE19975 [Drosophila yakuba]

gi|195591356 **Mass:** 57134 **Score:** 88 **Matches:** 3(3) **Sequences:** 3(3)
catalase [Drosophila simulans]

---

**3.** gi|195022939 **Mass:** 57059 **Score:** 86 **Matches:** 2(2) **Sequences:** 2(2) **emPAI:** 0.12
GH14374 [Drosophila grimshawi]

| Query | Observed | Mr(expt) | Mr(calc) | Delta | Miss | Score | Expect | Rank | Unique | Peptide |
|---|---|---|---|---|---|---|---|---|---|---|
| 12891 | 619.3951 | 1236.7757 | 1236.5986 | 0.1771 | 0 | 45 | 0.051 | 1 | U | R.DAASNQLLDYK.N |
| 15036 | 721.9436 | 1441.8726 | 1440.6481 | 1.2245 | 0 | 83 | 6.4e-006 | 1 | | R.FSTVGGESGSADTAR.D |

**Proteins matching the same set of peptides:**

gi|195378516 **Mass:** 57117 **Score:** 86 **Matches:** 2(2) **Sequences:** 2(2)
GJ11574 [Drosophila virilis]

---

**4.** gi|8444 **Mass:** 62291 **Score:** 84 **Matches:** 2(2) **Sequences:** 2(2) **emPAI:** 0.11
unnamed protein product [Drosophila melanogaster]

| Query | Observed | Mr(expt) | Mr(calc) | Delta | Miss | Score | Expect | Rank | Unique | Peptide |
|---|---|---|---|---|---|---|---|---|---|---|
| 12806 | **615.2691** | **1228.5236** | **1228.6775** | **-0.1539** | **0** | **64** | **0.0013** | **1** | **U** | **K.SNILVATDVAAR.G** |
| 13278 | **640.6392** | **1279.2638** | **1278.7296** | **0.5342** | **0** | **62** | **0.0014** | **1** | **U** | **R.GDGPIALVLAPTR.E** |

**Proteins matching the same set of peptides:**

gi|24644479 **Mass:** 78500 **Score:** 84 **Matches:** 2(2) **Sequences:** 2(2)
Rm62, isoform A [Drosophila melanogaster]

gi|24644481 **Mass:** 62436 **Score:** 84 **Matches:** 2(2) **Sequences:** 2(2)
Rm62, isoform D [Drosophila melanogaster]

gi|24644483 **Mass:** 62791 **Score:** 84 **Matches:** 2(2) **Sequences:** 2(2)
Rm62, isoform E [Drosophila melanogaster]

gi|24644485 **Mass:** 62829 **Score:** 84 **Matches:** 2(2) **Sequences:** 2(2)
Rm62, isoform C [Drosophila melanogaster]

gi|194745414 **Mass:** 77004 **Score:** 84 **Matches:** 2(2) **Sequences:** 2(2)
GF18634 [Drosophila ananassae]

gi|194898941 **Mass:** 78362 **Score:** 84 **Matches:** 2(2) **Sequences:** 2(2)
GG10666 [Drosophila erecta]

gi|195144578 **Mass:** 71625 **Score:** 84 **Matches:** 2(2) **Sequences:** 2(2)
GL23489 [Drosophila persimilis]

gi|195343855 **Mass:** 78805 **Score:** 84 **Matches:** 2(2) **Sequences:** 2(2)
GM10578 [Drosophila sechellia]
gi|195453112 **Mass:** 81548 **Score:** 84 **Matches:** 2(2) **Sequences:** 2(2)
GK14214 [Drosophila willistoni]
gi|195502160 **Mass:** 78418 **Score:** 84 **Matches:** 2(2) **Sequences:** 2(2)
GE24123 [Drosophila yakuba]
gi|198452778 **Mass:** 79875 **Score:** 84 **Matches:** 2(2) **Sequences:** 2(2)
GA10214 [Drosophila pseudoobscura pseudoobscura]
gi|383292518 **Mass:** 50067 **Score:** 84 **Matches:** 2(2) **Sequences:** 2(2)
Rm62, isoform J [Drosophila melanogaster]
gi|383292519 **Mass:** 53163 **Score:** 84 **Matches:** 2(2) **Sequences:** 2(2)
Rm62, isoform K [Drosophila melanogaster]
gi|383292520 **Mass:** 51999 **Score:** 84 **Matches:** 2(2) **Sequences:** 2(2)
Rm62, isoform L [Drosophila melanogaster]
gi|383292521 **Mass:** 52514 **Score:** 84 **Matches:** 2(2) **Sequences:** 2(2)
Rm62, isoform M [Drosophila melanogaster]
gi|383873392 **Mass:** 33568 **Score:** 84 **Matches:** 2(2) **Sequences:** 2(2)
MIP33508p1 [Drosophila melanogaster]
gi|195109284 **Mass:** 79484 **Score:** 84 **Matches:** 2(2) **Sequences:** 2(2)
GI23184 [Drosophila mojavensis]

---

**5.** gi|287945 **Mass:** 53487 **Score:** 63 **Matches:** 2(2) **Sequences:** 2(2) **emPAI:** 0.13
ATP synthase beta subunit [Drosophila melanogaster]

| Query | Observed | Mr(expt) | Mr(calc) | Delta | Miss | Score | Expect | Rank | Unique | Peptide |
|---|---|---|---|---|---|---|---|---|---|---|
| 10435 | **517.1084** | **1032.2022** | **1032.5604** | **-0.3582** | **0** | **46** | **0.035** | **1** | **U** | **K.VLDTGYPIR.I** |
| 14074 | **684.3671** | **1366.7196** | **1366.7456** | **-0.0260** | **0** | **61** | **0.0019** | **1** | **U** | **R.IINVIGEPIDER.G** |

**Proteins matching the same set of peptides:**
gi|24638766 **Mass:** 54074 **Score:** 63 **Matches:** 2(2) **Sequences:** 2(2)
ATP synthase-beta, isoform A [Drosophila melanogaster]
gi|38048209 **Mass:** 21234 **Score:** 63 **Matches:** 2(2) **Sequences:** 2(2)
similar to Drosophila melanogaster ATPsyn-beta, partial [Drosophila yakuba]
gi|38048553 **Mass:** 21774 **Score:** 63 **Matches:** 2(2) **Sequences:** 2(2)
similar to Drosophila melanogaster ATPsyn-beta, partial [Drosophila yakuba]
gi|194770714 **Mass:** 54254 **Score:** 63 **Matches:** 2(2) **Sequences:** 2(2)
GF21882 [Drosophila ananassae]
gi|195064082 **Mass:** 54196 **Score:** 63 **Matches:** 2(2) **Sequences:** 2(2)
GH23965 [Drosophila grimshawi]

gi|195134016 **Mass:** 54192 **Score:** 63 **Matches:** 2(2) **Sequences:** 2(2)
GI14045 [Drosophila mojavensis]
gi|195355680 **Mass:** 41109 **Score:** 63 **Matches:** 2(2) **Sequences:** 2(2)
GM13022 [Drosophila sechellia]
gi|195450700 **Mass:** 54274 **Score:** 63 **Matches:** 2(2) **Sequences:** 2(2)
GK13685 [Drosophila willistoni]
gi|195469417 **Mass:** 54104 **Score:** 63 **Matches:** 2(2) **Sequences:** 2(2)
ATPsyn-beta [Drosophila yakuba]
gi|195564346 **Mass:** 47247 **Score:** 63 **Matches:** 2(2) **Sequences:** 2(2)
ATPsyn-beta [Drosophila simulans]

---

**6.** gi|19921848 **Mass:** 59396 **Score:** 62 **Matches:** 2(2) **Sequences:** 2(2) **emPAI:** 0.11
CG8258 [Drosophila melanogaster]

| Query | Observed | Mr(expt) | Mr(calc) | Delta | Miss | Score | Expect | Rank | Unique | Peptide |
|---|---|---|---|---|---|---|---|---|---|---|
| 11419 | 560.9957 | 1119.9768 | 1119.5560 | 0.4208 | 0 | 46 | 0.056 | 1 | U | R.ALDDAINNFK.C |
| 15559 | 749.0649 | 1496.1153 | 1495.7406 | 0.3747 | 0 | 57 | 0.0039 | 1 | U | R.LGITTAEISDGYEK.A |

---

**7.** gi|24655737 **Mass:** 50115 **Score:** 62 **Matches:** 2(2) **Sequences:** 2(2) **emPAI:** 0.14
beta-Tubulin at 56D, isoform B [Drosophila melanogaster]

| Query | Observed | Mr(expt) | Mr(calc) | Delta | Miss | Score | Expect | Rank | Unique | Peptide |
|---|---|---|---|---|---|---|---|---|---|---|
| 13754 | 668.3984 | 1334.7823 | 1334.6904 | 0.0919 | 0 | 53 | 0.012 | 1 | U | R.IMNTYSVVPSPK.V |
| 16093 | 801.2383 | 1600.4620 | 1600.8131 | -0.3511 | 0 | 50 | 0.017 | 1 | U | R.AVLVDLEPGTMDSVR.S |

**Proteins matching the same set of peptides:**
gi|24655741 **Mass:** 51264 **Score:** 62 **Matches:** 2(2) **Sequences:** 2(2)
beta-Tubulin at 56D, isoform A [Drosophila melanogaster]
gi|188529361 **Mass:** 20477 **Score:** 62 **Matches:** 2(2) **Sequences:** 2(2)
beta-tubulin [Drosophila silvestris]
gi|194881355 **Mass:** 53339 **Score:** 62 **Matches:** 2(2) **Sequences:** 2(2)
GG20906 [Drosophila erecta]
gi|306922435 **Mass:** 51322 **Score:** 62 **Matches:** 2(2) **Sequences:** 2(2)
GH12877p [Drosophila melanogaster]

---

**8.** gi|24651407 **Mass:** 63075 **Score:** 60 **Matches:** 1(1) **Sequences:** 1(1) **emPAI:** 0.05
prolyl-4-hydroxylase-alpha EFB [Drosophila melanogaster]

| Query | Observed | Mr(expt) | Mr(calc) | Delta | Miss | Score | Expect | Rank | Unique | Peptide |
|---|---|---|---|---|---|---|---|---|---|---|
| 16080 | 799.1600 | 1596.3055 | 1595.7791 | 0.5264 | 0 | 60 | 0.0017 | 1 | U | R.LQDTYQLDTSSVAR.G |

**Proteins matching the same set of peptides:**

gi|125772807 **Mass:** 62641 **Score:** 60 **Matches:** 1(1) **Sequences:** 1(1)
GA15946 [Drosophila pseudoobscura pseudoobscura]

gi|194765194 **Mass:** 62567 **Score:** 60 **Matches:** 1(1) **Sequences:** 1(1)
GF22904 [Drosophila ananassae]

gi|194905436 **Mass:** 62737 **Score:** 60 **Matches:** 1(1) **Sequences:** 1(1)
GG11753 [Drosophila erecta]

gi|195055779 **Mass:** 55833 **Score:** 60 **Matches:** 1(1) **Sequences:** 1(1)
GH14110 [Drosophila grimshawi]

gi|195159323 **Mass:** 55783 **Score:** 60 **Matches:** 1(1) **Sequences:** 1(1)
GL13463 [Drosophila persimilis]

gi|195341536 **Mass:** 63009 **Score:** 60 **Matches:** 1(1) **Sequences:** 1(1)
GM12882 [Drosophila sechellia]

gi|195391754 **Mass:** 55603 **Score:** 60 **Matches:** 1(1) **Sequences:** 1(1)
GJ24502 [Drosophila virilis]

gi|195452726 **Mass:** 62836 **Score:** 60 **Matches:** 1(1) **Sequences:** 1(1)
GK14136 [Drosophila willistoni]

gi|195505190 **Mass:** 55776 **Score:** 60 **Matches:** 1(1) **Sequences:** 1(1)
GE10881 [Drosophila yakuba]

gi|195575089 **Mass:** 62991 **Score:** 60 **Matches:** 1(1) **Sequences:** 1(1)
GD21521 [Drosophila simulans]

---

**9.** gi|195385695 **Mass:** 47177 **Score:** 60 **Matches:** 2(2) **Sequences:** 1(1) **emPAI:** 0.07
GJ11603 [Drosophila virilis]

| Query | Observed | Mr(expt) | Mr(calc) | Delta | Miss | Score | Expect | Rank | Unique | Peptide |
|---|---|---|---|---|---|---|---|---|---|---|
| 13981 | 680.0380 | 1358.0614 | 1358.7228 | -0.6614 | 1 | 54 | 0.0044 | 1 | U | -.MSKQLADLQPTK.F 13982 |

---

**10.** gi|195438453 **Mass:** 56787 **Score:** 52 **Matches:** 1(1) **Sequences:** 1(1) **emPAI:** 0.06
GK24169 [Drosophila willistoni]

| Query | Observed | Mr(expt) | Mr(calc) | Delta | Miss | Score | Expect | Rank | Unique | Peptide |
|---|---|---|---|---|---|---|---|---|---|---|
| 12945 | 622.3690 | 1242.7234 | 1241.6364 | 1.0870 | 0 | 52 | 0.018 | 1 | U | K.NQDVAAVQEIR.E |

**Proteins matching the same set of peptides:**

gi|17137450 **Mass:** 55970 **Score:** 52 **Matches:** 1(1) **Sequences:** 1(1)
regulatory particle non-ATPase 3 [Drosophila melanogaster]

gi|19527921 **Mass:** 55992 **Score:** 52 **Matches:** 1(1) **Sequences:** 1(1)
AT15146p [Drosophila melanogaster]

gi|125986955 **Mass:** 56336 **Score:** 52 **Matches:** 1(1) **Sequences:** 1(1)
GA10344 [Drosophila pseudoobscura pseudoobscura]

gi|170280395 **Mass:** 26416 **Score:** 52 **Matches:** 1(1) **Sequences:** 1(1)
diphenol oxidase A2, partial [Drosophila erecta]
gi|170280397 **Mass:** 26437 **Score:** 52 **Matches:** 1(1) **Sequences:** 1(1)
diphenol oxidase A2, partial [Drosophila mauritiana]
gi|170280399 **Mass:** 26398 **Score:** 52 **Matches:** 1(1) **Sequences:** 1(1)
diphenol oxidase A2, partial [Drosophila melanogaster]
gi|170280401 **Mass:** 26430 **Score:** 52 **Matches:** 1(1) **Sequences:** 1(1)
diphenol oxidase A2, partial [Drosophila santomea]
gi|170280403 **Mass:** 26509 **Score:** 52 **Matches:** 1(1) **Sequences:** 1(1)
diphenol oxidase A2, partial [Drosophila sechellia]
gi|170280405 **Mass:** 26414 **Score:** 52 **Matches:** 1(1) **Sequences:** 1(1)
diphenol oxidase A2, partial [Drosophila simulans]
gi|170280407 **Mass:** 26458 **Score:** 52 **Matches:** 1(1) **Sequences:** 1(1)
diphenol oxidase A2, partial [Drosophila teissieri]
gi|194879789 **Mass:** 56011 **Score:** 52 **Matches:** 1(1) **Sequences:** 1(1)
GG21155 [Drosophila erecta]
gi|195156337 **Mass:** 56318 **Score:** 52 **Matches:** 1(1) **Sequences:** 1(1)
GL26159 [Drosophila persimilis]
gi|195345003 **Mass:** 56051 **Score:** 52 **Matches:** 1(1) **Sequences:** 1(1)
GM17319 [Drosophila sechellia]
gi|195484349 **Mass:** 56025 **Score:** 52 **Matches:** 1(1) **Sequences:** 1(1)
GE13227 [Drosophila yakuba]
gi|195580065 **Mass:** 56028 **Score:** 52 **Matches:** 1(1) **Sequences:** 1(1)
GD24178 [Drosophila simulans]

---

**11.** gi|3108349 **Mass:** 57437 **Score:** 50 **Matches:** 1(1) **Sequences:** 1(1) **emPAI:** 0.06
pyruvate kinase [Drosophila melanogaster]

| Query | Observed | Mr(expt) | Mr(calc) | Delta | Miss | Score | Expect | Rank | Unique | Peptide |
|---|---|---|---|---|---|---|---|---|---|---|
| 11674 | **571.3002** | **1140.5859** | **1140.6026** | **-0.0167** | **0** | **50** | **0.031** | **1** | **U** | **R.GDLGIEIPAEK.V** |

**Proteins matching the same set of peptides:**
gi|24648964 **Mass:** 55024 **Score:** 50 **Matches:** 1(1) **Sequences:** 1(1)
pyruvate kinase, isoform B [Drosophila melanogaster]
gi|27819773 **Mass:** 57438 **Score:** 50 **Matches:** 1(1) **Sequences:** 1(1)
RH07636p [Drosophila melanogaster]
gi|28571814 **Mass:** 57404 **Score:** 50 **Matches:** 1(1) **Sequences:** 1(1)
pyruvate kinase, isoform A [Drosophila melanogaster]
gi|38047755 **Mass:** 19755 **Score:** 50 **Matches:** 1(1) **Sequences:** 1(1)

similar to Drosophila melanogaster PyK, partial [Drosophila yakuba]
gi|194744590 **Mass:** 57203 **Score:** 50 **Matches:** 1(1) **Sequences:** 1(1)
GF18439 [Drosophila ananassae]
gi|194911138 **Mass:** 57502 **Score:** 50 **Matches:** 1(1) **Sequences:** 1(1)
GG11123 [Drosophila erecta]
gi|195053328 **Mass:** 57522 **Score:** 50 **Matches:** 1(1) **Sequences:** 1(1)
GH20329 [Drosophila grimshawi]
gi|195056081 **Mass:** 57549 **Score:** 50 **Matches:** 1(1) **Sequences:** 1(1)
GH13278 [Drosophila grimshawi]
gi|195112292 **Mass:** 57581 **Score:** 50 **Matches:** 1(1) **Sequences:** 1(1)
GI22374 [Drosophila mojavensis]
gi|195145138 **Mass:** 57003 **Score:** 50 **Matches:** 1(1) **Sequences:** 1(1)
GL24201 [Drosophila persimilis]
gi|195330979 **Mass:** 57418 **Score:** 50 **Matches:** 1(1) **Sequences:** 1(1)
GM26420 [Drosophila sechellia]
gi|195391442 **Mass:** 102583 **Score:** 50 **Matches:** 1(1) **Sequences:** 1(1)
GJ24408 [Drosophila virilis]
gi|195453308 **Mass:** 57517 **Score:** 50 **Matches:** 1(1) **Sequences:** 1(1)
GK14262 [Drosophila willistoni]
gi|195502545 **Mass:** 57418 **Score:** 50 **Matches:** 1(1) **Sequences:** 1(1)
PyK [Drosophila yakuba]
gi|198452351 **Mass:** 57033 **Score:** 50 **Matches:** 1(1) **Sequences:** 1(1)
GA26534 [Drosophila pseudoobscura pseudoobscura]

---

**12.** gi|195434210 **Mass:** 21374 **Score:** 49 **Matches:** 1(1) **Sequences:** 1(1) **emPAI:** 0.16
GK19056 [Drosophila willistoni]

| Query | Observed | Mr(expt) | Mr(calc) | Delta | Miss | Score | Expect | Rank | Unique | Peptide |
|---|---|---|---|---|---|---|---|---|---|---|
| 12267 | 590.4731 | 1178.9317 | 1178.5866 | 0.3451 | 0 | 49 | 0.013 | 1 | U | K.YQLIQMNNR.S |

---

**13.** gi|21356731 **Mass:** 58448 **Score:** 49 **Matches:** 1(1) **Sequences:** 1(1) **emPAI:** 0.06
CG5112 [Drosophila melanogaster]

| Query | Observed | Mr(expt) | Mr(calc) | Delta | Miss | Score | Expect | Rank | Unique | Peptide |
|---|---|---|---|---|---|---|---|---|---|---|
| 11365 | 558.2457 | 1114.4769 | 1113.6142 | 0.8627 | 0 | 49 | 0.022 | 1 | U | K.SAVELAQQIR.E |

**Proteins matching the same set of peptides:**
gi|125776626 **Mass:** 58393 **Score:** 49 **Matches:** 1(1) **Sequences:** 1(1)
GA18668 [Drosophila pseudoobscura pseudoobscura]
gi|194908530 **Mass:** 58327 **Score:** 49 **Matches:** 1(1) **Sequences:** 1(1)

GG11409 [Drosophila erecta]

gi|195152431 **Mass:** 58361 **Score:** 49 **Matches:** 1(1) **Sequences:** 1(1)
GL22143 [Drosophila persimilis]

gi|195349457 **Mass:** 58431 **Score:** 49 **Matches:** 1(1) **Sequences:** 1(1)
GM10248 [Drosophila sechellia]

gi|195453896 **Mass:** 58368 **Score:** 49 **Matches:** 1(1) **Sequences:** 1(1)
GK12847 [Drosophila willistoni]

gi|195504320 **Mass:** 58393 **Score:** 49 **Matches:** 1(1) **Sequences:** 1(1)
GE23605 [Drosophila yakuba]

gi|195573909 **Mass:** 58441 **Score:** 49 **Matches:** 1(1) **Sequences:** 1(1)
GD21221 [Drosophila simulans]

---

**14.** gi|195395688 **Mass:** 58173 **Score:** 49 **Matches:** 1(1) **Sequences:** 1(1) **emPAI:** 0.06
GJ10965 [Drosophila virilis]

| Query | Observed | Mr(expt) | Mr(calc) | Delta | Miss | Score | Expect | Rank | Unique | Peptide |
|---|---|---|---|---|---|---|---|---|---|---|
| 11365 | 558.2457 | 1114.4769 | 1113.6506 | 0.8263 | 1 | 49 | 0.022 | 1 | U | K.SAVELAKQIR.E |

---

**15.** gi|195434441 **Mass:** 51967 **Score:** 48 **Matches:** 1(1) **Sequences:** 1(1) **emPAI:** 0.06
GK14789 [Drosophila willistoni]

| Query | Observed | Mr(expt) | Mr(calc) | Delta | Miss | Score | Expect | Rank | Unique | Peptide |
|---|---|---|---|---|---|---|---|---|---|---|
| 14049 | **683.3903** | **1364.7660** | **1364.6295** | **0.1364** | **0** | **48** | **0.041** | **1** | **U** | **K.MDSHFNAFQLR.N** |

---

**16.** gi|194763631 **Score:** 48 **Matches:** 1(1) **Sequences:** 1(1) **emPAI:** 0.03
GF20995 [Drosophila ananassae]

| Query | Observed | Mr(expt) | Mr(calc) | Delta | Miss | Score | Expect | Rank | Unique | Peptide |
|---|---|---|---|---|---|---|---|---|---|---|
| 13278 | 640.6392 | 1279.2638 | 1278.7296 | 0.5342 | 0 | 48 | 0.036 | 2 | U | R.GEGPVALVLAPTR.E |

**Proteins matching the same set of peptides:**
gi|195438679 **Score:** 48 **Matches:** 1(1) **Sequences:** 1(1)

---

**17.** gi|194901836 **Mass:** 5994 **Score:** 47 **Matches:** 1(1) **Sequences:** 1(1) **emPAI:** 0.60
GG18651 [Drosophila erecta]

| Query | Observed | Mr(expt) | Mr(calc) | Delta | Miss | Score | Expect | Rank | Unique | Peptide |
|---|---|---|---|---|---|---|---|---|---|---|
| 13236 | **638.0350** | **1274.0554** | **1273.5760** | **0.4794** | **1** | **47** | **0.046** | **1** | **U** | **K.KGGGGGGGGGGGGSGNK.N** |

---

**18.** gi|24648478 **Mass:** 101593 **Score:** 46 **Matches:** 1(1) **Sequences:** 1(1) **emPAI:** 0.03
KaiRIA [Drosophila melanogaster]

| Query | Observed | Mr(expt) | Mr(calc) | Delta | Miss | Score | Expect | Rank | Unique | Peptide |
|---|---|---|---|---|---|---|---|---|---|---|
| 13749 | 668.1045 | 1334.1944 | 1334.6790 | -0.4845 | 1 | 46 | 0.043 | 1 | U | R.SRNSSQSIESLK.T |

**Proteins matching the same set of peptides:**

gi|61675663 **Mass:** 101564 **Score:** 46 **Matches:** 1(1) **Sequences:** 1(1)
RE24732p [Drosophila melanogaster]

gi|194899712 **Mass:** 101589 **Score:** 46 **Matches:** 1(1) **Sequences:** 1(1)
GG15269 [Drosophila erecta]

gi|195353927 **Mass:** 86610 **Score:** 46 **Matches:** 1(1) **Sequences:** 1(1)
GM23173 [Drosophila sechellia]

gi|195498265 **Mass:** 101563 **Score:** 46 **Matches:** 1(1) **Sequences:** 1(1)
GE25051 [Drosophila yakuba]

gi|195569399 **Mass:** 101626 **Score:** 46 **Matches:** 1(1) **Sequences:** 1(1)
GD20046 [Drosophila simulans]

---

**19.** gi|195023900 **Mass:** 190030 **Score:** 46 **Matches:** 1(1) **Sequences:** 1(1) **emPAI:** 0.02
GH20904 [Drosophila grimshawi]

| Query | Observed | Mr(expt) | Mr(calc) | Delta | Miss | Score | Expect | Rank | Unique | Peptide |
|---|---|---|---|---|---|---|---|---|---|---|
| 12330 | 593.1379 | 1184.2612 | 1184.6302 | -0.3690 | 1 | 46 | 0.029 | 1 | U | K.VFHLGGGGDAKK.K |

---

**20.** gi|195488195 **Mass:** 66917 **Score:** 46 **Matches:** 1(1) **Sequences:** 1(1) **emPAI:** 0.05
GE14061 [Drosophila yakuba]

| Query | Observed | Mr(expt) | Mr(calc) | Delta | Miss | Score | Expect | Rank | Unique | Peptide |
|---|---|---|---|---|---|---|---|---|---|---|
| 15913 | 775.3869 | 1548.7593 | 1549.7554 | -0.9961 | 1 | 46 | 0.039 | 1 | U | R.MEPMKDLMAVLEK.L |

---

**21.** gi|195126431 **Mass:** 66814 **Score:** 46 **Matches:** 1(1) **Sequences:** 1(1) **emPAI:** 0.05
GI13073 [Drosophila mojavensis]

| Query | Observed | Mr(expt) | Mr(calc) | Delta | Miss | Score | Expect | Rank | Unique | Peptide |
|---|---|---|---|---|---|---|---|---|---|---|
| 10435 | 517.1084 | 1032.2022 | 1032.5604 | -0.3582 | 0 | 46 | 0.035 | 1 | U | K.VIDTGYPIR.V |

**Proteins matching the same set of peptides:**

gi|195442758 **Mass:** 74458 **Score:** 46 **Matches:** 1(1) **Sequences:** 1(1)
GK24170 [Drosophila willistoni]

---

**22.** gi|16554779 **Mass:** 15812 **Score:** 44 **Matches:** 1(0) **Sequences:** 1(0) **emPAI:** 0.22
no on or off transient A [Drosophila virilis]

| Query | Observed | Mr(expt) | Mr(calc) | Delta | Miss | Score | Expect | Rank | Unique | Peptide |
|---|---|---|---|---|---|---|---|---|---|---|

| | | | | | | | | | |
|---|---|---|---|---|---|---|---|---|---|
| 15089 | 723.9612 | 2168.8617 | 2168.9154 | -0.0537 | 1 | 44 | 0.086 | 1 | U | R.ARGGGGGGGGGGGGGGGGGGGGGGGGGGGGGGGR.D |

---

**Mascot:** http://www.matrixscience.com/
